# Generative Modelling of Oncogene-carrying Extrachromosomal Circular DNA Biogenesis and Dynamics in Cells

**DOI:** 10.1101/2024.04.18.590030

**Authors:** János Haskó, Weijia Feng, Aram Arshadi, Doron Tolomeo, Chuang Sun Hembo, Trine Skov Petersen, Wei Lv, Peng Han, Yuchen Zeng, Fei Wang, Lars Bolund, Lin Lin, Birgitte Regenberg, Clelia Tiziana Storlazzi, Yonglun Luo

## Abstract

Extrachromosomal circular DNAs (ecDNA) are focal gene amplifications frequently associated with cancer development and often indicating a poor prognosis. To understand the early dynamics of oncogene-carrying ecDNAs, we previously developed CRISPR-C, a tool for precise ecDNA generation by deleting specific chromosomal regions. Here, we adapted CRISPR-C to recreate tumor ecDNAs. This method also allowed us to enhance ecDNA generation efficiency by directly delivering Cas9 protein and sgRNAs as a ribonucleoprotein complex. By using the modified CRISPR-C, we successfully generated ecDNAs carrying oncogenes (*EGFR, CDK4, MDM2, MYC, MYCN, FGFR2, ABCB1,* and *DHFR*) in various human cell types. Furthermore, we demonstrated that our method could generate chimeric ecDNAs composed of target sequences from distant intra or inter-chromosomal regions. Using these generative ecDNA cell models, we studied the oncogene ecDNA expression and stability. The *MDM2* expression was increased after CRISPR-C, while *CDK4* was decreased indicating genomic-context dependent effect. The copy number of CRISPR-C generated *CDK4* was ecDNA increased in cells after a long period of treatment with the *CDK4* inhibitor palbociclib. Unlike CDK4, the CRISPR-C generated *ABCB1* ecDNA was unstable in cells under normal growth conditions, but is stably retained when the cells were treated with colcemid, a recognized substrate for ABCB1. We thus provide valuable tools and an attractive platform for studying ecDNA biogenesisy and in vitro drug screening on ecDNA stability.

## INTRODUCTION

Increased number of oncogenes in somatic tissues is a hallmark of cancer. Oncogenes are often found on extrachromosomal circular DNA (ecDNA). EcDNA is defined as large, amplified, circular DNA particles that originate from one or more chromosomes (1,2). Unlike other extrachromosomal circular DNA of small size (eccDNA), which are commonly found throughout somatic tissues (3,4), ecDNA is restricted to tumors and typically carries one or several oncogenes in high copy numbers (1,5–7). Presumably, ecDNA plays a significant role in inducing genomic rearrangements in cancer cells. This can occur through the pseudo-stable circularization of DNA fragments, leading to excretion, reinsertion, inversion, and loss (1,7). Alternatively, they remain stable as ecDNAs for extended periods or over multiple cell divisions (2).

Extensive studies on clinical cancer samples and analysis of cancer-genome databases, such as The Cancer Genome Atlas (TCGA) and Pan-Cancer Analysis of Whole Genomes (PCAWG), have reported a high prevalence of ecDNAs across a wide range of cancer types, and shown that ecDNAs are often associated with poor prognoses (8,9). Many genomic regions show recurrent amplification on circular DNA, several of which encode one or more of the 24 most commonly amplified oncogenes (1,9). Examples of these are *MYC* (*10*), *MYCN* (*11*), and cell cycle-regulating kinase genes like *CDK4* (9), which significantly enhance cell proliferation. Drug resistance is associated with circular amplicons carrying *ABCB1* (12) and *DHFR*(13) and could allow cancer cells to acquire resistance to chemotherapy. Apart from the increased oncogene copy number, the expression of genes associated with ecDNA is significantly altered due to epigenetic changes (14–16). This includes increased accessibility to the transcription machinery and the acquisition of spatially juxtaposed enhancer elements through enhancer hijacking (16–18). Due to all of these factors, ecDNAs are attractive targets for treating a range of tumors.

However, current studies of ecDNA are based on correlation, and cell models that can recapitulate causation between circular DNA and tumorigenesis are largely missing. Cancer cell lines with ecDNAs have been used to study their segregation (16,19), but they have several disadvantages. Firstly, they have undergone secondary mutations that may cover the effect of ecDNAs. Second, there may be epigenetic changes that allow the ecDNA to be expressed at high level in the cells. Therefore, the initial consequences of DNA circularization and the cellular changes that allow for high levels of expression from ecDNA cannot be studied in the established cancer cell lines.

To study newly formed ecDNAs, we developed a CRISPR-based gene editing technique for the *de novo* generation of circular DNA in cells, which we will refer to as CRISPR-C in this paper (20). This method involves the delivery of two specific CRISPR guide RNA molecules (sgRNAs or gRNA) and the Cas9 protein into cells using plasmid-based expression vectors, resulting in the creation of ecDNA from defined chromosomal DNA fragments (20). CRISPR-C demonstrates the capability to generate ecDNAs of various sizes, ranging from 2 Kb to 207 Kb, and even up to the size of a whole ring chromosome structure from chromosome 18. Our previous CRISPR-C method suffers from limitations including low efficiency in inducing double-strand breaks (DSBs), which lowers the efficiency of ecDNA generation, and constraints related to the precise timing of ecDNA formation due to the prolonged presence of CRISPR molecular components expressed from the plasmid vectors.

In this study, we present a more efficient version of CRISPR-C, referred to as CRISPR-C 2.0, which utilizes the delivery of two ready-to-use synthetic sgRNAs and the Cas9 protein in the form of ribonucleoproteins (RNPs) through optimized nucleofection. This new approach allows rapid, highly simultaneous induction of ecDNA generation in human cells. We successfully targeted eight distinct chromosomal regions that are frequently amplified into circular amplicons, thereby inducing the formation of oncogene-harboring ecDNAs ranging from 100 Kb to 1 Mb in various cell lines, including both commonly used immortalized cell lines, primary non-cancer cells, and cancer cell lines. Furthermore, we monitored the persistence of ecDNAs after formation, specifically focusing on those harboring *CDK4* and *MDM2* oncogenes, associated with cell growth, and the drug resistance related gene *ABCB1*. Collectively, we provide novel methods for the generating and modeling of ecDNAs *de novo*, allowing for future studies of oncogene-harboring ecDNAs that can address how they are formed and are modified to become stable in response to selection pressures.

## MATERIALS AND METHODS

### Cell culture

Human embryonic kidney 293T (HEK293T) cells, HeLaS3 cervical adenocarcinoma cells (referred to as HeLa in this manuscript) and U2OS osteosarcoma cells (generous gifts from the laboratory of Christian Kanstrup Holm, Aarhus University, Aarhus, Denmark) and primary human skin fibroblasts (annotated as KU-H132, acquired form Copenhagen University in a frame of collaboration) were maintained in Dulbeccós modified Eaglés medium (DMEM) (Gibco; Thermo Fisher Scientific, Inc., Waltham, MA, USA) supplemented with 10% fetal bovine serum (FBS) (Gibco; Thermo Fisher), 1% GlutaMAX (Gibco; Thermo Fisher Scientific) and penicillin/streptomycin (100 units penicillin and 0.1mg streptomycin/ml, Gibco; Thermo Fisher Scientific) at 37 °C under 5% CO_2_. MCF7 triple-negative breast cancer cells were grown in Roswell Park Memorial Institute (RPMI) 1640 Medium with supplements as above at 37 °C under 5% CO_2_. Glioblastoma cell lines (GSC23 and I1552 cells a gift from the laboratory of Peter C. Scacheri, Case Western Reserve University, Cleveland, Ohio, USA) were cultured in Neurobasal-A medium (Gibco; Thermo Fisher Scientific) completed with B-27 supplement (50X), minus vitamin A, 20 ng/ml epidermal growth factor (EGF) and 20 ng/ml basic fibroblast growth factor (bFGF) (Invitrogen, Thermo Fisher Scientific) as neurospheres. Human astrocytes (Gift from the laboratory of Anders Nykjær, Aarhus University, Aarhus, Denmark) were cultured on poly-l-lysine (Sigma)-coated dishes and kept in astrocyte basal medium (ScienCell, Carlsbad, CA, USA) completed with 2% fetal bovine serum and 1% ml of astrocyte growth supplement (ScienCell, Carlsbad, CA, USA) at 37 °C under 5% CO_2_.

### CRISPR-C 1.0: eccDNA generation by plasmid vectors

eccDNA generation from the eccDNA biosensor cassette using plasmid-based vectors (CRISPR-C 1.0) was performed as described previously (20). Briefly, 50,000 HEK293-TRE-ECC, U2OS-TRE-ECC, and HeLa-TRE-ECC cells were seeded on 24-well plates 18-24 h before transfection. Cells were transfected with the following mixture: 50 μL OptiMEM (Gibco, Thermo Fisher Scientific, Inc., Waltham, MA, USA), 60ng L2-gRNA-plasmid, 60ng R2-gRNA-plasmid 180 ng Cas9 plasmid with 1,5µl X-tremeGene9 transfection reagent (Roche, Mannheim, Germany), simultaneously by adding the transfection mixture gently in a drop-wise manner to the cells. The pMaxGFP plasmid (3486 bp, Lonza, Amaxa GmbH) was delivered to the cells of the transfection control group by X-tremeGene transfection reagent to evaluate transfection efficiency.

### Single guide RNA (sgRNA) design for CRISPR-C 2.0

We designed CRISPR-C sgRNAs to generate ecDNAs for eight selected oncogenes. For each gene, we evaluated the presence of predicted enhancers 700 Kb upstream and downstream of the transcription start site (TSS) using the integrated GeneCards database and other reported putative enhancers (11). For most oncogene, we designed CRISPR-C sgRNAs to generated ecDNA of three different sizes, taking the abundance of enhancers within the ecDNA into consideration. The CRISPOR online tool (http://crispor.tefor.net/) was utilized to design sgRNAs to generate *MYC-* or *EGFR-*containing ecDNAs. We used our online software tool and webserver CRISPRon(V1.0) (https://rth.dk/resources/crispr/crispron/) to design sgRNAs with high predicted efficiency for the other oncogenes. The sgRNAs with the highest predicted editing efficiency were further filtered for low off target effect by using our other novel webtool, CRISPRoff (v1.2beta) (https://rth.dk/resources/crispr/crisproff/) as is indicated in **Supplementary Table S1** along with the designed sgRNAs. The selected sgRNAs were acquired from Synthego, (Redwood City, CA, USA) containing site-specific chemical modifications to ensure the stability and the decreased activation of the cellular innate immunity.

### Generation of ecDNAs by CRISPR-C 2.0

Cells were trypsinized and centrifuged at 400g for 5 min, followed by washing in PBS and resuspension in Opti-MEM (Gibco, Thermo Fisher Scientific, Inc., Waltham, MA, USA) to the cell density of 10^4^ cells/μL. RNP complex was generated by mixing 0.6 µL of each sgRNA (3.2 µg/µL) targeting the 5’ and 3’ terminals of the selected chromosomal region for ecDNA generation with 0.6 µl Alt-R SpCas9 (10 µg/µL, IDT, Leuven, Belgium) and incubating for 10-30 min at room temperature. 20 µL of cell suspension (200,000 cells) was mixed with the RNP-containing solution and electroporated using the 4D-Nucleofector X Unit (Lonza, Basel, Switzerland) with the CM138 nucleofection program. Nucleofected cells were transferred into 3 mL cell culture medium and seeded into three wells of a 12-well cell-culture plate.

### Flow cytometry analysis of eccDNA biosensor cell lines

One, two, three and five days after induction of either CRISPR-C 1.0 with transfection of plasmid vectors or CRISPR-C 2.0 by nucleofection with RNP complexes of eccDNA biosensor cassettes, respectively in HEK293T-TRE-ECC, HeLa-TRE-ECC, U2OS-TRE-ECC cell lines, cells were harvested and washed with 500 µl PBS and resuspended in 110 µl of PBS. Flow cytometry measurements were performed using a Quanteon analyzer, and data analysis was performed using FlowJo_V10.8.1.

### Polymerase chain reaction (PCR)

Adherent cells were pelleted and resuspended in 100 μL lyses buffer (50 mM KCl, 1.5 mM MgCl_2_, 0.5% NP40 and 0,5% Tween20, 10mM Tris, pH 8.5) containing 5 μL proteinase K (18.7 mg/ml, Roche) and incubated for 30 min at 65 °C followed by the heat-inactivation of proteinase K at 95 °C 10 min. The cell lysate was stored at −20 °C until the PCR assay was performed. PCR experiments were carried out with high fidelity AccuPrime™ Pfx DNA Polymerase (Invitrogene, Thermo Fisher Scientific, Inc., Waltham, MA, USA) in Veriti™ 96-Well Fast Thermal Cycler. All oligonucleotide primers (listed in **Supplementary Table S4**) were obtained from Merck KGaA, Darmstadt, Germany.

### Droplet digital PCR (ddPCR) analysis of ecDNA circular junction site

The circular junction site of ecDNA was quantified using ddPCR analysis. Adherent cells were harvested, pelleted, and washed with PBS. DNA extraction was performed by incubating the cells in a lysis buffer containing Tris (10 mM), EDTA (1 mM), NaCl (50 mM), SDS (0.5%), and Proteinase K (194 μg/ml, recombinant, PCR grade, Roche) for 2 hours at 56 °C to achieve cell lysis and protein digestion. Subsequently, 330 μL of 6M NaCl was added per milliliter of cell lysate, followed by centrifugation (5 minutes, 13,000 rpm, 4°C). The DNA precipitate was obtained by adding ice-cold absolute ethanol and then washed with 70% ice-cold ethanol.

For ddPCR analysis, 1.5 μL of purified genomic DNA (100 ng/μL) was utilized as a template. This DNA was mixed with 12 μL of the ddPCR supermix for probes (without dUTP, Bio-Rad, #186-3026). Additionally, 0.6 μL of PrimeTime qPCR Primers with Hex-conjugated probe (IDT) for the reference site at the human *TERT* gene and 0.6 μL of PrimeTime qPCR Primers with FAM-conjugated probe (IDT) to detect the CRISPR-C *CDK4* 508 Kb ecDNA circular junction site and 0.12 μl of Hind III Fast Digest enzyme (FD0504) were added. Nuclease-free water was added to achieve a final reaction volume of 24 μL. ddPCR droplets were produced using the QX200 Droplet generator (Bio-Rad) from Bio-Rad, following the manufacturer’s guidelines. The ddPCR reaction was carried out using the following cycling protocol: initial denaturation at 95 °C for 10 minutes; 50 cycles of denaturation at 94 °C for 30 s, annealing at 60 °C for 30 s, extension at 72 °C for 2 minutes, and a final extension step at 98 °C for 10 minutes. Droplets were analyzed using a QX200 Droplet Digital PCR reader (Bio-Rad). Quantification of the target and reference DNA copies within the droplets was performed using the Quantasoft Analysis Pro (version 1.0.596, Bio-Rad). Circular junction site relative frequency was calculated as the percentage of target sequence copies relative to that of the reference sequence copy. The sequences of the various primer/probe assays employed are provided in **Supplementary Table S4**.

### Evaluation of nucleofection efficiency

Nucleofection efficiency was evaluated by delivering 1.6 µg EGFP mRNA generating a positive control group. EGFP mRNA was generated by in vitro transcription (IVT) using the MEGAscript kit (Thermo Fischer Scientific, Inc., Waltham, MA, USA) and a T7-EGFP plasmid linearized by BbsI enzyme (Thermo Fischer Scientific). Cells were cultured under the same condition as their CRISPR edited counterpart for 24 h, then incubated with 1 µg/ml Hoechst33342 (Thermo Fischer Scientific) for 2 hrs to label their nuclei. Live cells were visualized by Olympus Scan^R fluorescent microscope using a UPLSAPO 10x dry objective.

### Evaluation of CRISPR-Cas9 editing efficiency

To evaluate the cleavage efficiency of each Cas9 sgRNA, HEK293T cells were nucleofected with CRISPR/Cas9 ribonucleoprotein (RNP) complexes containing the sgRNAs (single type of sgRNA per nucleofection group) enlisted in **Supplementary Table S1**. To form RNPs, 0.6 µL of sgRNA (3.2µg/µL) was mixed with 0.6 µL Alt-R S.p. Cas9 Nuclease V3 (10 µg/µL, IDT) and 0.6 µL nuclease free water (Synthego) and incubated for 10-30 min at room temperature. HEK293T cells were harvested, washed with PBS and resuspended in 20 μL of Opti-MEM (Gibco). A total of 200,000 cells were nucleofected using the 4D-Nucleofector X Unit (nucleofection program CM138). Two days after nucleofection, cells were harvested and lysed to acquire DNA samples. Genomic regions of the sgRNA target sites of edited and unedited samples were amplified by PCR and sent for Sanger sequencing at Eurofins Genomics, DK, by following the manufacturer’s protocol of Mix2Seq Kit (Eurofins Genomics, DK). The analysis of the Sanger sequencing data and the calculation of CRISPR gene editing efficiency and assessment of the profiles of all different types of edits were performed by utilizing Snap gene viewer and the “ICE” (Inference of CRISPR Edits) webtool (https://ice.synthego.com/#/) (21).

### Genotyping of chromosomal scar and circular junction sites

Cells reaching 10^6^ cell number after the onset of CRISPR-C 2.0 were trypsinized and 250,000 cells were resuspended in 100 μL lysis buffer as described above. The target sites of each of the two CRISPR-C sgRNAs the DNA scar in place of the deleted DNA fragment and the circular junction site formed by the two connecting terminals were amplified as depicted in **Fig. 1A** and the **Supplemental Figures S3-12.** Amplified DNA products were separated by agarose gel electrophoresis and visualized by iBright CL1500 Imaging System (Invitrogen, Thermo Fisher Scientific, Inc., Waltham, MA, USA). Amplified regions of CRISPR-C unedited samples and the specific bands derived from the amplification of the deletion scar and circular junction sites were dissected and DNA products purified by NucleoSpin Gel and PCR Clean-up kit (MACHEREY-NAGEL Düren, Nordrhein-Westfalen, Germany) and sent for Sanger sequencing at Eurofins Genomics. The obtained sequences were visualized with Snap gene viewer and manually analyzed and classified.

**Figure 1:**
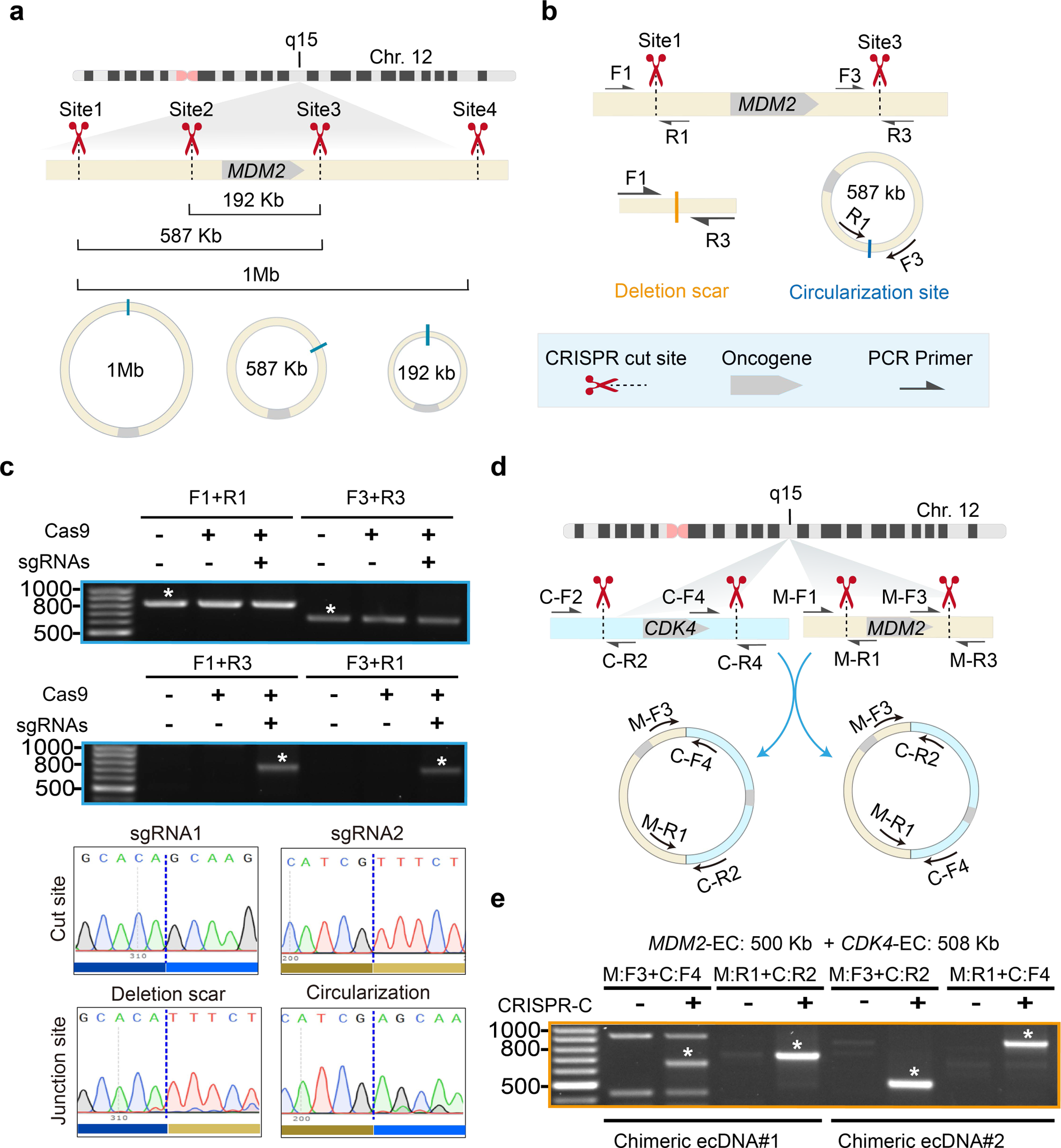
Generation oncogene-harboring large size ecDNAs with improved CRISPR-C 2.0. **(a)** Model of CRISPR-C to generate different sizes (1 Mb, 587 Kb and 192 Kb) of ecDNAs carrying *MDM2*. Scissor shape symbols indicate the location of the CRISPR induced DSBs (GRCh38). Blue bar, circular junction site **(b)** Graphical illustration of the CRISPR-C based generation of 578 Kb size *MDM2* carrying ecDNA and the created deletion site on chromosome 12. Orange bar: deletion scar, blue bar: circular junction site. Gray arrows indicate map positions of the forward (F1, F3) and reverse (R1, R3) primers used in PCR assays to amplify the unedited sgRNA mapping sites (F1+R1, F3+R3), the deletion scars on chromosome 12 (F1+R3), and the circular junction site on the ecDNAs (F3+R1). **(c)** Validation of the *MDM2* 578 Kb size ecDNA circular junction site and deletion scar at chromosome 12 of origin by PCR and Sanger sequencing. White asterisk indicates the sequenced PCR products. **(d)** Scheme of *MDM2* and *CDK4* harboring chimeric ecDNA derived from two separate fragments of the same chromosome (chromosome 12) by CRISPR-C upon the introduction of sgRNAs targeting four cleavage sites. **(e)** Inverse PCR based genotyping of circular junction site by PCR and Sanger sequencing. Asterisks indicate the targeted PCR products characterized by Sanger sequencing.

### Fluorescent in situ hybridization (FISH)

U2OS, MCF7, and human skin fibroblasts, were analyzed by FISH to detect deletions or ecDNAs, generated via the CRISPR-C 2.0 system, encompassing *CDK4*, *MDM2*, and *ABCB1* oncogenes. For the *MDM2* and *CDK4* genes, two Bacterial Artificial Chromosome (BAC) clones encompassing the whole target region were selected according to the GRCh38/hg38 human reference genome from the UCSC Human Genome Browser (http://genome.ucsc.edu) (**Supplementary Table S6**), respectively labeled with Cy3- and Cy5-conjugated dUTP by nick translation. Conversely, for *ABCB1*, a single probe directly labelled with Cy3-dUTP was used. Additionally, whole chromosome painting (WCP) probes, derived from flow-sorted chromosomes and amplified by degenerated oligonucleotide probe PCR, were directly labeled with FITC-dUTP and used to identify the original mapping position of target genes. The lack of target gene signals by BAC probes, or the presence of fainter signals compared to those on homologous chromosomes, indicated full or partial oncogene deletion.

FISH was performed as previously described (22). Digital images were obtained using a Leica DM-RXA2 epifluorescence microscope equipped with a cooled CCD camera (Princeton Instruments, Boston, MA). Cy3 (red), FITC (green), Cy5 (infra-red), and DAPI (blue) fluorescence signals were detected using specific filters. Images were separately recorded as gray-scale images. Pseudocoloring and merging of images were performed using the Adobe Photoshop software.

### MTT assay, Formazan quantification, cell counting and Formazan imaging

6 days and 20 days after CRISPR-C induction of three different sizes of *CDK4* and *MDM2* ecDNA formation (triplicates of each group) in U2OS cells, cells were seeded in 24-well plates at 2 × 10^4^ cells/100 µl and cultured at 37 °C, 5% CO_2_ for 24 hours into a 96 well 96-Well, Nunclon Delta-Treated, Flat-Bottom Microplate (Thermo Scientific). On the following day 10 µl MTT labeling reagent (5 mg/ml) of the Cell Proliferation Kit I (Roche) was added and cells were incubated at 37 °C, 5% CO_2_ for 3 hrs followed by the addition of solubilization solution of the MTT kit and further incubation for 4 hours at 37 °C, 5% CO_2_. The absorbance was read at 570 nm using 630 nm as a reference wavelength on a SpectraMax iD5 Multi-Mode Microplate Reader (Molecular Devices). At each time point 10^5^ cells were lysed and genotyped using inverse PCR for detection of the presence of circular junction sites.

### Cell proliferation assay with EdU

To measure DNA synthesis, 3 × 10^5^ cells were seeded into T-25 cell culture flasks 6 days after CRISPR-C generation of three different sizes of *CDK4* and *MDM2* ecDNAs (triplicates of each group) in U2OS cells. 18-24 hrs after Click-iT® EdU was added to the culture media to the final concentration of 10 µM and incubated it for 1.5 h. After incubation cells were harvested and fixed and the labelling of EdU incorporation was conducted by Click-iT™ EdU Alexa Fluor™ 488 Flow Cytometry Assay Kit (Molecular Probes). EdU-Alexa Fluo 488 positive cells were detected by flow cytometry on Quanteon analyzer and data analysis was performed using FlowJo_V10.8.1.

### Generation of palbociclib and colcemid resistance cell-populations

Three different sizes of *CDK4* harboring ecDNAs were generated by CRISPR-C 2.0 in U2OS cells (unedited control, 1 Mb, 508 Kb and 201 Kb size *CDK4* coding region carrying ecDNA). 5 days after the onset of CRISPR-C 2.0. U2OS cells were subjected to 1 µM palbociclib (Sigma) treatment for 30 days and 3 µM palbociclib treatment for 70 days with medium replaced in every 3^rd^ day. Persistence of CRISPR-C:*ABCB1*-ecDNA deletion and circular junction sites were screened using PCR.

Three different sizes of *ABCB1* harboring ecDNA were generated by CRISPR-C 2.0 in MCF7 cells (unedited control, 1 Mb, 520 Kb and 244 Kb size *ABCB1* coding region carrying ecDNA). 5 days after the onset of CRISPR-C 2.0, MCF7 cells were subjected to 5 ng/ml colcemid (KaryoMAX™ Colcemid™ Solution in PBS, Gibco, Thermo Fisher Scientific) treatment for 14 days and 10 ng/ml colcemid treatment for 45 days with the medium replaced in every 3^rd^ day. In parallel, each nucleofection group was cultured in normal medium without colcemid selection pressure. Persistence of CRISPR-C:*ABCB1*-ecDNA deletion and circular junction sites were screened using PCR.

### RNA sequencing and differentially expressed genes analysis

5×10^5^ cells were collected every six days at five consecutive time points over 30 days following the *CDK4* 508 Kb, the *MDM2* 582 Kb and the chimeric ecDNA from the same *CDK4* and *MDM2* region generation by CRISPR-C 2.0, and total RNA was isolated by the miRNeasy Tissue/Cells Advanced Micro Kit (QIAGEN, Cat.no.217684,Germany). After RNA purification, 800ng RNA was used per sample as input for the RNA library preparation using NEBNext® rRNA Depletion Kit v2 (NEB, E7405L, USA) and NEBNext® Ultra™ II Directional RNA Library Prep Kit for Illumina (NEB, E7760S, USA) following the manufacturer’s recommendations. Each library was ligated with different indexes using NEBNext® Multiplex Oligos for Illumina (NEB, E6440S, USA) kit and amplified with 11 PCR cycles according to the instructions. The quality and quantity of the cDNA libraries were evaluated using the Qubit Fluorometer (Invitrogen, Q32857,USA) and the Bioanalyzer 2100 (Agilent). The RNA libraries were sequenced on an Illumina (Novaseq 6000, SP flow cell, Rigshospitalet, Copenhagen, Denmark) platform with 150 bp paired-end reads.

The quality control (QC) of the raw sequences was performed using the FastQC software (v0.12.0). Low-quality reads and adapters were removed by Fastp (v0.23.3). The processed reads were aligned to the hg38 reference genome by Hisat2 (2.2.1). After the alignment, the subsequent mapped reads were quantified by featureCounts (2.0.0).

DESeq2 (1.38.3) was used to normalize and perform differential expression analysis on the quantified reads, Genes with an absolute value of logarithmic fold change (log2FC) > 0.58 were qualified for the final set of differentially expressed genes (DEGs). Gene Set Enrichment Analysis (GSEA) identified enriched gene sets from the normalized count data and Kyoto Encyclopedia of Genes and Genomes (KEGG) enrichment analysis was implemented by the cluster profile R package. Terms with a corrected p value < 0.05 were considered significantly enriched by DEGs.

### Data availability

The raw RNA sequencing data are accessible to the public as of the publication date through the SRA database (BioProject number PRJNA1062599). The eccBiosensor plasmids are available through the public plasmid repository of the Luo lab (https://www.addgene.org/Yonglun_Luo/; plasmid ID: 140832 and 140833). All cell lines with the corresponding oncogene-carrying ecDNA can be requested from the Luo lab under the condition of proper MTA being signed.

### Statistics

GraphPad Prism 9 (GraphPad Software, Inc., USA) was utilized for the visualization of flow cytometry, MTT viability, and ddPCR data, with results presented as mean ± SD. Statistical analyses were performed using one-way ANOVA followed by Tukey’s multiple comparisons test, comparing each CRISPR-C treated sample group to the Cas9-treated control groups or to different time points.

For RNAseq analysis, significance across all cases was assessed using two-tailed paired t-tests, with a significance threshold set at p-value ≤0.05. Data plots were generated using GraphPad Prism 8.

## RESULTS

### CRISPR-C 2.0 is a highly efficient and precise molecular tool for DNA circularization in cells

In our previously introduced CRISPR-C system (CRISPR-C 1.0)(20), human cells transiently expressed the Cas9 protein and the two sgRNAs after transfection with the vector plasmids. However, CRISPR-C 1.0 is subject to limitations inherent in plasmid-based transient gene delivery strategies. These limitations include low delivery efficiency, which varies depending on the cell type, and uncertainty regarding the duration of vector retention and active transcription in cells.

To address these challenges, we implemented modifications and developed an upgraded version known as CRISPR-C 2.0. In this advanced version, we adopted the use of preassembled ribonucleoprotein complexes (RNPs), which have recently gained widespread application in CRISPR strategies. This approach offers high delivery, editing efficiency, and precise scalability (23,24). In CRISPR-C 2.0, two precisely defined double-stranded breaks (DSBs) are introduced by a pair of RNPs formed by Cas9 proteins and two synthetic sgRNA molecules. The assembled RNPs are delivered through optimized electroporation (see Materials and Methods section). CRISPR-C 2.0 represents a significant improvement over CRISPR-C 1.0 (P < 0.001) when tested with our eccDNA biosensor cells (detailed description in Moller et al. 2018 (20) and depicted at **Supplementary Figure S1a and b**), addressing the limitations associated with plasmid-based delivery and providing enhanced efficiency in gene editing.

Forty-eight hours after the CRISPR-C induction, the ecDNA biosensor line experienced a 7-8-fold increase in ecDNA generation and inversion formation frequency when utilizing the RNP-based method, in contrast to the plasmid-based delivery system (**Supplementary Figure S1c**). Additionally, it was observed that the fluorescent signals manifested 24 hours earlier with CRISPR-C 2.0 compared to CRISPR-C 1.0. Regarding the fluorescent cell quantity and median fluorescence intensity of the fluorescent cells, a more rapid increase followed by a steady decline was noted when employing the RNP-based delivery system, as depicted in **Supplementary Figure S1d**. These findings indicate the superior and more synchronized ecDNA generation capability of the CRISPR-C 2.0, highlighting its promptness compared to the delayed response associated with plasmid-based delivery.

### Generation of oncogene-carrying ecDNAs with CRISPR-C 2.0

Using CRISPR-C 2.0, we sought to recapitulate the ecDNA formation from specific chromosome regions (**Supplementary Table S1**) in a high number of cells. To achieve this, we designed sgRNAs targeting neighboring regions of eight selected oncogenes, known to be recurrently amplified on circular amplicons and well studied in the field of mammalian circular DNA biology. These oncogenes include *EGFR, CDK4, MDM2, MYC, MYCN, FGFR2, ABCB1* and *DHFR.* The *EGFR, CDK4, MDM2, MYC, MYCN, FGFR2* are oncogenes recurrently amplified on circular amplicons, and *ABCB1* and *DHFR* whose increased copy number and expression play a role in chemotherapeutic drug resistance during cancer development (**Supplementary Figure S2**).

We designed CRISPR-C gRNAs using the CRISPRon deep-learning tool to select gRNAs with high efficiency and specificity (25). For each oncogene, we generated ecDNAs of diverse size spanning from 100 Kb to 1 Mb as detailed in **Supplementary Table S2** encompassing the size range of ecDNA harbored by cancer cells (5,14,15). The editing efficiency of each designed sgRNA was assessed by Sanger sequencing and the ICE (Inference of CRISPR Edits) algorithm, which showed an average of 70% efficiency (**Supplementary Table S1**). Using these sgRNAs, we performed CRISPR-C 2.0 to induce two DSBs and generate ecDNAs containing the oncogenes of interest. This encompassed the broad size spectrum of oncogene ecDNAs detected in cancer cells (14,15). In the case of six selected regions (the carrying region of *ABCB1, CDK4, FGFR2, MDM2, MYC* and *MYCN* genes), we generated three different sizes of ecDNAs as the graphical design shows in **Figure 1a**, one size of *EGFR* and *DHFR* harboring regions, respectively (**Supplementary Table S2**).

To detect the generation of ecDNAs in HEK293T cells, we used PCR assays with four appropriate primer combinations to amplify the each sgRNA target sites, the deletion scars generated after the displacement of the sequence located between the two CRISPR cut sites, and the junction sites formed after the circularization of the deleted DNA fragment. **Figure 1b** illustrates the combination of primer pairs used for PCR detection and Sanger sequencing analysis (Figure 1c), through the example of CRISPR-C 2.0-induced generation of a 587 Kb ecDNA containing *MDM2*. Our results confirmed circular junction sites and an equal number of deletion scars through PCR and Sanger sequencing (**Supplementary Table S3; Supplementary Figures S3-S10.**).

### Unique indel profiles at the CRISPR-C generated ecDNA junctions

The generation of ecDNAs using CRISPR-C involved the DSB repair machinery in cells. We analyzed the formed junction sites using Sanger sequencing. We observed that ∼37% of the circular junction sites and ∼42% of the deletion scar sites retained the original sequence, showing a preference for indel-free ligation of the break sites (non-homologous blunt end joining, NHEBJ). It seems to be a highly conserved repair phenotype, which we previously harnessed for predictable gene therapy (26). Approximately 37% of the circular junction sites and ∼21% of the deletion scar sites had 1 or 2 bases insertion or deletion, where the junction site sequence could be matched with the original sequences. However, in ∼26% of the circular junction sites and ∼32% of the deletion sites the Sanger sequencing profiles showed strong disturbances and no resemblance to the original sequence after the breakpoint induction (**Supplementary Table S4**).

The signature of accurate error-free ligation (27,28) or small base indels at the ligation sites indicates that non-homologous end-joining (NHEJ) (29) was the dominant form of junction formation in the majority of the CRISPR-C 2.0 induced rearrangements. Furthermore, the high prevalence of mixed indel pattern in many instances is the signature of an error-prone repair mechanism, where we cannot exclude the involvement of other error-prone repair mechanisms, such as microhomology-mediated end-joining (MMEJ) and single-strand annealing (SSA) besides the NHEJ (30). Interestingly, nearly all the utilized sgRNAs introduced several indels demonstrated by the ICE analysis (**Supplementary Figure S13-S20**). However, Sanger sequencing of the junction sites indicates that blunt end joining contributes significantly to the circle formation events. These results imply that the formation of large-size DNA circles might use a different mechanism than DNA end joining.

### Broad applicability of CRISPR-C 2.0 in human cell lines

To demonstrate the broad applicability of CRISPR-C 2.0 as a tool for generating large ecDNAs, we conducted ecDNA generation experiments in various human cancer cell lines, including HeLa, U2OS, MCF7, primary human skin fibroblasts, and human astrocytes (**Supplementary Table S2**). To assess nucleofection efficiency, a fraction of each listed cell type was nucleofected with EGFP-coding mRNA.(**Supplementary Figure S21**). The nucleofection efficiency exceeded 90% in all cell types, indicating the high efficiency of our delivery method. We then performed CRISPR-C 2.0 to generate ecDNAs using the designed CRISPR-C sgRNA pairs and validated the results by PCR in the U2OS cell line to demonstrate the applicability of CRISPR-C in cancer cell lines. We also generated CRISPR-C:*ABCB1* (31–33) ecDNA in the MCF7 and HeLa cells, CRISPR-C: *DHFR* (2) ecDNA in HeLa cells, and CRISPR-C: *CDK4* and CRISPR-C: *EGFR* (7,34) in human glioblastoma cell lines, due to their relevance in previous studies. These were all validated by PCR. **Supplementary Table S2** summarizes the tested CRISPR-C-induced ecDNA formations in these cell lines.

### Validation of CRISPR-C generated ecDNA with FISH

To further validate extrachromosomal nature of the CRISPR-C 2.0 generated ecDNAs, we performed FISH assays on metaphase cells oncogene specific and whole chromosome painting (WCP) probes. This analysis was conducted on U2OS osteosarcoma cells one week after the CRISPR-C 2.0 procedure targeting genomic regions of *CDK4* and *MDM2*. In detail, to detect *CDK4* regions, two overlapping BAC clones were used for FISH analysis: RP11-936I7 (blue) and RP11-846E20 (red) (**Supplementary Table S6, Supplementary Figure S22a**). As shown in the left panel of **Figure 2a**, the unedited U2OS cells had three copies of *CDK4* (as colocalizing FISH blue and red signals), with one copy on chromosome 12 displaying normal WCP green staining, another revealing an aberrant form of chromosome 12, and one copy situated on a different chromosome lacking WCP 12 staining. The signals of both probes are absent in one of the two chromosomes 12/derivative chromosome 12 [der(12)] for the 1 Mb and 508 Kb ecDNAs (**Figure 2a**, two middle panels, **Supplementary Figure S23**b and c) in 23.5% and 43.7% of the analyzed metaphases, respectively, indicating the deletion of the target regions (**Supplementary Table S5**). For the shorter ecDNA (201 Kb), a faint red signal for RP11-846E20 (red) indicated the retained region of the scar on the der (12) (**Figure 2a**, right panel, **Supplementary Figure S23d**). In this case, we detected the deletion in 53.5% of the observed cells. The extrachromosomal signals corresponding to the target locus were observed in two cells per CRISPR-C group for the 508 Kb, and 201 Kb ecDNAs, where the same cells also showed the deletion of one copy of the same region, affirming the deletion of the targeted region from the chromosome and the retention of it in the cell as an extrachromosomal copy (**Figure 2c, Supplementary Figure S25a-b**).

**Figure 2:**
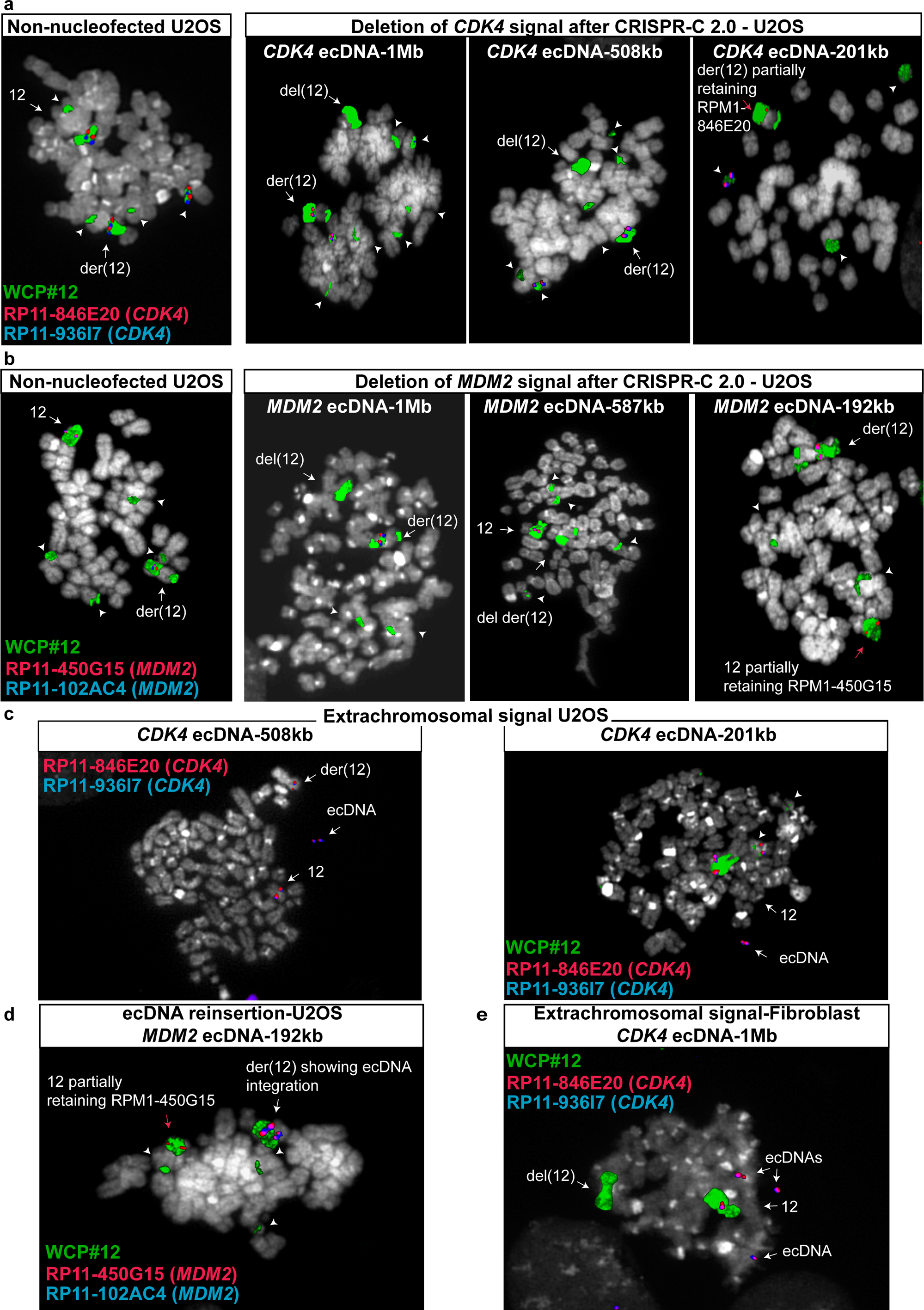
CRISPR-C induced chromosome alterations targeting *CDK4* and *MDM2* containing regions. DNA-metaphase-FISH detection of rearrangements in U2OS cells and skin fibroblast cells. Grey: DAPI staining of DNA; Green: WCP of chromosome 12; Red and Blue: BAC clones RP11-846E20 and RP11-936I7 respectively for CDK4, (a) and RP11-450G15 and RP11-1024C4 respectively for *MDM2*, (b) containing regions (details in **Supplementary Table S6**). **(a), (b)** CRISPR-C 2.0 instigated deletion of 1 Mb, 508 Kb and 201 Kb *CDK4* containing regions and 1 Mb, 500 Kb and 150 Kb *MDM2* carrying fragments in U2OS cells. **(c)** *CDK4* positive ecDNA signal in *CDK4*:CRISPR-C 2.0 treated U2OS cells (508 Kb and 201 Kb). **(d)** Reinsertion of 192 Kb *MDM2* containing region upon CRISPR-C 2.0 induction in U2OS. **(e)** Three *CDK4* positive ecDNA signals in CDK4:CRISPR-C 2.0 treated fibroblast cells, and one deletion on normal chromosome 12 of the target genomic region (1 Mb).

To detect *MDM2* regions, we used the BAC clones RP11-450G15 (red) and RP11-1024C4 (blue) in FISH assays (**Supplementary Figure 22b**, **Supplementary Table S6**). As shown in Figure 2b (left panel) and **Supplementary Figure S24**, the unedited cells showed two copies of *MDM2* (colocalizing blue and red signals) on two chromosome 12, exhibiting normal or abnormal chromosome 12-specific WCP green staining. As shown in **Figure 2b** (two middle panels) and **Supplementary Figure S24** second and third panel, the deletion of the genomic regions corresponding to the 1 Mb (reported in 11.1% of cells) and 587 Kb (affecting the 11.7% of cells) was detected by the lack of signal for both probes in one of the two chromosomes 12/der(12) (**Figure 2b, Supplementary Table S5**). Similarly to what was shown for the shortest ecDNA containing CDK4 (**Figure 2a**), also for the shortest ecDNA containing *MDM2* (192 Kb), a faint red signal for RP11-450G15 (red) indicated the undeleted region of the scar on the normal chromosome 12 in the 47.3% of observed cells (**Figure 2B**, right panel and **Supplementary Figure S24d**, **Supplementary Table S5**). Additionally, we observed the translocation of a 192 Kb fragment carrying the *MDM2* gene, indicating that the generated extrachromosomal fragment can be reintegrated onto chromosomes (**Figure 2d, Supplementary Figure S25c**). Also, the low number of extrachromosomal signals compared to deletion events shows that most of the generated fragments are cleared out from the cells (**Supplementary Table S5)** and only a fraction of them were sustained for a longer time period.

Since the genome of most cancer cell lines is heavily rearranged, including that of U2OS cells, which might interfere with the precise quantification through FISH, we employed skin fibroblasts from healthy human donors with a diploid genome to circumvent these challenges and enhance the accuracy of our analyses. As expected, we observed two *CDK4* or *MDM2* genomic loci (**Supplementary Figure S26a, b**) located on the chromosome 12 of the unedited cells. We detected deletions of the targeted region in three out of the six CRISPR-C groups (**Supplementary Table S5**). Additionally, we observed the deletion of the 1 Mb region carrying the *CDK4* gene from one chromosome 12 and the presence of three extra-chromosomal copies of the targeted fragment in the same cells, indicating copy number amplification of the generated excised genomic fragment (**Figure 2e, Supplementary Figure S26c, Supplementary Table S5**).

### Modelling of chimeric ecDNA with CRISPR-C 2.0

While specific cancer cell types have been reported to harbor large circular DNAs derived from a single chromosomal region (35), the sequences of the majority of described ecDNAs are typically derived from more than 20 different genomic regions (7,36). To investigate whether our system can trigger the generation of chimeric ecDNAs, we selected target regions around *CDK4* and *MDM2* to produce ecDNA containing both fragments. Notably, these two genes are frequently coamplified in neuroblastoma (37) and glioblastoma (7) on the same circular amplicon. We co-delivered four sgRNAs and Cas9 protein as RNPs into the U2OS osteosarcoma cell line to generate two approximately 500 Kb long chromosomal fragments located on chromosome 12. We anticipated the ligation of the two cleaved fragments in two possible ways (**Figure 1d**), resulting in four different circular junction sites and two forms of chimeric circular DNA. Each junction site was detected by PCR and confirmed by Sanger sequencing (**Figure 1e, Supplementary Figure S10b and c**).

To assess the ability of our CRISPR-C 2.0 system to induce the formation of ecDNAs derived from different chromosomes, we targeted genomic regions of chromosome 12 carrying the *CDK4* gene and chromosome 7 harboring the *EGFR* gene, as these genes are known to co-amplify frequently in glioblastoma (38). HEK-293T cells were co-nucleofected with two combinations of sgRNAs: combination #1 involved CRISPR-C sgRNAs for generating a 668 Kb ecDNA of *EGFR* and a CRISPR-C sgRNA for generating a 508 Kb ecDNA of *CDK4*, while combination #2 involved CRISPR-C sgRNAs for generating a 668 Kb ecDNA of *EGFR* and a CRISPR-C sgRNA for generating a 201 Kb ecDNA of *CDK4*. In total, we expected the formation of four ecDNAs and eight circular junction sites. Indeed, we successfully detected these using PCR and Sanger sequencing as depicted in detail in **Supplementary Figures S11-12**.

### The effect of targeted generation of pan-oncogene carrying ecDNAs on cell function and transcriptome

The current body of evidence supports the notion that oncogene-carrying circular amplicons present in high copy number exert diverse effects on the functionality and transcriptomics of the host cells (10,16). Nevertheless, our understanding of how newly formed ecDNAs harboring oncogenes impact gene expression and cellular activity in cancer cells remains limited. After confirming the presence of unique junction circular sites and detecting ecDNA generation along with other chromosomal rearrangements using FISH (**Figure 3a**), we examined the cellular effects of CRISPR-C targeting of *CKD4* and *MDM2* genes, both of which play crucial roles in cell cycle regulation and growth.

**Figure 3.**
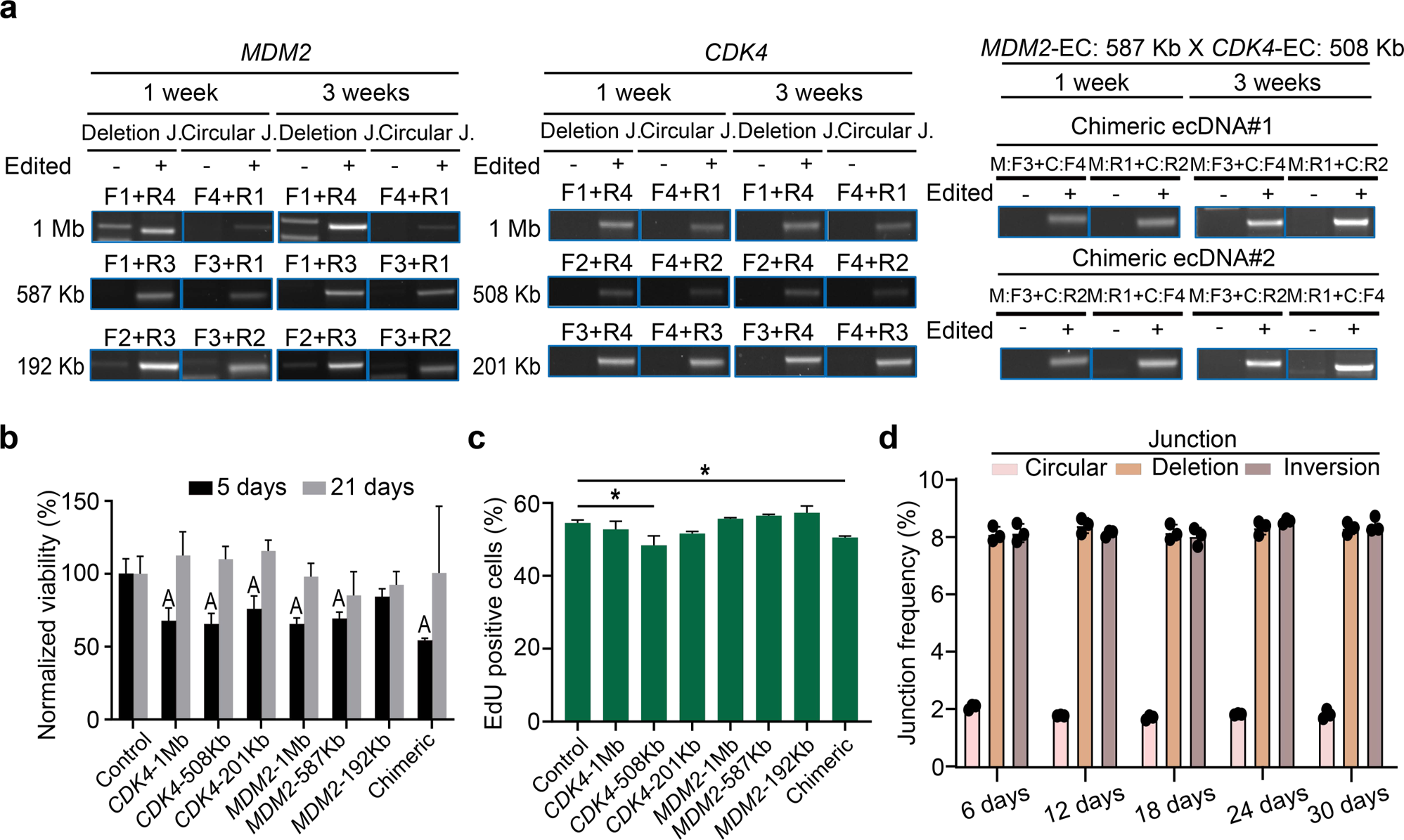
Persistence and cellular effect of CRISPR-C 2.0 generated pan-cancer oncogene. **(a)** Assessment of the presence of junction sites formed after deletion and DNA fragment circularization in U2OS samples after *CDK4-MDM2*:CRISPR-C 2.0. treatment followed over one month sampled at each passage through five passages. **(b)** Histograms of MTT viability assays normalized to measured viability of the unedited control group performed five and 21 days after the *CDK4-MDM2*:CRISPR-C 2.0 procedure in U2OS cells. “A” represents statistically significant difference related to control with p<0,05 Error bars: SD.; n=3. **(c)** Cell proliferation level detected by Click-iT EdU cell proliferation assay one week after *CDK4-MDM2*:CRISPR-C 2.0 in U2OS cells. (*) indicates statistical p < 0,05 compared to control. Error bars: SD.; n=3, **(d)** Quantification of ecDNA junction sites, deletion site and inversion of CRIPSR-C targeted *CDK4* 508Kb by ddPCR. Frequency is quantified by comparing to the copy number of the TERT reference gene n=3, error bars: SD; black dots, individual sample values.

Analysis of the MTT assay data revealed a substantial reduction in viability across all treated cell groups (except the 192 Kb *MDM2* ecDNA) five days post CRISPR-C, followed by a subsequent restoration of viability three weeks after treatment (**Figure 3b**). Moreover, during the initial week following CRISPR-C treatment, we observed a notable decline in cell proliferation specifically within the 508 Kb *CDK4* circle and chimeric circle groups (**Figure 3c**). Our PCR genotyping result confirmed the presence of the deletion scar and circular junction sites even after the restoration of viability (**Figure 3a**). We designed a ddPCR assay specific for the circular junction site to quantify the circularized fragments in cells. Interestingly the frequency of the circularized fragment did not change significantly over one month, including the two time-points of the viability assessment (**Figure 3D**). Due to the low fraction of ecDNA in cells, these dynamic changes in viability might be attributed to frequent deletions occurring within the genomic regions, leading to the sole overgrowth of cells that retain ecDNAs carrying *CDK4* and/or *MDM2*, or the proliferation of cells unaffected by the CRISPR-C intervention.

Next, we examined the transcriptional consequences of CRISPR-C induction of about half-megabase size *CDK4*, *MDM2* and their chimeric circle for one month following the CRISPR-C procedure. Firstly, we monitored the temporal changes in transcript levels of the targeted genes carried by the CRISPR-C circles. We observed a general down-regulation in the transcriptional activity of the genes located within the CRISPR-C target regions across most experimental groups (**Figure 4a**), indicating prevalent gene deletions at that particular time point which aligned with the findings from the cell viability and proliferation assays. Intriguingly, at the initial time point in the chimeric ecDNA group the transcriptional level of *MDM2* exhibited a significant up-regulation. This result was consistently observed in three parallel experiments, suggesting a potential influence arising from the formation of the chimeric ecDNA structure.

**Figure 4.**
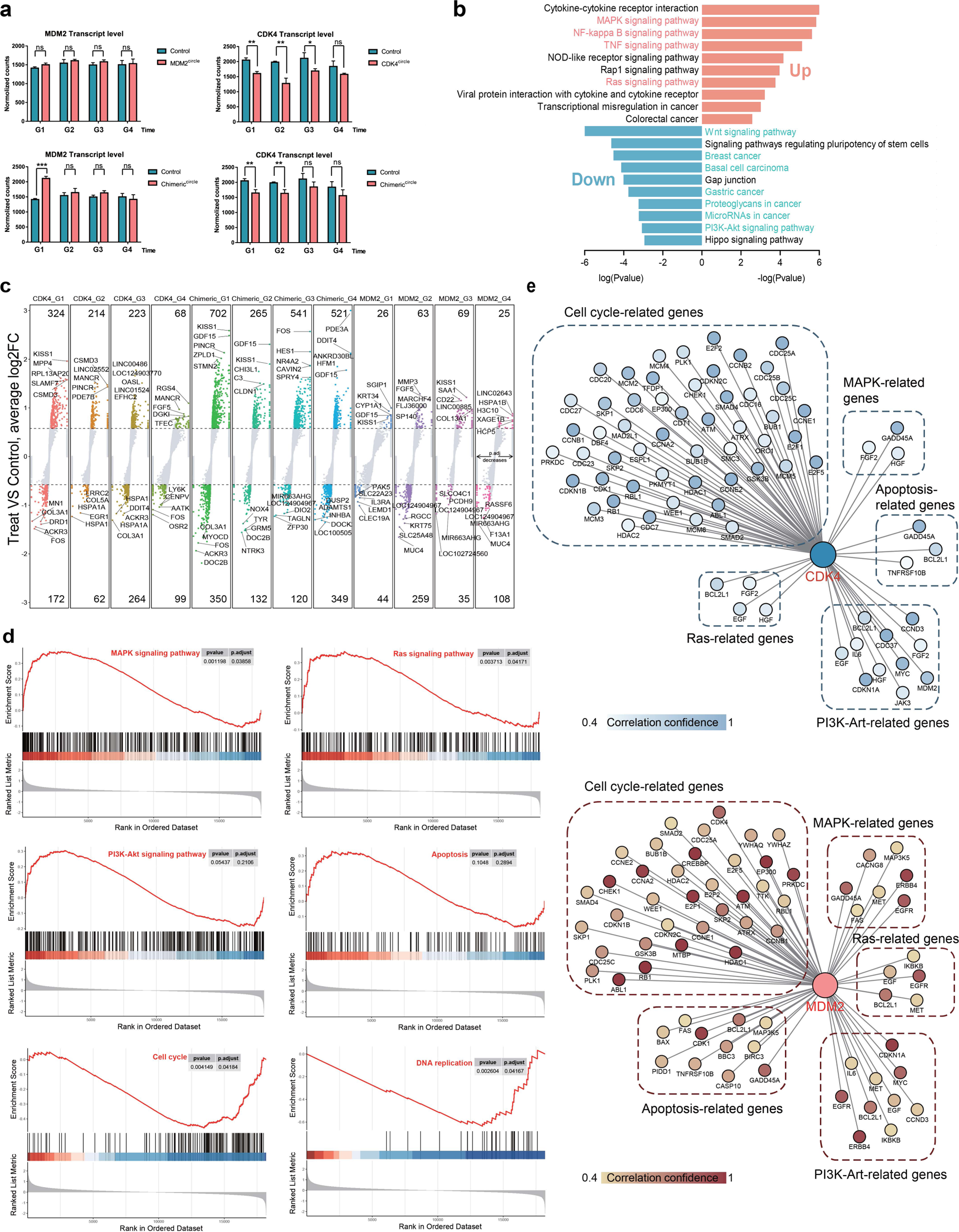
Characterization of RNA profiles in the *CDK4* ecDNA, *MDM2* ecDNA and *CDK4-MDM2* chimeric ecDNA cell lines. **(a)** Comparative gene expression profiles of *MDM2* and *CDK4* in control and CRISPR-C edited cell lines. Transcript levels are quantified using normalized counts. **(b)** Volcano plot of differentially expressed genes (DEGs) in three CRISPR-C treatments cell lines compared to control cell lines across the four cell generations. **(c)** KEGG analysis of DEGs in the first generation of the chimeric ecDNA cell line. Left: significant KEGG enrich terms of up-regulated genes. Right: significant KEGG enrich terms of down-regulated genes. The top 20 significant KEGG terms were visualized. **(d)** Enrichment score of the KEGG pathways analyzed by GSEA based on RNA-seq datasets. **(e)** Correlation of *MDM2* and *CDK4* expression with the expression profiles of genes related to “Apoptosis”, “MAPK Signaling pathway”, “Ras Signaling pathway”, “Cell cycle” and “PI3K-Art” pathways based on RNA-seq data. The color shade represents the strength of the correlation.

We then conducted differential gene expression analyses within each experimental group. Remarkably, at the initial time point, the chimeric ecDNA group displayed the highest number of both up-regulated and down-regulated genes (**Figure 4B**). Based on these findings, we proceeded with further analysis to unravel the functional implications of these gene expression changes focusing on the chimeric ecDNA group at the first time point. We conducted comprehensive analyses, including the Kyoto Encyclopedia of Genes and Genomes (KEGG) enrichment analysis and gene set enrichment analysis (GSEA), to unveil and comprehend the enriched biological pathways and processes linked to the differentially expressed genes.

The KEGG analysis revealed that the upregulated genes were predominantly enriched in inflammatory pathways implicated in the development and progression of cancer, including MAPK, TNF, Ras, and NF-kappa B signaling pathways (**Figure 4C**). Concurrently, GSEA revealed the activation of these key pathways including MAPK, TNF, Ras, PI3K-Akt, and Apoptosis (**Figure 4D** and **Supplementary Figure S29**). Each of these signaling pathways is intricate and interconnected, and therefore dysregulation or aberrant activation of these pathways can lead to pathological conditions and impact normal cellular function. Furthermore, the KEGG analysis demonstrated that the down-regulated genes were prominently enriched in pathways associated with cell growth and cancer, including the Wnt signaling pathway (identified as the top pathway among the down-regulated genes in the KEGG enrichment analysis), which plays a crucial role in maintaining cellular homeostasis, driving oncogenic transformation, facilitating tumor progression, and promoting metastasis. GSEA analysis demonstrated suppression of DNA replication and Cell cycle pathways (**Figure 4D**), which aligns with the findings from the cell proliferation assay (**Figure 4C**). Moreover, The String database revealed a strong correlation between the expression of *MDM2* and *CDK4* and the expression of several genes enriched in these cellular pathways (**Figure 4E**). Our results indicate a comprehensive context-dependent effect of CRISPR-C-generated oncogene ecDNA on their transcriptional activities.

### Persistence of CRISPR-C 2.0 generated ecDNA in cells

To examine the persistence of CRISPR-C 2.0-generated ecDNAs containing oncogenes in cells, we monitored the presence of ecDNAs harboring *CDK4* (**Supplementary Figure S4**), *MDM2* (**Supplementary Figure S3**), or both genes (**Figure 1e and e**). These two pan-oncogenes were selected because of their frequent recurrence and significant role in cell-growth. Cells were cultured for 30 days after CRIPSR-C 2.0 induction of ecDNA generation. Cell samples were taken at each of the five passages of the investigation period. The presence of the CRISPR-C 2.0 generated deletion sites and circular sites were tracked with PCR assays. Both, the deletion site and the circular junction site were detected at each timepoint from all of the three different sizes of *CDK4* (**Supplementary Figure S30**) or *MDM2* (**Supplementary Figure S31**) harboring ecDNA and chimeric DNAs (**Supplementary Figure S32**) indicating the retention of the generated ecDNAs over 30 days.

To quantify the frequency of the fragments originated by the circularization and retained in the cells, we performed ddPCR (**Supplementary Figure S33**) on DNA samples purified from subset of cells collected at each passage as described above. We selected the 508 Kb *CDK4* ecDNA for this study, as we could successfully confirm the occurrence of ecDNA by FISH. Our results indicate that only a small percentage (2%) of the target sequences were present in the cells as a circularized fragment, in agreement with our FISH-based ecDNA assay. The frequency of deletion and inversion of the target sequence was also low at approximately 8 %, when assessed by ddPCR. Interestingly the frequency of the unique circular junction sites deletions and inversions did not show any prominent changes over the investigated period (**Figure 3D**).

### Drug selection induces retention and copy-number increase of CRISPR-C targeted pan-oncogene regions

Our ddPCR results suggest that ecDNAs may endure/persist in cells during the initial period following the introduction of the two DSBs without additional selection pressure. These findings, coupled with the broader understanding of ecDNA, prompt the question of whether these ecDNAs exhibit prolonged persistence within cells and whether their copy number undergoes alterations under certain selection pressure. To get further insight into the early dynamics of ecDNAs, we followed the persistence of different CRISPR-C generated ecDNAs in the presence or absence of drug selection.

The *CDK4* coding region was chosen because it is consistently identified as the most frequently occurring gene on circular amplicons in various cancers, and it is known to promote cancer growth. We hypothesize that the introduction of a CDK4 inhibitor as an additional selection pressure may impact the retention of the CRISPR-C targeted sequence. If cells adapt to the inhibition by activating alternative signaling pathways, we expect the elimination of the *CDK4*-harboring fragment (39). Conversely, cells may respond by increasing the expression of the blocked protein to restore their growth capacity, potentially involving the preservation or even an increase in the *CDK4* copy number (40). In this instance, the retention and possible copy number increase of the CRISPR-C generated ecDNA could be observed.

To test our hypothesis, we carried out CRISPR-C to induce the formation *CDK4*-carrying ecDNAs (**Supplementary Figure S4**). These cells were then cultured for a long period of time (100 days) in an escalating concentration of palbociclib, a specific inhibitor for cyclin-dependent kinase 4/6 (**Figure 5a**). First, we carried out PCR screening of the CRISPR-C circular junction sites. After one month all the tested junction sites were present in the cells, as it was observed previously in cells cultured without drug selection (**Figure 5b**). After further 70 days of both culturing under increased concentration of palbociclib and in the absence of it, the circular junction site and the deletion scar were detectable in all the three kinds of CRISPR-C ecDNAs. Notably, the 508 Kb ecDNA junction site was strongly detected in the drug-treated group. Using FISH analysis, we further confirmed the presence of *CDK4*-carrying ecDNAs in the 508 Kb and 204 Kb CRISPR-C *CDK4* groups (Figure 5c). It is important to mention that the visualized *CDK4* positive ecDNA copy number remained low. No cells with high copy-number increase was observed and no *CDK4*-positive extrachromosomal elements was observed in the drug-treated CRISPR-C non-edited group, indicating that the drug selection alone did not induce *CDK4* ecDNA formation (**Figure 5c, Supplementary Figure S27**).

**Figure 5.**
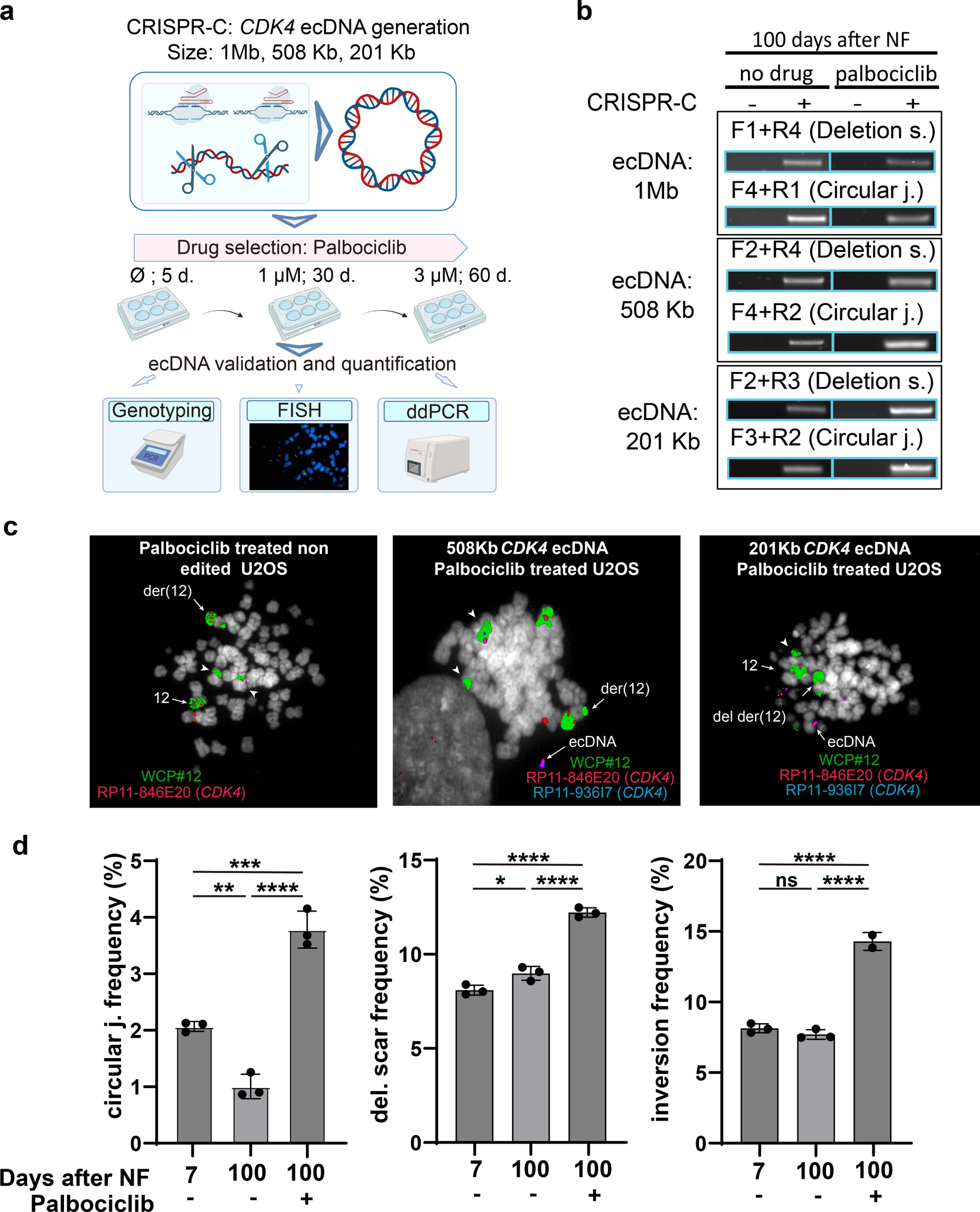
In vitro modelling of drug selection dynamics after *CDK4* carrying ecDNA generation by CRISPR-C. **(a)** Workflow of the assessment of drug-selection effect on CRISPR-C generated CDK4 ecDNA persistence. **(b)** Inverse PCR validation of the presence/absence of circular junction site after CRISPR-C 2.0 generation of three different sizes of ecDNA cultured under palbociclib drug selection for 100 days. **(c)** Metaphase FISH analysis of *CKD4* in U2OS treated with palbociclib, as indicated in the A panel of the figure. Blue and red signals: BAC clones RP11-846E20 and RP11-936I7 for the CDK4 region, respectively; Green: WCP of chromosome 12 DAPI: Grey. Left image: non-edited cells showing three copies of *CDK4* (as colocalizing blue and red signals). The central panel shows the presence of an extrachromosomal CDK4 signal after the induction of the CRISPR-C *CDK4* ecDNA of 508 Kb. The right panel shows the deletion of one original *CDK4* signal and an extrachromosomal CDK4 signal upon induction of the CRISPR-C *CDK4* ecDNA of 201 Kb. **(d)** Charts showing the frequencies of CRISPR-C *CDK4*-ecDNA 508 Kb circular junction, deletion scare, and inversion events assessed by ddPCR. Y axis depicts the frequency of the junction site relative to TERT gene locus copy number as a reference sequence. The asterisks of *, **, ***, and **** represent p values less than 0.05; 0.01, 0.001, and 0,0001 respectively. Error bars: SD; black dots=individual sample values.; ns, not significant; n=3.

To obtain a quantitative estimation of the effect of palbociclib treatment on CRISPR-C generated ecDNAs, ddPCR was utilized to assess the presence of the circular junction site of the 508 Kb CDK4 ecDNA and the formed deletion scars, as well as inversions. The frequency of the circular junction sites dropped by around 50% 100 days after the nucleofection compared to the cell state at day seven. However, the occurrence of the circular junction site almost doubled compared to the initial non-drug selected phase, and this frequency was about four times higher compared to the non-drug treated groups 100 days post nucleofection. Interestingly, the frequency of deletion and inversion did not change over time in the absence of Palbociclib. Still, both increased upon drug treatment, albeit to a lesser extent than the circular junction site frequency (**Figure 5D**). Our findings suggest that cells already possessing CDK4 harboring ecDNAs might increase copy number to adapt to the drug-induced pressure, while other cells adapt to these conditions using alternative mechanisms. Additionally, our observations align with previous hypotheses and findings, indicating that ecDNA is capable of more dynamic changes compared to those still linked to the chromosomal genome.

### Drug selection induces retention of CRISPR-C targeted multi-drug resistance gene region

In cancers, an elevated copy number of a gene itself cannot inherently enhance cell growth or survival unless subjected to additional selection pressure, such as for the *ABCB1* oncogene included in this study. The protein coded for by this gene is recognized for its involvement in conferring resistance to chemotherapeutic drugs in cancer cells and requires a substantial amount of energy to function (41). Therefore, cells are not inclined to maintain a high level of this protein unless under certain selection pressure. We hypothesize that without drug selection the CRISPR-C generated *ABCB1* ecDNAs would be eliminated from the cells, and would persist upon specific drug selection. To explore this, we induced the formation of three different sizes of CRISPR-C ecDNA carrying the *ABCB1* coding gene in the MCF7 human breast cancer cell line (**Supplementary Figure S5**) and invested their persistence.

After five days of the CRISPR-C procedure, each of the generated groups was treated with colcemid (also known as demecolcine), which is a cytostatic agent and a known substrate of the *ABCB1* protein (42,43), for a period of 60 days (**Figure 6a**). We monitored the presence of generated deletion sites and circular junction sites in both treated and untreated cells using PCR assays (**Figure 6b**). In all three edited groups, both the deletion scars and circular junction sites were detectable two weeks after CRISPR-C induction in both the treated and untreated groups. The circular junction sites of all three ecDNAs disappeared two months after nucleofection. However, the circular junction site of the 244 Kb ecDNA remained detectable in cells treated with colcemid as confirmed by PCRs and FISH experiments (**Figure 6b-c, Supplementary Figure S28**). Additionally, we observed that the deletion site was retained in all groups except the 244 Kb ecDNA in the absence of colcemid treatment. This finding suggests that drug selection might promote the persistence of ecDNAs carrying drug-esistance genes in cancer cells.

**Figure 6.**
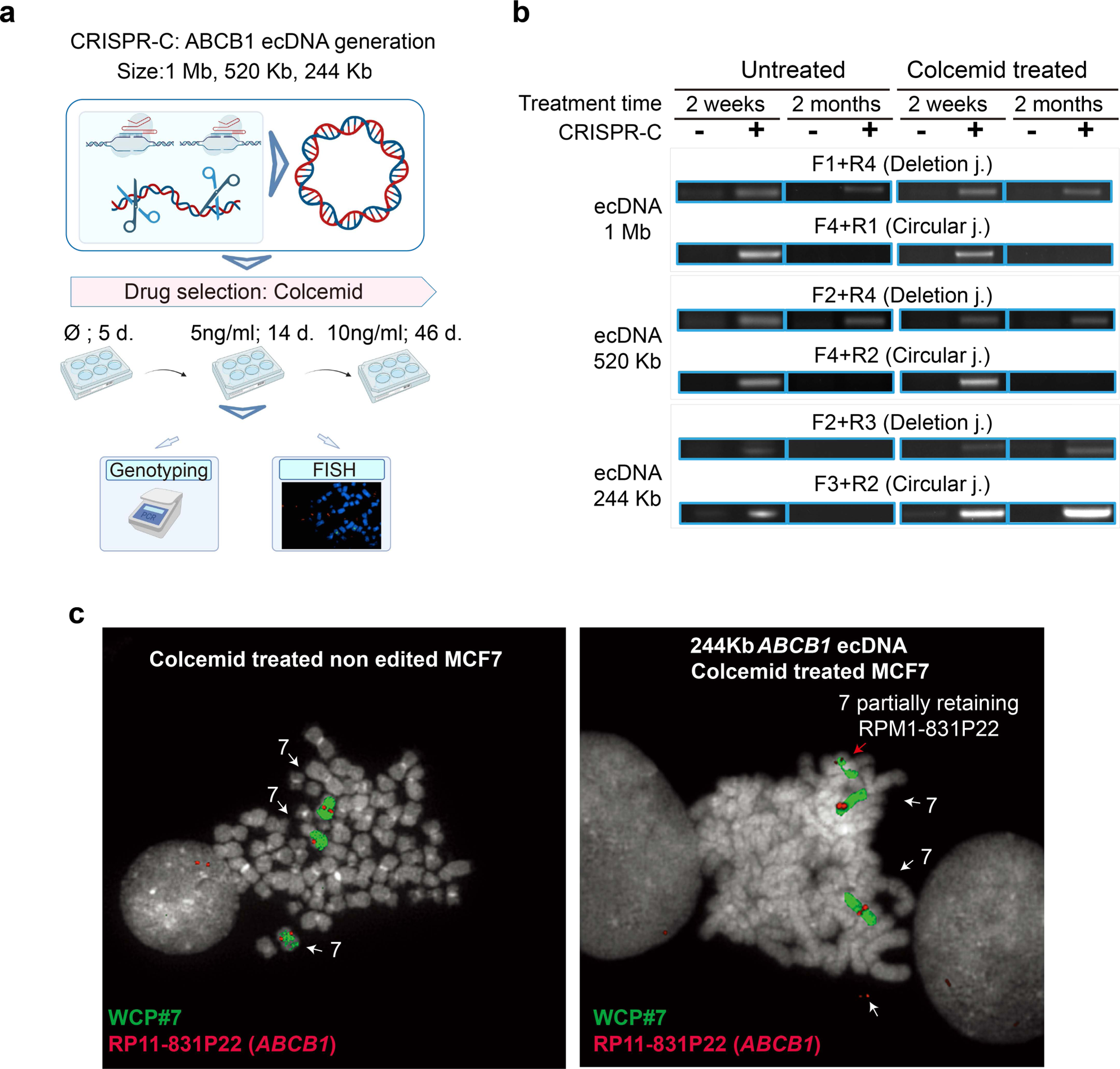
In vitro modelling of drug selection dynamics after *ABCB1* carrying ecDNA generation by CRISPR-C. **(a)** Workflow of the assessment of drug-selection effect on CRISPR-C generated ABCB1 ecDNA persistence. **(b)** PCR validation of the presence/absence of circular junction site and chromosomal scar after CRISPR-C 2.0 generation of ecDNAs of three different sizes. The results of PCR detection from non-treated (left side) or colcemid treated (right side) cells are shown. **(c)** Metaphase FISH analysis of *ABCB1* in MCF7 treated with colcemid, as indicated in the A panel. Red: signal of the RP11 encompassing the *ABCB1* gene, Green: WCP probe for chromosome 7. DAPI: Grey. Left panel: non-edited cells showing three red signals for *ABCB1*. Right panel: deletion of one *ABCB1* signal on one chromosome 7 and an extrachromosomal red signal upon induction of the CRISPR-C *ABCB1* ecDNA of 244 Kb.

## DISCUSSION

Here, we present here an improved version of CRISPR-C to recapitulate the generation of oncogene-carrying ecDNA in human cell lines. Compared to our previous version of CRISPR-C, the improved version (2.0) takes the advantages of *in silico* CRISPR design with deep-learning methods (25), chemical modification of gRNAs with enhanced intracellular stability (44), and efficient delivery of CRISPR-C components (Cas9 protein and sgRNAs). This indeed has substantially improved its efficiency. The efficiency CRISPR-C 2.0 was increased by seven folds over CRISPR-C 1.0 when applied in creating small extrachromosomal circular DNA (EGFP circle, 2 Kb) in cells. Although we did not perform pair-wise comparison of the generation of oncogene ecDNAs (up to 1 Mb) between the two versions of CRISPR-C, numerous results on CRISPR gene editing have consistently demonstrated that ribonucleoprotein (RNP) is the most efficient approach for *in vitro* CRISPR delivery (24). CRISPR-C 2.0 is a highly conventional and feasible approach for generating eccDNAs (small extrachromosomal circular DNA) and ecDNAs in cells (45). Although 60% of the cells were positive for the eccDNA (EGFP) with CRISPR-C 2.0, the efficiency of generating oncogene-harboring ecDNA (*CDK4*) was estimated at approximately 2% with ddPCR and FISH analysis, which highlights the need for further efficiency improvement. The genomic context plays an essential role in the efficiency of ecDNA generation with CRISPR-C. Rose et al. have recently explored the utilization of the 3D genome architecture but found no significant effect (46). Instead, CRISPR sgRNAs, genomic context (locus heterogeneity), and double-strand DNA break repair mechanisms play important roles in the efficiency of CRISPR-C-based ecDNA generation. Inhibition of one of the DSB repair protein DNA-PK has been demonstrated to promote ecDNA generation (46), promises further improvements of CRISPR-C by hijacking the DSB repair machineries in cells.

The ecDNA is a commonly detected genetic element in both normal and diseased (e.g., tumor) tissues. While most of them are detected across the entire genome, certain genomic regions have been found to be more vulnerable, such as gene-rich areas (3,47). In tumors, oncogene-harboring ecDNAs are commonly found to be positively selected and associated with tumor progression and evolution of resistance. Several mechanisms have been discovered for the association of ecDNA and tumor fitness, such as oncogene amplification and enhanced gene expression (1,9), enhancer hijacking (17), chromatin decompaction (14), genomic instability and rearrangement (8). We studied the effect of *CKD4* and *MDM2* CRISPR-C on gene expression in cell lines, but did not detect increased expression. However, this is not surprising due to the low circularization efficiency, and most importantly, the CRISPR-C is associated with gene deletion (knockouts) from the chromosome. An interesting finding is the transient increase of *MDM2* expression following the generation of *CKD4::MDM2* chimeric ecDNAs. Although we did not characterize the genomic and epigenomic architectures of the *CKD4::MDM2* chimeric ecDNAs and the resulting chromosomal scares in cells, our results provide a promising experimental strategy for exploring the evolution of ecDNA architectures in future studies.

Due to random/asymmetric segregation, oncogene-harboring ecDNAs have been thought to be associated with the rapid amplification of oncogenes in tumors (1,45). Our results suggest that ecDNA stability in tumor cells is more context-dependent. Under normal *in vitro* growth conditions, the CRISPR-C-generated *CDK4* and *MDM2* ecDNAs are stably maintained in cells. However, the *ABCB1*-harboring ecDNA is negatively selected. The addition of selection stresses, such as the *CDK4* inhibitor palbociclib and the cytostatic agent colcemid, can positively select cells with *CDK4* and *ABCB1* ecDNAs respectively. Interestingly, two out of three types of *ABCB1* ecDNA disappeared from the cells even under drug selection, suggesting a potential mechanism of context-dependent ecDNA stability. Previous studies have proposed the existence of adaptive mechanisms that increase the expression of the *ABCB1* gene without changes in the copy number, which may have also played a role in our cells’ adaptation to drug selection and reduced the importance of the generated *ABCB1*-harboring ecDNAs (48). It should be noted that cell lines used in the current study are derived from patient tumors. Therefore, the effect of the CRISPR-C generated ecDNAs on cell fitness could be affected by other genomic and chromosomal alterations. Since we have demonstrated that oncogene-harboring ecDNAs can be generated in normal cells (fibroblasts), we expect that the strategy will be exploited for *in vitro* (cultured cells) and even *ex vivo* (such as organoids)/*in vivo* (animal models) transformation of normal cells through oncogene circularization.

In conclusion, we present the development of an improved CRISPR-C for modeling the generation of oncogene-carrying ecDNA in human cells. Our method, allowing the genesis of ecDNAs from specific chromosomal regions also deriving from different chromosomes, can serve as a powerful molecular tool in studying the role, biogenesis, persistence, and increase in copies of circular DNA in cancer evolution.

## Supporting information

Source data for gel images in both figures and supplementary figures

## CONFLICT OF INTEREST

Authors declare no conflict of interest.

## AUTHOR CONTRIBUTION

Conceptualization, Y.L.; Methodology, J.H., W.F., A.A., Y.L.; Validation, W.F., A.A., C.S.H., T.S.P., D.T., W.L., P.H., Y.Z., F.W.,; Formal analysis, J.H., W.F., A.A., C.T.S., D.T.; Investigation, J.H., W.F., A.A., C.S.H., T.S.P., D.T., W.L., P.H., Y.Z., F.W.,; Resources, L.B., L.L., C.T.S., B.R., Y.L.; Writing—original draft preparation, J.H., Y.L.; Writing—review and editing, J.H., W.F., A.A., C.S.H., T.S.P., D.T., W.L., P.H., Y.Z., F.W., L.B., L.L., C.T.S., B.R., Y.L.; Visualization, J.H., W.F., A.A.; Supervision, C.T.S., B.R., Y.L.; Project administration, Y.L.; Funding acquisition, B.R., Y.L., C.T.S.; All authors have read and agreed to the published version of the manuscript.

## ACKNOWLEDGEMENT

We thank Roberta May Fisher for critical comments and language editing provided to this manuscript. This project is supported by the European Union’s Horizon 2020 research and innovation program under grant agreement No. 899417. Y.L. is supported by Danish Research Council (9041-00317B), European Union’s Horizon 2020 research and innovation program under grant agreement No 899417 (Y.L.), the Novo Nordisk Foundation (NNF21OC0068988; NNF21OC0071031), M-ERA.Net and the Innovation Fund Denmark (9355 PIECRISCI), and the Lundbeck Foundation Ascending Investigator award (R396-2022-350). C.T.S. is supported by the AIRC (Associazione Italiana per la Ricerca sul Cancro), AIRC IG no. 25706.

## Notes

### Competing Interest Statement

The authors have declared no competing interest.

## REFERENCES

1. Turner, K.M., Deshpande, V., Beyter, D., Koga, T., Rusert, J., Lee, C., Li, B., Arden, K., Ren, B., Nathanson, D.A. et al. (2017) Extrachromosomal oncogene amplification drives tumour evolution and genetic heterogeneity. Nature, 543, 122-+.

2. Shoshani, O., Brunner, S.F., Yaeger, R., Ly, P., Nechemia-Arbely, Y., Kim, D.H., Fang, R.X., CasPllon, G.A., Yu, M., Li, J.S.Z. et al. (2021) Chromothripsis drives the evoluPon of gene amplificaPon in cancer (vol 591, pg 137, 2021). Nature, 591, E19–E19.

3. Shibata, Y., Kumar, P., Layer, R., Willcox, S., Gagan, J.R., Griffith, J. D. and Dutta, A. (2012) Extrachromosomal microDNAs and chromosomal microdeletions in normal tissues. Science, 336, 82–86.

4. Moller, H.D., Mohiyuddin, M., Prada-Luengo, I., Sailani, M.R., Halling, J.F., Plomgaard, P., Maretty, L., Hansen, A.J., Snyder, M.P., Pilegaard, H. et al. (2018) Circular DNA elements of chromosomal origin are common in healthy human somatic tissue. Nature Communications, 9.

5. Hung, K.L., Luebeck, J., Dehkordi, S.R., Colon, C.I., Li, R., Tsz-Lo Wong, I., Coruh, C., Dharanipragada, P., Lomeli, S.H., Weiser, N.E. et al. (2022) Targeted profiling of human extrachromosomal DNA by CRISPR-CATCH. Nat Genet, 54, 1746-+.

6. Yi, E., Gujar, A.D., Guthrie, M., Kim, H., Zhao, D., Johnson, K.C., Amin, S.B., Costa, M.L., Yu, Q.R., Das, S. et al. (2022) Live-Cell Imaging Shows Uneven Segregation of Extrachromosomal DNA Elements and Transcriptionally Active Extrachromosomal DNA Hubs in Cancer. Cancer Discov, 12, 468–483.

7. deCarvalho, A.C., Kim, H., Poisson, L.M., Winn, M.E., Mueller, C., Cherba, D., Koeman, J., Seth, S., Protopopov, A., Felicella, M., et al. (2018) Discordant inheritance of chromosomal and extrachromosomal DNA elements contributes to dynamic disease evolution in glioblastoma. Nat Genet, 50, 708-+.

8. Koche, R.P., Rodriguez-Fos, E., Helmsauer, K., Burkert, M., MacArthur, I.C., Maag, J., Chamorro, R., Munoz-Perez, N., Puiggros, M., Dorado Garcia, H. et al. (2020) Extrachromosomal circular DNA drives oncogenic genome remodeling in neuroblastoma. Nat Genet, 52, 29–34.

9. Kim, H., Nguyen, N.P., Turner, K., Wu, S.H., Gujar, A.D., Luebeck, J., Liu, J.H., Deshpande, V., Rajkumar, U., Namburi, S. et al. (2020) Extrachromosomal DNA is associated with oncogene amplification and poor outcome across multiple cancers. Nat Genet, 52, 891-+.

10. Hung, K.L., Yost, K.E., Xie, L., Shi, Q., Helmsauer, K., Luebeck, J., Schopflin, R., Lange, J.T., Chamorro Gonzalez, R., Weiser, N.E., et al. (2021) ecDNA hubs drive cooperative intermolecular oncogene expression. Nature, 600, 731–736.

11. Helmsauer, K., Valieva, M.E., Ali, S., Chamorro Gonzalez, R., Schopflin, R., Roefzaad, C., Bei, Y., Dorado Garcia, H., Rodriguez-Fos, E., Puiggros, M., et al. (2020) Enhancer hijacking determines extrachromosomal circular MYCN amplicon architecture in neuroblastoma. Nat Commun, 11, 5823.

12. Schoenlein, P.V., Shen, D.W., Barrett, J.T., Pastan, I. and Gottesman, M.M. (1992) Double Minute Chromosomes Carrying the Human Multidrug Resistance-1 and Resistance-2 Genes Are Generated from the Dimerization of Submicroscopic Circular Dnas in Colchicine-Selected Kb Carcinoma-Cells. Mol Biol Cell, 3, 507–520.

13. Singer, M.J., Mesner, L.D., Friedman, C.L., Trask, B.J. and Hamlin, J.L. (2000) Amplification of the human dihydrofolate reductase gene via double minutes is initiated by chromosome breaks. P Natl Acad Sci USA, 97, 7921–7926.

14. Wu, S., Turner, K.M., Nguyen, N., Raviram, R., Erb, M., Santini, J., Luebeck, J., Rajkumar, U., Diao, Y., Li, B. et al. (2019) Circular ecDNA promotes accessible chromatin and high oncogene expression. Nature, 575, 699–703.

15. Hung, K.L., Luebeck, J., Dehkordi, S.R., Colon, C.I., Li, R., Wong, I.T., Coruh, C., Dharanipragada, P., Lomeli, S.H., Weiser, N.E. et al. (2022) Targeted profiling of human extrachromosomal DNA by CRISPR-CATCH. Nat Genet, 54, 1746–1754.

16. Zhu, Y., Gujar, A.D., Wong, C.H., Tjong, H., Ngan, C.Y., Gong, L., Chen, Y.A., Kim, H., Liu, J., Li, M. et al. (2021) Oncogenic extrachromosomal DNA functions as mobile enhancers to globally amplify chromosomal transcription. Cancer Cell, 39, 694–707 e697.

17. Morton, A.R., Dogan-Artun, N., Faber, Z.J., MacLeod, G., Bartels, C.F., Piazza, M.S., Allan, K.C., Mack, S.C., Wang, X., Gimple, R.C. et al. (2019) Functional Enhancers Shape Extrachromosomal Oncogene Amplifications. Cell, 179, 1330–1341 e1313.

18. Hung, K.L., Yost, K.E., Xie, L.Q., Shi, Q.M., Helmsauer, K., Luebeck, J., Schopflin, R., Lange, J.T., Gonzalez, R.C., Weiser, N.E., et al. (2021) ecDNA hubs drive cooperative intermolecular oncogene expression. Nature, 600, 731-+.

19. Lange, J.T., Rose, J.C., Chen, C.Y., Pichugin, Y., Xie, L., Tang, J., Hung, K.L., Yost, K.E., Shi, Q., Erb, M.L. et al. (2022) The evolutionary dynamics of extrachromosomal DNA in human cancers. Nat Genet, 54, 1527–1533.

20. Moller, H.D., Lin, L., Xiang, X., Petersen, T.S., Huang, J.R., Yang, L.H., Kjeldsen, E., Jensen, U.B., Zhang, X.Q., Liu, X. et al. (2018) CRISPR-C: circularization of genes and chromosome by CRISPR in human cells. Nucleic Acids Res, 46.

21. Conant, D., Hsiau, T., Rossi, N., Oki, J., Maures, T., Waite, K., Yang, J., Joshi, S., Kelso, R., Holden, K. et al. (2022) Inference of CRISPR Edits from Sanger Trace Data. CRISPR J, 5, 123–130.

22. Storlazzi, C.T., Von Steyern, F.V., Domanski, H.A., Mandahl, N. and Mertens, F. (2005) Biallelic somatic inactivation of the NF1 gene through chromosomal translocations in a sporadic neurofibroma. Int J Cancer, 117, 1055–1057.

23. Ramakrishna, S., Kwaku Dad, A.B., Beloor, J., Gopalappa, R., Lee, S.K. and Kim, H. (2014) Gene disruption by cell-penetrating peptide-mediated delivery of Cas9 protein and guide RNA. Genome Res, 24, 1020–1027.

24. Zhang, S., Shen, J., Li, D. and Cheng, Y. (2021) Strategies in the delivery of Cas9 ribonucleoprotein for CRISPR/Cas9 genome editing. Theranostics, 11, 614–648.

25. Xiang, X., Corsi, G.I., Anthon, C., Qu, K., Pan, X., Liang, X., Han, P., Dong, Z., Liu, L., Zhong, J. et al. (2021) Enhancing CRISPR-Cas9 gRNA efficiency prediction by data integration and deep learning. Nat Commun, 12, 3238.

26. Xiang, X., Zhao, X., Pan, X., Dong, Z., Yu, J., Li, S., Liang, X., Han, P., Qu, K., Jensen, J.B. et al. (2021) Efficient correction of Duchenne muscular dystrophy mutations by SpCas9 and dual gRNAs. Mol Ther Nucleic Acids, 24, 403–415.

27. Stinson, B.M., Moreno, A.T., Walter, J.C. and Loparo, J.J. (2020) A Mechanism to Minimize Errors during Non-homologous End Joining. Mol Cell, 77, 1080–1091 e1088.

28. Chakraborty, A., Tapryal, N., Venkova, T., Horikoshi, N., Pandita, R.K., Sarker, A.H., Sarkar, P.S., Pandita, T.K. and Hazra, T.K. (2016) Classical non-homologous end-joining pathway utilizes nascent RNA for error-free double-strand break repair of transcribed genes. Nat Commun, 7, 13049.

29. Song, B., Yang, S., Hwang, G.H., Yu, J. and Bae, S. (2021) Analysis of NHEJ-Based DNA Repair aler CRISPR-Mediated DNA Cleavage. Int J Mol Sci, 22.

30. Caracciolo, D., Riillo, C., Di Martino, M.T., Tagliaferri, P. and Tassone, P. (2021) Alternative Non-Homologous End-Joining: Error-Prone DNA Repair as Cancer’s Achilles’ Heel. Cancers (Basel), 13.

31. Sanchez, A.M., Barrett, J.T. and Schoenlein, P.V. (1998) Fractionated ionizing radiation accelerates loss of amplified MDR1 genes harbored by extrachromosomal DNA in tumor cells. Cancer Res, 58, 3845–3854.

32. Schoenlein, P.V., Shen, D.W., Barrett, J.T., Pastan, I. and Gottesman, M.M. (1992) Double minute chromosomes carrying the human mulPdrug resistance 1 and 2 genes are generated from the dimerization of submicroscopic circular DNAs in colchicine-selected KB carcinoma cells. Mol Biol Cell, 3, 507–520.

33. Reed, K., Hembruff, S.L., Laberge, M.L., Villeneuve, D.J., Cote, G.B. and Parissenti, A.M. (2008) Hypermethylation of the ABCB1 downstream gene promoter accompanies ABCB1 gene amplification and increased expression in docetaxel-resistant MCF-7 breast tumor cells. Epigenetics, 3, 270–280.

34. Purshouse, K., Friman, E.T., Boyle, S., Dewari, P.S., Grant, V., Hamdan, A., Morrison, G.M., Brennan, P.M., Beentjes, S.V., Pollard, S.M. et al. (2022) Oncogene expression from extrachromosomal DNA is driven by copy number amplification and does not require spatial clustering in glioblastoma stem cells. Elife, 11.

35. L’Abbate, A., Tolomeo, D., Cifola, I., Severgnini, M., Turchiano, A., Augello, B., Squeo, G., D’Addabbo, P., Traversa, D., Daniele, G., et al. (2018) Correction: MYC-containing amplicons in acute myeloid leukemia: genomic structures, evolution, and transcriptional consequences. Leukemia, 32, 2304.

36. Luebeck, J., Coruh, C., Dehkordi, S.R., Lange, J.T., Turner, K.M., Deshpande, V., Pai, D.A., Zhang, C., Rajkumar, U., Law, J.A. et al. (2020) AmpliconReconstructor integrates NGS and optical mapping to resolve the complex structures of focal amplifications. Nat Commun, 11, 4374.

37. Amoroso, L., Ognibene, M., Morini, M., Conte, M., Di Cataldo, A., Tondo, A., D’Angelo, P., Castellano, A., Garaventa, A., Lasorsa, V.A., et al. (2020) Genomic coamplification of CDK4/MDM2/FRS2 is associated with very poor prognosis and atypical clinical features in neuroblastoma patients. Genes Chromosomes Cancer, 59, 277–285.

38. Walentynowicz, K.A., Engelhardt, D., Cristea, S., Yadav, S., Onubogu, U., Salatino, R., Maerken, M., Vincentelli, C., Jhaveri, A., Geisberg, J. et al. (2023) Single-cell heterogeneity of EGFR and CDK4 co-amplification is linked to immune infiltration in glioblastoma. Cell Rep, 42, 112235.

39. Chang, C.M. and Lam, H.Y.P. (2023) Mechanism of CDK4/6 Inhibitor Resistance in Hormone Receptor-positive Breast Cancer and Alternative Treatment Strategies. Anticancer Res, 43, 5283–5298.

40. Olanich, M.E., Sun, W., Hewitt, S.M., Abdullaev, Z., Pack, S.D. and Barr, F.G. (2015) CDK4 Amplification Reduces Sensitivity to CDK4/6 Inhibition in Fusion-Positive Rhabdomyosarcoma. Clin Cancer Res, 21, 4947–4959.

41. Skinner, K.T., Palkar, A.M. and Hong, A.L. (2023) Genetics of ABCB1 in Cancer. Cancers (Basel), 15.

42. Loo, T.W., Bartlett, M.C. and Clarke, D.M. (2003) Substrate-induced conformational changes in the transmembrane segments of human P-glycoprotein. Direct evidence for the substrate-induced fit mechanism for drug binding. J Biol Chem, 278, 13603–13606.

43. Loo, T.W., Bartlett, M.C. and Clarke, D.M. (2003) Drug binding in human P-glycoprotein causes conformational changes in both nucleotide-binding domains. J Biol Chem, 278, 1575–1578.

44. Hendel, A., Bak, R.O., Clark, J.T., Kennedy, A.B., Ryan, D.E., Roy, S., Steinfeld, I., Lunstad, B.D., Kaiser, R.J., Wilkens, A.B. et al. (2015) Chemically modified guide RNAs enhance CRISPR-Cas genome editing in human primary cells. Nat Biotechnol, 33, 985–989.

45. Noer, J.B., Horsdal, O.K., Xiang, X., Luo, Y. and Regenberg, B. (2022) Extrachromosomal circular DNA in cancer: history, current knowledge, and methods. Trends Genet, 38, 766–781.

46. Rose, J.C., Wong, I.T.-L., Daniel, B., Jones, M.G., Yost, K.E., Hung, K.L., Curtis, E.J., Mischel, P.S. and Chang, H.Y. (2023) Disparate pathways for extrachromosomal DNA biogenesis and genomic DNA repair. bioRxiv, 2023.2010.2022.563489.

47. Dillon, L.W., Kumar, P., Shibata, Y., Wang, Y.H., Willcox, S., Griffth, J.D., Pommier, Y., Takeda, S. and Dutta, A. (2015) Production of Extrachromosomal MicroDNAs Is Linked to Mismatch Repair Pathways and Transcriptional Activity. Cell Rep, 11, 1749–1759.

48. Calcagno, A.M. and Ambudkar, S.V. (2010) Molecular mechanisms of drug resistance in single-step and multi-step drug-selected cancer cells. Methods Mol Biol, 596, 77–93.

