## Supplementary Figures for "Generative Modelling of Oncogene-carrying Extrachromosomal Circular DNA Biogenesis and Dynamics in Cells"

#### SUPPLEMENTARY FIGURE LEGENDS

**Supplementary Figure S1. RNP-based eccDNA generation is highly efficient compared to plasmid-based system in eccDNA biosensor line.** (a) Graphical description of TRE-ECC dual fluorescent eccDNA biosensor system after CRISPR-C induced two double-strand breaks resulting in circularization of the cleaved fragment leading to EGFP expression or inversion leading to mCherry expression. TRE-ECC biosensor cassette was inserted into the genome of HEK293T cells as a single copy. (b) Timeline of plasmid- (CRISPR-C 1.0) vs RNP-based (CRISPR-C 2.0) eccDNA induction and assessment by flowcytometry. (c) Comparison of eccDNA and inversion generation efficiency of CRISPR-C 1.0 and CRISPR-C 2.0 in HEK293T TRE-ECC cell line 48 h after CRISPR-C induction by flowcytometry (fluorescent cell quantity),*** represent p values less than 0.001. (d) Dynamics of fluorescent cell quantity (±SD) and median fluorescent intensity of GFP and mCherry positive cells over time after plasmid vs RNP based CRISPR-C.

**Supplementary Figure S2.** Schematic drawing of the location of recurrent ecDNA carried oncogens in humans.

**Supplementary Figure S3. Genotyping of CRISPR-C generated *MDM2* carrying ecDNA.** (a) Graphical illustration of CRISPR-C to generate different sizes (1Mb, 587 Kb and 192 Kb) of ecDNAs carrying the *MDM2* oncogene. Validation of 1Mb - chr12 $^{68,371,015-69,389,817}$ (b), 578 Kb - chr12 $^{68,371,015-68,958,626}$ (PCR gel is identical to the one represented at Figure 1 c) (c) and 192 Kb - chr12 $^{68,766,906-68,958,645}$ (d) size ecDNA deletion scar and circularization ligation site by PCR and sanger sequencing. White asterisk indicates the sequenced PCR products.

**Supplementary Figure S4. Genotyping of CRISPR-C generated *CDK4* carrying ecDNA.** (a) Graphical illustration of CRISPR-C to generate different sizes (1Mb, 508 Kb and 201 Kb) of ecDNAs carrying the *CDK4* oncogene. Validation of 1Mb – chr12 $^{57,613,237-58,609,860}$ (b), 508Kb – chr12 $^{57,613,237-58,121,411}$ (c) and 201 Kb – chr12 $^{57,613,237-57,814,432}$ (d) size ecDNA deletion scar and circularization ligation site by PCR and sanger sequencing. White asterisk indicates the sequenced PCR products.

**Supplementary Figure S5. Genotyping of CRISPR-C generated *ABCB1* carrying ecDNA.** (a) Graphical illustration of CRISPR-C to generate different sizes (1Mb, 520 Kb, 244 Kb) of ecDNAs carrying the *ABCB1* oncogene. Validation of 1Mb chr7 $^{87,217,400-88,262,995}$ (b), 520 Kb chr7 $^{87,217,400-87,737,965}$ (c) and 244 Kb chr7 $^{87,493076-87,737,965}$ (d) size ecDNA deletion scar and circularization ligation site by PCR and sanger sequencing. White asterisk indicates the sequenced PCR products.

**Supplementary Figure S6. Genotyping of CRISPR-C generated *MYC* carrying ecDNA.** (a) Graphical illustration of CRISPR-C to generate different sizes (1Mb, 600 Kb and 100 Kb) of ecDNAs carrying the *MYC* oncogene. Validation of 1Mb chr8 $^{127,250,015\text{-}128,249,709}$ (b), 600 Kb chr8 $^{127,250,015\text{-}127,849,549}$ (c) and 100 Kb chr8$^{127,649,837\text{-}127,749,332}$ (d) size ecDNA deletion scar and circularization ligation site by PCR and sanger sequencing. White asterisk indicates the sequenced PCR products.

**Supplementary Figure S7. Genotyping of CRISPR-C generated *MYCN* carrying ecDNA.** (a) Graphical illustration of CRISPR-C to generate different sizes (1Mb, 505 Kb and 239 Kb) of ecDNAs carrying the *MYCN* oncogene. Validation of 1Mb chr2 $^{15,333,415\text{-}16,332,075}$ (b), 535 Kb chr2 $^{15,749,854\text{-}16,254,983}$ (c) and 239 Kb chr2 $^{15,890,833\text{-}16,129,387}$ (d) size ecDNA deletion scar and circularization ligation site by PCR and sanger sequencing. White asterisk indicates the sequenced PCR products.

**Supplementary Figure S8. Genotyping of CRISPR-C generated *FGFR2*carrying ecDNA.** (a) Graphical illustration of CRISPR-C to generate different sizes (1Mb, 537 Kb and 173 Kb) of ecDNAs carrying the *FGFR2* oncogene. Validation of 1Mb chr10 $^{120,997,077\text{-} 121,982,362}$ (b), 537 Kb chr10 $^{121,445,321\text{-} 121,982,381}$ (c) and 173 Kb chr10 $^{121,445,321\text{-}121,618,535}$ (d) size ecDNA deletion scar and circularization ligation site by PCR and sanger sequencing. White asterisk indicates the sequenced PCR products.

**Supplementary Figure S9. Genotyping of CRISPR-C generated *EGFR* carrying ecDNA.** (a) Graphical illustration of CRISPR-C to generate 668 Kb size ecDNAs carrying the *EGFR* oncogene. Validation of 668Kb EGFR chr2 $^{54,589,038\text{-} 55,257,084}$ ecDNA deletion scar and circularization ligation site by PCR (b) and sanger sequencing (c) . White asterisk indicates the sequenced PCR products.

**Supplementary Figure S10. Genotyping of CRISPR-C generated chimeric *CDK4* and *MDM2* carrying ecDNA with intrachromosomal origin.** (a) Graphical illustration of CRISPR-C generation of DNA fragments (508Kb chr12: $^{57,613,237\text{-}58,121,411}$ and 587 Kb chr12 $^{68,371,015\text{-}68,958,626}$ ) carrying the *CDK4* or the *MDM2* genes both derived from chromosome 12, where the two DNA fragments can ligate and circularize in two possible ways. (b, c) PCR and Sanger sequencing validation of the 2X2 possible ligation sites of the chimeric circles (PCR gel is identical to the one represented at Figure 1 e).

**Supplementary Figure S11. Genotyping of CRISPR-C generated chimeric *CDK4* and *EGFR* carrying 1.2 Mb size ecDNA with interchromosomal origin.** (a) Graphical illustration of CRISPR-C generation of DNA fragments (508Kb chr12 $^{57,613,237\text{-}58,121,411}$ and 668 Kb chr2 $^{54,589,038\text{-} 55,257,084}$) carrying the *CDK4* or the *EGFR* genes both derived from chromosome 12 and chromosome 7, where the two DNA fragment can ligate and circularize in two possible ways. PCR primers are labelled as black arrows. (b, c) PCR and Sanger sequencing validation of the 2X2 possible ligation sites of the chimeric circles.

**Supplementary Figure S12. Genotyping of CRISPR-C generated chimeric *CDK4* and *EGFR* carrying 869 Kb size ecDNA with inter-chromosomal origin.** (a) Graphical illustration of CRISPR-C generation of DNA fragments (201 Kb, chr12$^{57,613,237\text{-}57,814,432}$ and 668 Kb, chr2 $^{54,589,038\text{-}\ 55,257,084}$) carrying the *CDK4* or the *EGFR* genes both derived from chromosome 12 and chromosome 7, where the two DNA fragments can ligate and circularize in two possible ways. PCR primers are labelled as black arrows. (b, c) PCR and Sanger sequencing validation of the 2X2 possible ligation sites of the chimeric circles.

**Supplementary Figure S13. ICE analysis of *ABCB1* carrying CRISPR-C ecDNA generation sgRNAs.** (a) Sanger sequence display depicting both edited and wild-type (control) sequences within the vicinity of the guide sequence. (b) Indel plot displaying the inferred distribution of indels in the edited set of cells by each sgRNAs.

**Supplementary Figure S14. ICE analysis of *CDK4* carrying CRISPR-C ecDNA generation sgRNAs.** (a) Sanger sequence display depicting both edited and wild-type (control) sequences within the vicinity of the guide sequence. (b) Indel plot displaying the inferred distribution of indels in the edited set of cells by each sgRNAs.

**Supplementary Figure S15. ICE analysis of *DHFR* carrying CRISPR-C ecDNA generation sgRNAs.** (a) Sanger sequence display depicting both edited and wild-type (control) sequences within the vicinity of the guide sequence. (b) Indel plot displaying the inferred distribution of indels in the edited set of cells by each sgRNAs

**Supplementary Figure S16. ICE analysis of *EGFR* carrying CRISPR-C ecDNA generation sgRNAs.** (a) Sanger sequence display depicting both edited and wild-type (control) sequences within the vicinity of the guide sequence. (b) Indel plot displaying the inferred distribution of indels in the edited set of cells by each sgRNAs.

**Supplementary Figure S17. ICE analysis of *FGFR2* carrying CRISPR-C ecDNA generation sgRNAs.** (a) Sanger sequence display depicting both edited and wild-type (control) sequences within the vicinity of the guide sequence. (b) Indel plot displaying the inferred distribution of indels in the edited set of cells by each sgRNAs.

**Supplementary Figure S18. ICE analysis of *MDM2* carrying CRISPR-C ecDNA generation sgRNAs.** (a) Sanger sequence display depicting both edited and wild-type (control) sequences within the vicinity of the guide sequence. (b) Indel plot displaying the inferred distribution of indels in the edited set of cells by each sgRNAs.

**Supplementary Figure S19. ICE analysis of *MYC* carrying CRISPR-C ecDNA generation sgRNAs.** (a) Sanger sequence display depicting both edited and wild-type (control) sequences within the vicinity of the guide sequence. (b) Indel plot displaying the inferred distribution of indels in the edited set of cells by each sgRNAs.

**Supplementary Figure S20. ICE analysis of *MYCN* carrying CRISPR-C ecDNA generation sgRNAs.** (a) Sanger sequence display depicting both edited and wild-type (control) sequences within the vicinity of the guide sequence. (b) Indel plot displaying the inferred distribution of indels in the edited set of cells by each sgRNAs.

**Supplementary Figure S21. Fluorescent microscopy based investigation of the efficiency of electroporation based delivery.** 24 h after electroporation of control cells with eGFP IVT mRNA. White scalebar: 50 µm.

**Supplementary Figure S22. Genomic map of BAC probes and ecDNAs.** Genomic maps showing the relative positions of the BAC probes used in FISH experiments, and ecDNAs induced by CRISR-C 2.0, according to UCSC genome browser (GRCh38/hg38, https://genome.ucsc.edu/). The genomic regions harboring *CDK4* (a), *MDM2* (b), and *ABCB1* (c) genes are reported.

**Supplementary Figure S23. DNA-metaphase-FISH detection of *CDK4* region deletion in U2OS cells.** Grey: DAPI staining of DNA; Green: WCP of chromosome 12; Red and Blue: BAC clones RP11-846E20 and RP11-936I7 respectively for *CDK4*. Non-merged images of Figure 2 a.

**Supplementary Figure S24. DNA-metaphase-FISH detection of *MDM2* region deletion in U2OS cells.** Grey: DAPI staining of DNA; Green: WCP of chromosome 12; Red and Blue: BAC clones RP11-450G15 and RP11-1024C4 respectively for *MDM2*. Non-merged images of Figure 2 b.

**Supplementary Figure S25. DNA-metaphase-FISH detection of *CDK4* and *MDM2* region located on extrachromosomal copy or reinsertion and their deletion in U2OS cells.** Grey: DAPI staining of DNA; Green: WCP of chromosome 12; Red and Blue: BAC clones RP11-846E20 and RP11-936I7 respectively for *CDK4*, and RP11-450G15 and RP11-1024C4 respectively for *MDM2*. Non-merged images of Figure 2 c, d.

**Supplementary Figure S26. DNA-metaphase-FISH detection of *CDK4* region on extrachromosomal copy and its deletion in fibroblast cells.** Grey: DAPI staining of DNA; Green: WCP of chromosome 12; Red and Blue: BAC clones RP11-846E20 and RP11-936I7 respectively for *CDK4*, and RP11-450G15 and RP11-1024C4 respectively for *MDM2*. Non-merged and merged images of CRISPR unedited fibroblast cells and non-merged images of Figure 2 e.

**Supplementary Figure S27. DNA-metaphase-FISH detection of *CDK4* region on extrachromosomal copy and its deletion in U2OS cells after CRISPR-C followed by palbociclib treatment** as indicated in Figure 5. a. Grey: DAPI staining of DNA; Green: WCP of chromosome 12; Red and Blue: BAC clones RP11-846E20 and RP11-936I7 respectively for *CDK4*, and RP11-450G15 and RP11-1024C4 respectively for *MDM2*. Non-merged images of Figure 5 c

**Supplementary Figure S28. DNA-metaphase-FISH detection of *ABCB1* region on extrachromosomal copy and its deletion in MCF7 cells after CRISPR-C followed by colcemid treatment as indicated** on Figure 6.a. Red: signal of the RP11 encompassing the *ABCB1* gene, Green: WCP probe for chromosome 7. DAPI: Grey. Non-merged images of Figure 6 c

**Supplementary Figure S29.** Gene Set Enrichment Analysis (GSEA) of differentially expressed genes (DEGs) in a comparison between the first generation of control and chimeric ecDNA cell lines. The GSEA Enrichment Score curves of TNF signaling pathway (left) and Wnt signaling pathway (right) are shown here.

**Supplementary Figure S30.** Following persistence of deletion scar and circular ligation site of *CDK4* harboring 508 Kb CRISPR-C generated ecDNA by iPCR over 30 days.

**Supplementary Figure S31.** Following persistence of deletion scar and circular ligation site of *MDM2* harboring 587 Kb CRISPR-C generated ecDNA by iPCR over 30 days.

**Supplementary Figure S32.** Following persistence of circular ligation site of *CDK4* and *MDM2* harboring chimeric ecDNA harboring 1,1 Mb CRISPR-C generated ecDNA by PCR over 30 days.

**Supplementary Figure S33. ddPCR assessment of *CDK4* harboring 508 Kb ecDNA circularization, deletion scar formation and inversion formation frequency.** a. Graphical illustration of the ddPCR setup indicating the location of the site-specific primers and probes. b. representative plots of the obtained ddPCR data with indication of the analyzed clusters.

#### Figure S1

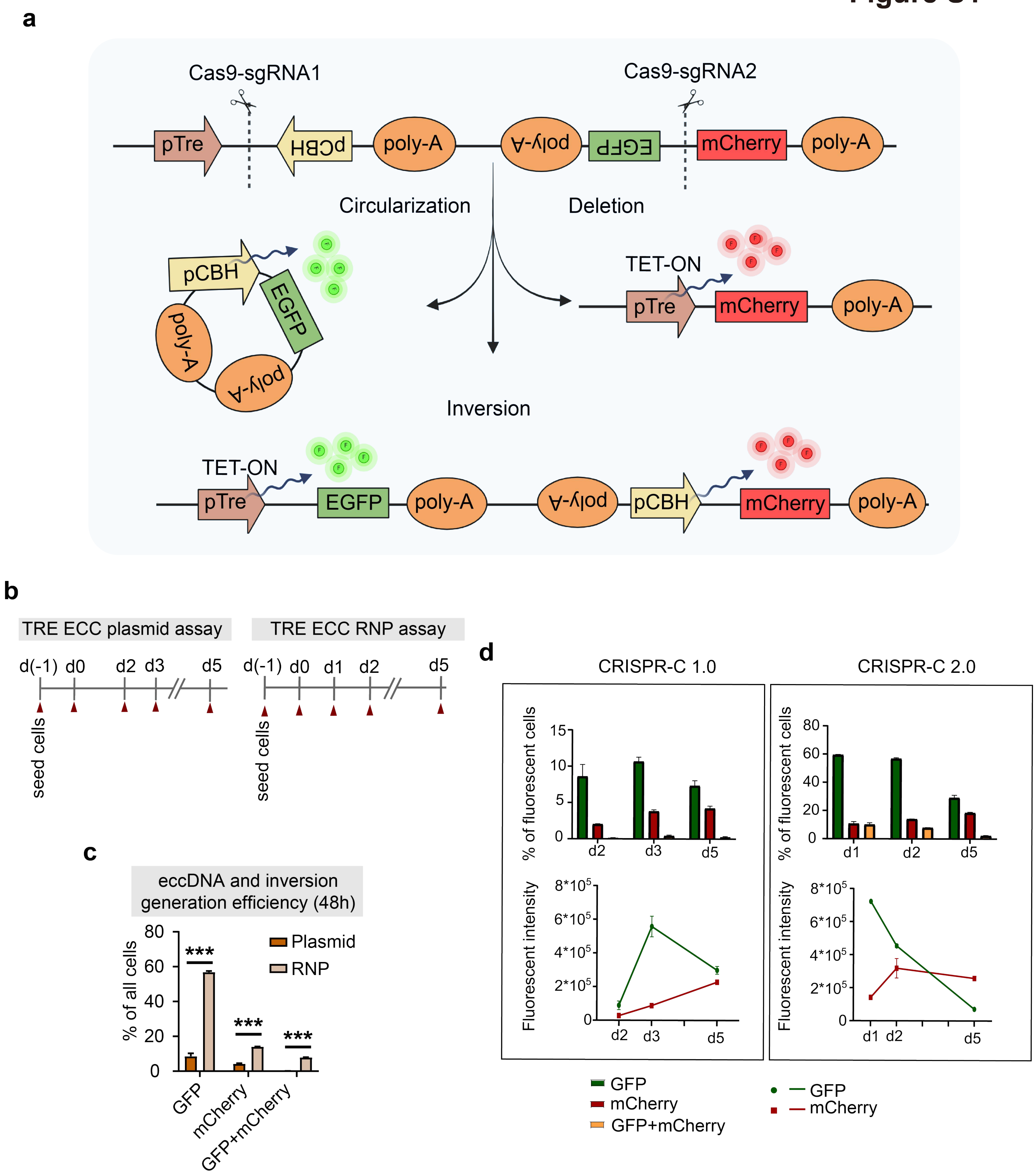
a
Cas9-sgRNA1
Cas9-sgRNA2
pTre
pCBH
poly-A
poly-A
EGFP
mCherry
poly-A
Circularization
Deletion
pCBH
EGFP
poly-A
poly-A
TET-ON
pTre
mCherry
poly-A
Inversion
TET-ON
pTre
EGFP
poly-A
poly-A
pCBH
mCherry
poly-A
b
TRE ECC plasmid assay
d(-1) d0 d2 d3 d5
seed cells
TRE ECC RNP assay
d(-1) d0 d1 d2 d5
seed cells
c
eccDNA and inversion generation efficiency (48h)
% of all cells
80
60
40
20
0
***
***
***
Plasmid
RNP
GFP
mCherry
GFP+mCherry
d
CRISPR-C 1.0
CRISPR-C 2.0
% of fluorescent cells
15
10
5
0
d2
d3
d5
% of fluorescent cells
80
60
40
20
0
d1
d2
d5
Fluorescent intensity
8*10^5
6*10^5
4*10^5
2*10^5
0
d2
d3
d5
Fluorescent intensity
8*10^5
6*10^5
4*10^5
2*10^5
0
d1
d2
d5
GFP
mCherry
GFP+mCherry
GFP
mCherry

### Figure S2

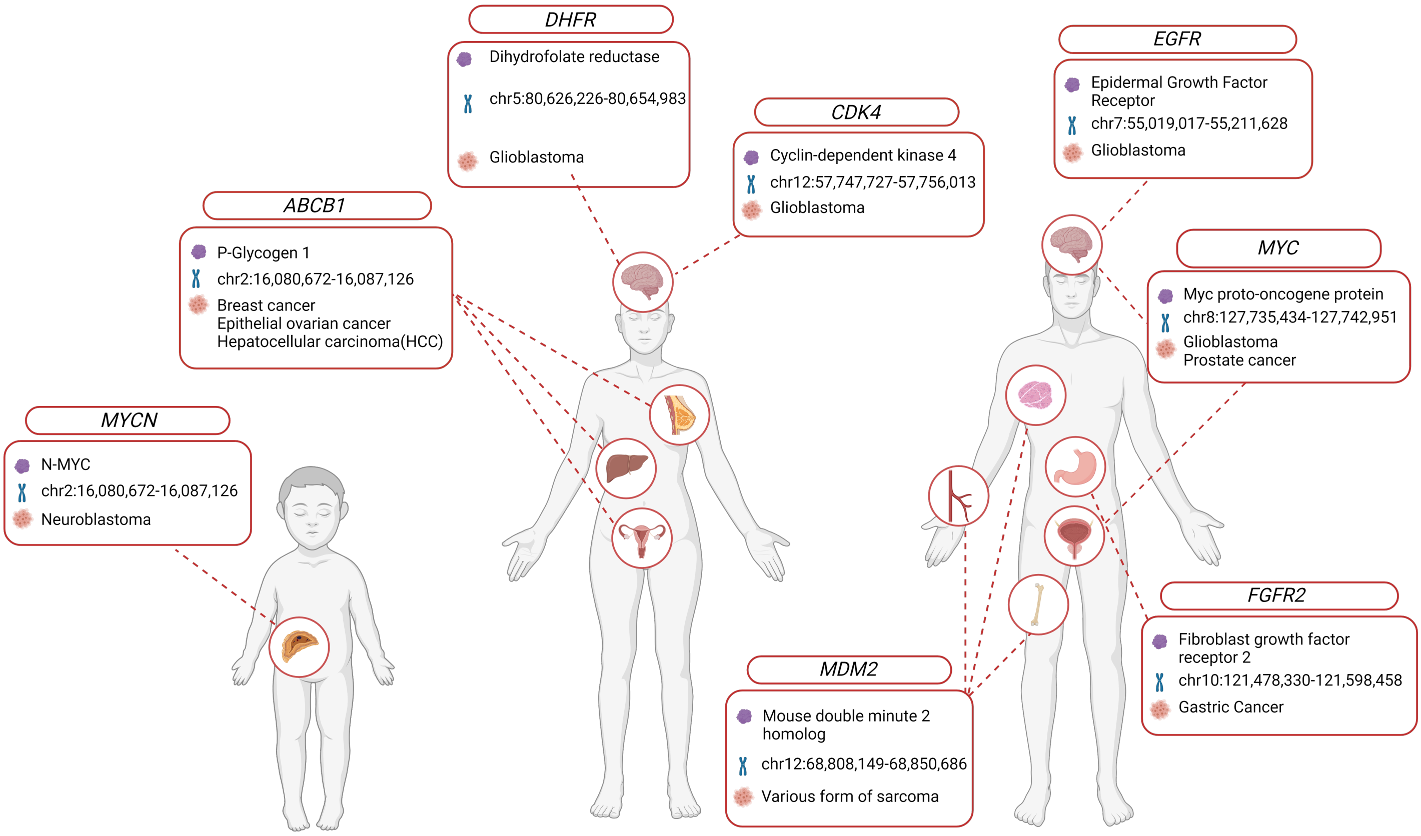
DHFR
Dihydrofolate reductase
chr5:80,626,226-80,654,983
Glioblastoma
CDK4
Cyclin-dependent kinase 4
chr12:57,747,727-57,756,013
Glioblastoma
EGFR
Epidermal Growth Factor Receptor
chr7:55,019,017-55,211,628
Glioblastoma
ABCB1
P-Glycogen 1
chr2:16,080,672-16,087,126
Breast cancer
Epithelial ovarian cancer
Hepatocellular carcinoma(HCC)
MYC
Myc proto-oncogene protein
chr8:127,735,434-127,742,951
Glioblastoma
Prostate cancer
MYCN
N-MYC
chr2:16,080,672-16,087,126
Neuroblastoma
FGFR2
Fibroblast growth factor receptor 2
chr10:121,478,330-121,598,458
Gastric Cancer
MDM2
Mouse double minute 2 homolog
chr12:68,808,149-68,850,686
Various form of sarcoma

### Figure S3

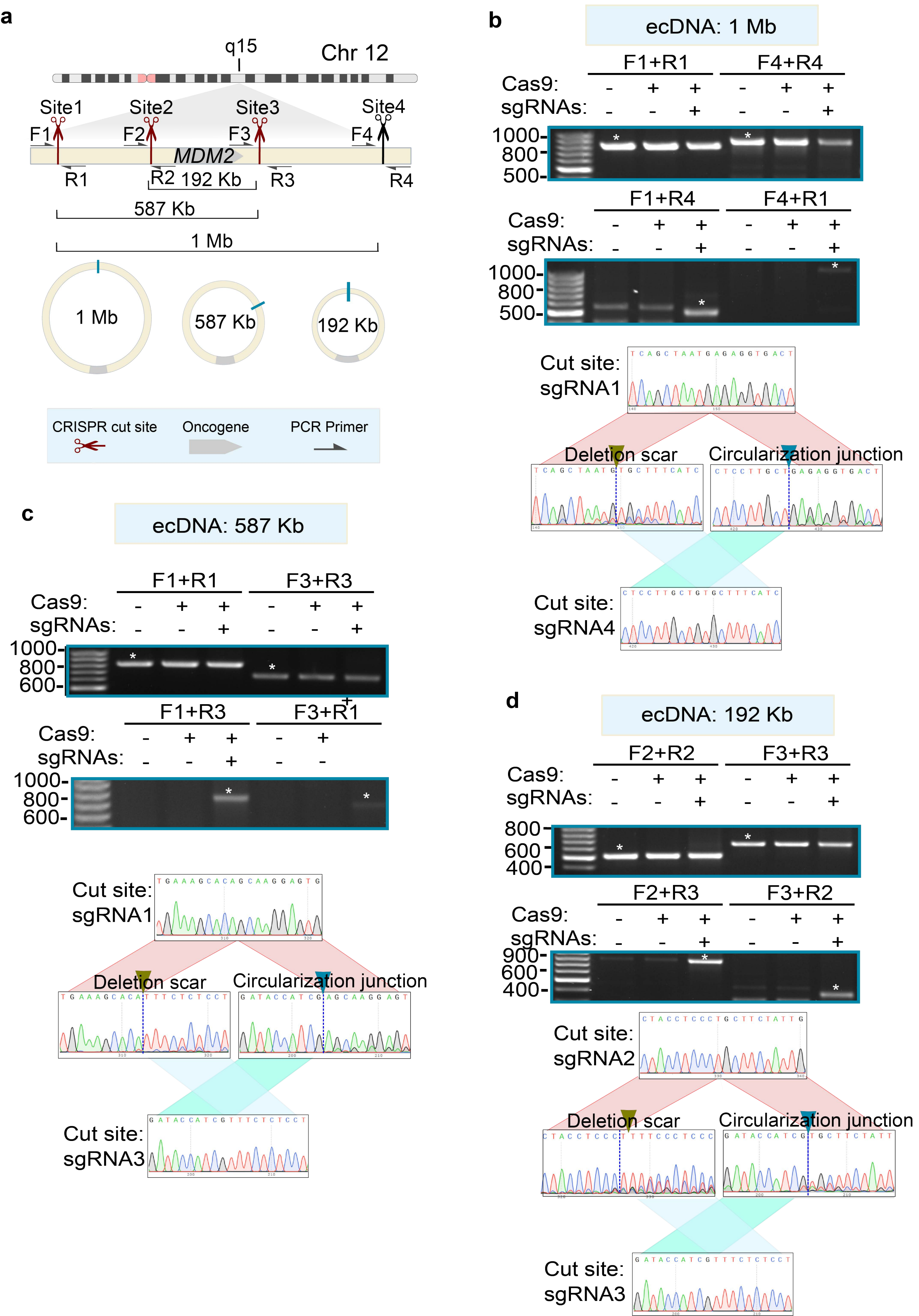
a
q15
Chr 12
Site1
Site2
Site3
Site4
F1
F2
F3
F4
MDM2
R1
R2
192 Kb
R3
R4
587 Kb
1 Mb
1 Mb
587 Kb
192 Kb
CRISPR cut site
Oncogene
PCR Primer
b
ecDNA: 1 Mb
F1+R1
F4+R4
Cas9:
sgRNAs:
1000-
800-
500-
F1+R4
F4+R1
Cut site: sgRNA1
Deletion scar
Circularization junction
Cut site: sgRNA4
c
ecDNA: 587 Kb
F1+R1
F3+R3
1000-
800-
600-
F1+R3
F3+R1
Cut site: sgRNA1
Deletion scar
Circularization junction
Cut site: sgRNA3
d
ecDNA: 192 Kb
F2+R2
F3+R3
800-
600-
400-
F2+R3
F3+R2
900-
Cut site: sgRNA2
Deletion scar
Circularization junction
Cut site: sgRNA3

### Figure S4

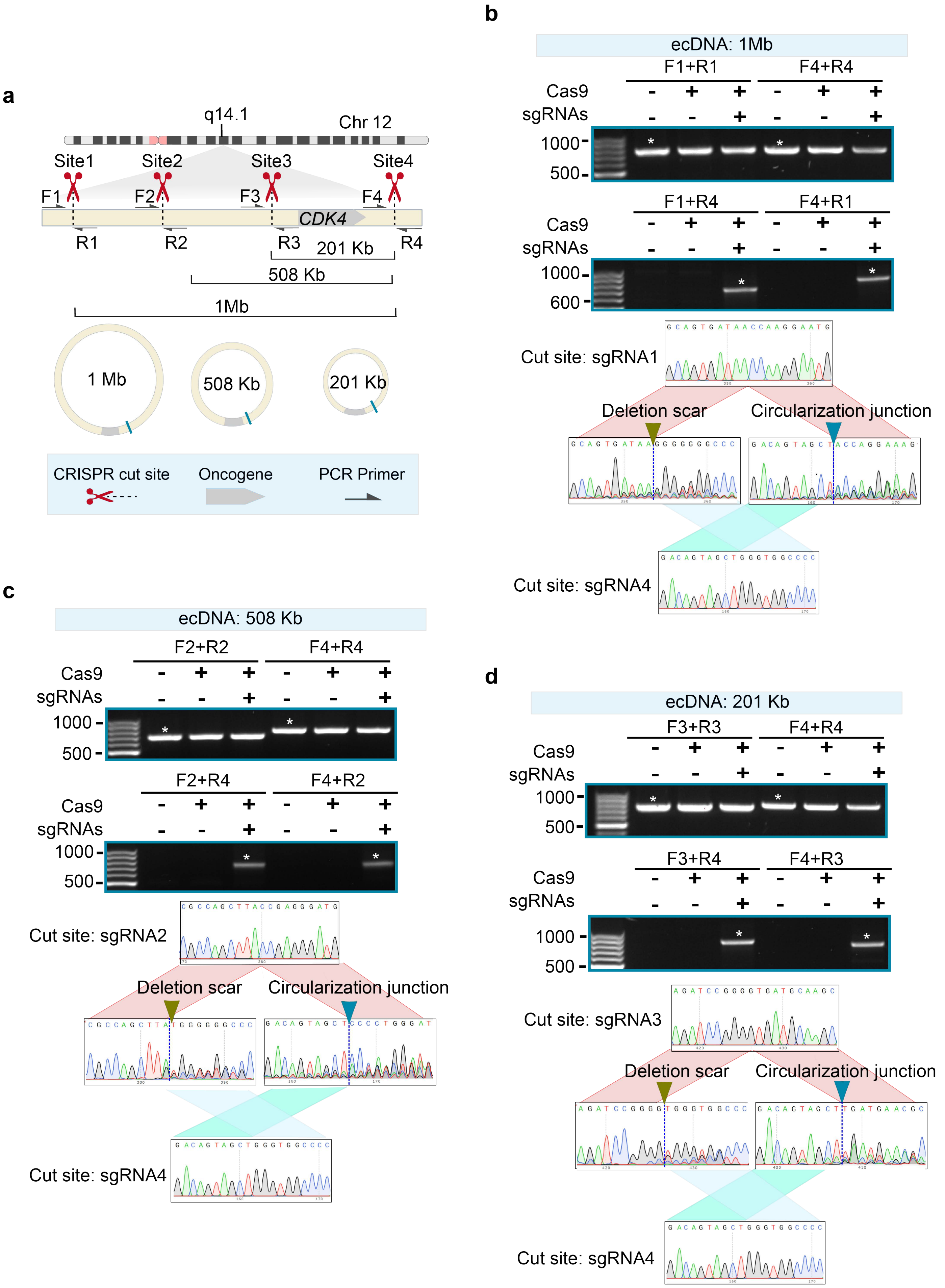
a
q14.1
Chr 12
Site1
Site2
Site3
Site4
F1
F2
F3
F4
CDK4
R1
R2
R3
R4
201 Kb
508 Kb
1Mb
1 Mb
508 Kb
201 Kb
CRISPR cut site
Oncogene
PCR Primer
b
ecDNA: 1Mb
F1+R1
F4+R4
Cas9
sgRNAs
1000
500
F1+R4
F4+R1
Cas9
sgRNAs
1000
600
Cut site: sgRNA1
Deletion scar
Circularization junction
Cut site: sgRNA4
c
ecDNA: 508 Kb
F2+R2
F4+R4
Cas9
sgRNAs
1000
500
F2+R4
F4+R2
Cas9
sgRNAs
1000
500
Cut site: sgRNA2
Deletion scar
Circularization junction
Cut site: sgRNA4
d
ecDNA: 201 Kb
F3+R3
F4+R4
Cas9
sgRNAs
1000
500
F3+R4
F4+R3
Cas9
sgRNAs
1000
500
Cut site: sgRNA3
Deletion scar
Circularization junction
Cut site: sgRNA4

### Figure S5

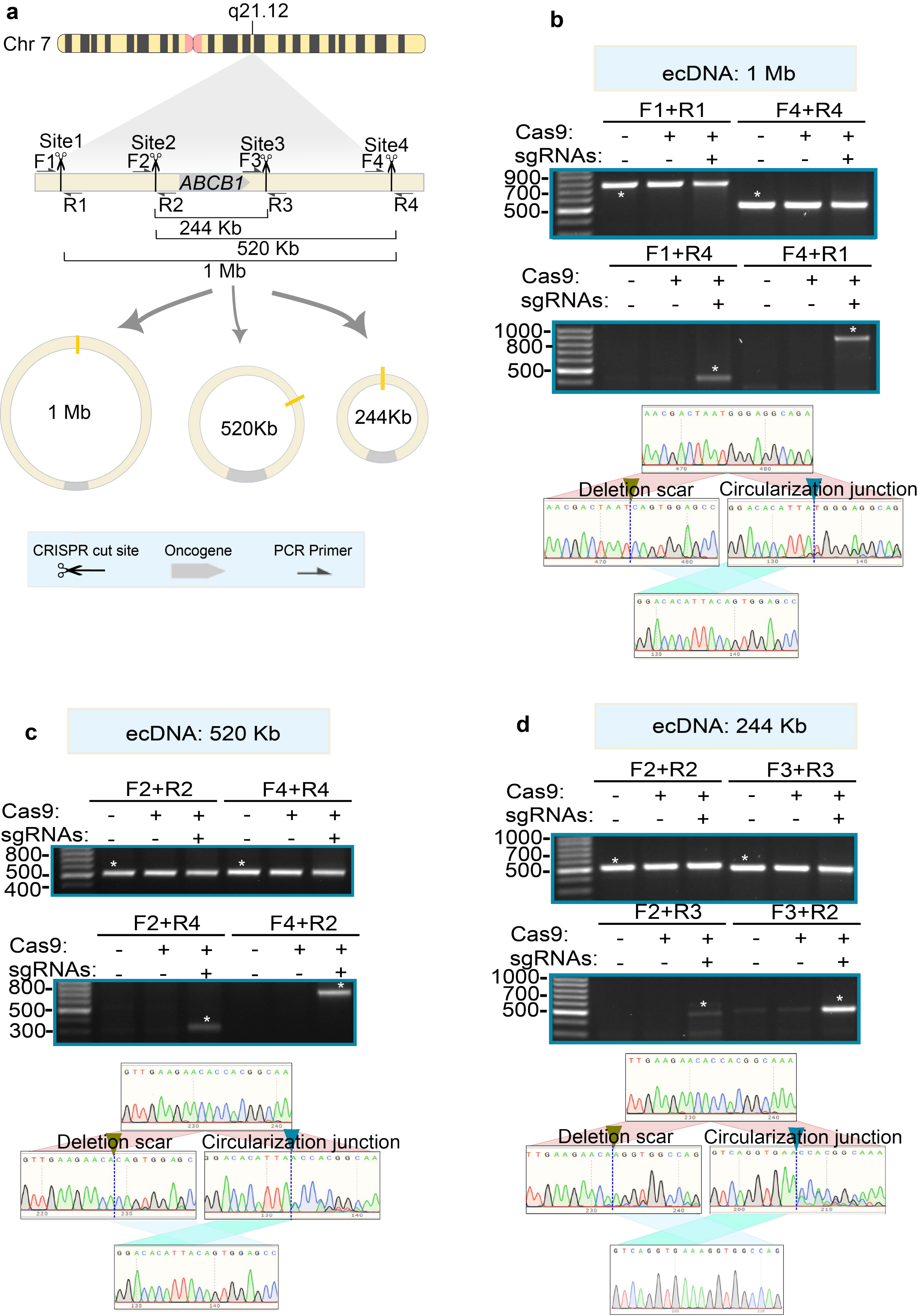
a
q21.12
Chr 7
Site1
Site2
Site3
Site4
F1
F2
F3
F4
ABCB1
R1
R2
R3
R4
244 Kb
520 Kb
1 Mb
1 Mb
520Kb
244Kb
CRISPR cut site
Oncogene
PCR Primer
b
ecDNA: 1 Mb
F1+R1
F4+R4
Cas9:
sgRNAs:
900-
700-
500-
F1+R4
F4+R1
Cas9:
sgRNAs:
1000-
800-
500-
Deletion scar
Circularization junction
c
ecDNA: 520 Kb
F2+R2
F4+R4
Cas9:
sgRNAs:
800-
500-
400-
F2+R4
F4+R2
Cas9:
sgRNAs:
800-
500-
300-
Deletion scar
Circularization junction
d
ecDNA: 244 Kb
F2+R2
F3+R3
Cas9:
sgRNAs:
1000-
700-
500-
F2+R3
F3+R2
Cas9:
sgRNAs:
1000-
700-
500-
Deletion scar
Circularization junction

### Figure S6

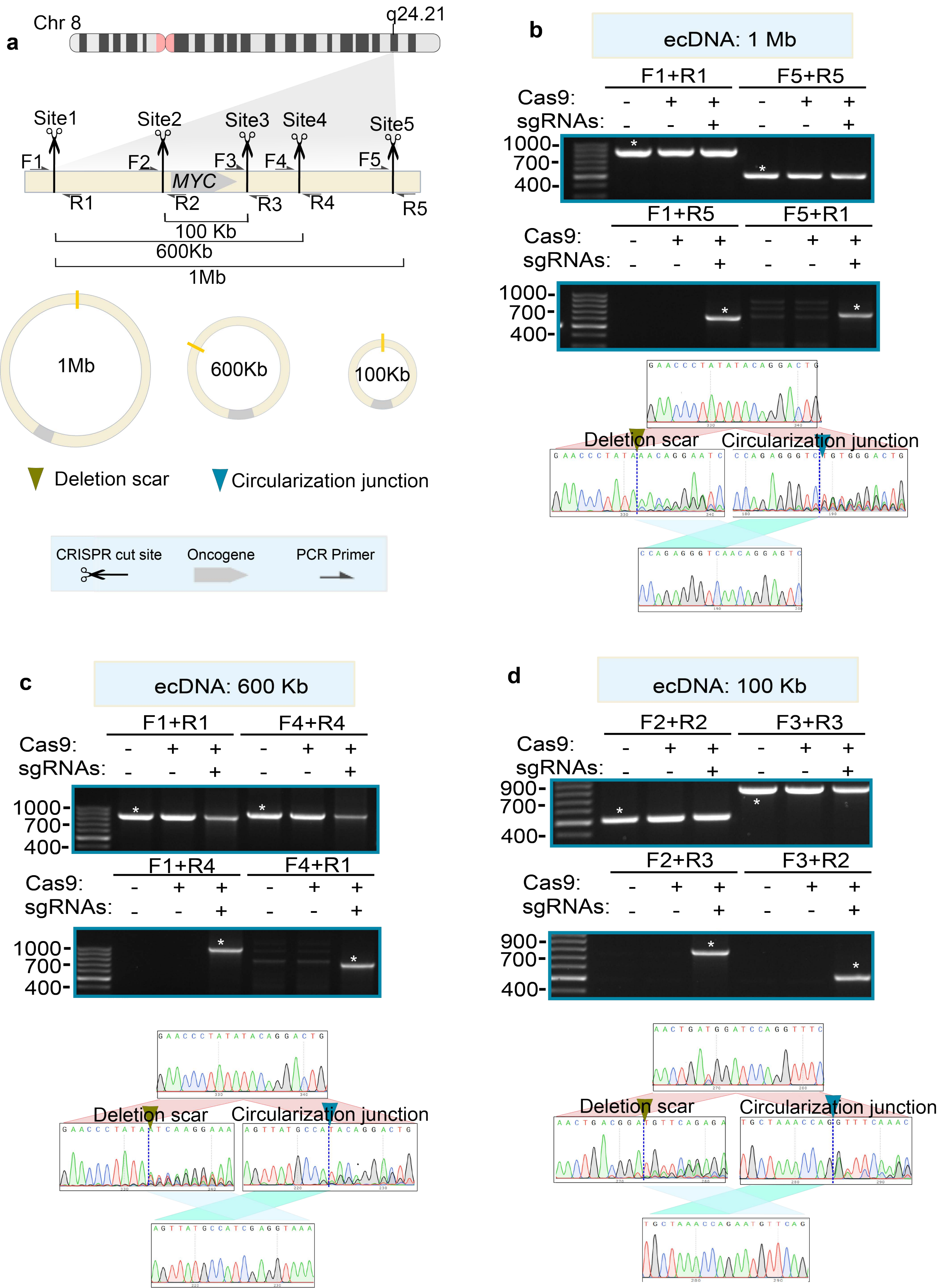

a
Chr 8
q24.21
Site1
Site2
Site3
Site4
Site5
F1
F2
F3
F4
F5
MYC
R1
R2
R3
R4
R5
100 Kb
600Kb
1Mb
1Mb
600Kb
100Kb
Deletion scar
Circularization junction
CRISPR cut site
Oncogene
PCR Primer
b
ecDNA: 1 Mb
F1+R1
F5+R5
Cas9:
sgRNAs:
1000-
700-
400-
F1+R5
F5+R1
Cas9:
sgRNAs:
1000-
700-
400-
Deletion scar
Circularization junction
c
ecDNA: 600 Kb
F1+R1
F4+R4
Cas9:
sgRNAs:
1000-
700-
400-
F1+R4
F4+R1
Cas9:
sgRNAs:
1000-
700-
400-
Deletion scar
Circularization junction
d
ecDNA: 100 Kb
F2+R2
F3+R3
Cas9:
sgRNAs:
900-
700-
400-
F2+R3
F3+R2
Cas9:
sgRNAs:
900-
700-
400-
Deletion scar
Circularization junction

### Figure S7

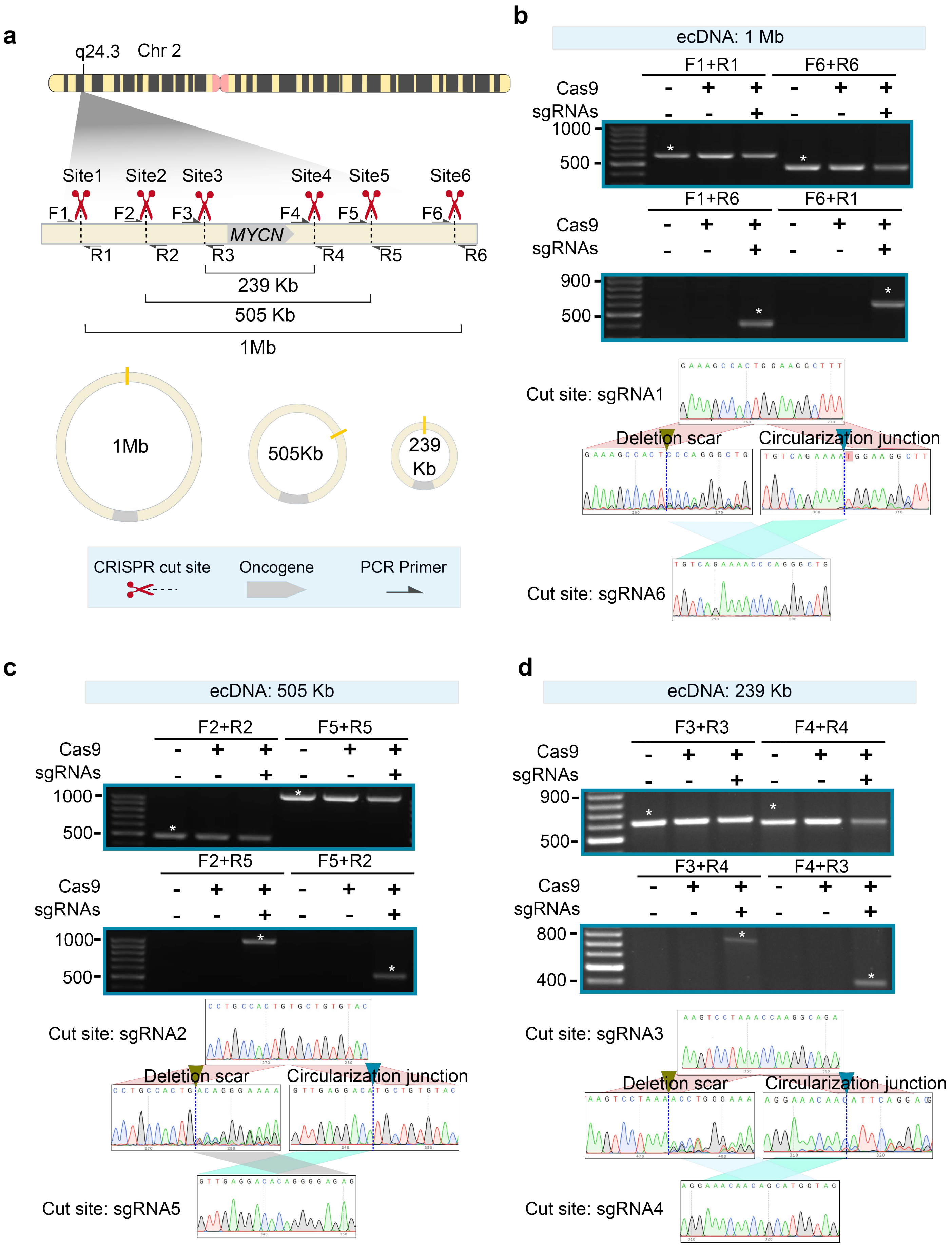
a
q24.3
Chr 2
Site1
Site2
Site3
Site4
Site5
Site6
F1
F2
F3
F4
F5
F6
MYCN
R1
R2
R3
R4
R5
R6
239 Kb
505 Kb
1Mb
1Mb
505Kb
239 Kb
CRISPR cut site
Oncogene
PCR Primer
b
ecDNA: 1 Mb
F1+R1
F6+R6
Cas9
sgRNAs
1000
500
F1+R6
F6+R1
900
500
Cut site: sgRNA1
Deletion scar
Circularization junction
Cut site: sgRNA6
c
ecDNA: 505 Kb
F2+R2
F5+R5
F2+R5
F5+R2
1000
500
Cut site: sgRNA2
Cut site: sgRNA5
d
ecDNA: 239 Kb
F3+R3
F4+R4
F3+R4
F4+R3
900
500
800
400
Cut site: sgRNA3
Cut site: sgRNA4

### Figure S8

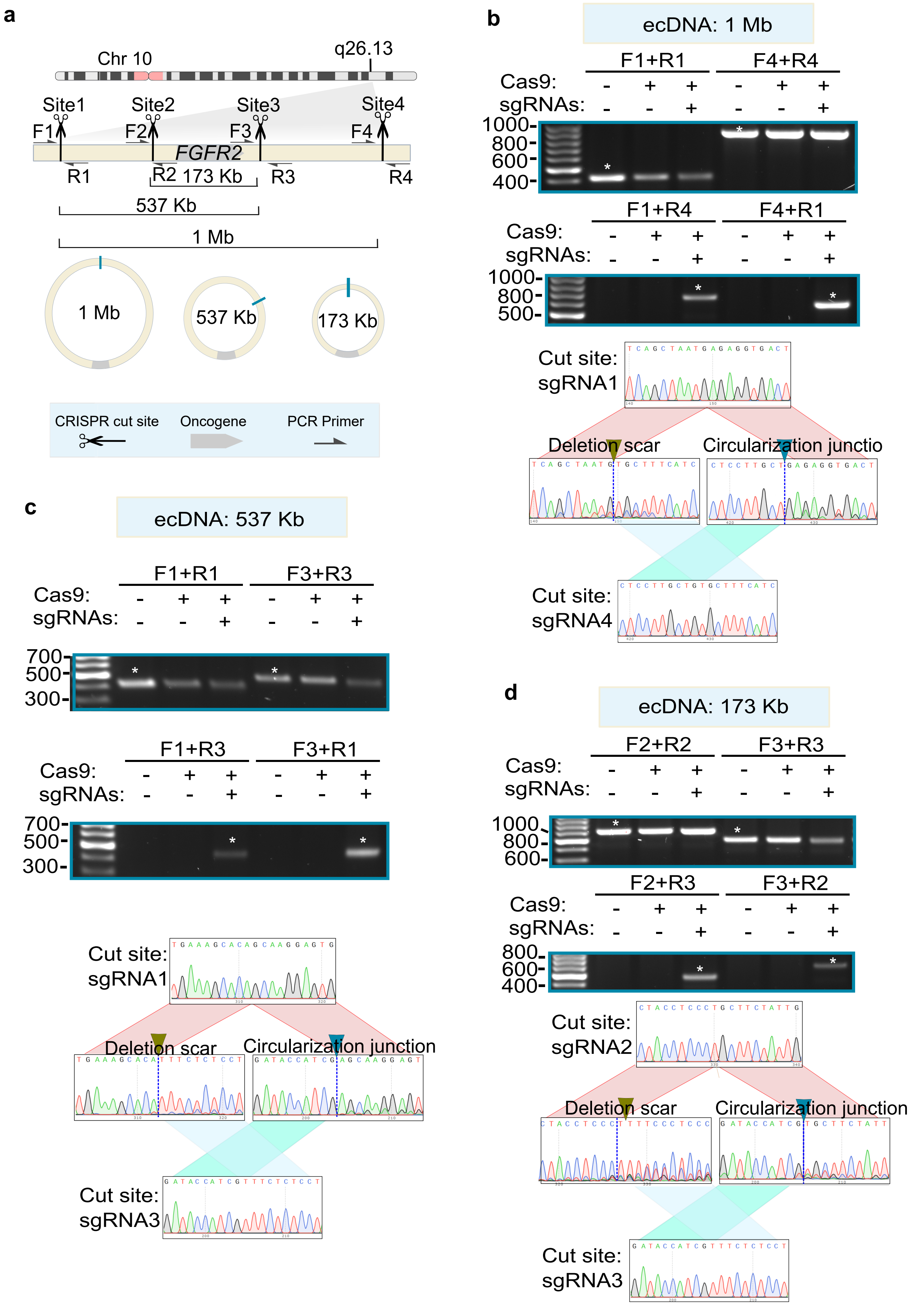
a
Chr 10
q26.13
Site1
Site2
Site3
Site4
F1
F2
F3
F4
FGFR2
R1
R2
R3
R4
173 Kb
537 Kb
1 Mb
1 Mb
537 Kb
173 Kb
CRISPR cut site
Oncogene
PCR Primer
b
ecDNA: 1 Mb
F1+R1
F4+R4
Cas9:
sgRNAs:
1000
800
600
400
F1+R4
F4+R1
Cas9:
sgRNAs:
1000
800
500
Cut site: sgRNA1
Deletion scar
Circularization junctio
Cut site: sgRNA4
c
ecDNA: 537 Kb
F1+R1
F3+R3
Cas9:
sgRNAs:
700
500
300
F1+R3
F3+R1
Cas9:
sgRNAs:
700
500
300
Cut site: sgRNA1
Deletion scar
Circularization junction
Cut site: sgRNA3
d
ecDNA: 173 Kb
F2+R2
F3+R3
Cas9:
sgRNAs:
1000
800
600
F2+R3
F3+R2
Cas9:
sgRNAs:
800
600
400
Cut site: sgRNA2
Deletion scar
Circularization junction
Cut site: sgRNA3

### Figure S9

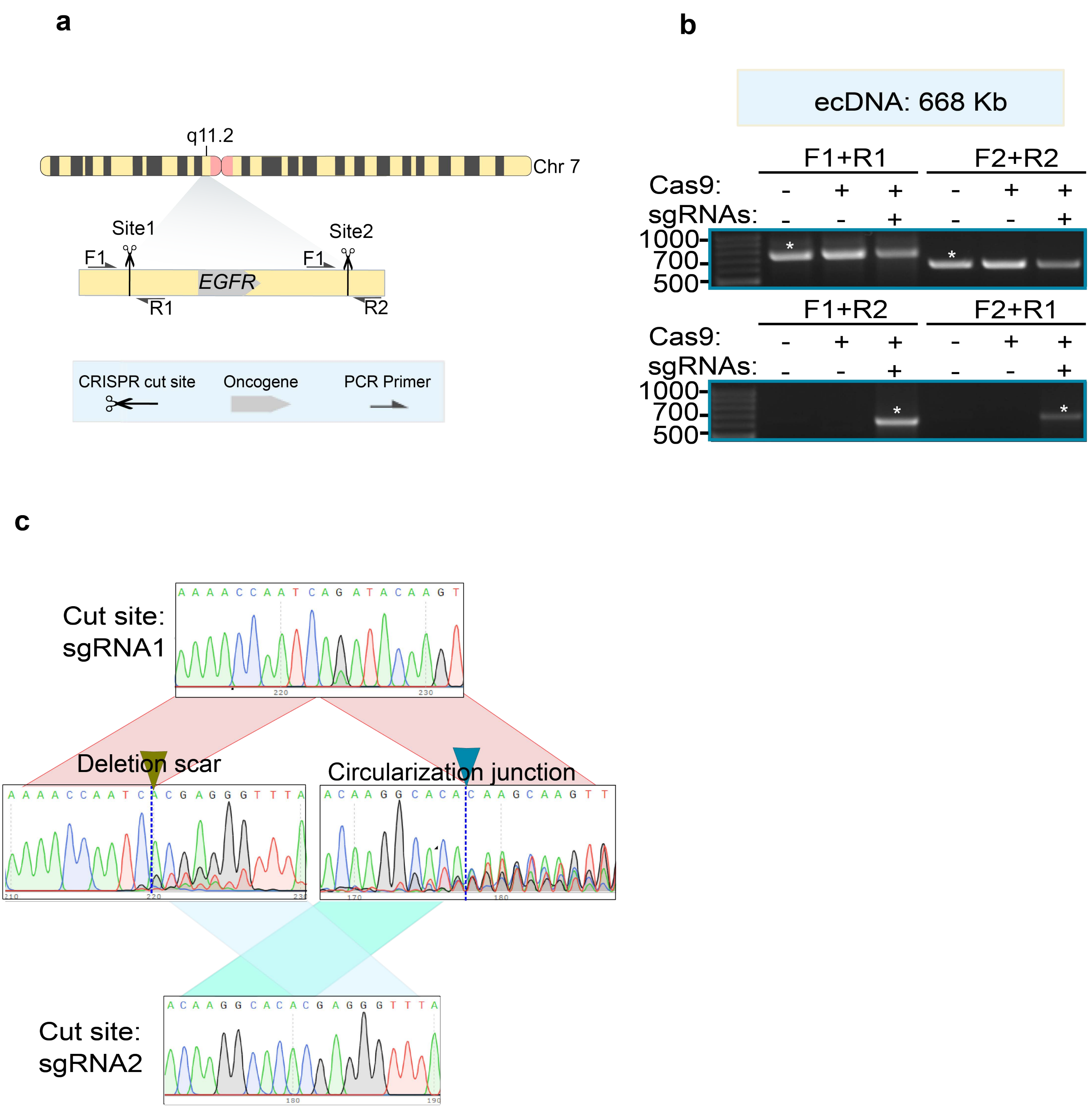

a
q11.2
Chr 7
Site1
Site2
F1
F1
EGFR
R1
R2
CRISPR cut site
Oncogene
PCR Primer
b
ecDNA: 668 Kb
F1+R1
F2+R2
Cas9: - + + - + +
sgRNAs: - - + - - +
1000-
700-
500-
F1+R2
F2+R1
Cas9: - + + - + +
sgRNAs: - - + - - +
1000-
700-
500-
c
Cut site: sgRNA1
A A A A C C A A T C A G A T A C A A G T
Deletion scar
A A A A C C A A T C A C G A G G G T T T A
Circularization junction
A C A A G G C A C A C A A G C A A G T T
Cut site: sgRNA2
A C A A G G C A C A C G A G G G T T T A

### Figure S10

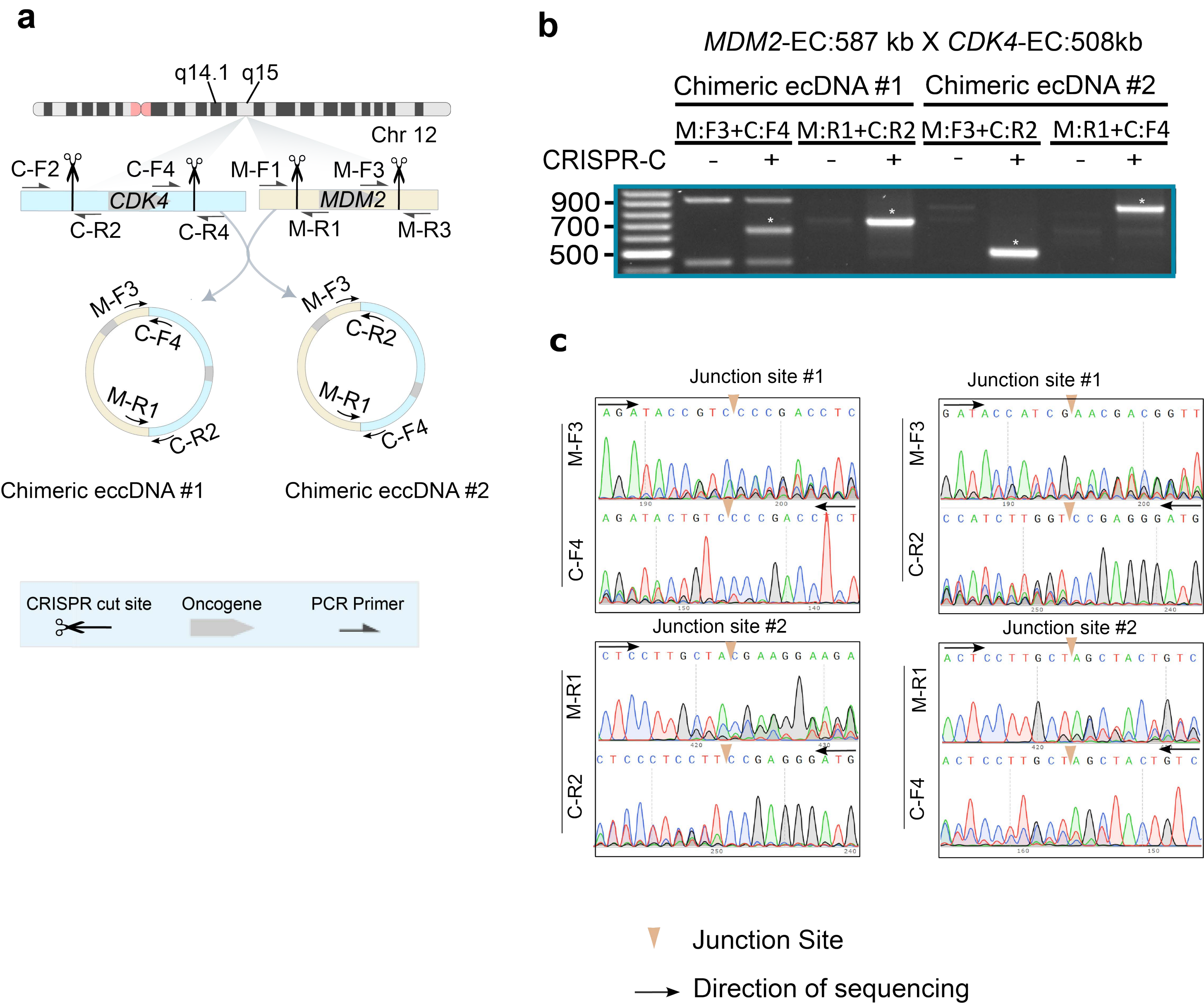

a
q14.1
q15
Chr 12
C-F2
C-F4
M-F1
M-F3
CDK4
MDM2
C-R2
C-R4
M-R1
M-R3
Chimeric eccDNA #1
Chimeric eccDNA #2
CRISPR cut site
Oncogene
PCR Primer
b
MDM2-EC:587 kb X CDK4-EC:508kb
Chimeric ecDNA #1
Chimeric ecDNA #2
M:F3+C:F4
M:R1+C:R2
M:F3+C:R2
M:R1+C:F4
CRISPR-C
900
700
500
c
Junction site #1
Junction site #2
M-F3
C-F4
C-R2
M-R1
Junction Site
Direction of sequencing

### Figure S11

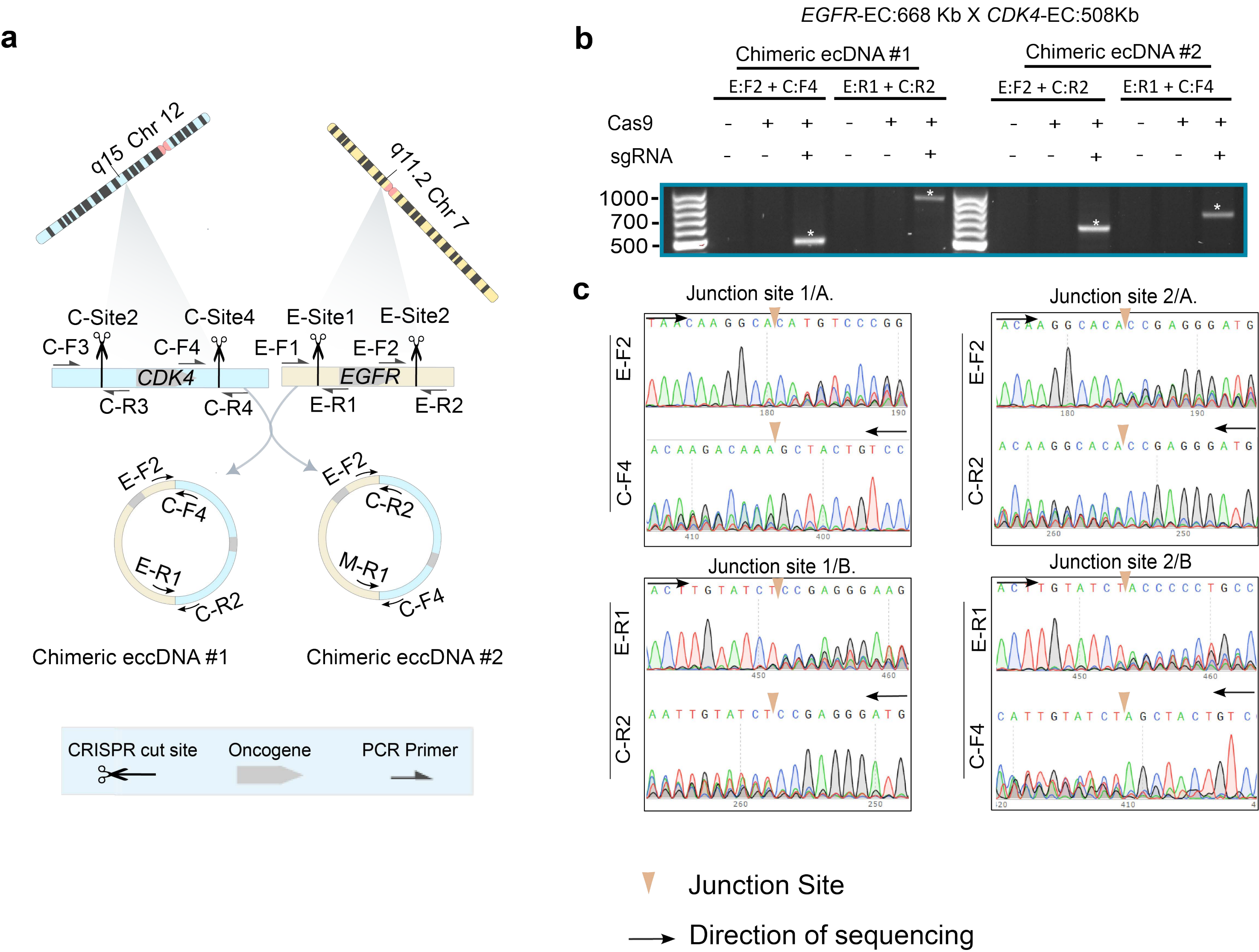

a
q15 Chr 12
q11.2 Chr 7
C-Site2
C-Site4
E-Site1
E-Site2
C-F3
C-F4
E-F1
E-F2
CDK4
EGFR
C-R3
C-R4
E-R1
E-R2
Chimeric eccDNA #1
Chimeric eccDNA #2
CRISPR cut site
Oncogene
PCR Primer
b
EGFR-EC:668 Kb X CDK4-EC:508Kb
Chimeric ecDNA #1
Chimeric ecDNA #2
E:F2 + C:F4
E:R1 + C:R2
E:F2 + C:R2
E:R1 + C:F4
Cas9
sgRNA
1000
700
500
c
Junction site 1/A.
Junction site 2/A.
Junction site 1/B.
Junction site 2/B
Junction Site
Direction of sequencing

### Figure S12

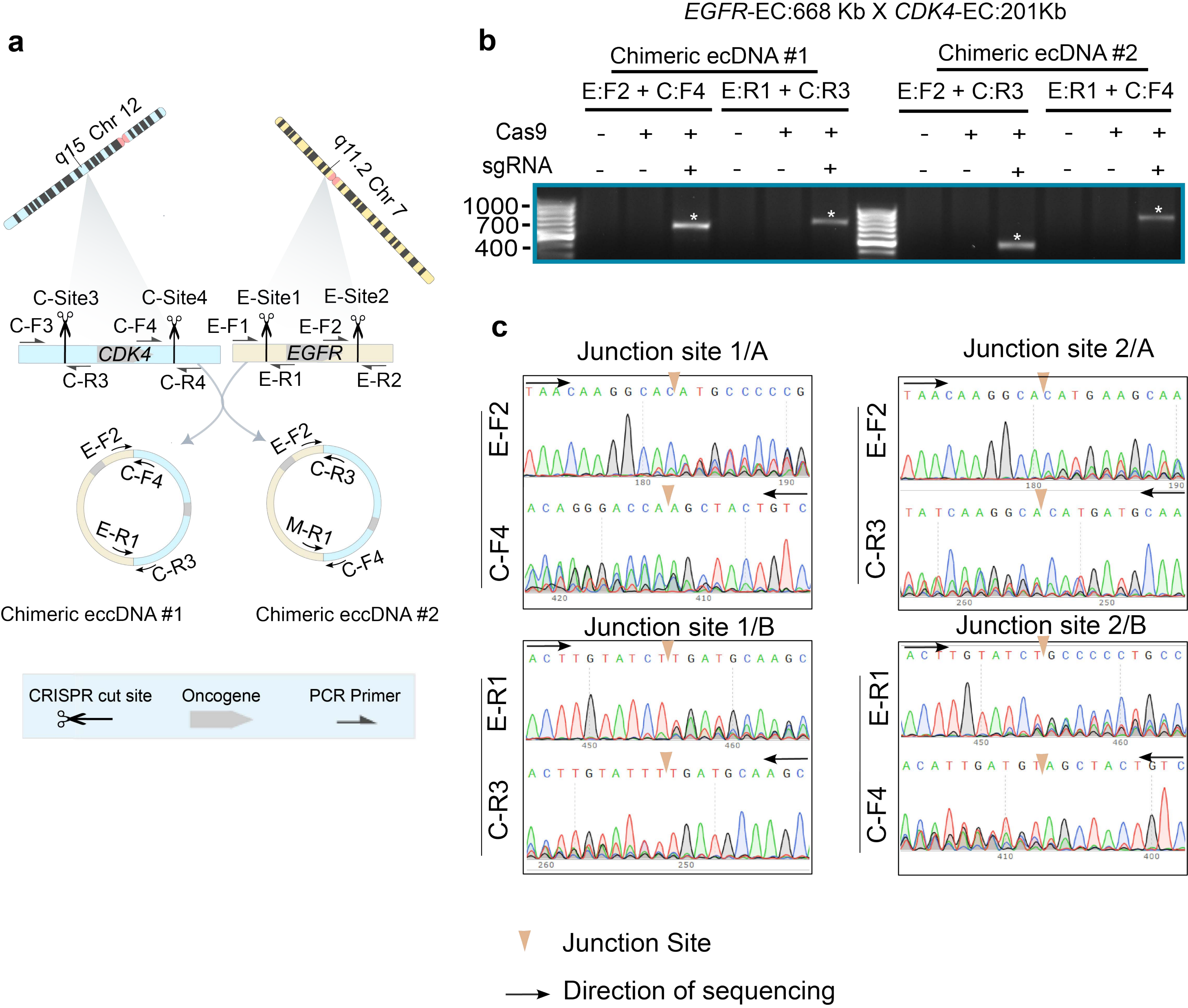

a
q15 Chr 12
q11.2 Chr 7
C-Site3
C-Site4
E-Site1
E-Site2
C-F3
C-F4
E-F1
E-F2
CDK4
EGFR
C-R3
C-R4
E-R1
E-R2
E-F2
C-F4
E-R1
C-R3
E-F2
C-R3
M-R1
C-F4
Chimeric eccDNA #1
Chimeric eccDNA #2
CRISPR cut site
Oncogene
PCR Primer
b
EGFR-EC:668 Kb X CDK4-EC:201Kb
Chimeric ecDNA #1
Chimeric ecDNA #2
E:F2 + C:F4
E:R1 + C:R3
E:F2 + C:R3
E:R1 + C:F4
Cas9
sgRNA
1000
700
400
c
Junction site 1/A
Junction site 2/A
Junction site 1/B
Junction site 2/B
E-F2
C-F4
E-F2
C-R3
E-R1
C-R3
E-R1
C-F4
Junction Site
Direction of sequencing

### Figure S13

#### CRISPR-C: *ABCB1*

a.

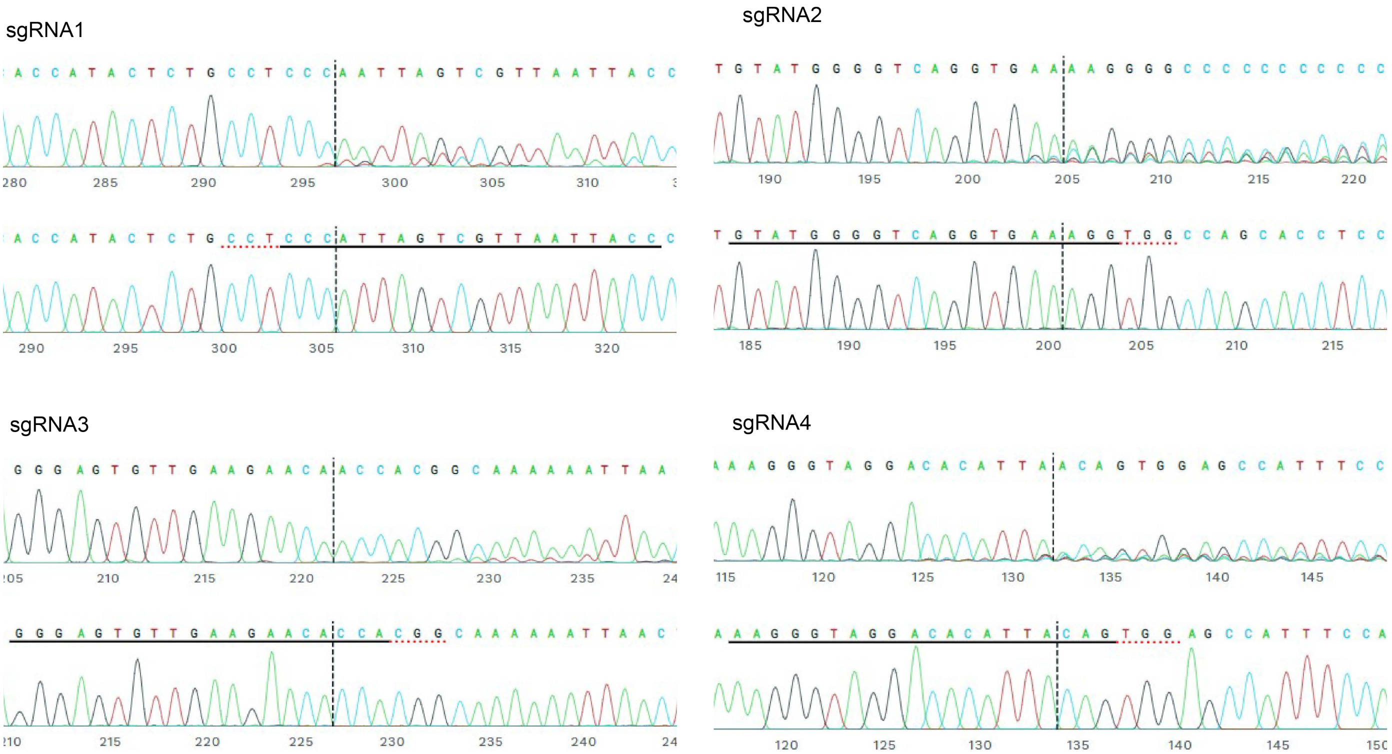


b.

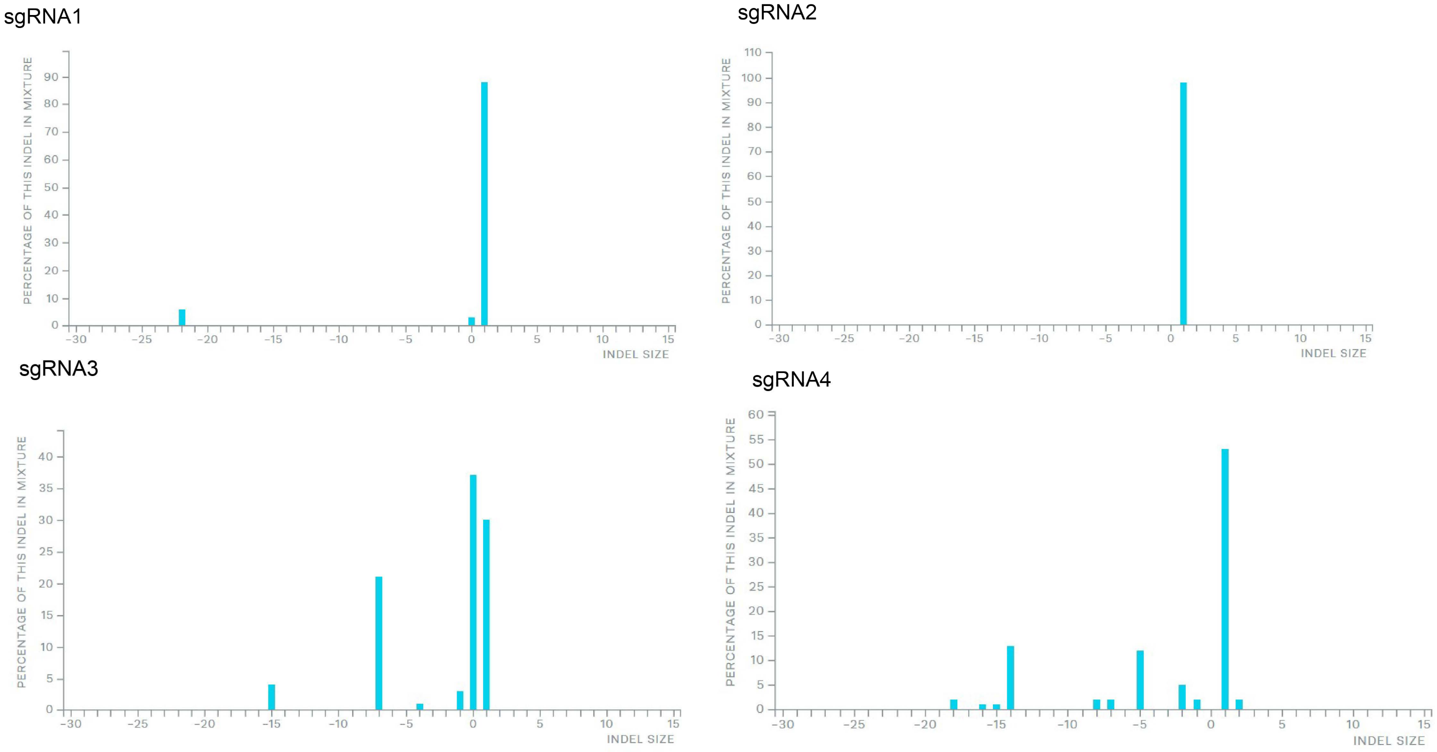

**CRISPR-C: *CDK4***

### Figure S14

**a**

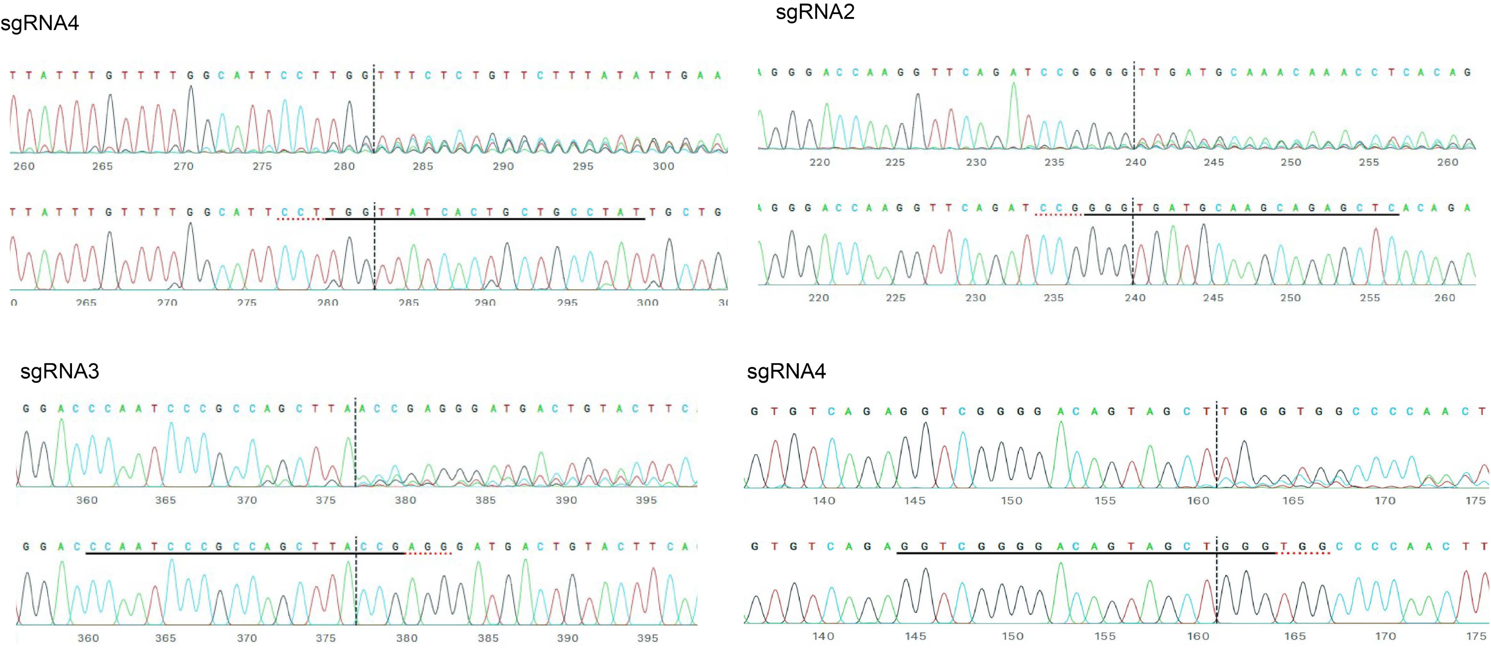


**b**

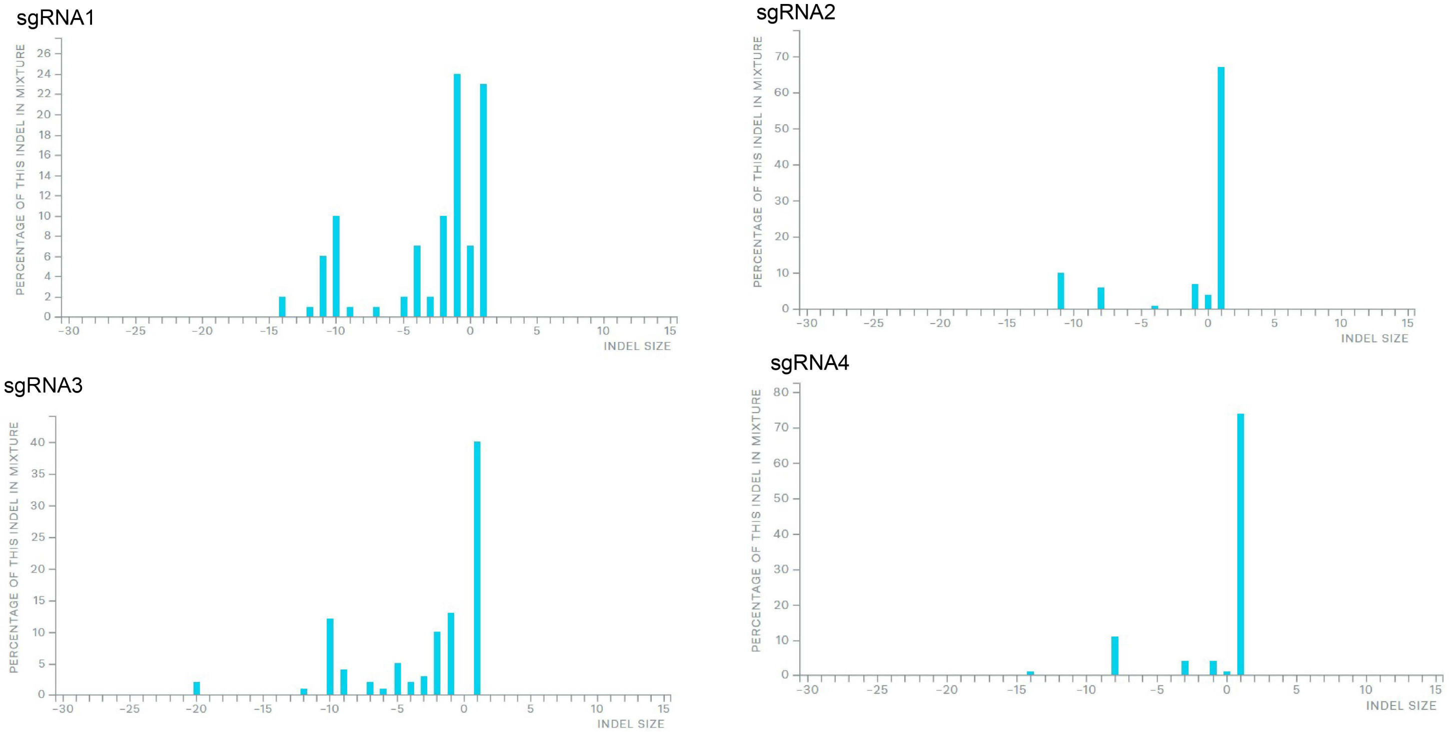

### Figure S15

### CRISPR-C: *DHFR*

## a

sgRNA1

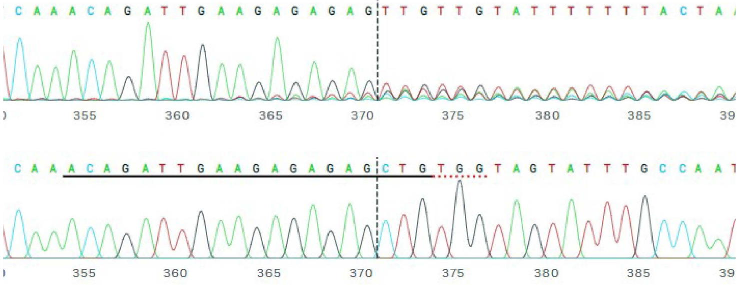

sgRNA2

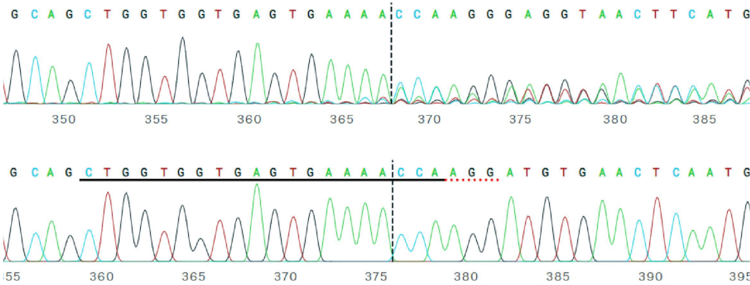

## b

sgRNA1

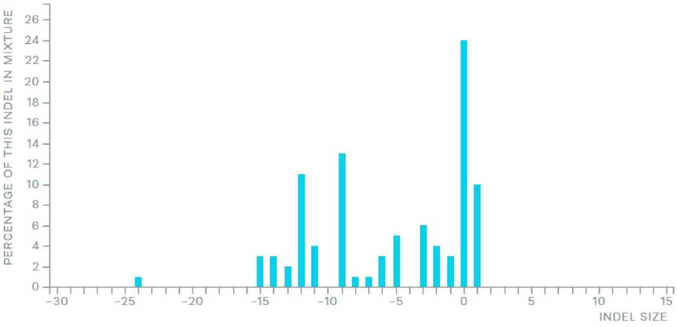


sgRNA2

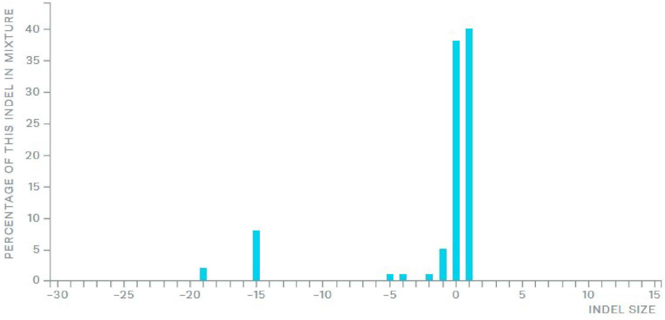

### CRISPR-C: *EGFR*

### Figure S16

**a**

sgRNA1

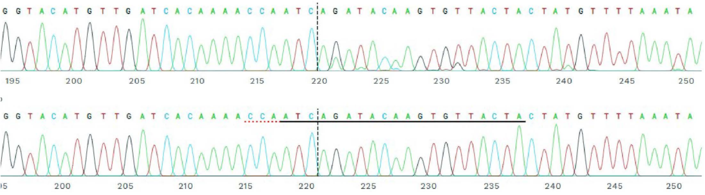

sgRNA2

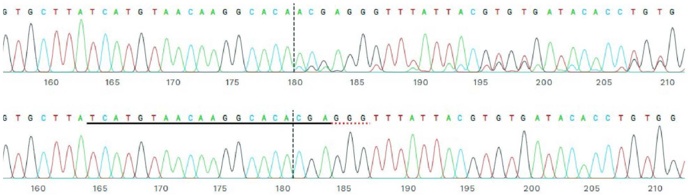

**b**

sgRNA1

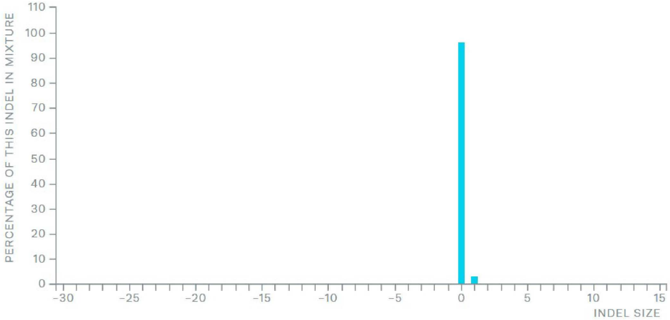


sgRNA2

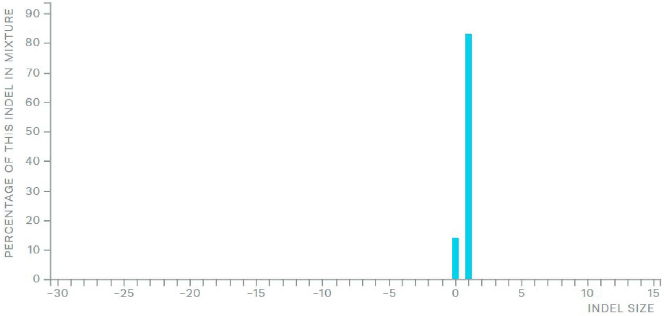

CRISPR-C: *FGFR2*

**Figure S17**

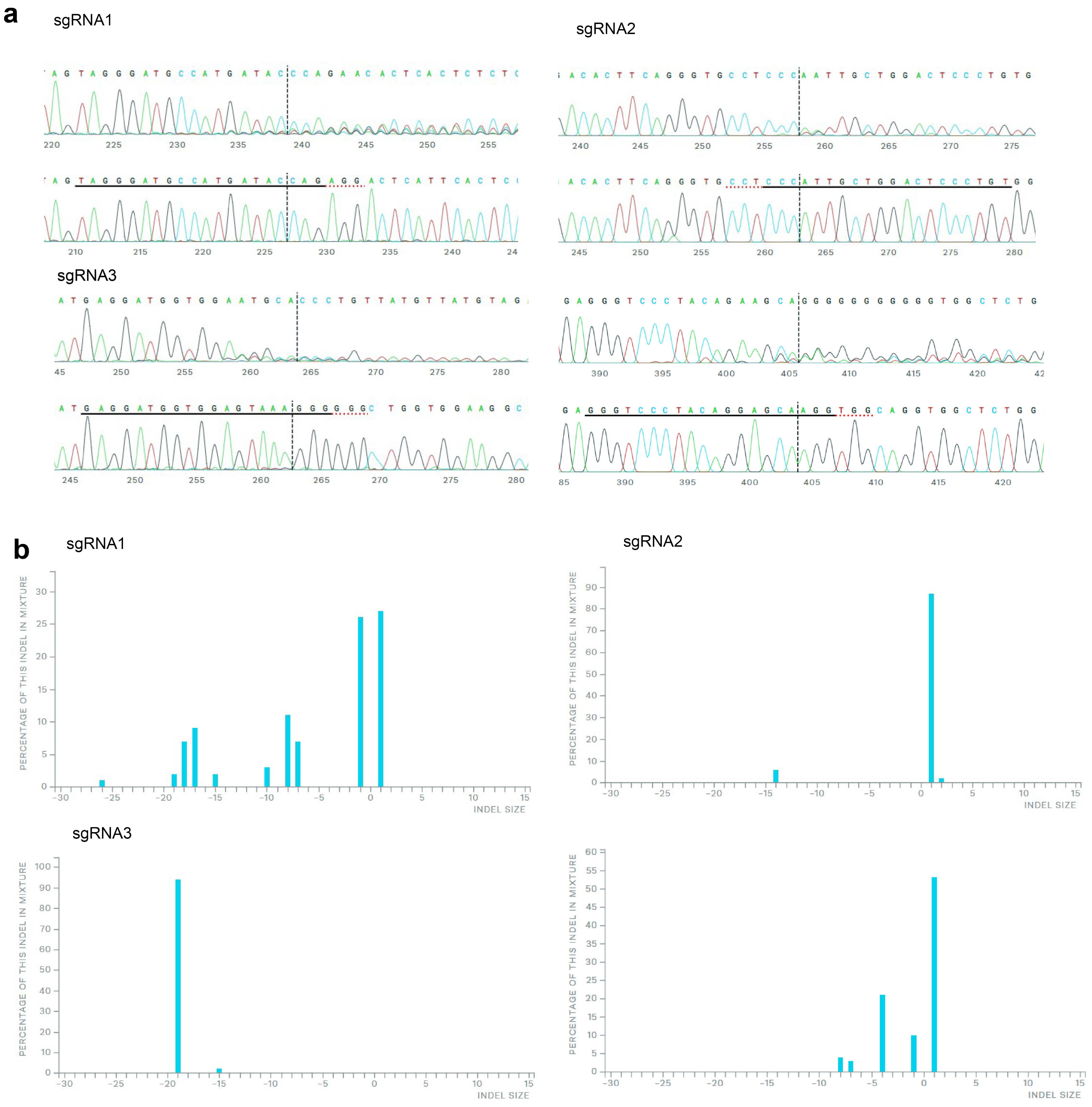

CRISPR-C: *MDM2*

### Figure S18

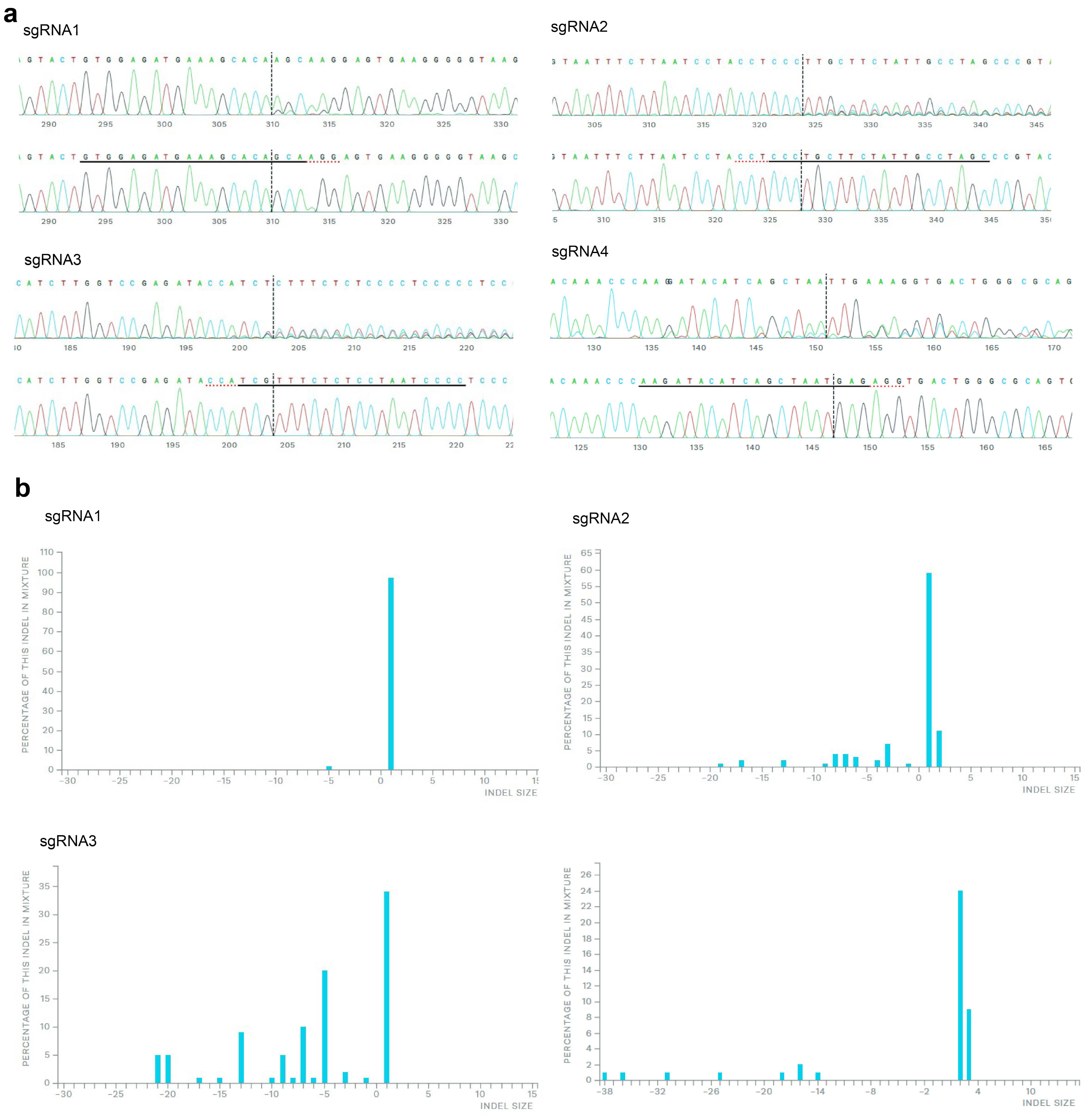

### CRISPR-C: *MYC*

**Figure S19**

**a**

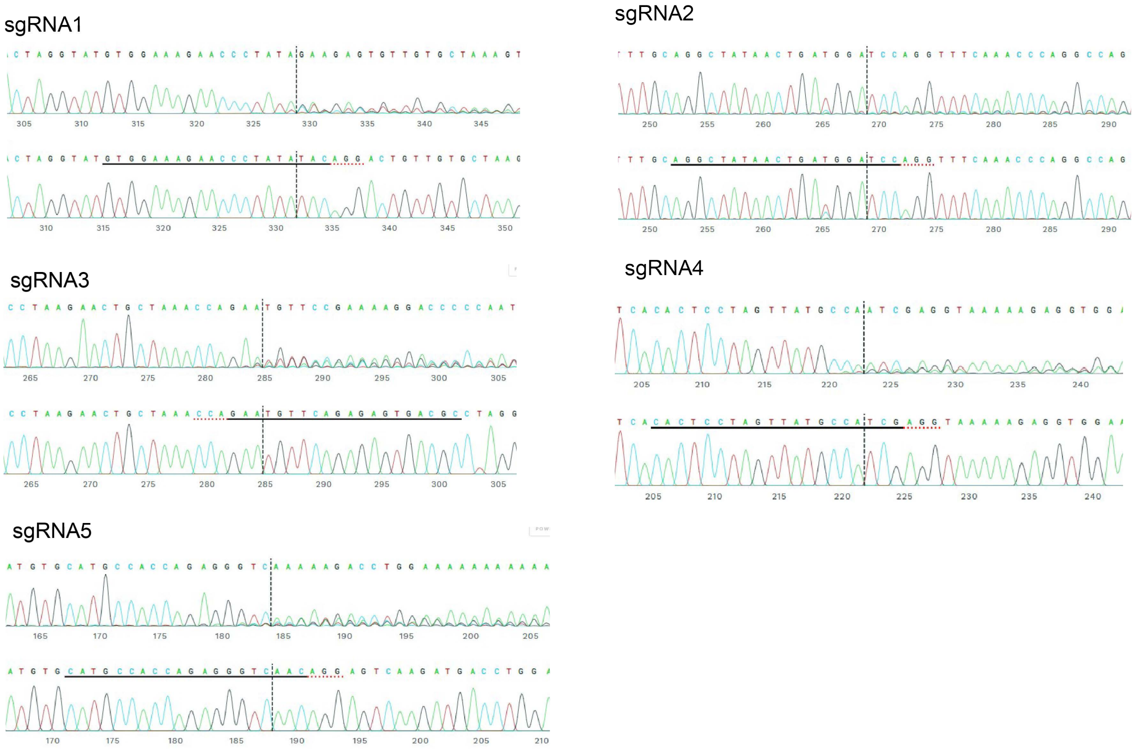


b

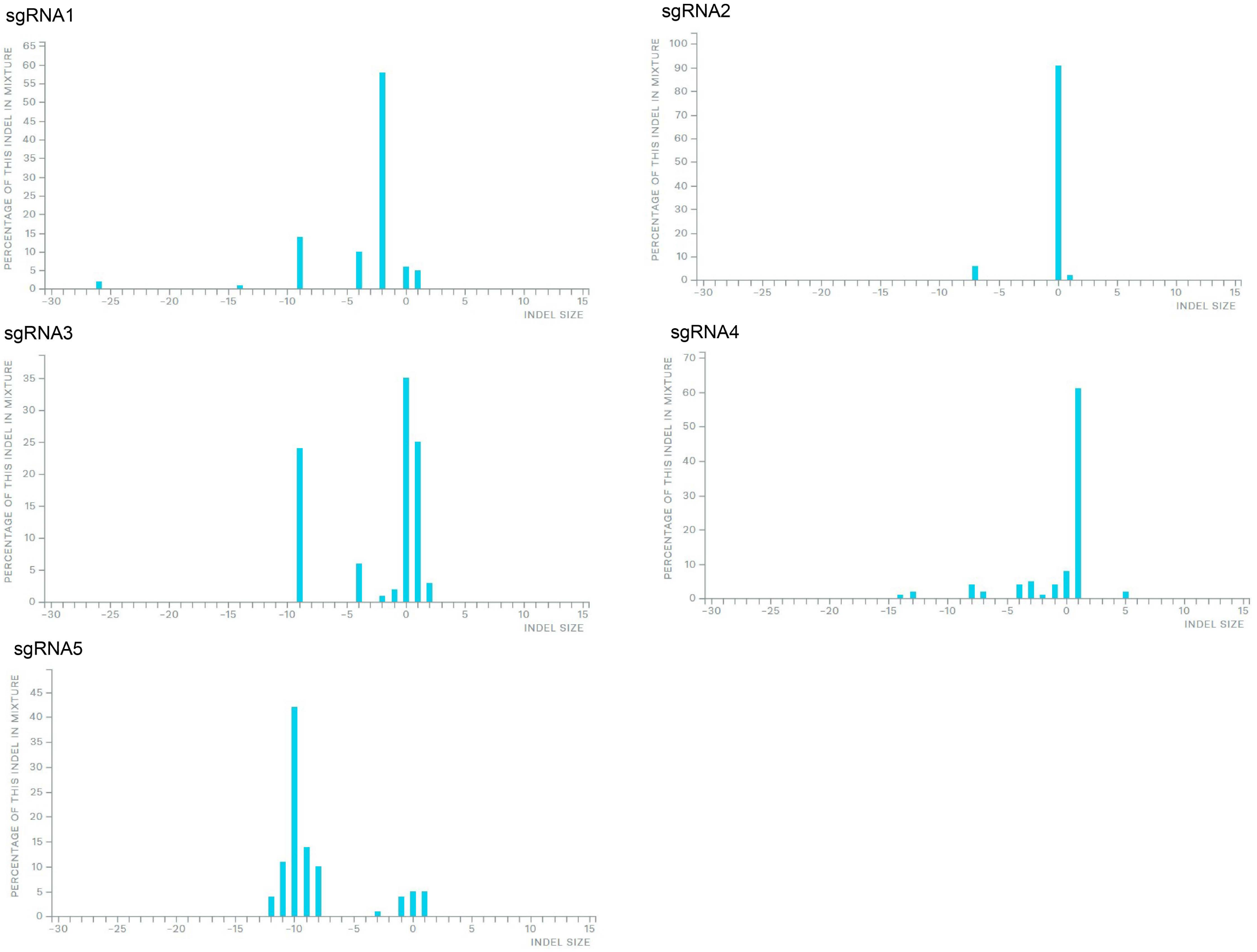

**CRISPR-C: *MYCN***

**Figure S20**

**a**

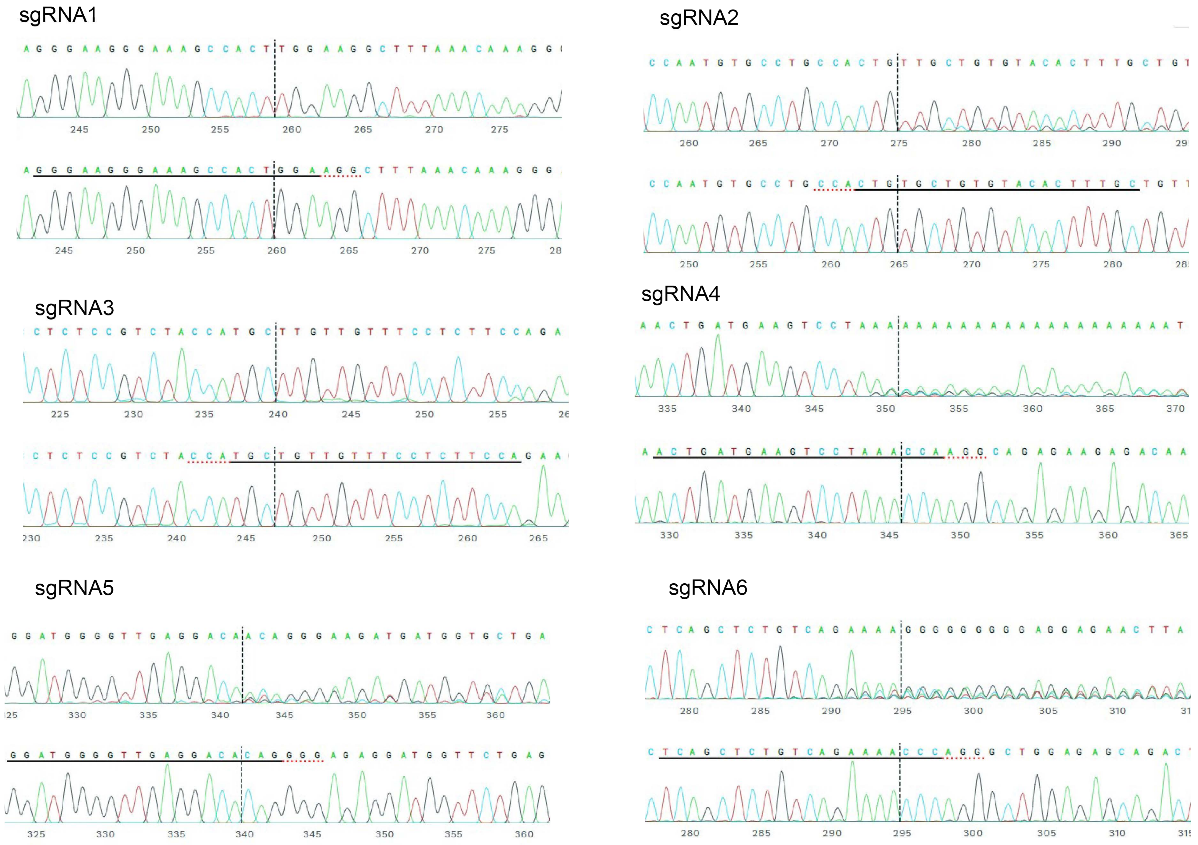


**b**

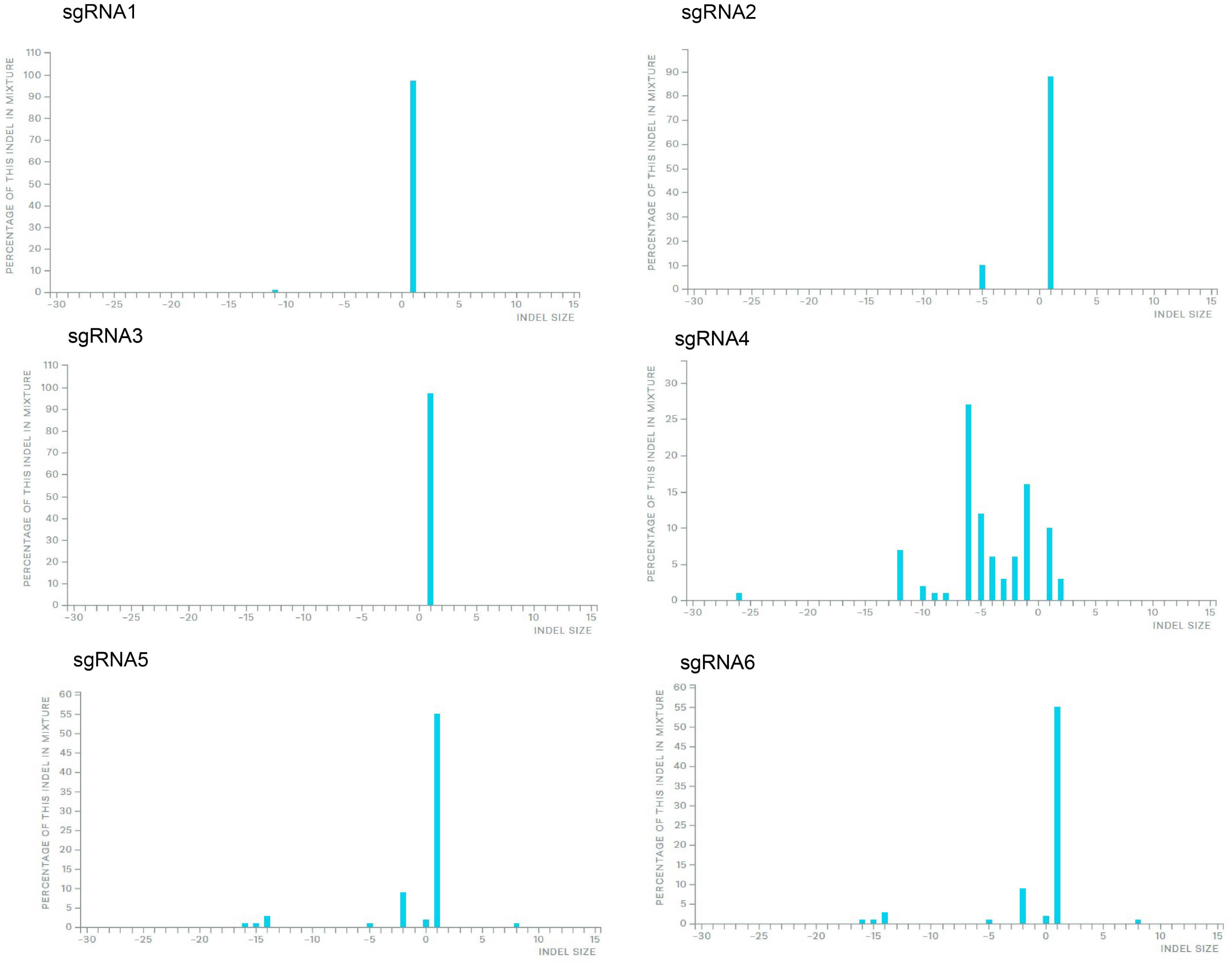

### Figure S21

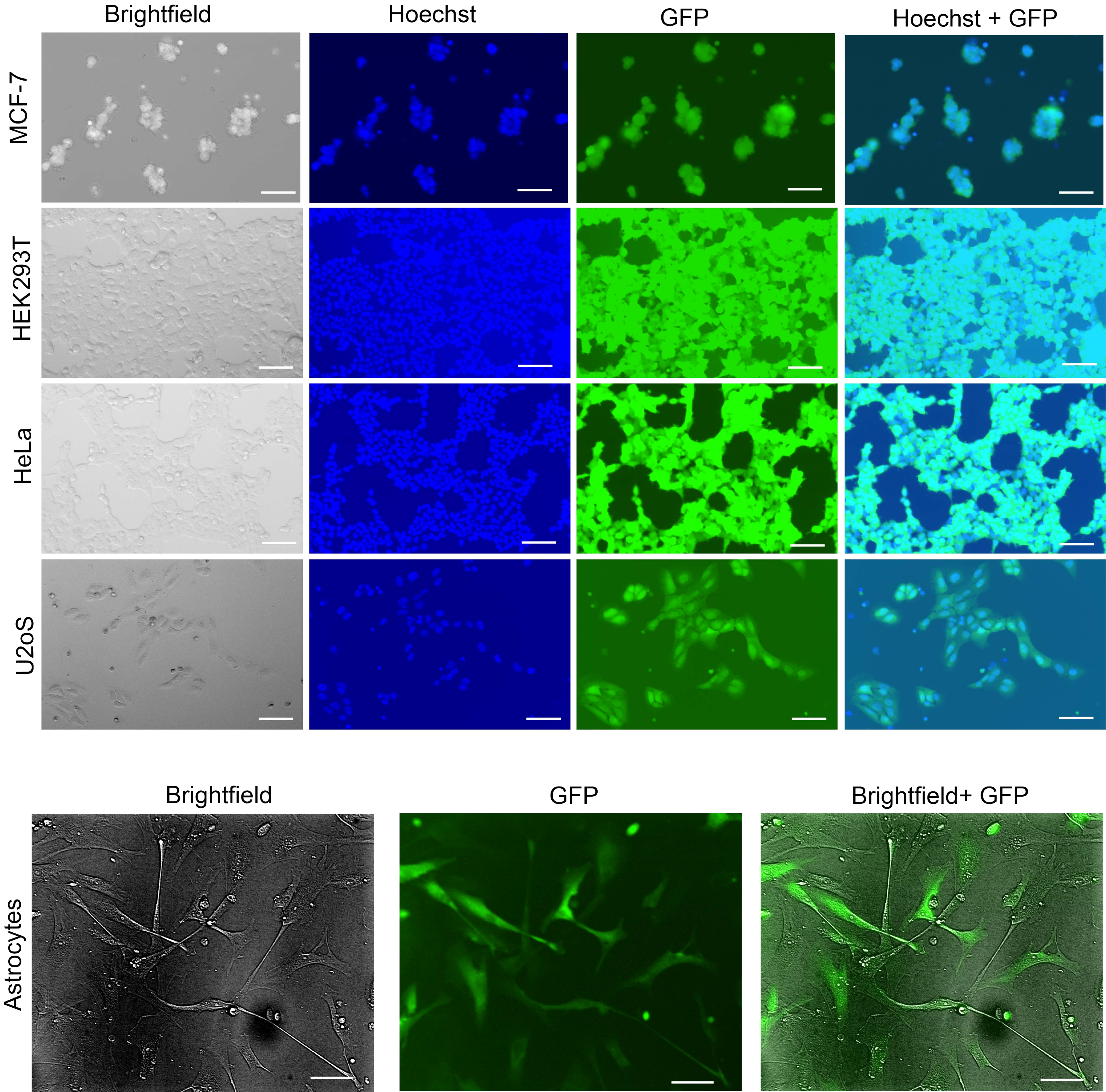
Brightfield
Hoechst
GFP
Hoechst + GFP
MCF-7
HEK293T
HeLa
U2oS
Brightfield
GFP
Brightfield+ GFP
Astrocytes

### Figure S22

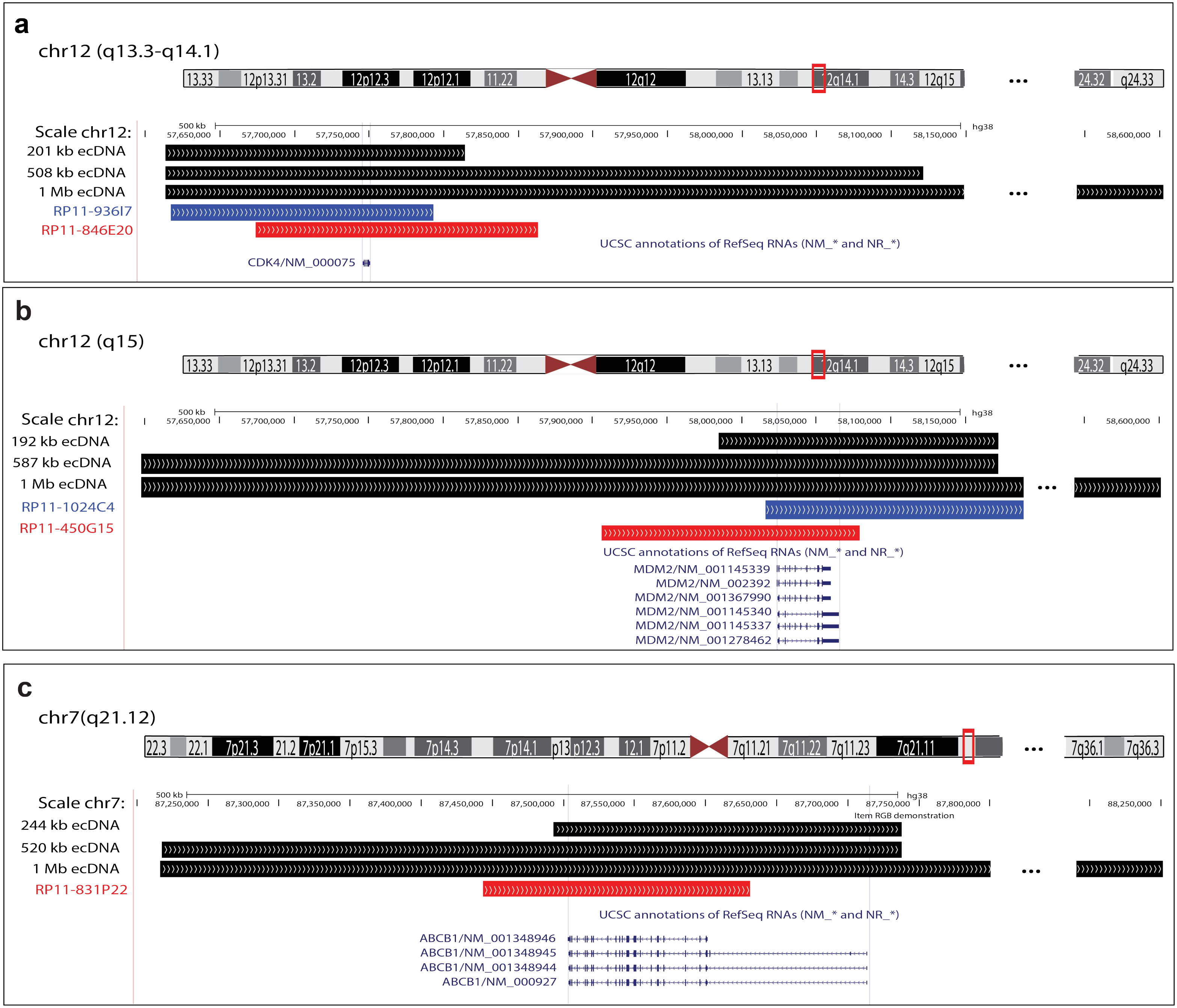
a
chr12 (q13.3-q14.1)
Scale chr12:
500 kb
hg38
201 kb ecDNA
508 kb ecDNA
1 Mb ecDNA
RP11-936I7
RP11-846E20
UCSC annotations of RefSeq RNAs (NM_* and NR_*)
CDK4/NM_000075
b
chr12 (q15)
Scale chr12:
192 kb ecDNA
587 kb ecDNA
1 Mb ecDNA
RP11-1024C4
RP11-450G15
UCSC annotations of RefSeq RNAs (NM_* and NR_*)
MDM2/NM_001145339
MDM2/NM_002392
MDM2/NM_001367990
MDM2/NM_001145340
MDM2/NM_001145337
MDM2/NM_001278462
c
chr7(q21.12)
Scale chr7:
Item RGB demonstration
244 kb ecDNA
520 kb ecDNA
1 Mb ecDNA
RP11-831P22
UCSC annotations of RefSeq RNAs (NM_* and NR_*)
ABCB1/NM_001348946
ABCB1/NM_001348945
ABCB1/NM_001348944
ABCB1/NM_000927

#### Figure S23

U2OS:

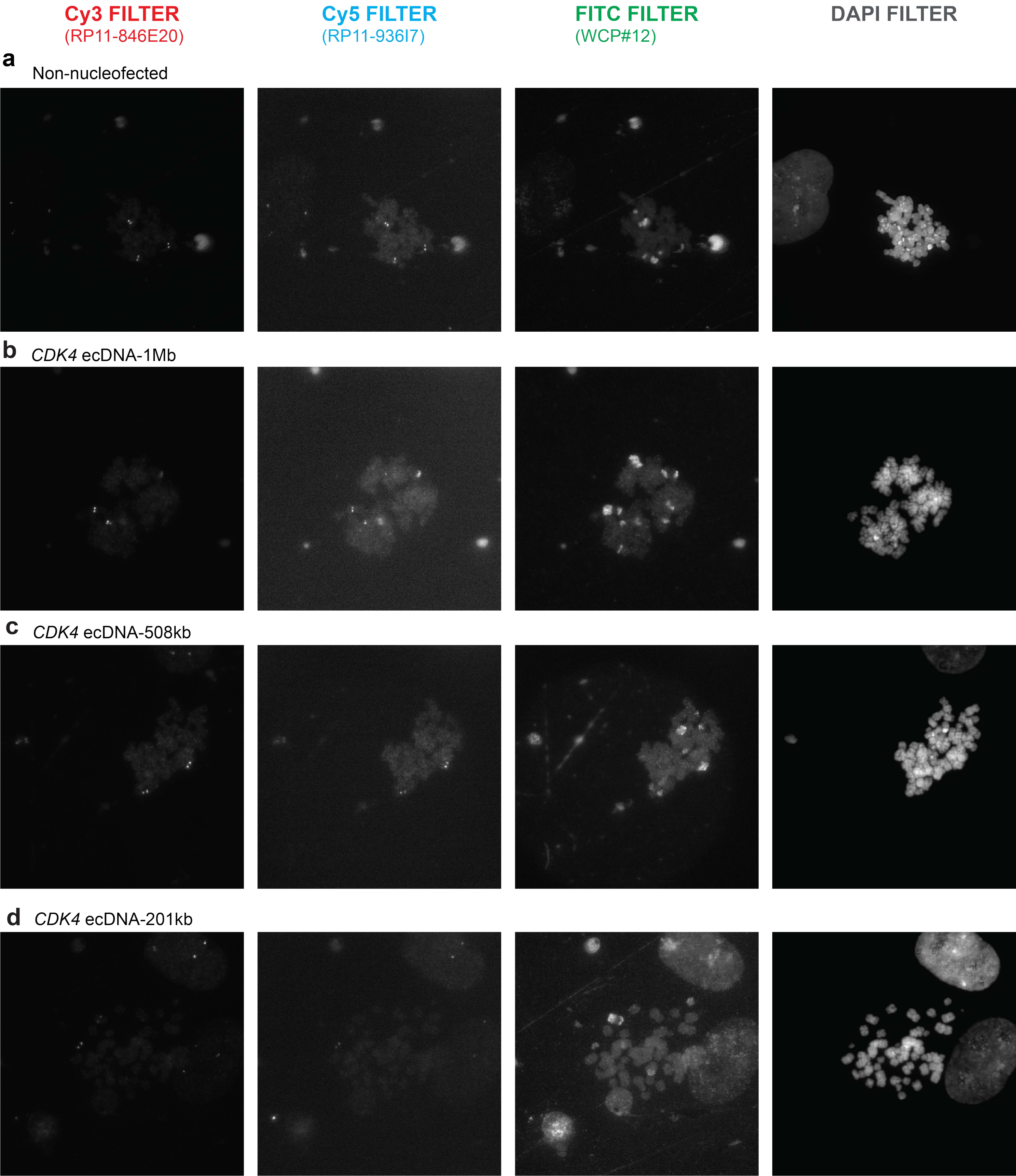

### Figure S24

**U2OS:**

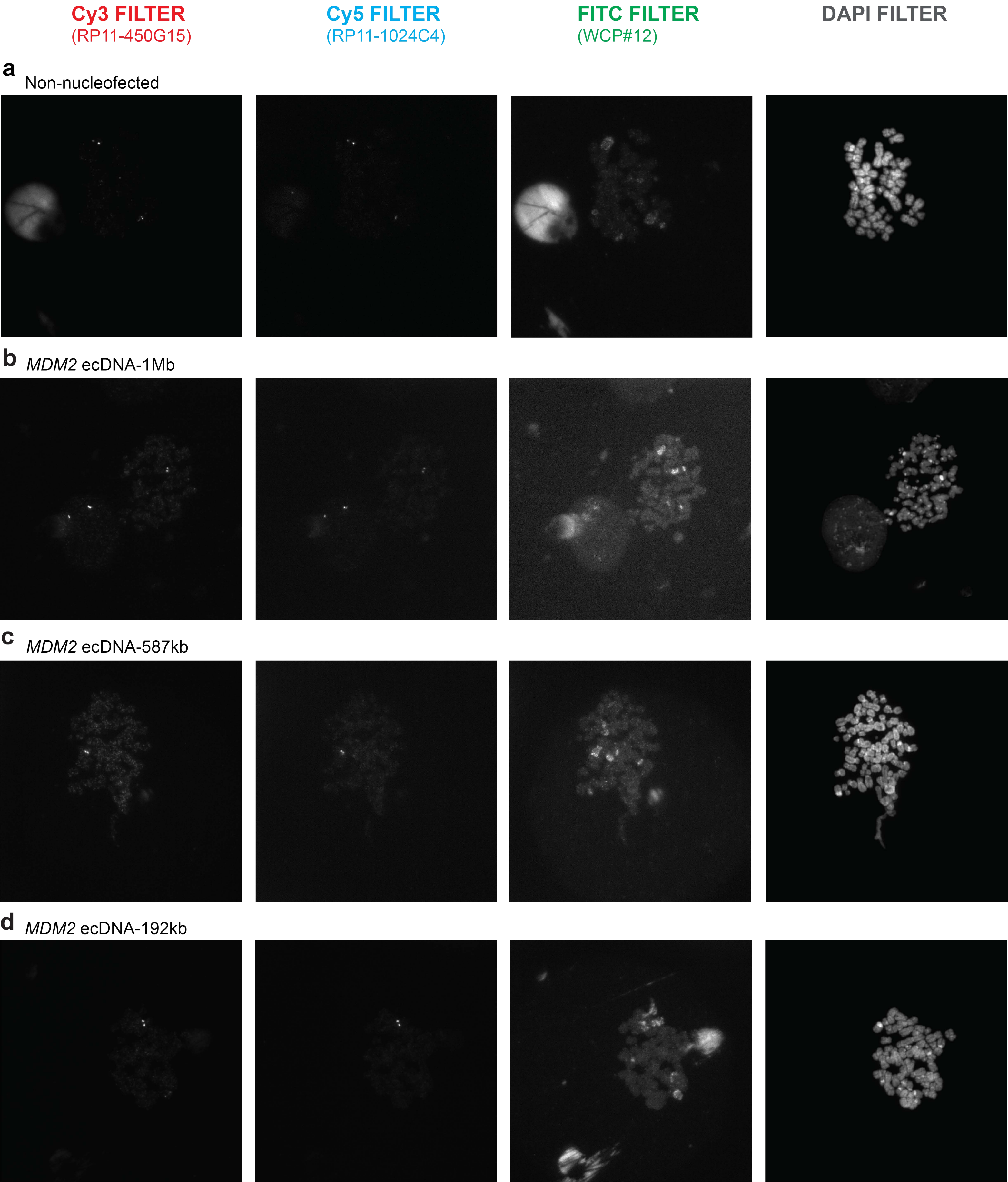

#### Figure S25

**U2OS:**

**a** **Cy3 FILTER** (RP11-846E20) | **Cy5 FILTER** (RP11-936I7) | **FITC FILTER** (WCP#12) | **DAPI FILTER**

*CDK4* ecDNA-508kb

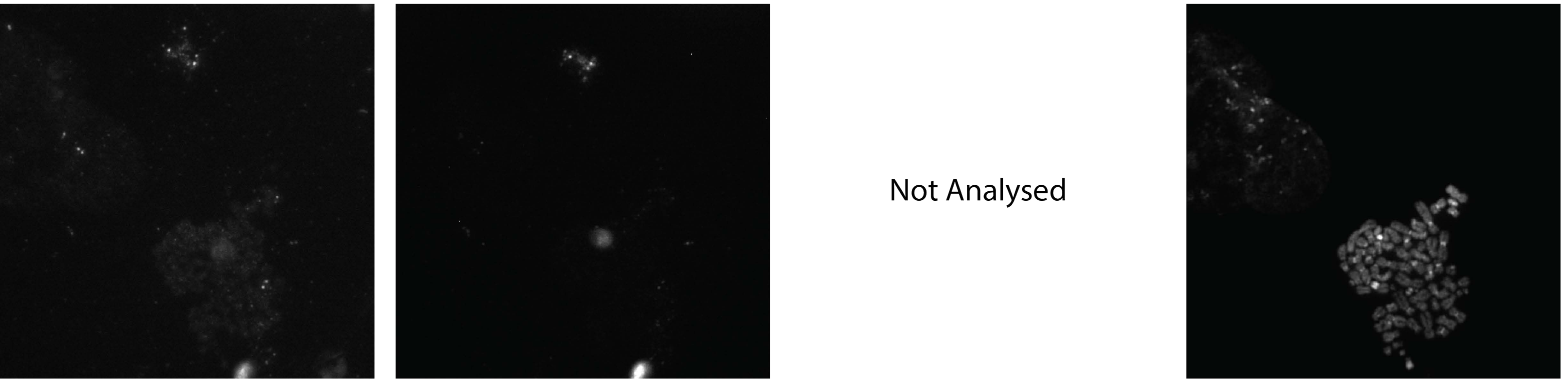


**b** **Cy3 FILTER** (RP11-846E20) | **Cy5 FILTER** (RP11-936I7) | **FITC FILTER** (WCP#12) | **DAPI FILTER**

*CDK4* ecDNA-201kb

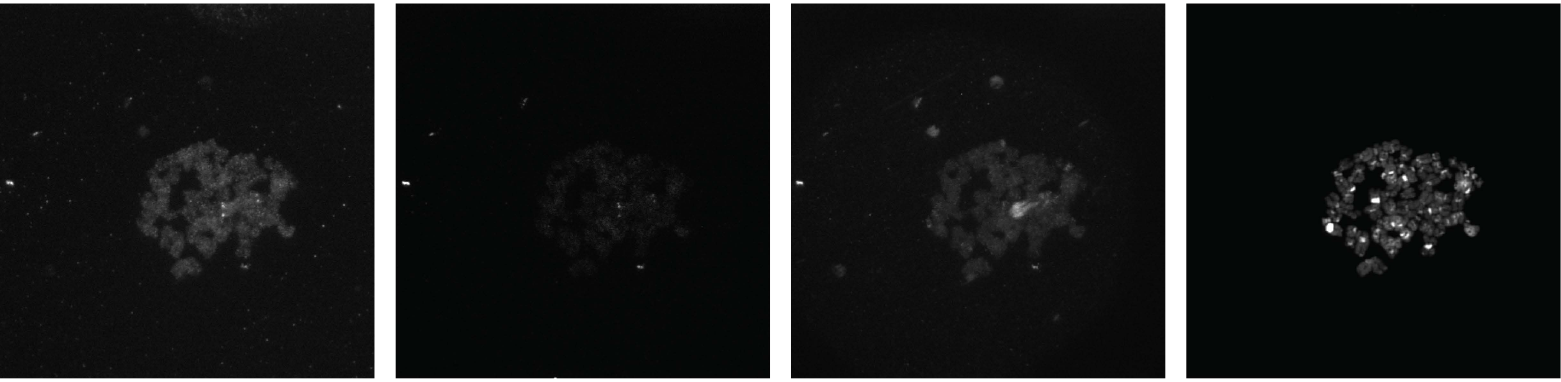

**c** **Cy3 FILTER** (RP11-450G15) | **Cy5 FILTER** (RP11-1024C4) | **FITC FILTER** (WCP#12) | **DAPI FILTER**

*MDM2* ecDNA-192kb

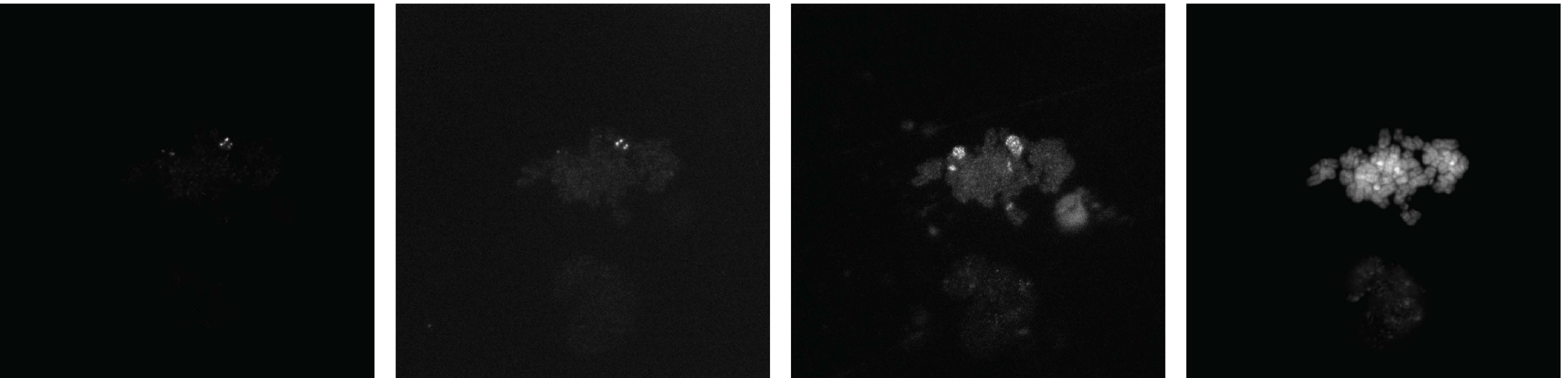

Fibroblast:

Figure S26

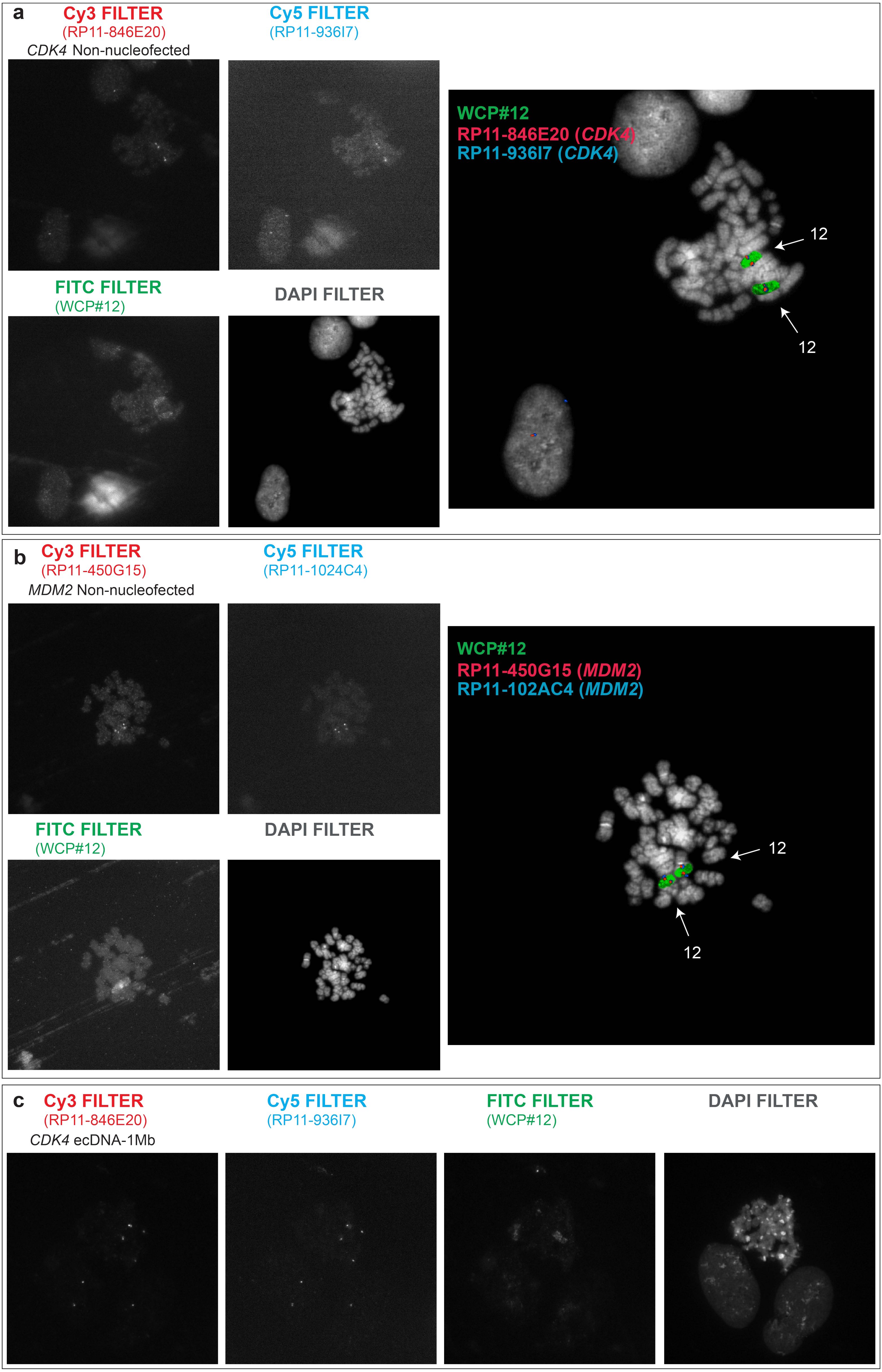

### Figure S27

**Palbociclib treated U2OS:**

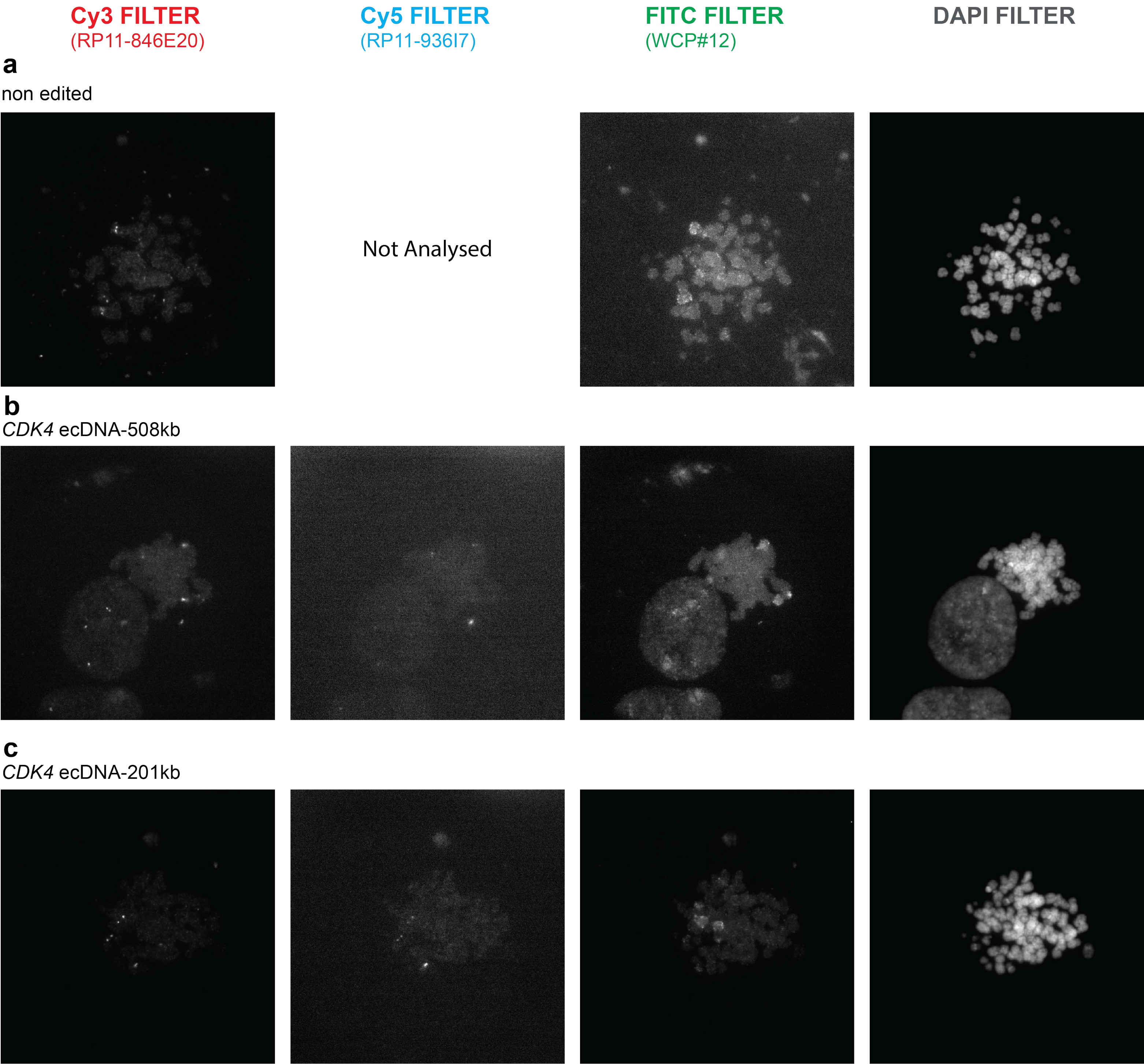

**Figure S28**

**Colcemid treated MCF7:**

**Cy3 FILTER** (RP11-831P22)

**FITC FILTER** (WCP7)

**DAPI FILTER**

**a**

non edited

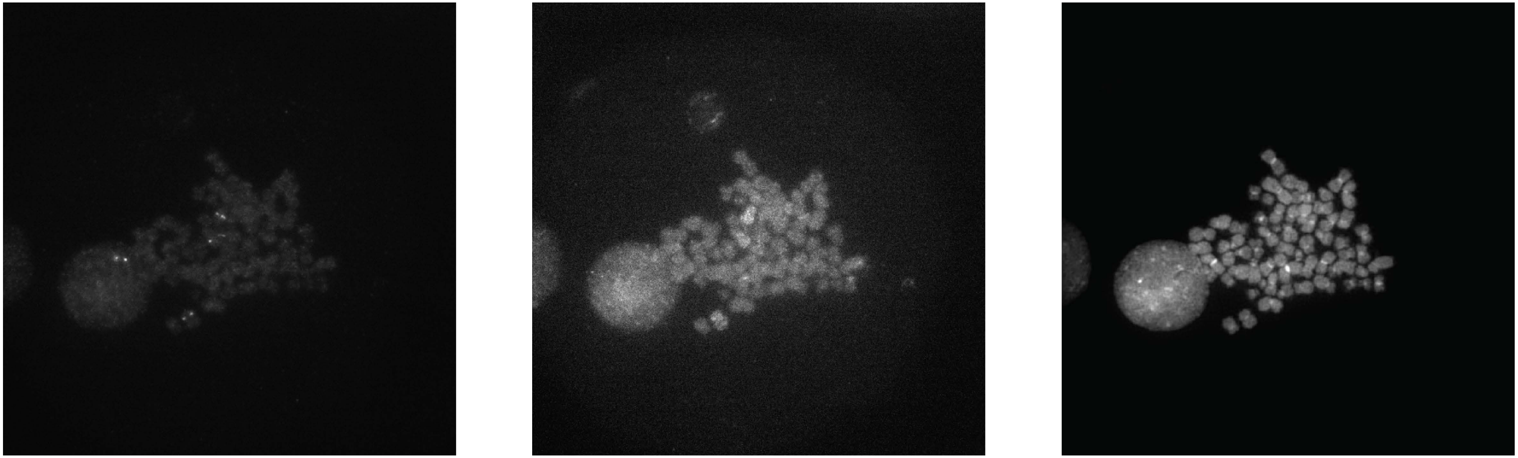

**b**

*ABCB1* ecDNA- 244kb

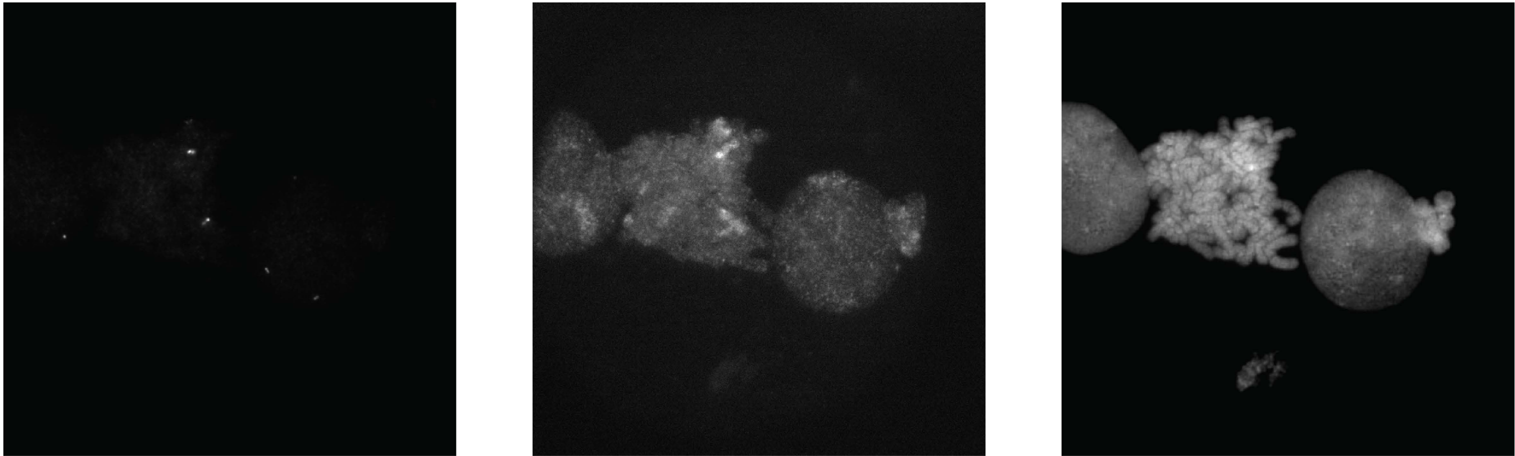

### Figure S29

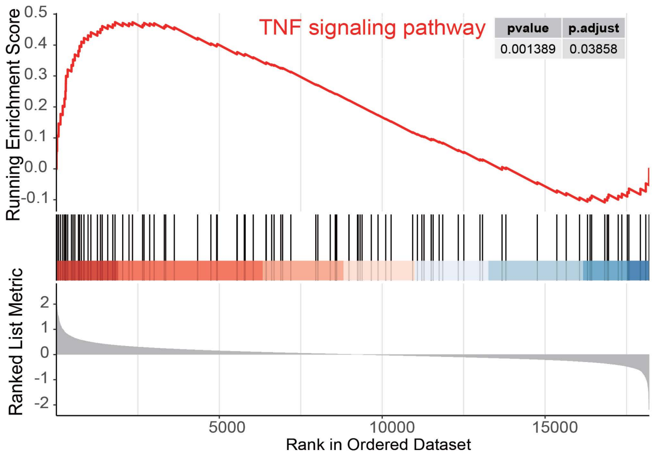
TNF signaling pathway
pvalue
p.adjust
0.001389
0.03858
Running Enrichment Score
0.5
0.4
0.3
0.2
0.1
0.0
-0.1
Ranked List Metric
2
1
0
-1
-2
5000
10000
15000
Rank in Ordered Dataset

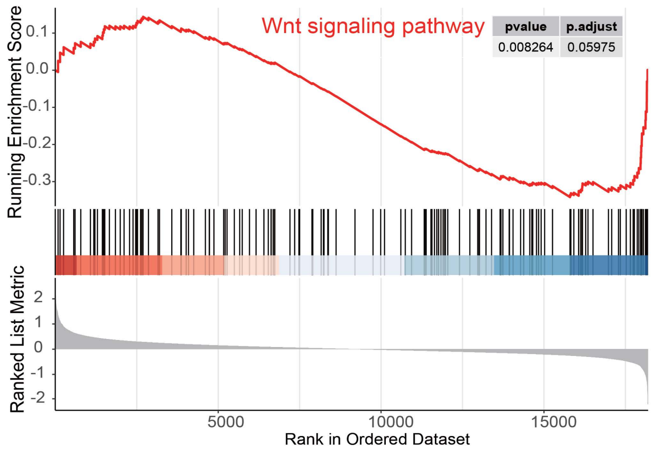
Wnt signaling pathway
pvalue
p.adjust
0.008264
0.05975
Running Enrichment Score
0.1
0.0
-0.1
-0.2
-0.3
Ranked List Metric
2
1
0
-1
-2
5000
10000
15000
Rank in Ordered Dataset

### Figure S30

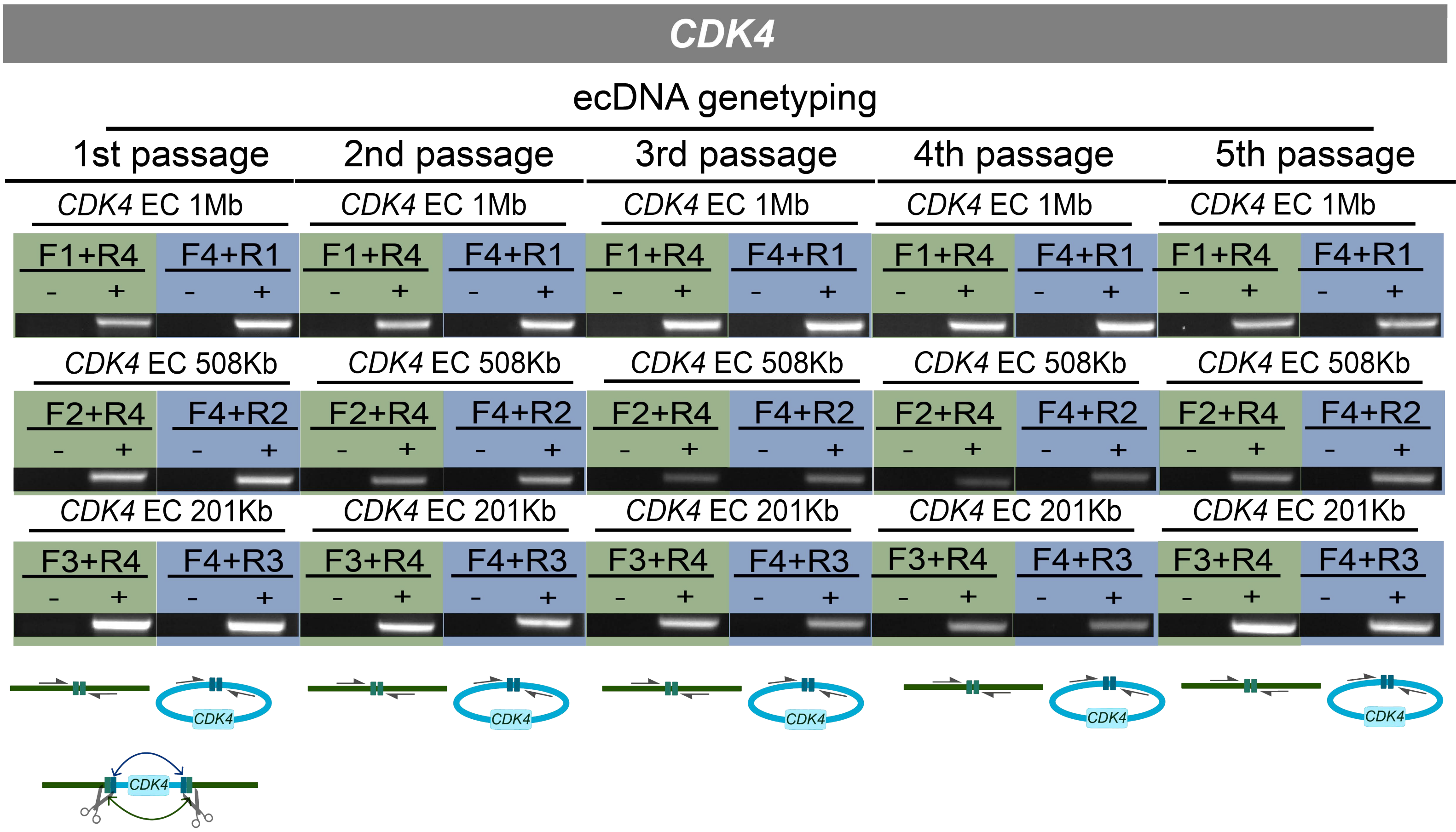


**CRISPR-C untreated: (-)** **CRISPR-C treated: (+)**

### Figure S31

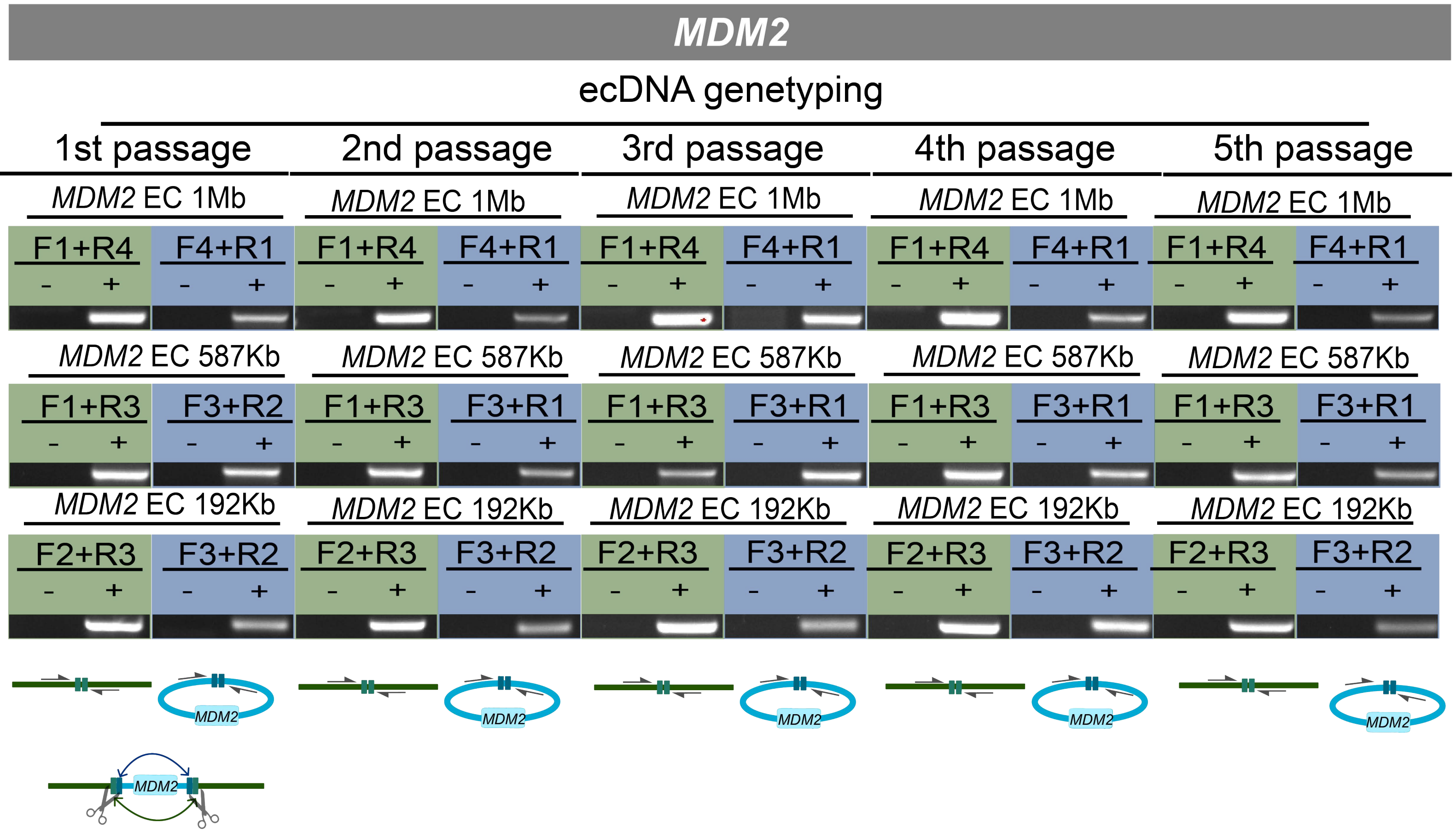


**CRISPR-C untreated: (-)** **CRISPR-C treated: (+)**

### Figure S32

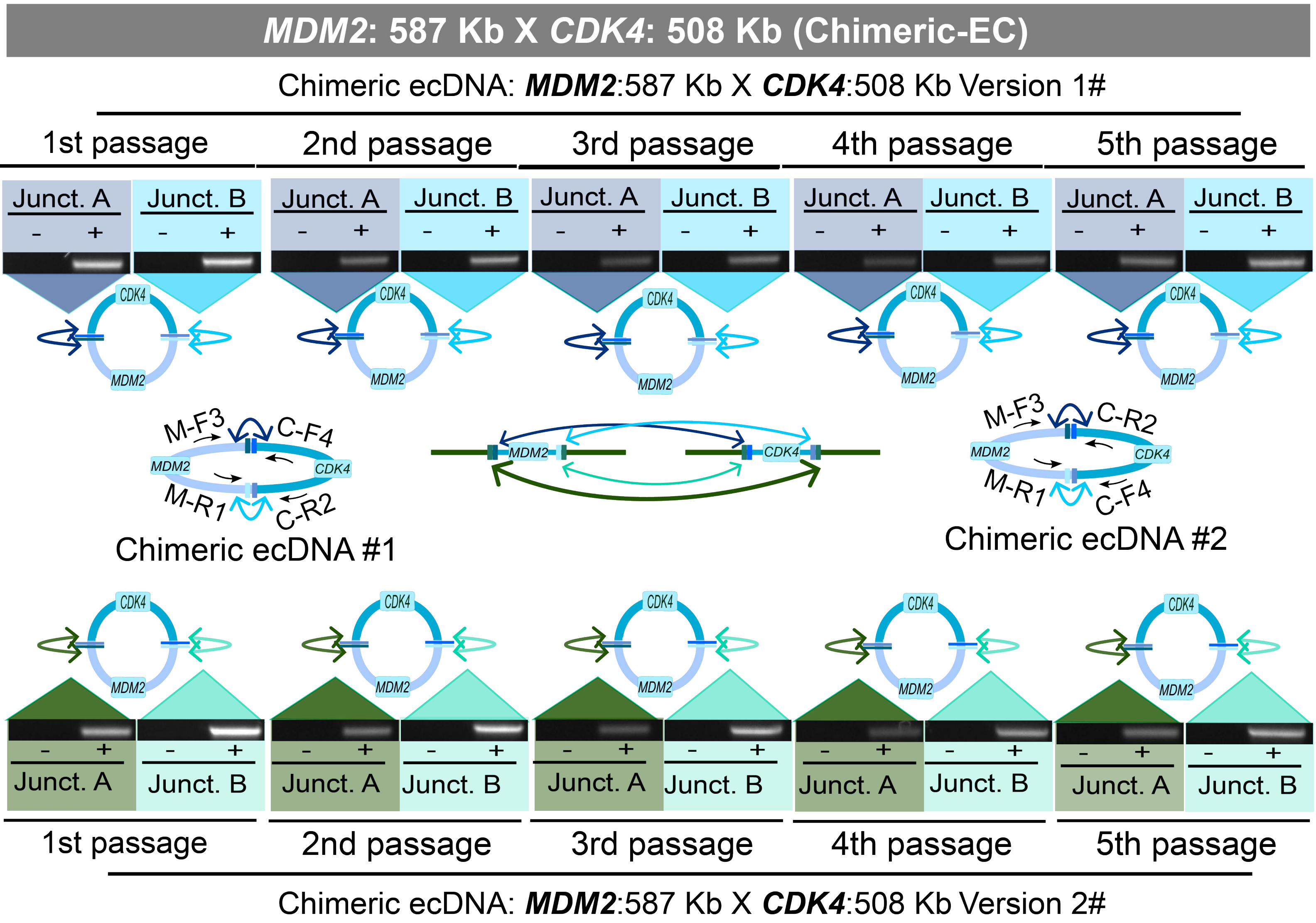


**CRISPR-C untreated: (-)** **CRISPR-C treated: (+)**

### Figure S33

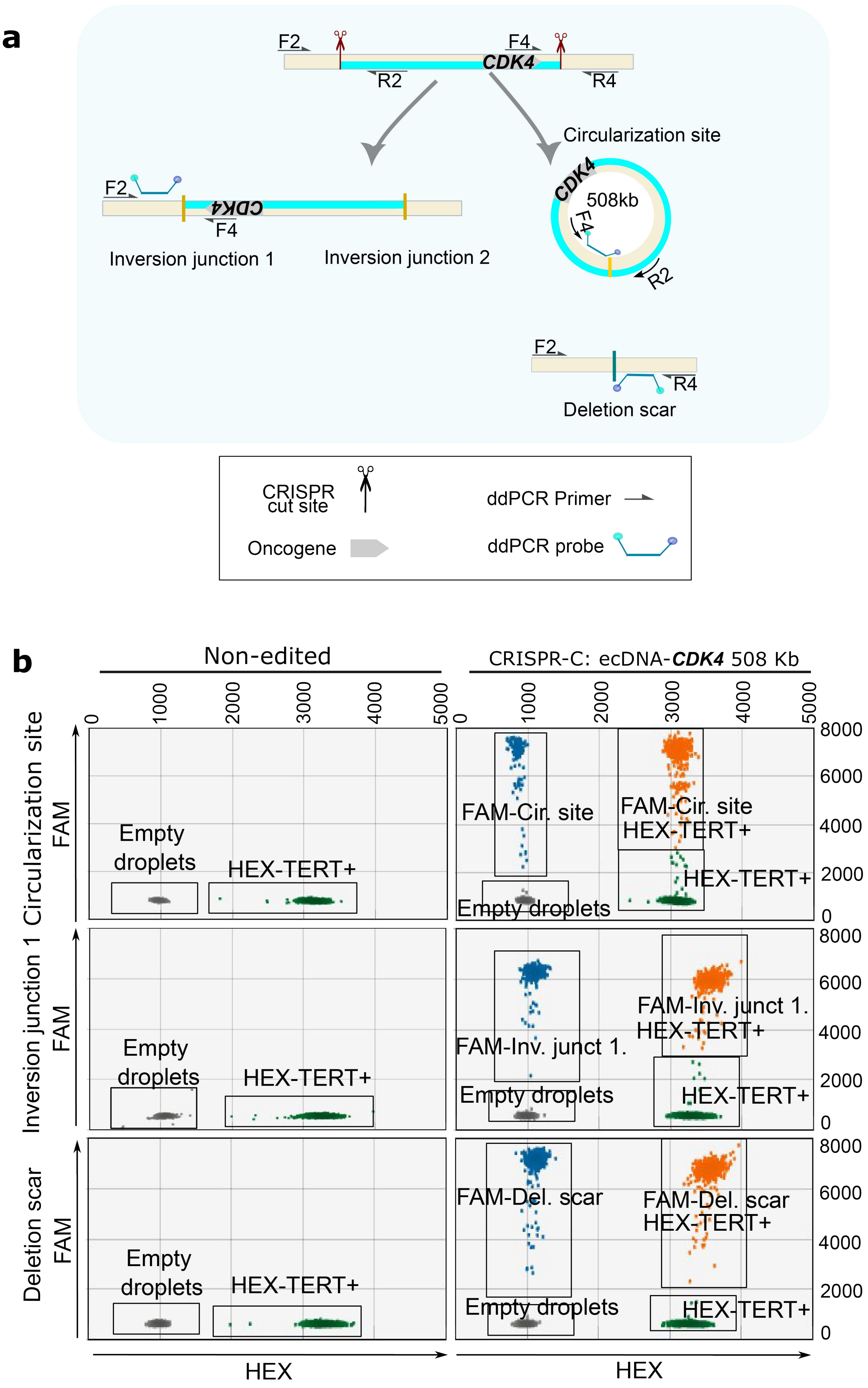
a
F2
R2
F4
CDK4
R4
Circularization site
CDK4
508kb
F4
R2
F2
CDK4
F4
Inversion junction 1
Inversion junction 2
F2
R4
Deletion scar
CRISPR cut site
ddPCR Primer
Oncogene
ddPCR probe
b
Non-edited
CRISPR-C: ecDNA-CDK4 508 Kb
Circularization site
FAM
Empty droplets
HEX-TERT+
FAM-Cir. site
FAM-Cir. site HEX-TERT+
HEX-TERT+
Empty droplets
Inversion junction 1
FAM
Empty droplets
HEX-TERT+
FAM-Inv. junct 1.
FAM-Inv. junct 1. HEX-TERT+
Empty droplets
HEX-TERT+
Deletion scar
FAM
Empty droplets
HEX-TERT+
FAM-Del. scar
FAM-Del. scar HEX-TERT+
Empty droplets
HEX-TERT+
HEX
HEX
