## Supplementary material for "Generative Modelling of Oncogene-carrying Extrachromosomal Circular DNA Biogenesis and Dynamics in Cells": Source data for gel images in both figures and supplementary figures

### Souce data for gel images

### Figure 1c

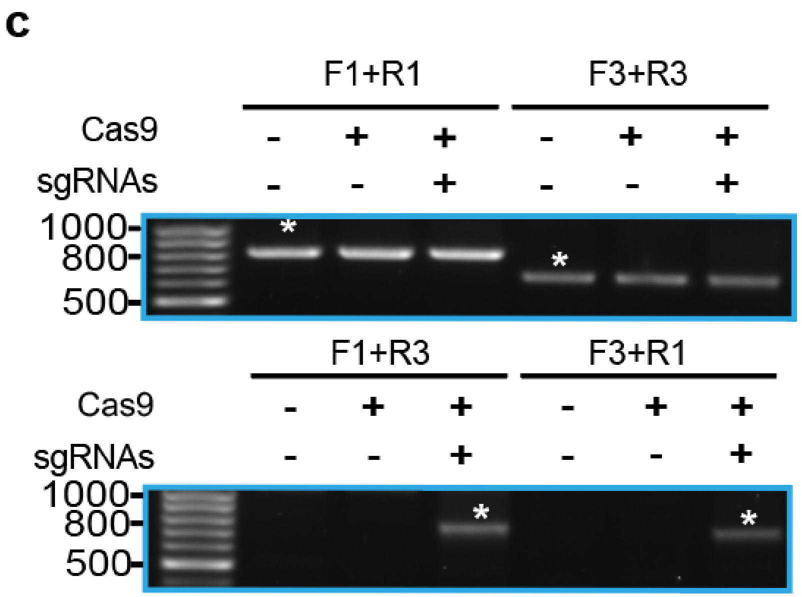
c
F1+R1
F3+R3
Cas9
sgRNAs
1000
800
500
F1+R3
F3+R1
Cas9
sgRNAs
1000
800
500

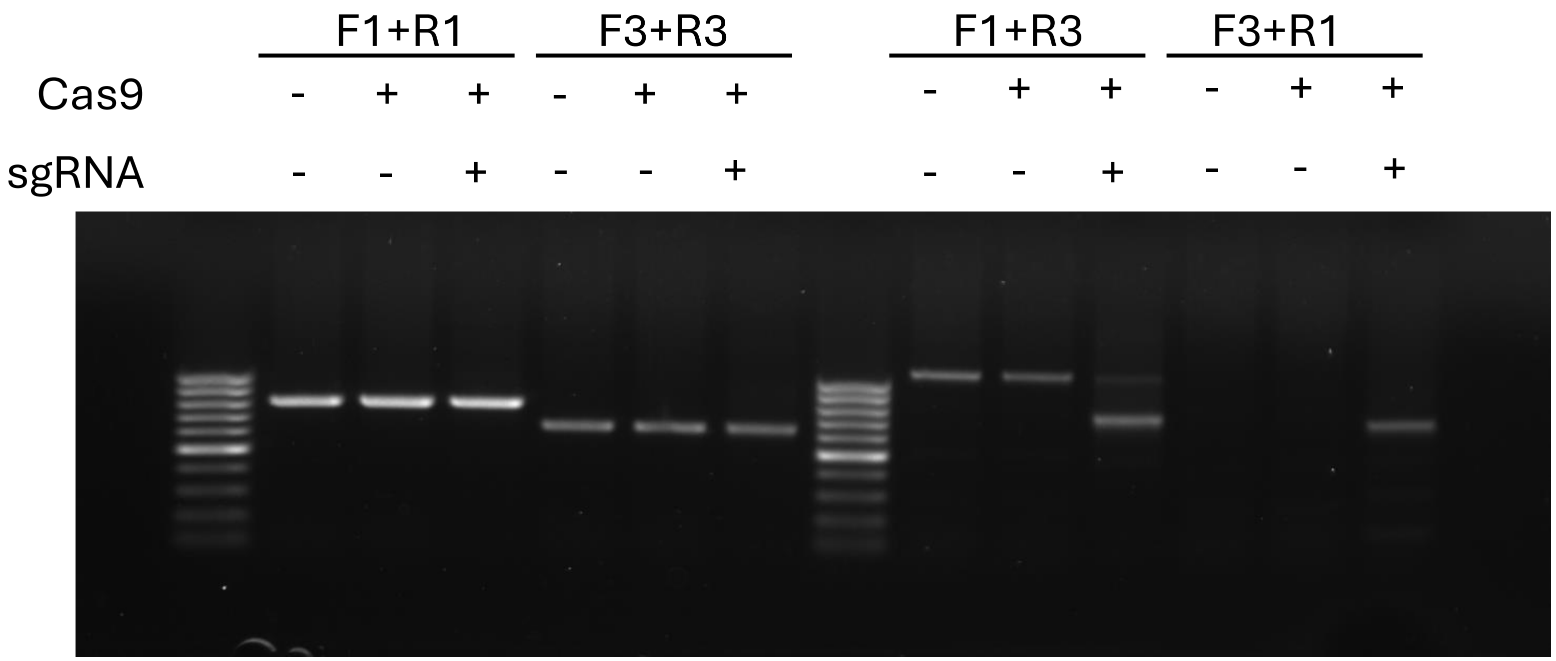
F1+R1
F3+R3
F1+R3
F3+R1
Cas9
sgRNA

### Figure 1e

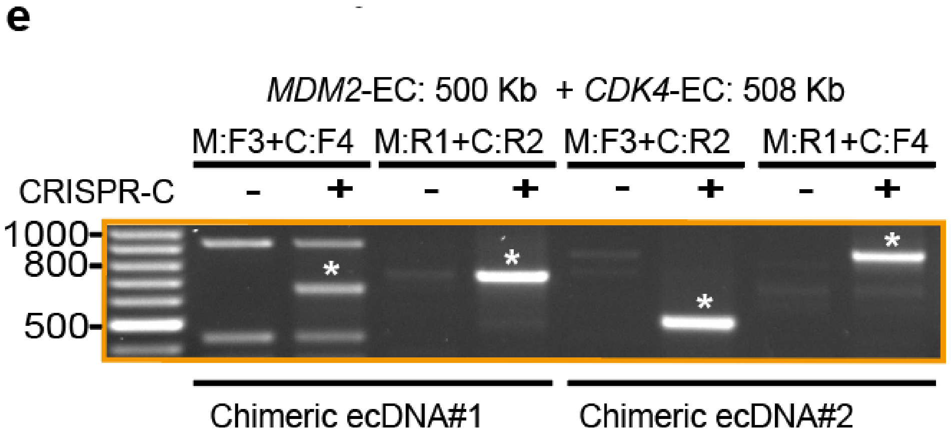
e
MDM2-EC: 500 Kb + CDK4-EC: 508 Kb
M:F3+C:F4
M:R1+C:R2
M:F3+C:R2
M:R1+C:F4
CRISPR-C
- + - + - + - +
1000
800
500
Chimeric ecDNA#1
Chimeric ecDNA#2

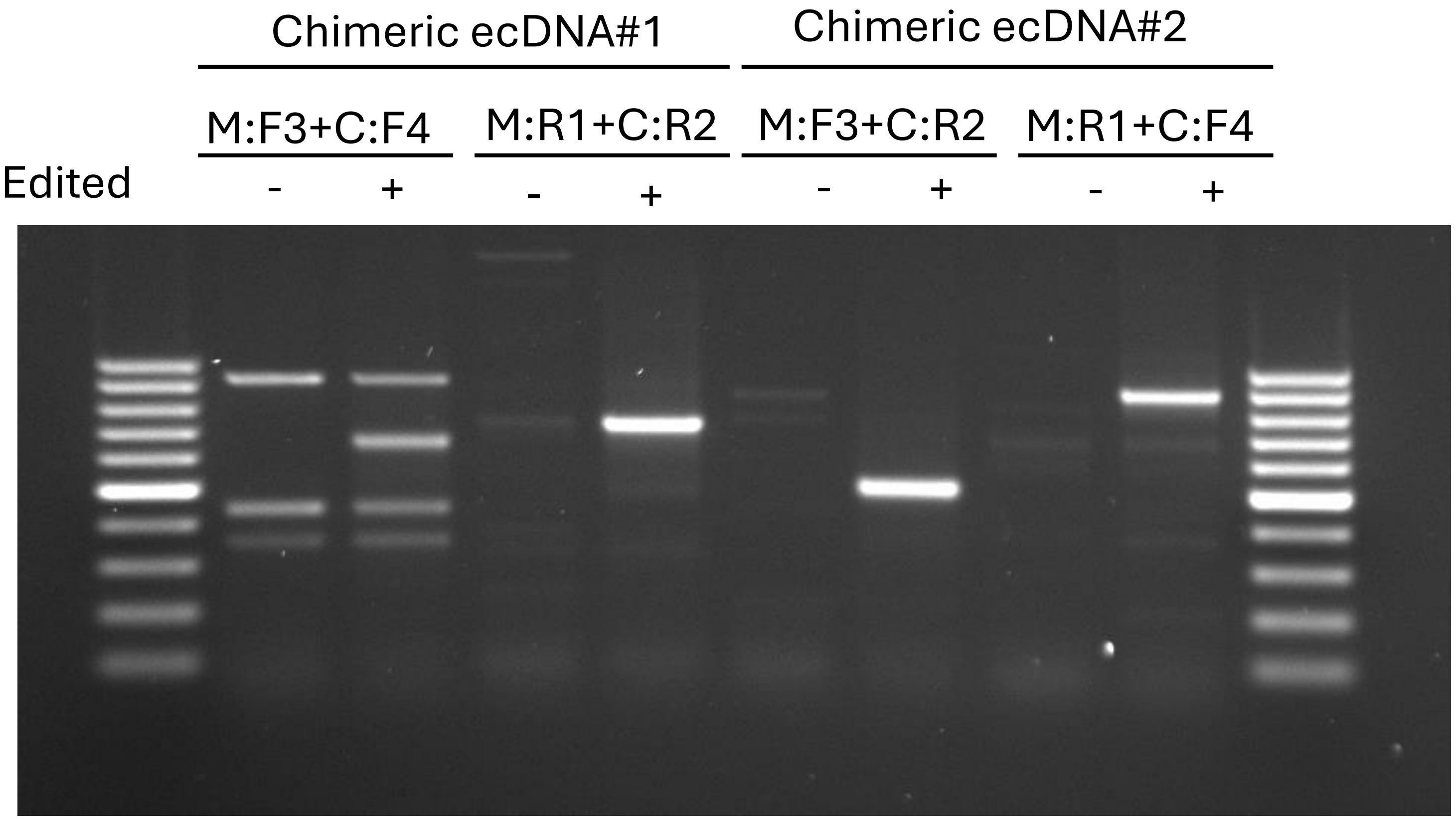
Chimeric ecDNA#1
Chimeric ecDNA#2
M:F3+C:F4
M:R1+C:R2
M:F3+C:R2
M:R1+C:F4
Edited
- + - + - + - +

### Figure 3a

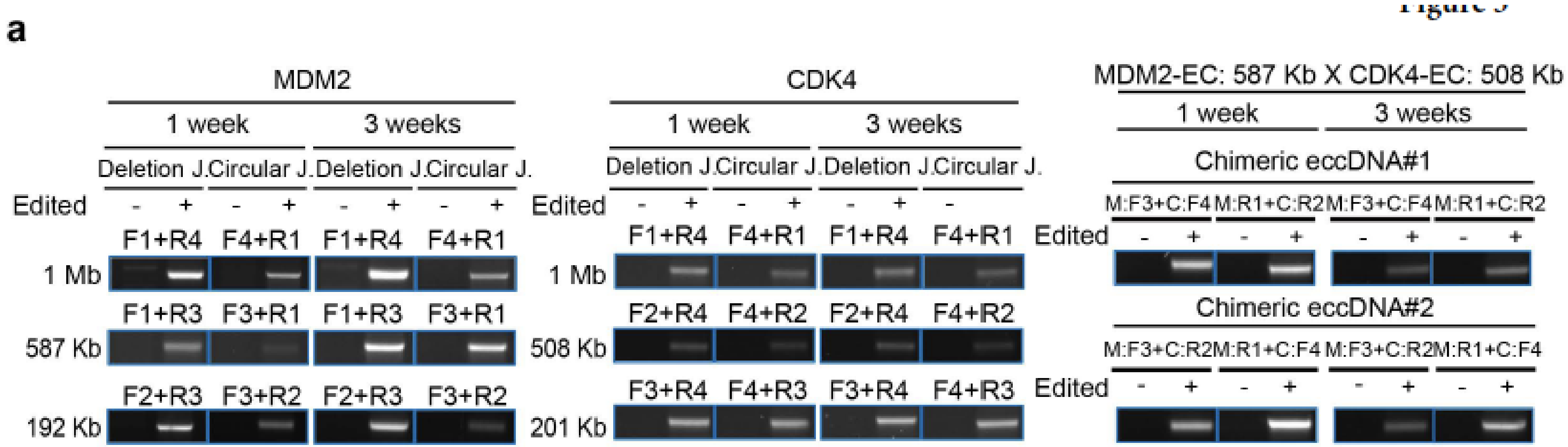

a
MDM2
1 week
3 weeks
Deletion J.
Circular J.
Edited
F1+R4
F4+R1
F1+R3
F3+R1
F2+R3
F3+R2
1 Mb
587 Kb
192 Kb
CDK4
F2+R4
F4+R2
F3+R4
F4+R3
508 Kb
201 Kb
MDM2-EC: 587 Kb X CDK4-EC: 508 Kb
Chimeric eccDNA#1
M:F3+C:F4
M:R1+C:R2
Chimeric eccDNA#2
M:F3+C:R2
M:R1+C:F4

### Figure 3a

#### *MDM2* 1 week 1Mb

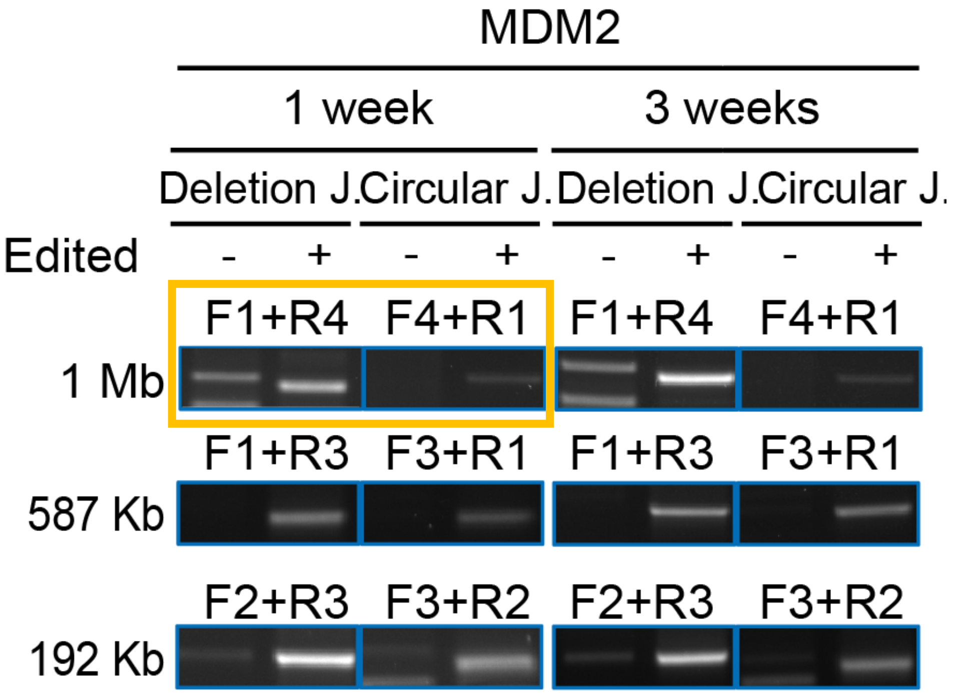


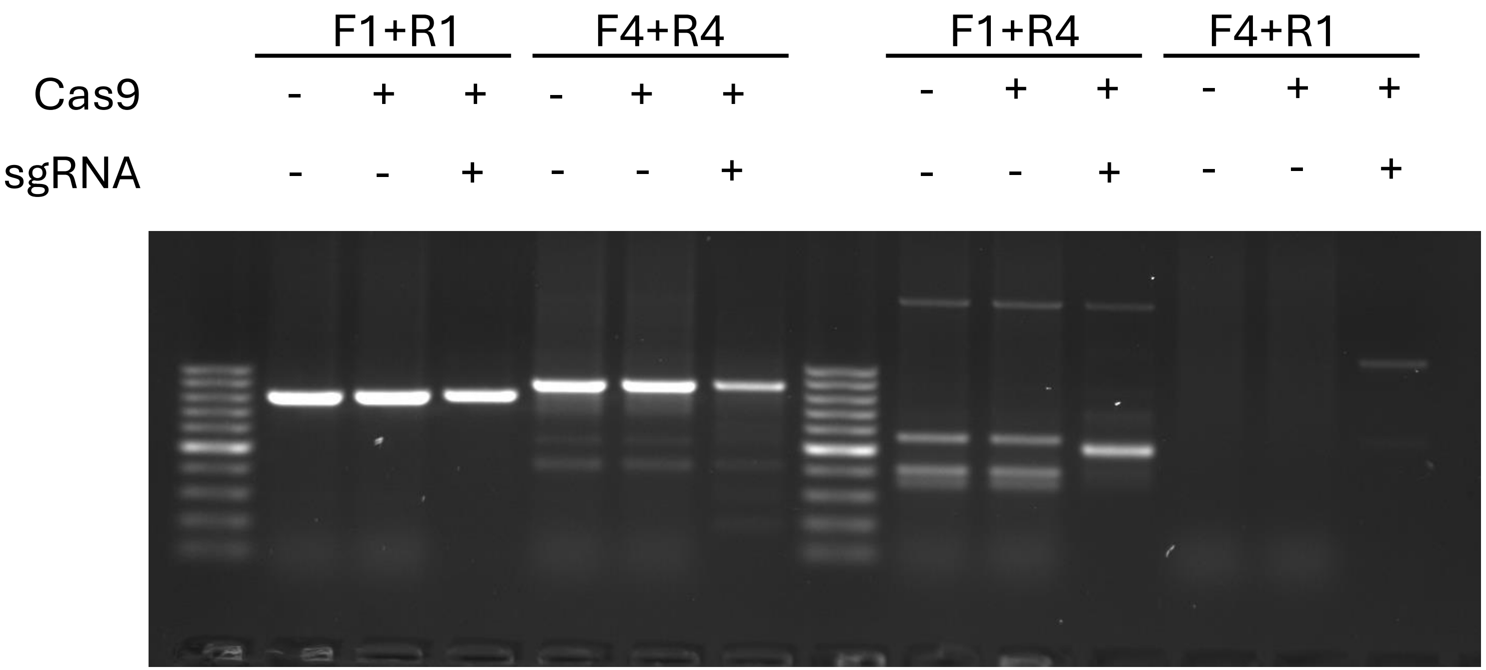

### Figure 3a

#### *MDM2* 1 week 587 Kb

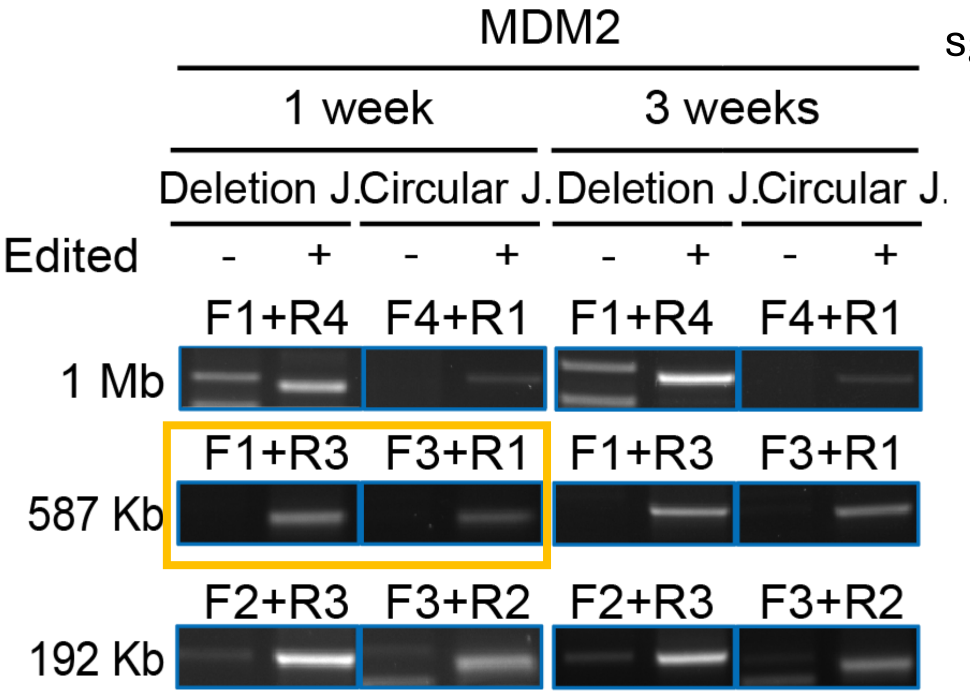


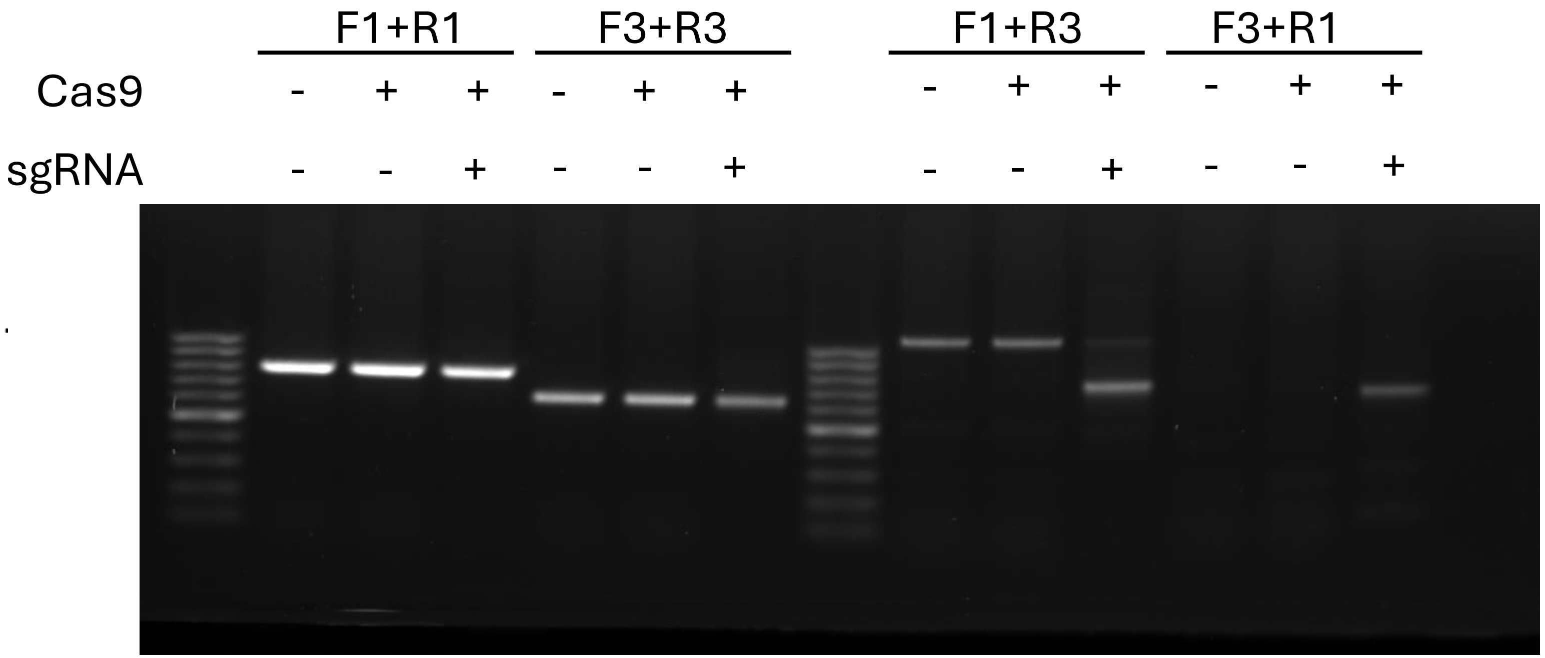

### Figure 3a

#### *MDM2* 1 week 192 Kb

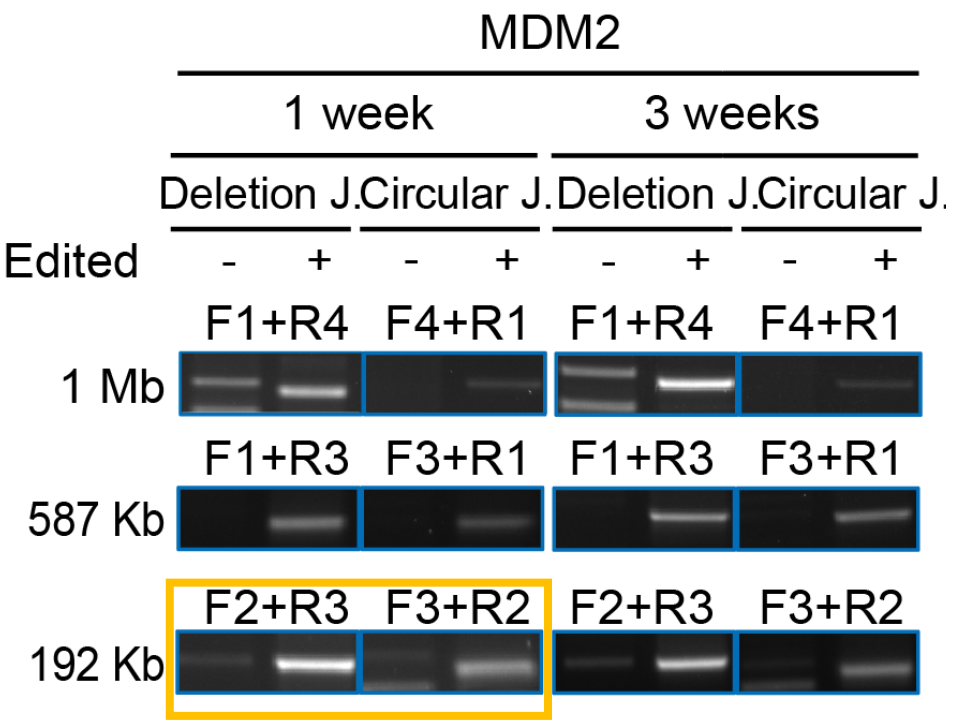


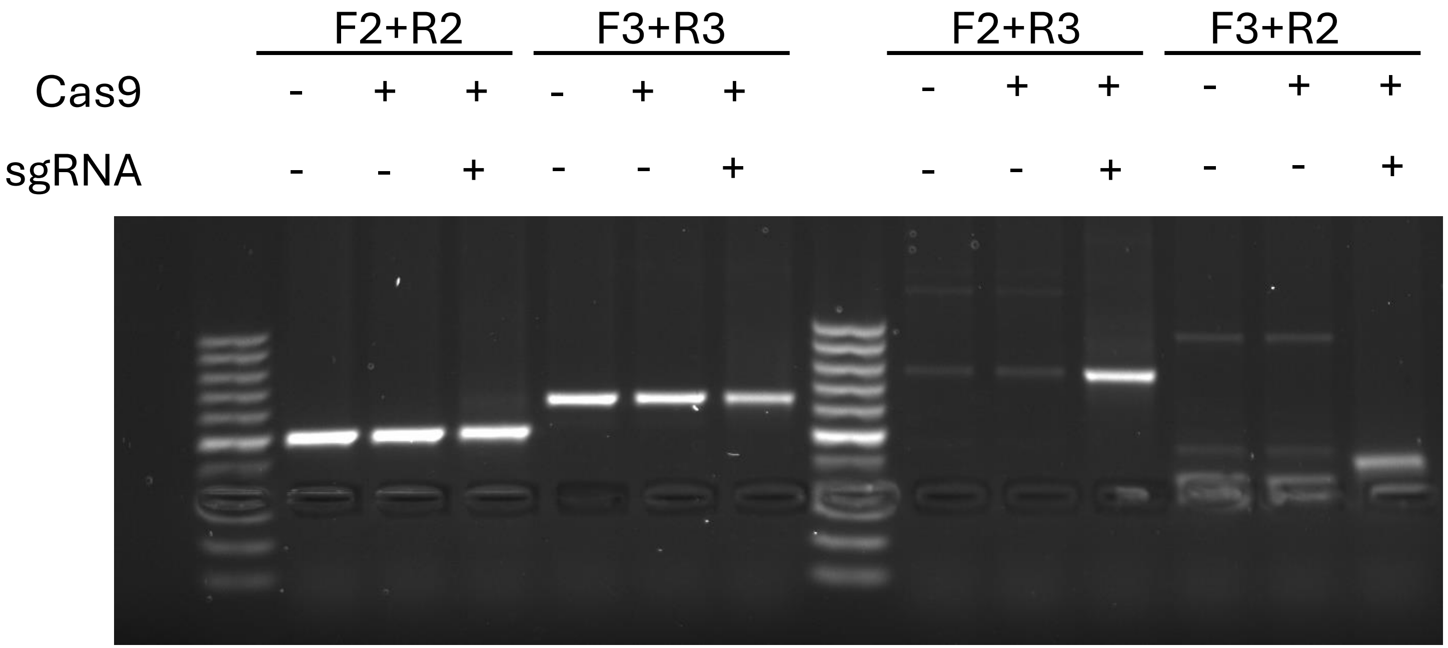

### Figure 3a

#### *MDM2* 3 weeks 1Mb

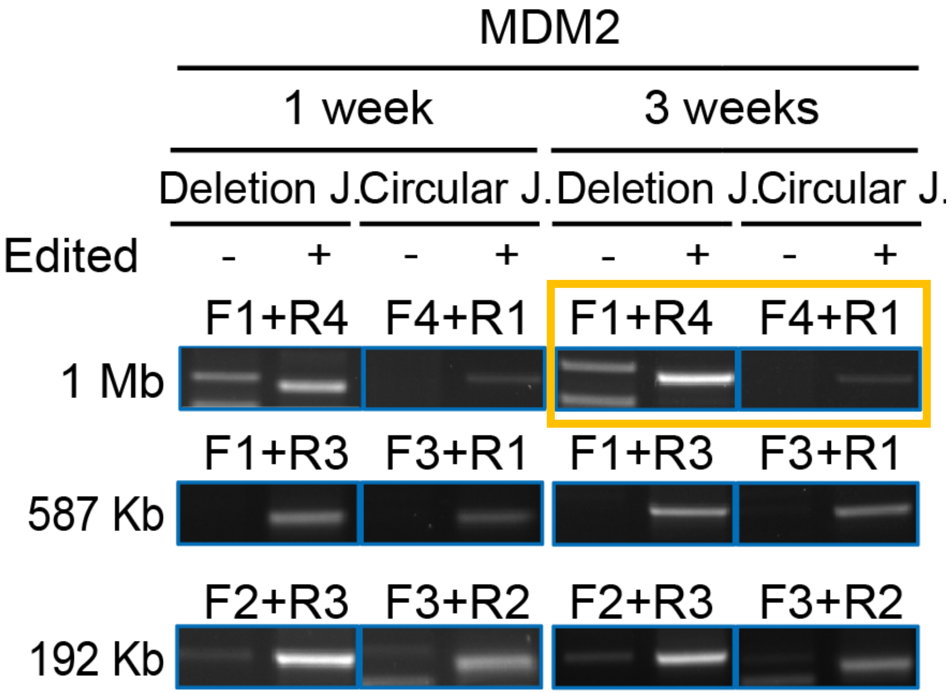


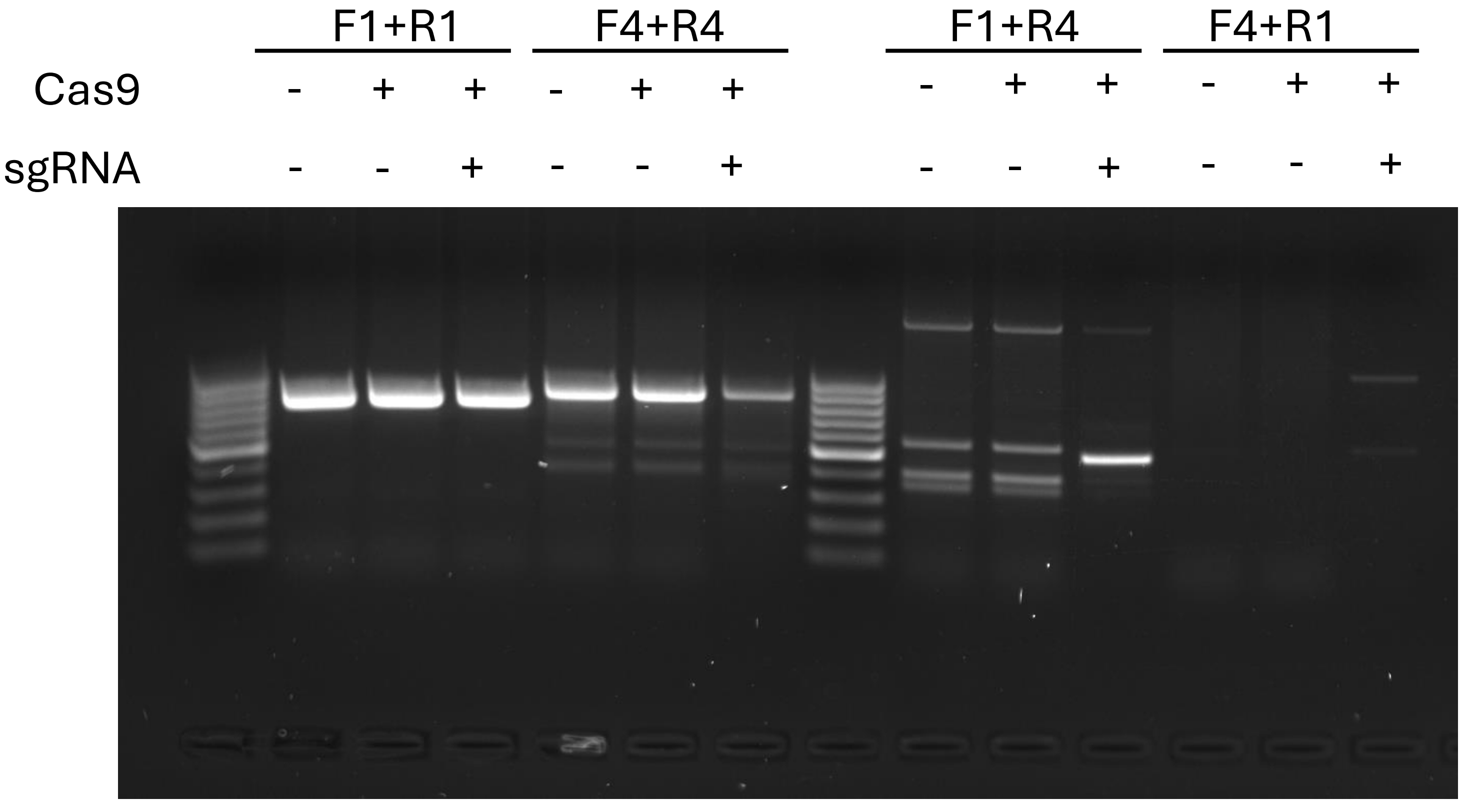

### Figure 3a

#### *MDM2* 3 weeks 587 Kb

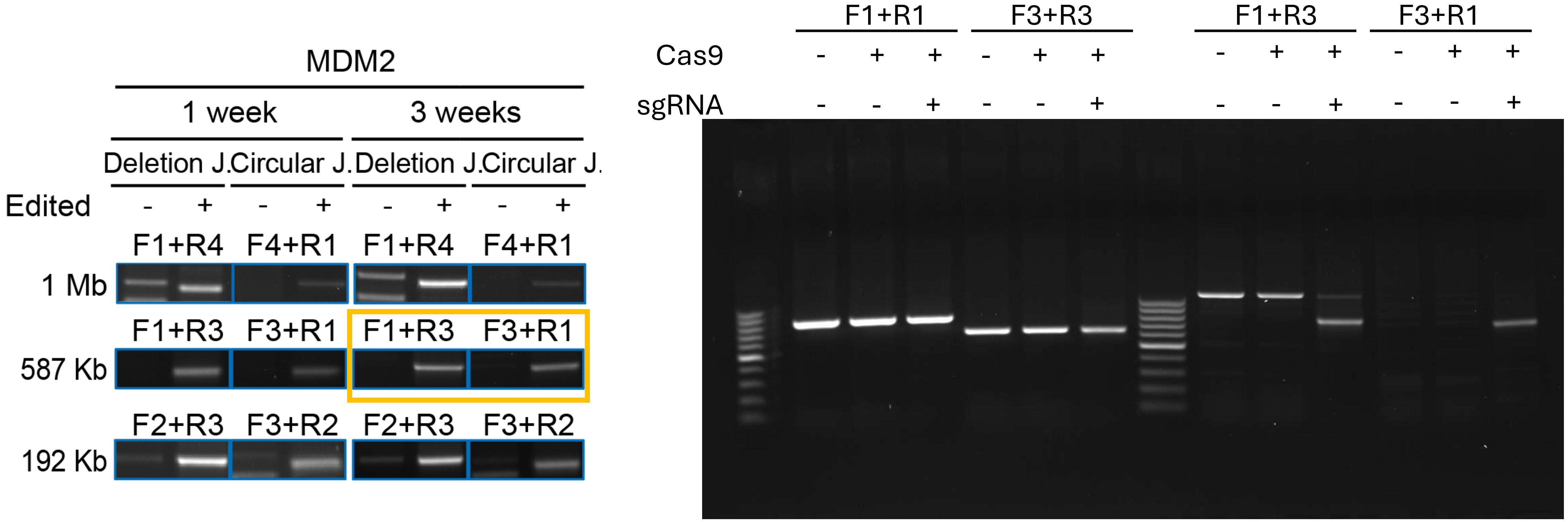

### Figure 3a

#### *MDM2* 3 weeks 192 Kb

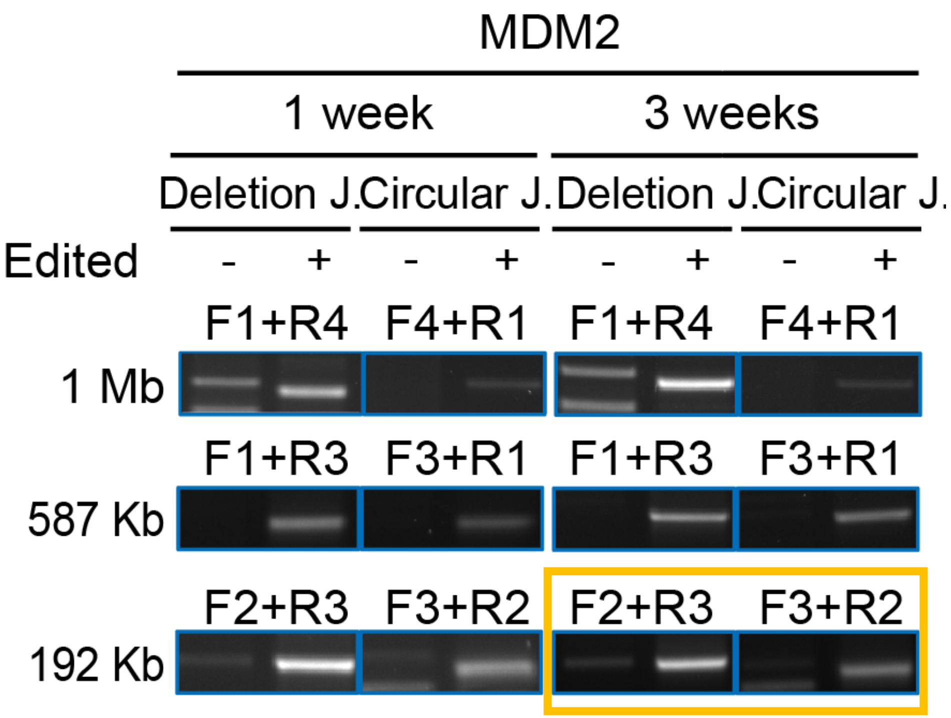


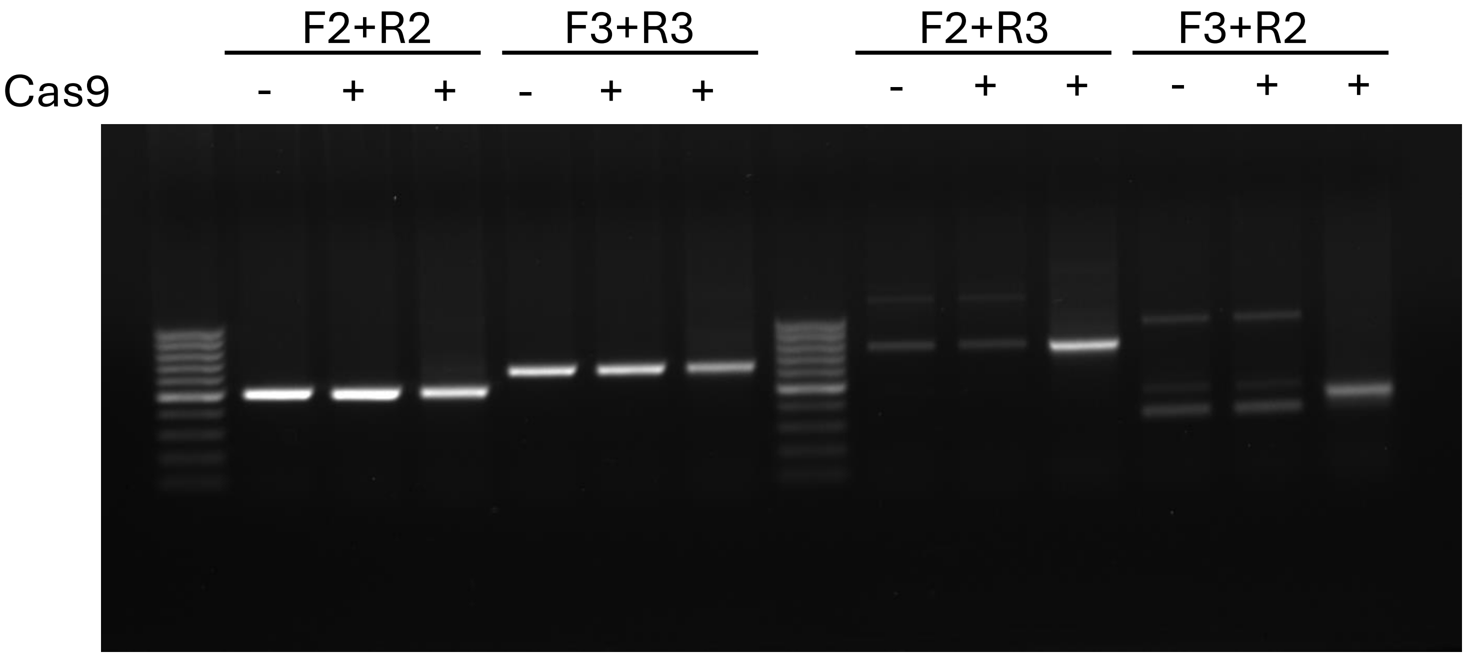

### Figure 3a

#### *CDK4* 1 week 1Mb

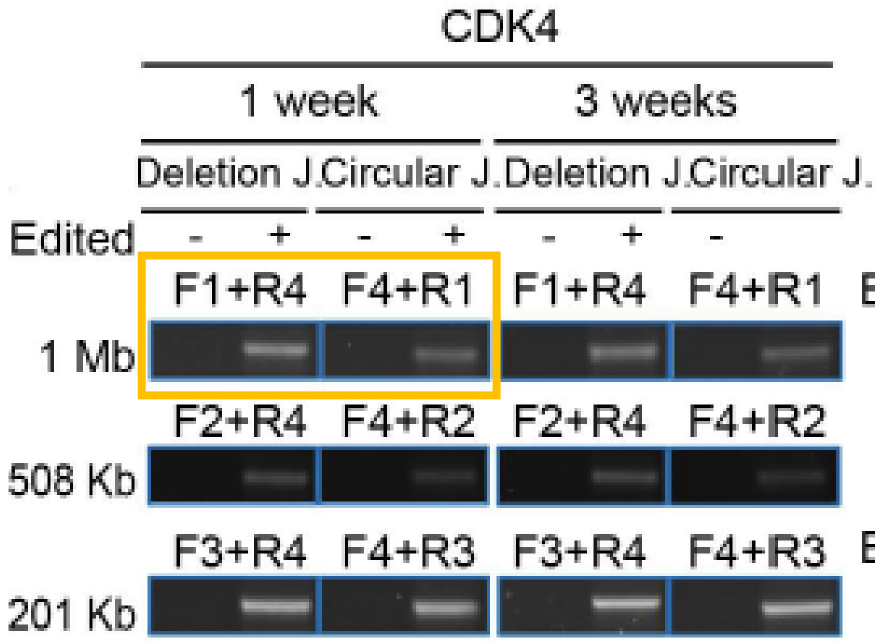


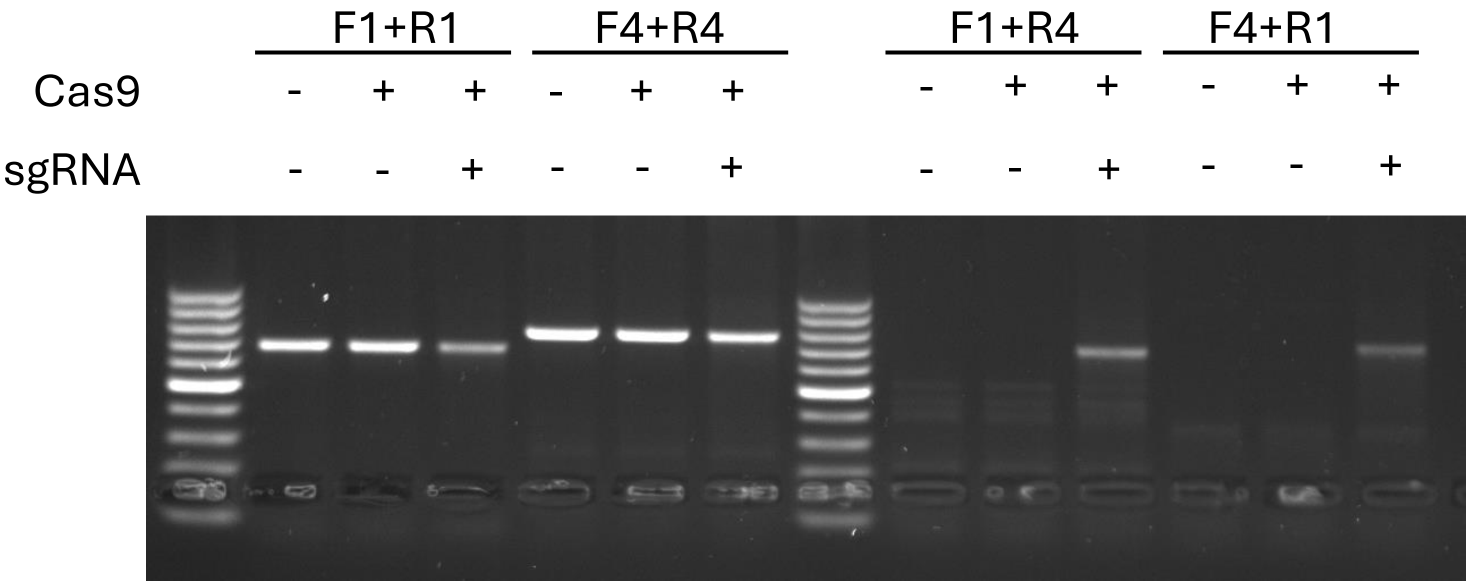

### Figure 3a

#### *CDK4* 1 week 508Kb

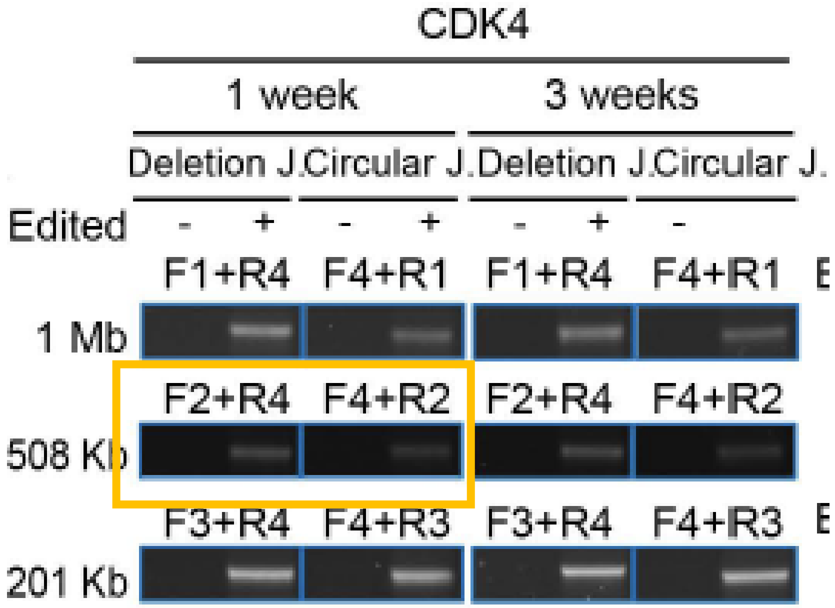


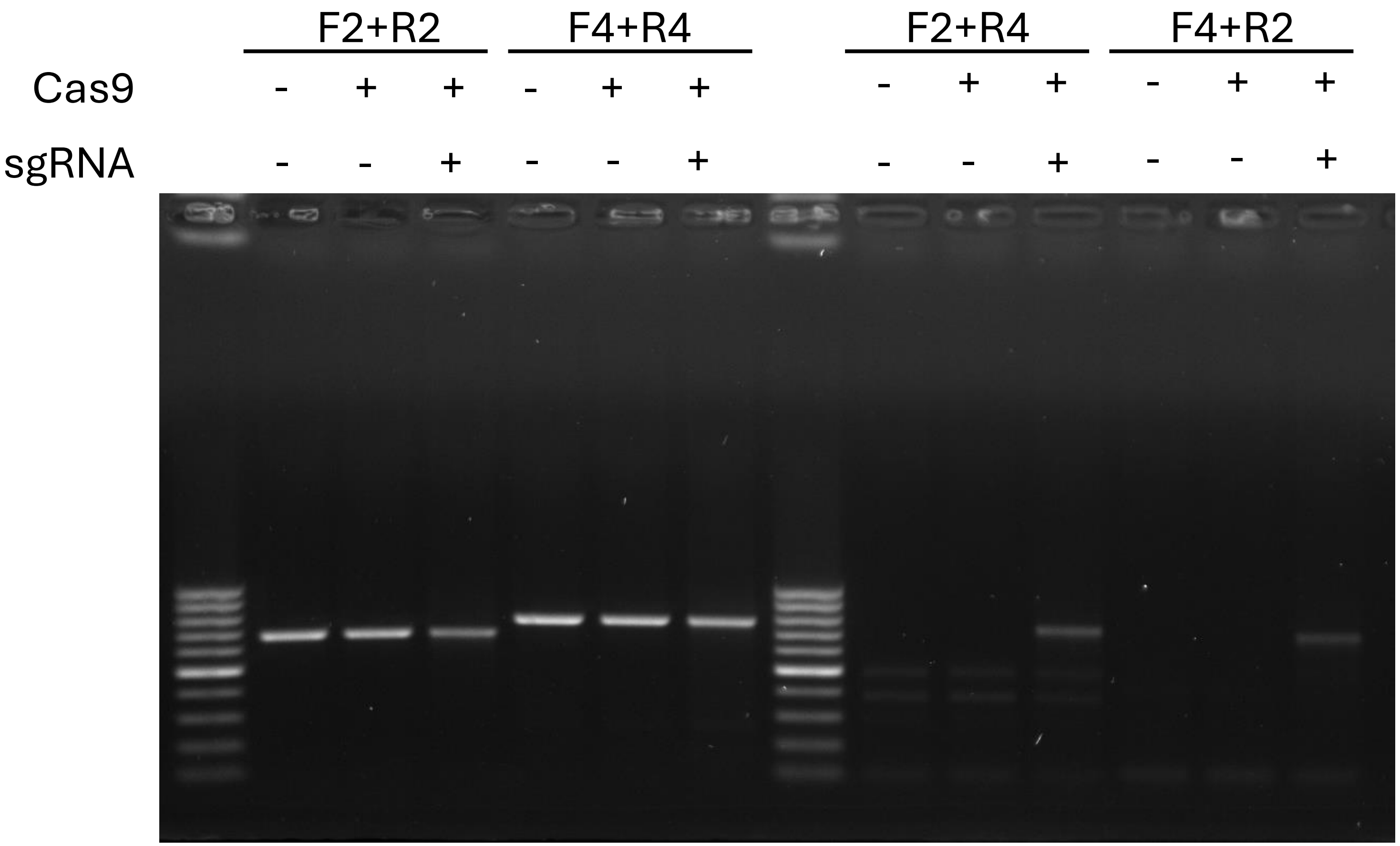

### Figure 3a

#### *CDK4* 1 week 201Kb

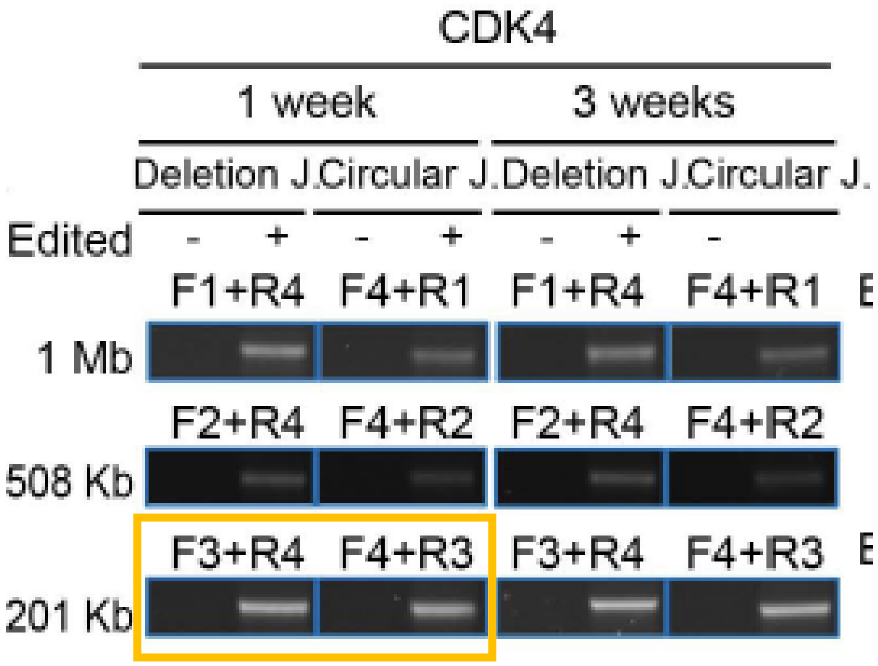


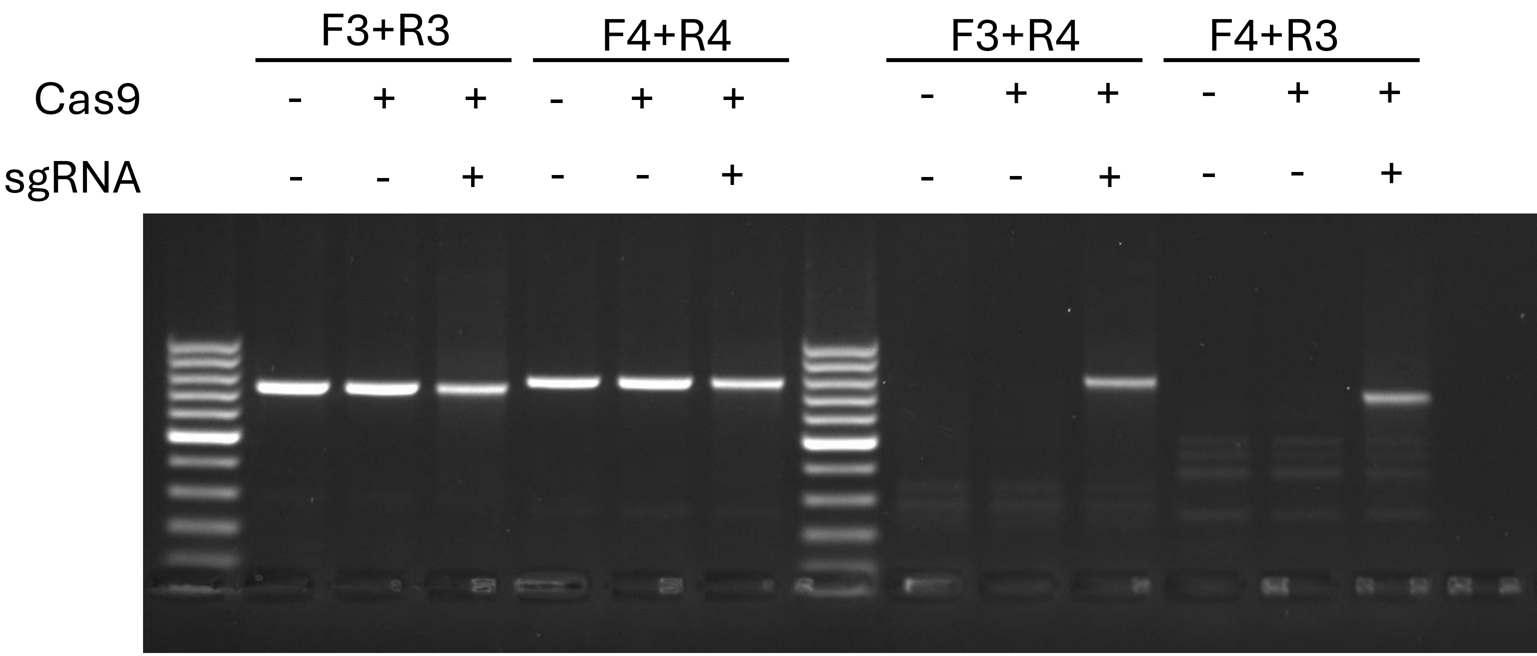

### Figure 3a

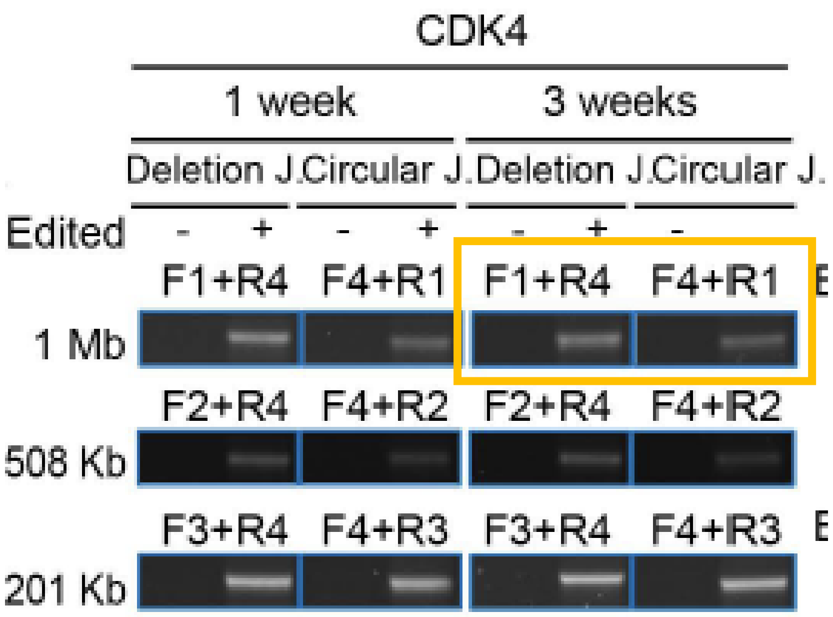


#### *CDK4* 3 weeks 1Mb

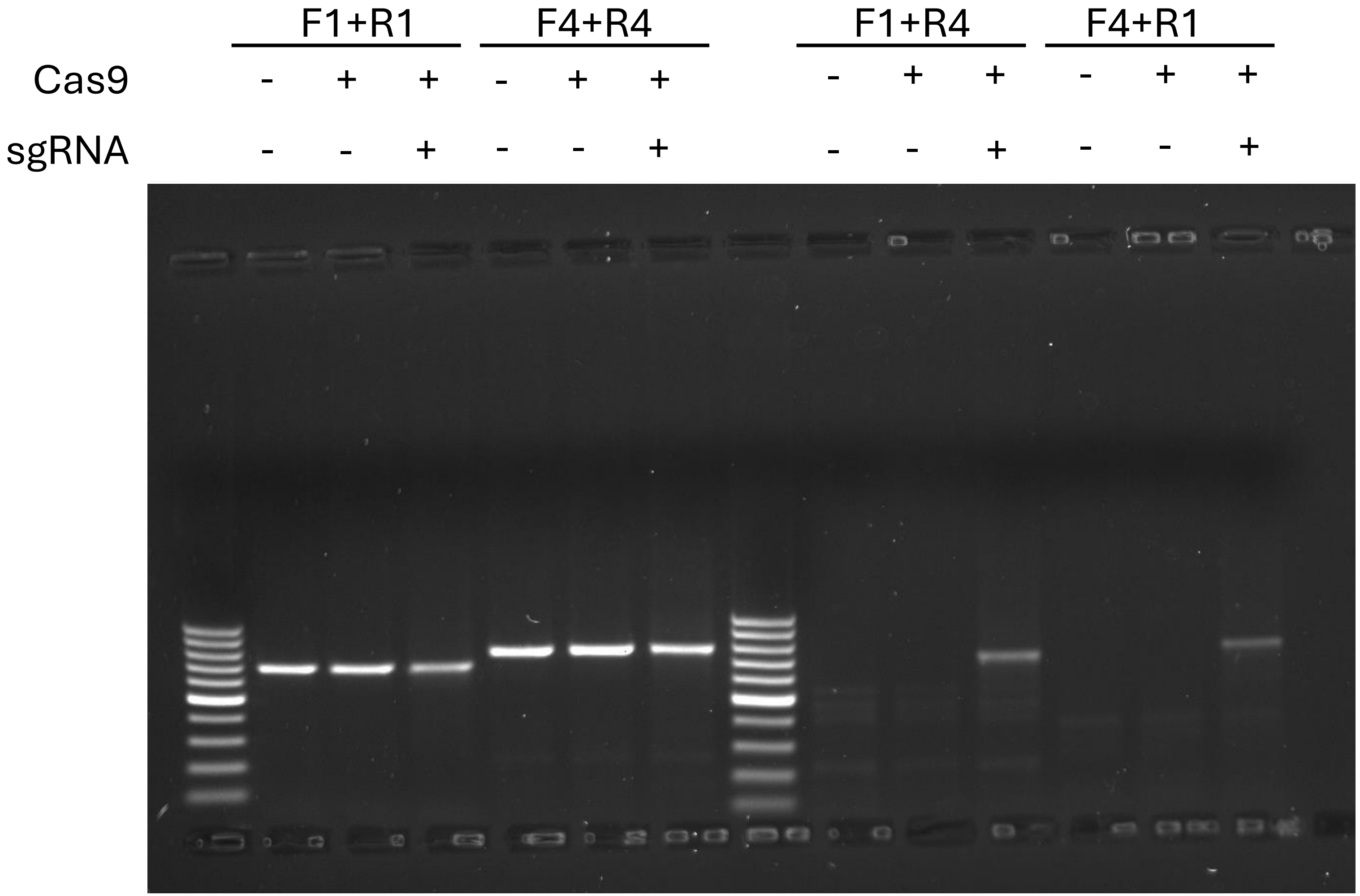

Figure 3a

### *CDK4* 3 weeks 508Kb

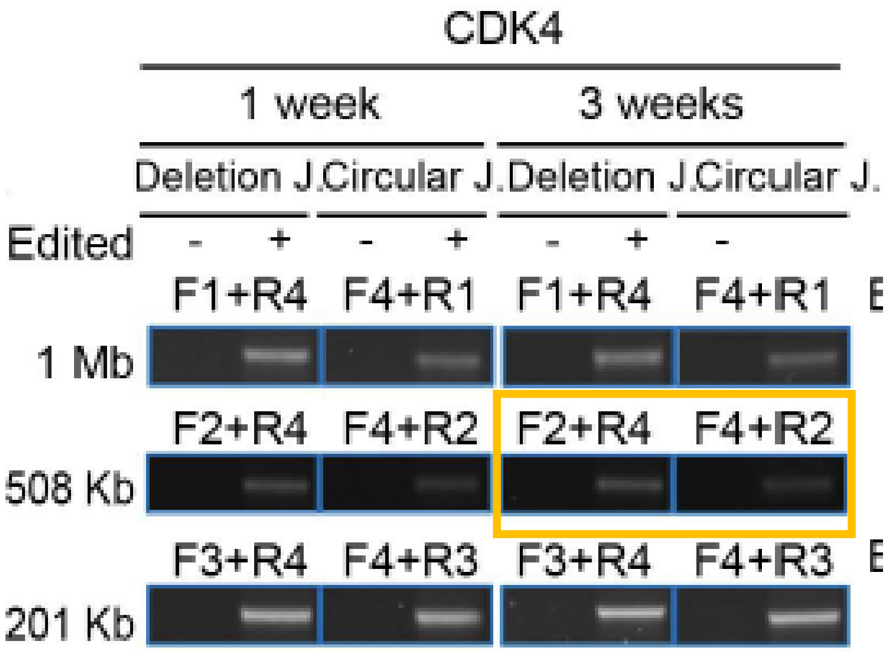


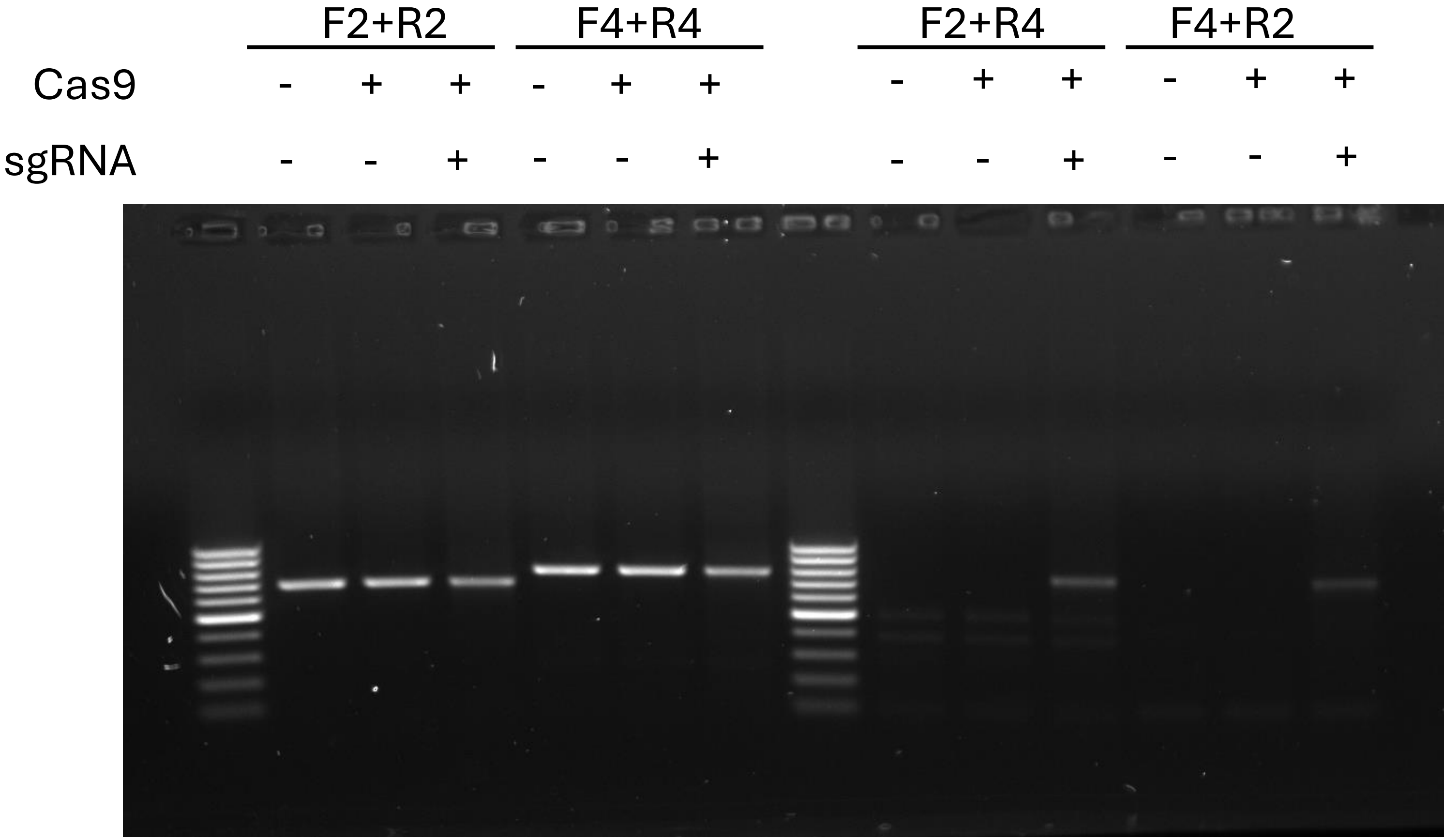

### Figure 3a

#### *CDK4* 3 weeks 201Kb

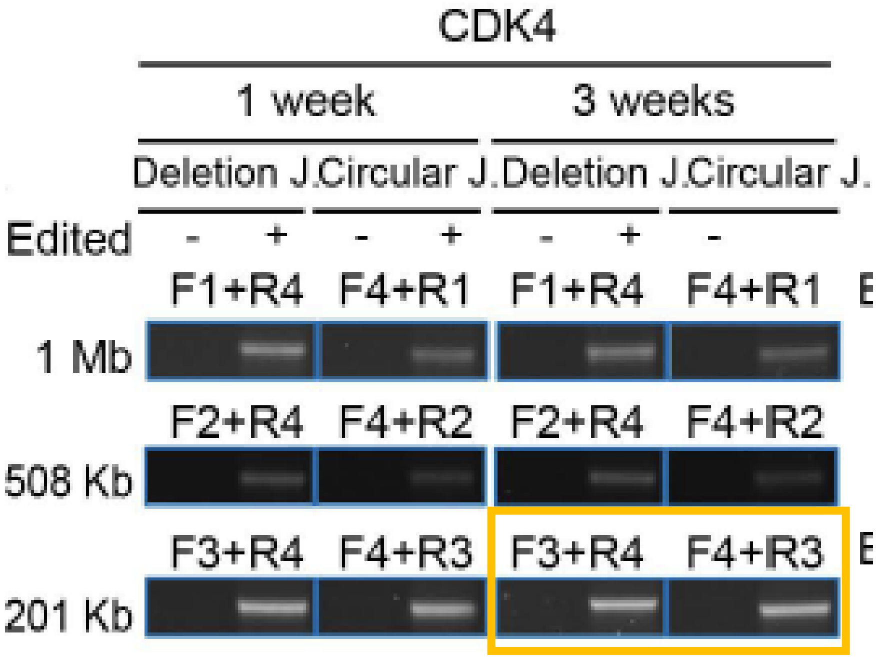


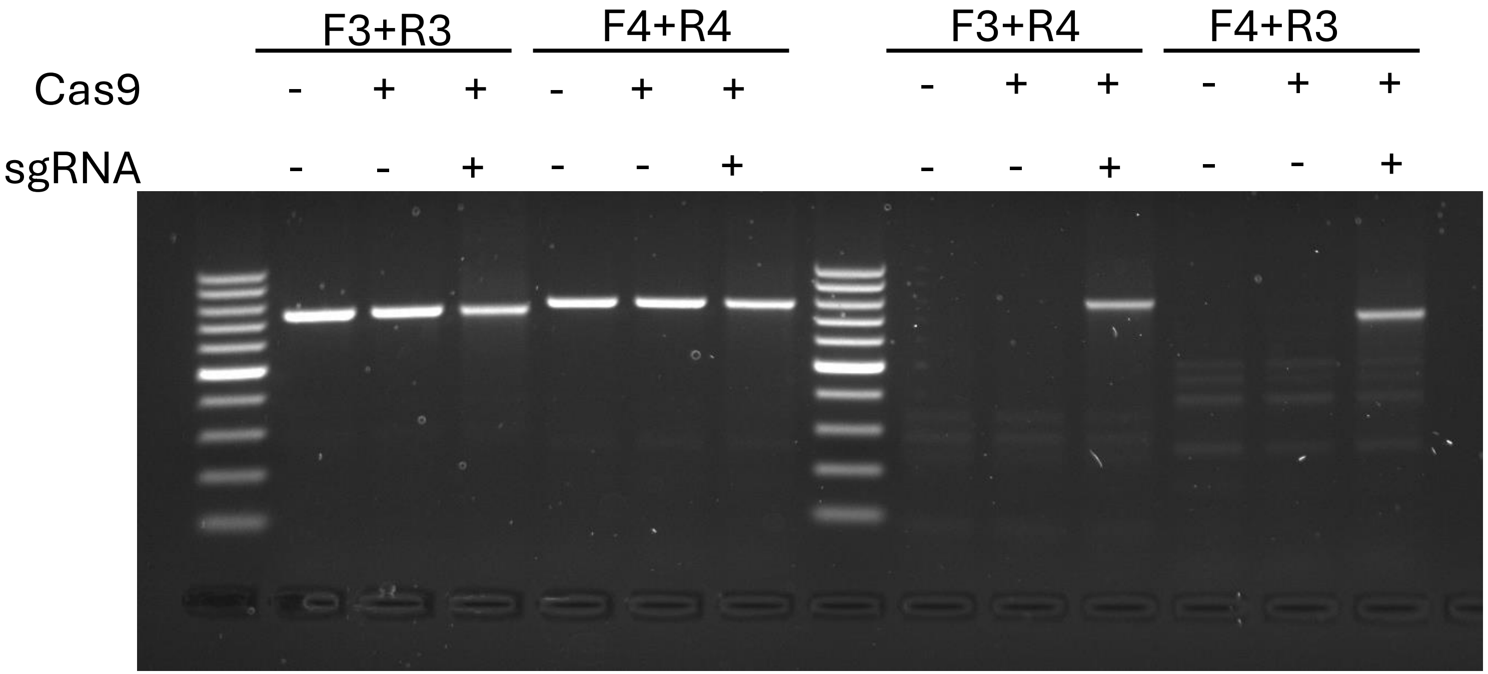

### Figure 3a

#### *MDM2*-EC: 587 Kb X *CDK4*-EC: 508 Kb – 1 week

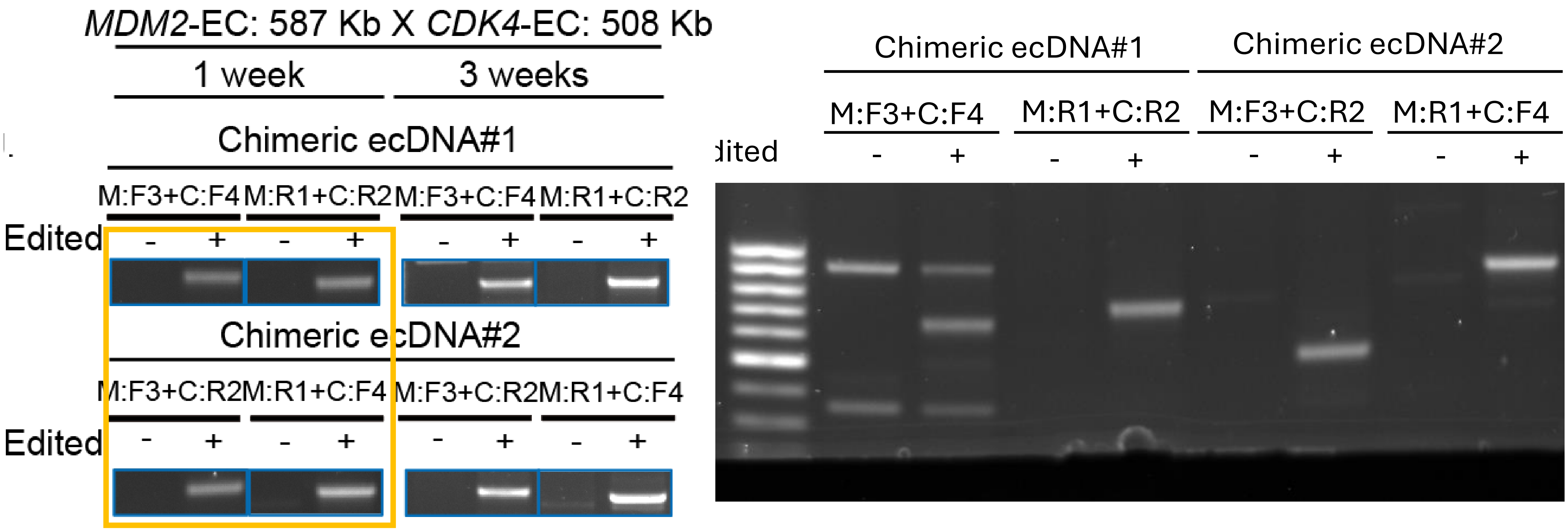

### Figure 3a

#### *MDM2*-EC: 587 Kb X *CDK4*-EC: 508 Kb – 3 weeks

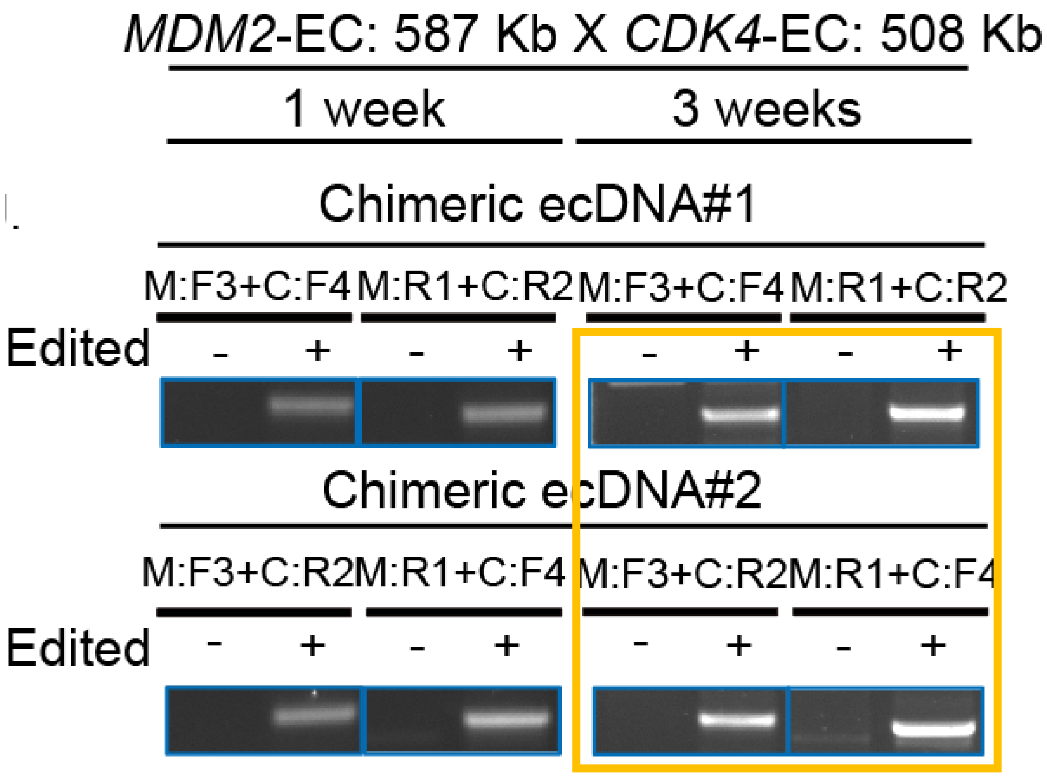


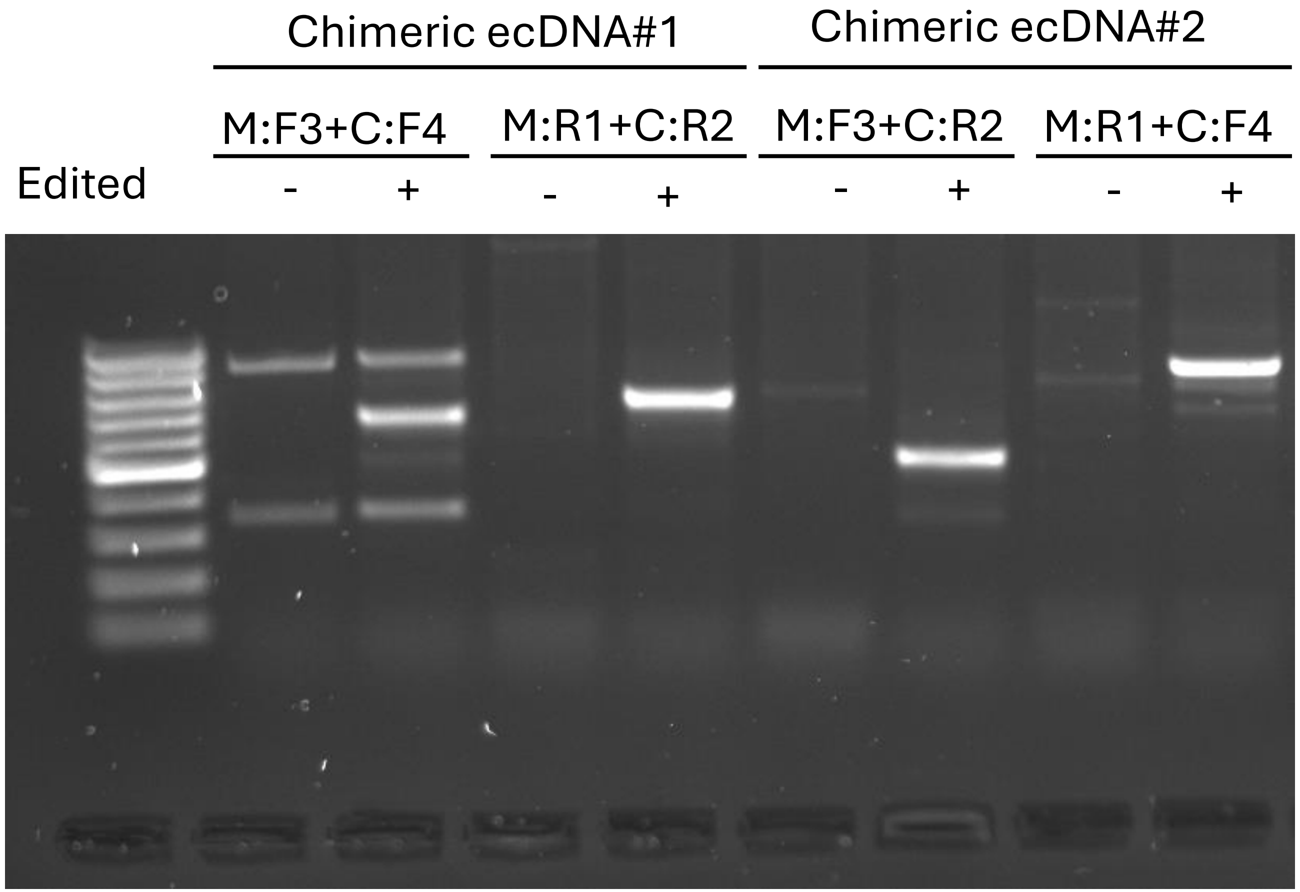

### Figure 5b

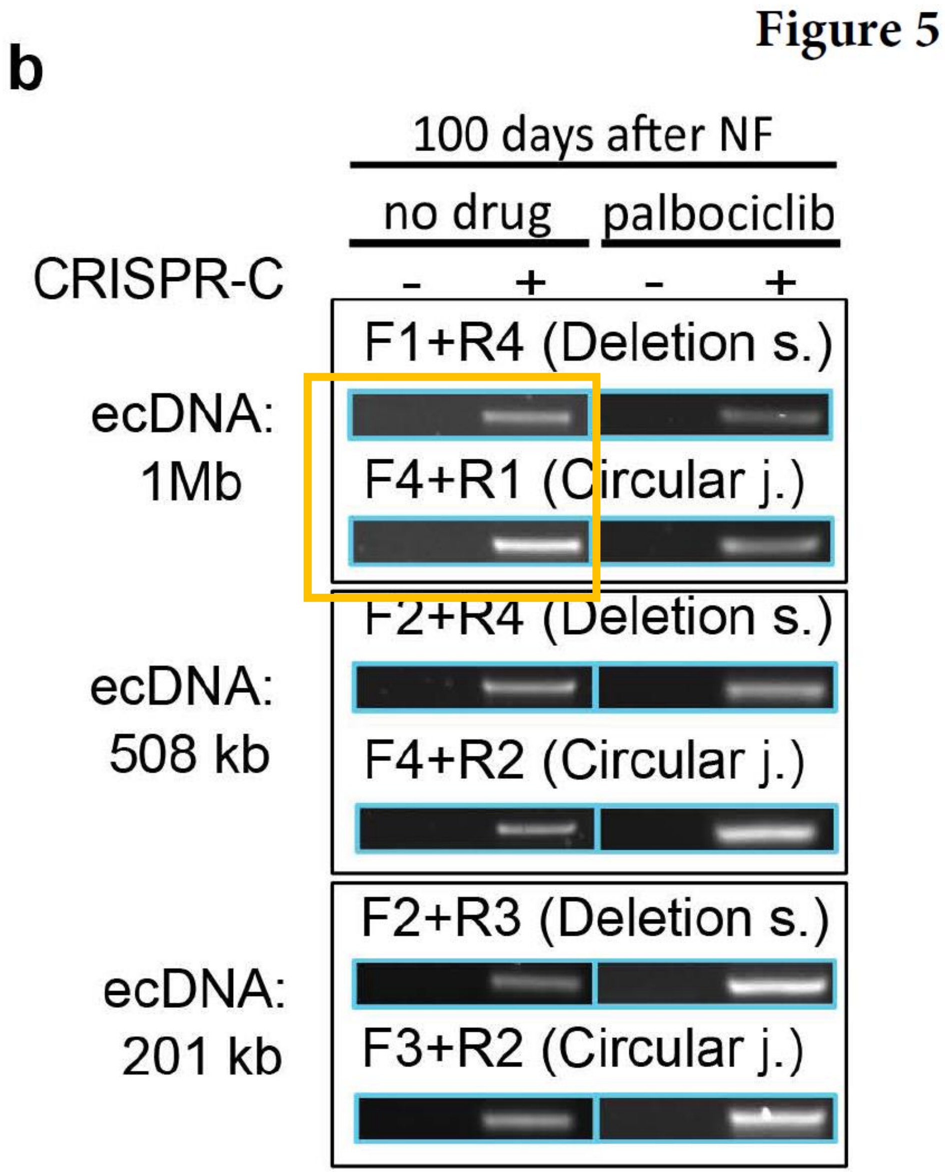


#### No drug CDK4 1Mb

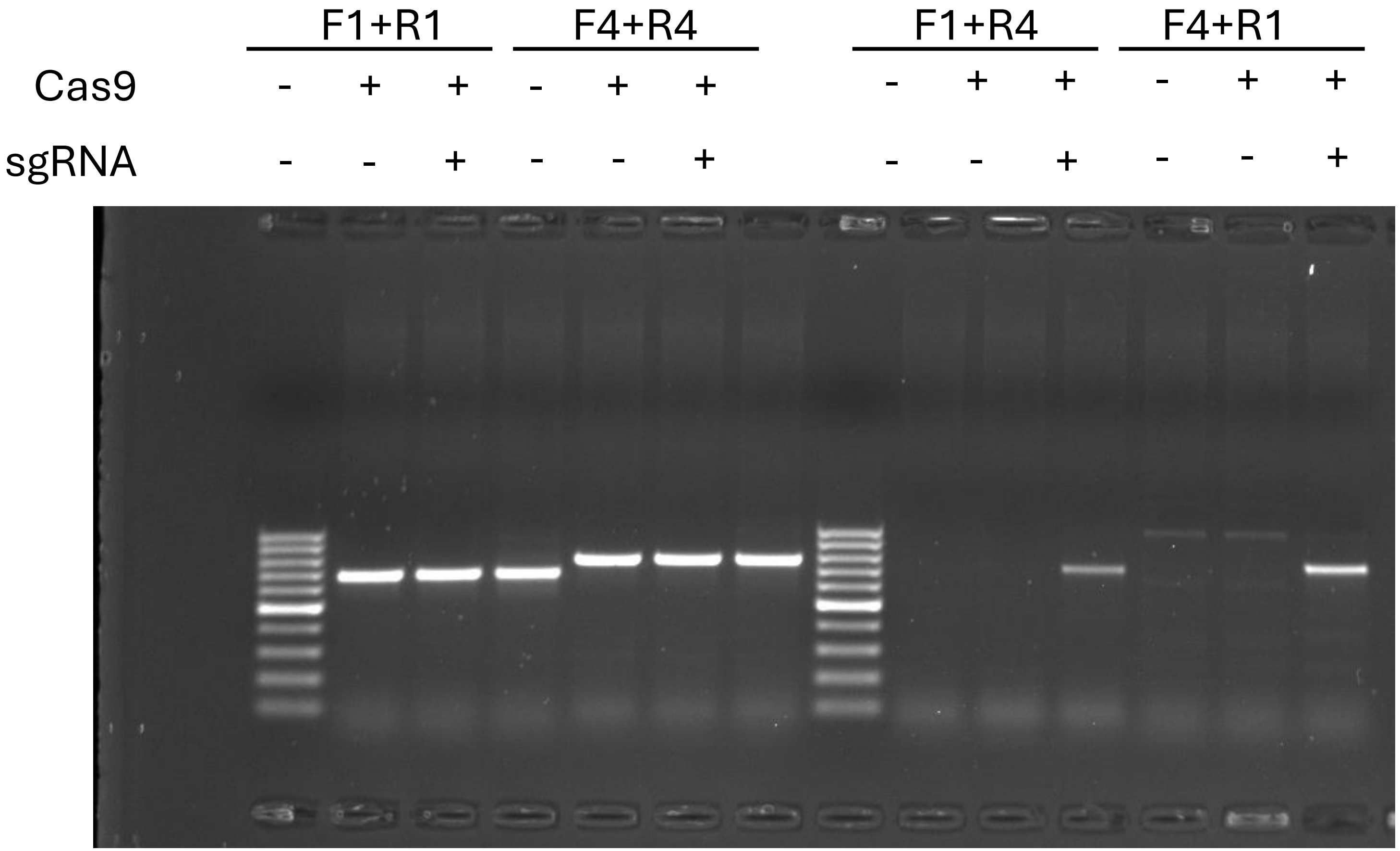

### Figure 5b

### No drug CDK4 508 Kb

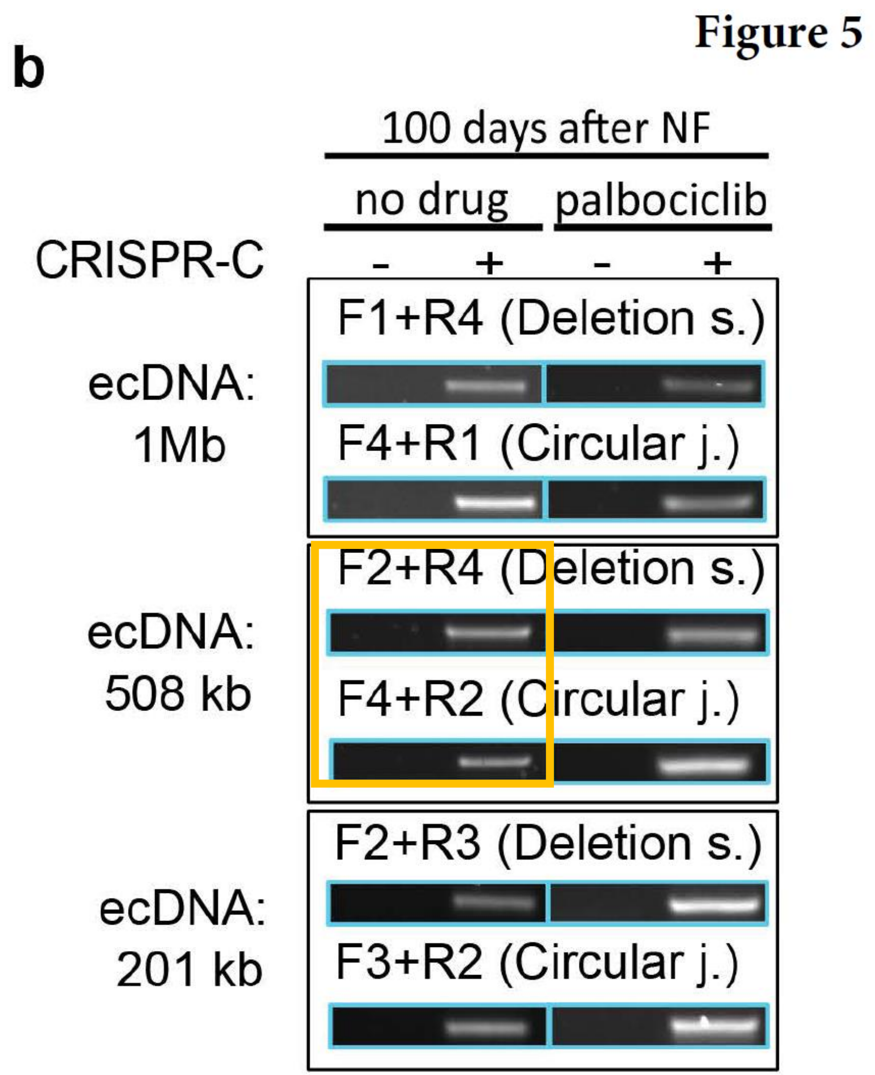


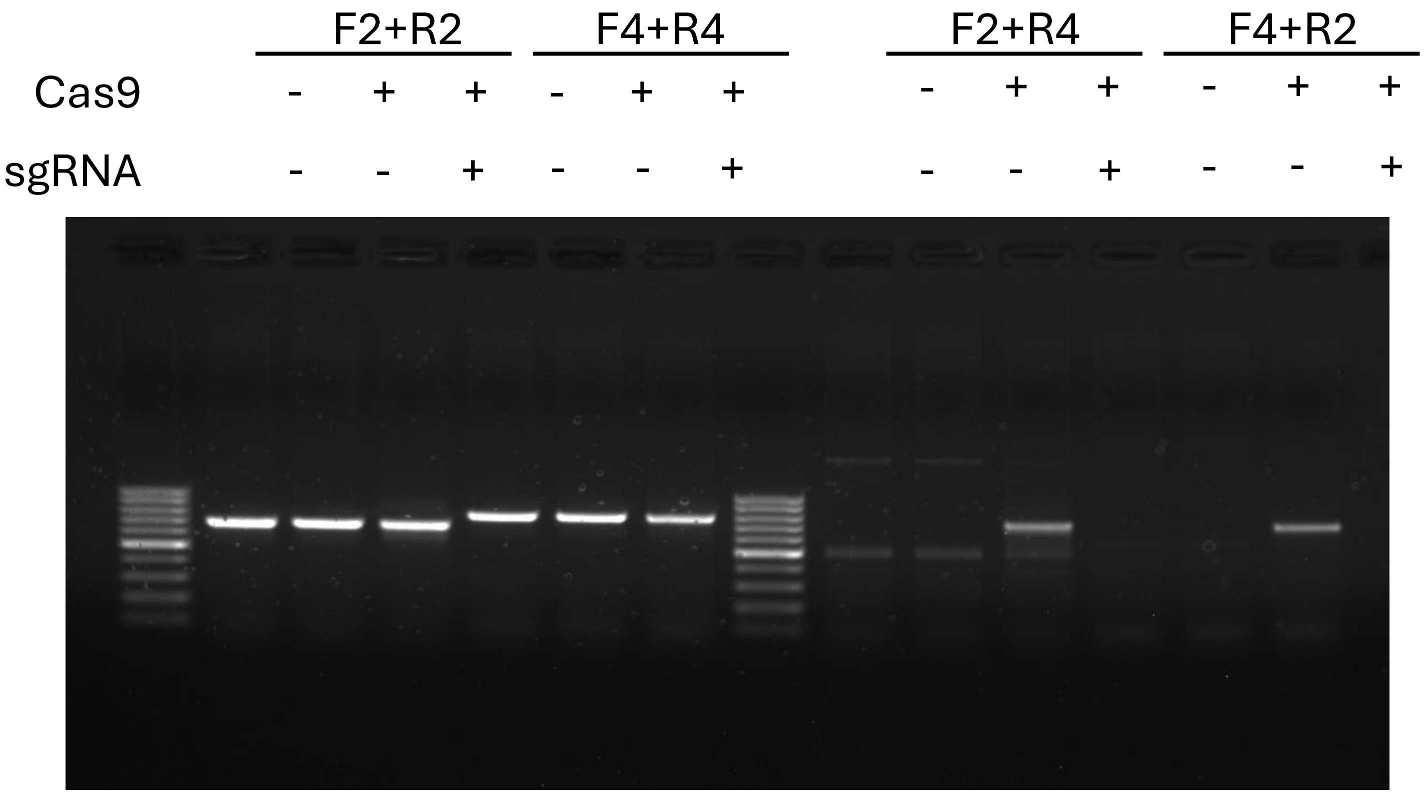

### Figure 5b

#### No drug CDK4 201 Kb

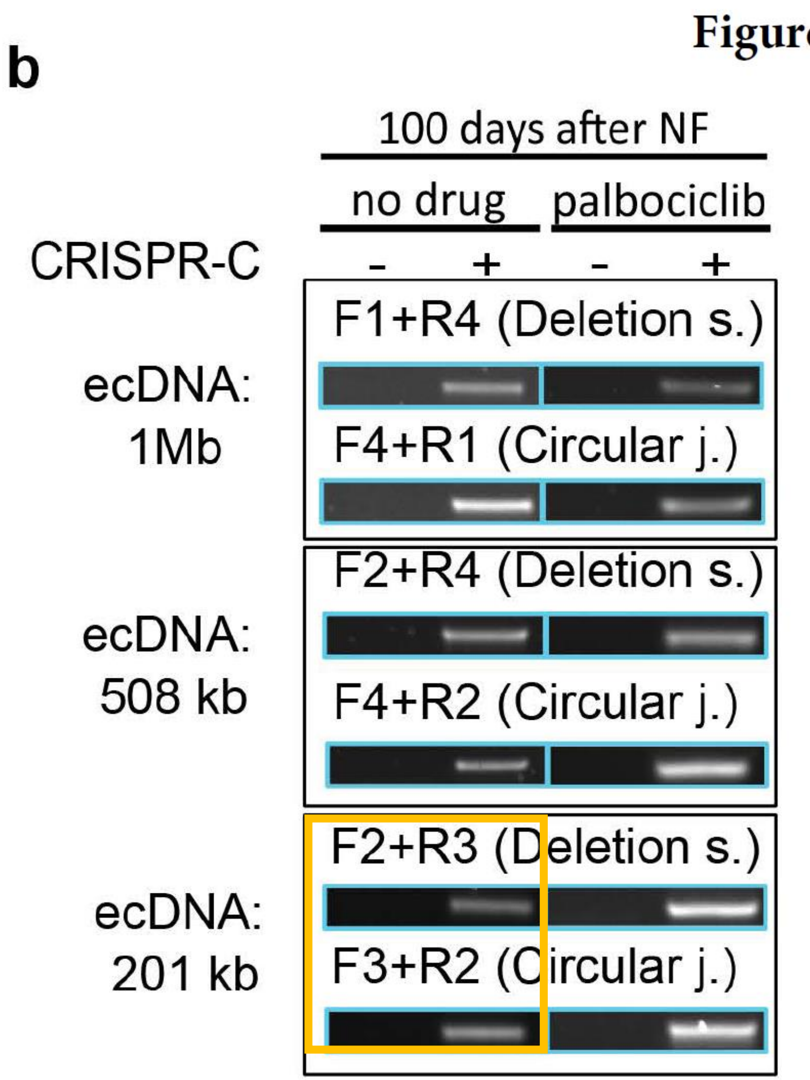


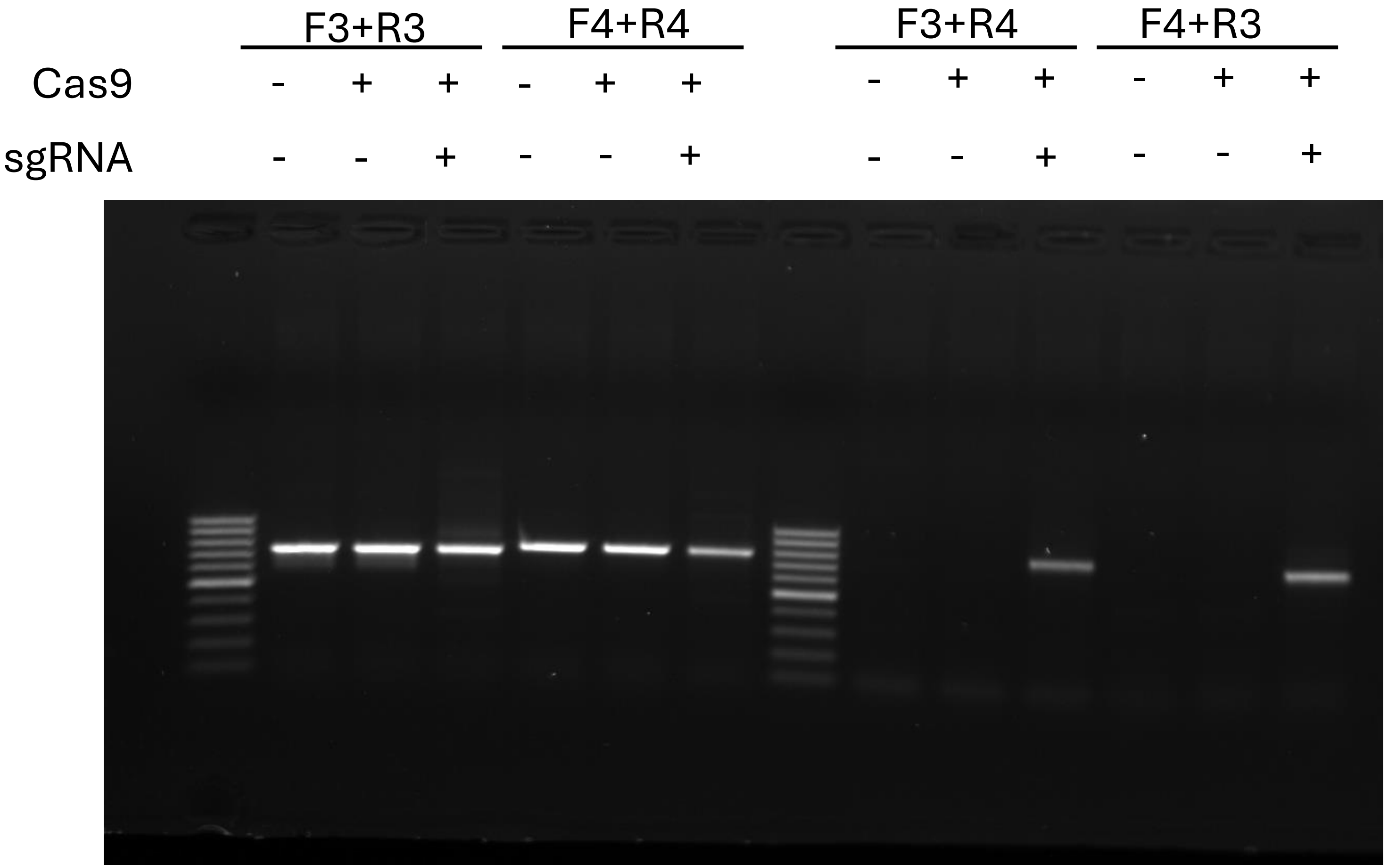

### Figure 5b

### Palbociclib CDK4 1Mb

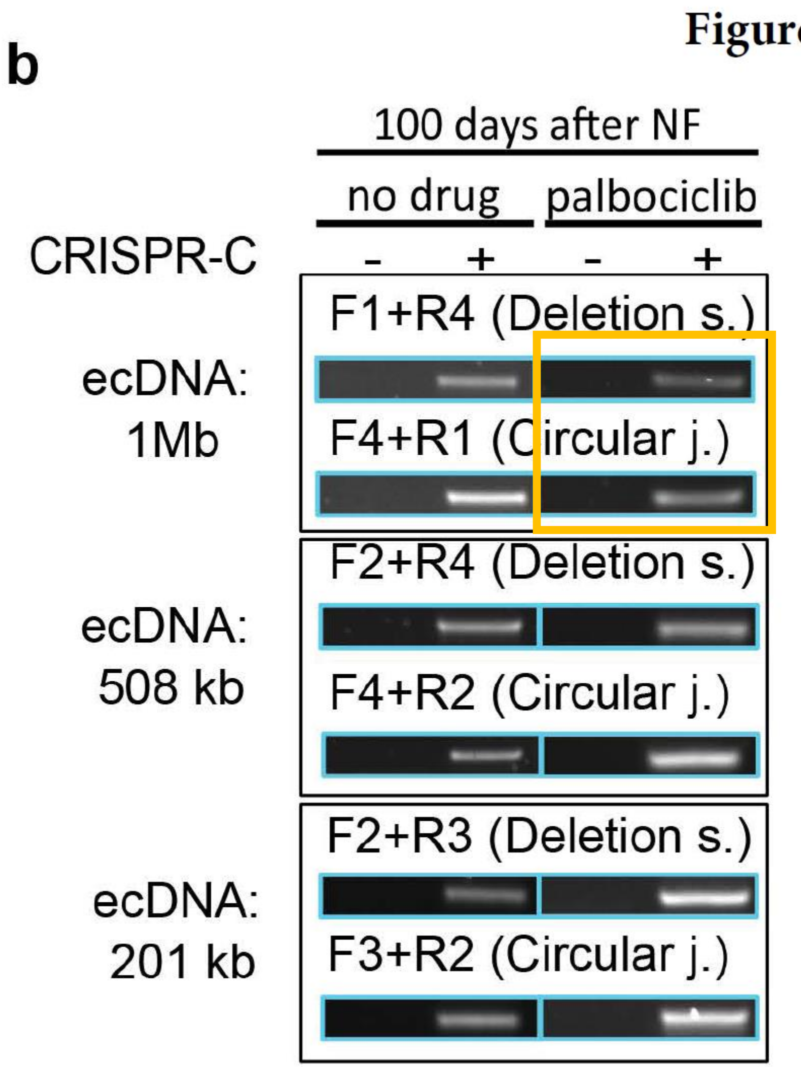


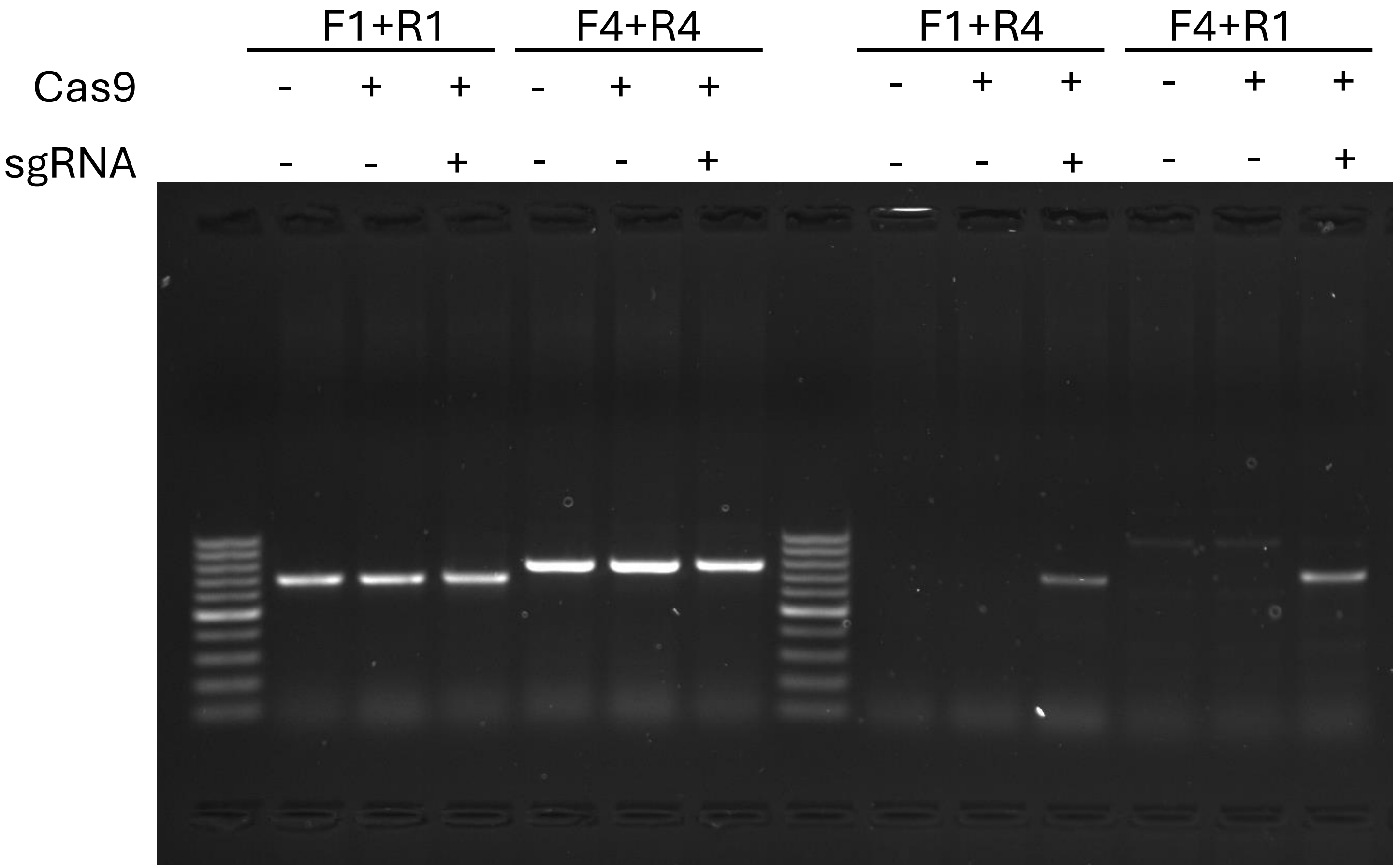

### Figure 5b

#### Palbociclib CDK4 508 Kb

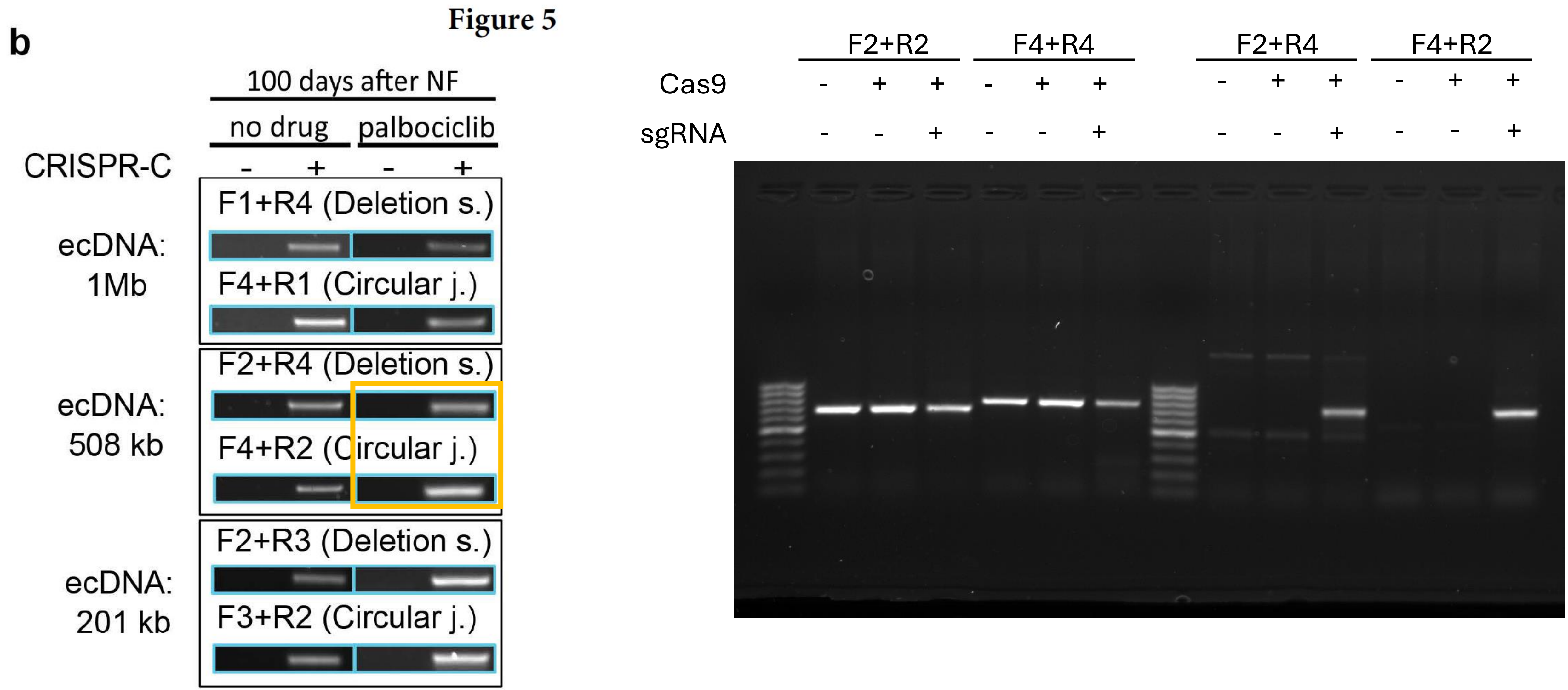

### Figure 5b

### Palbociclib CDK4 201 Kb

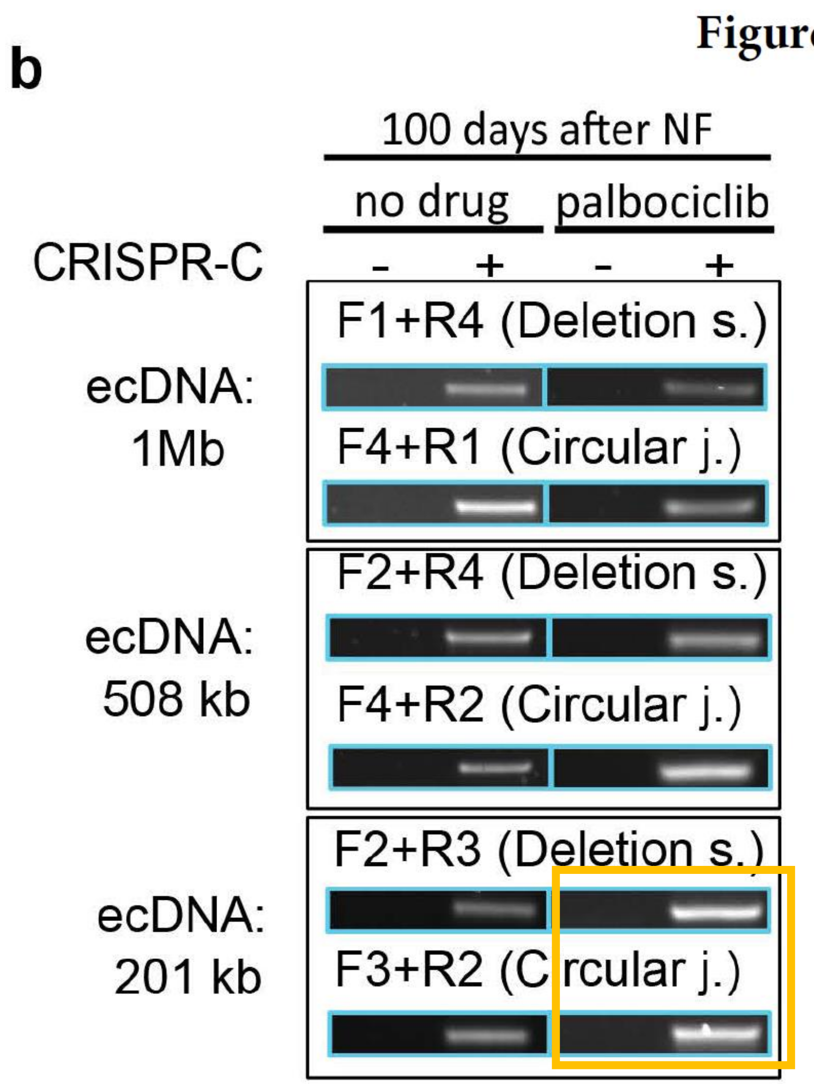


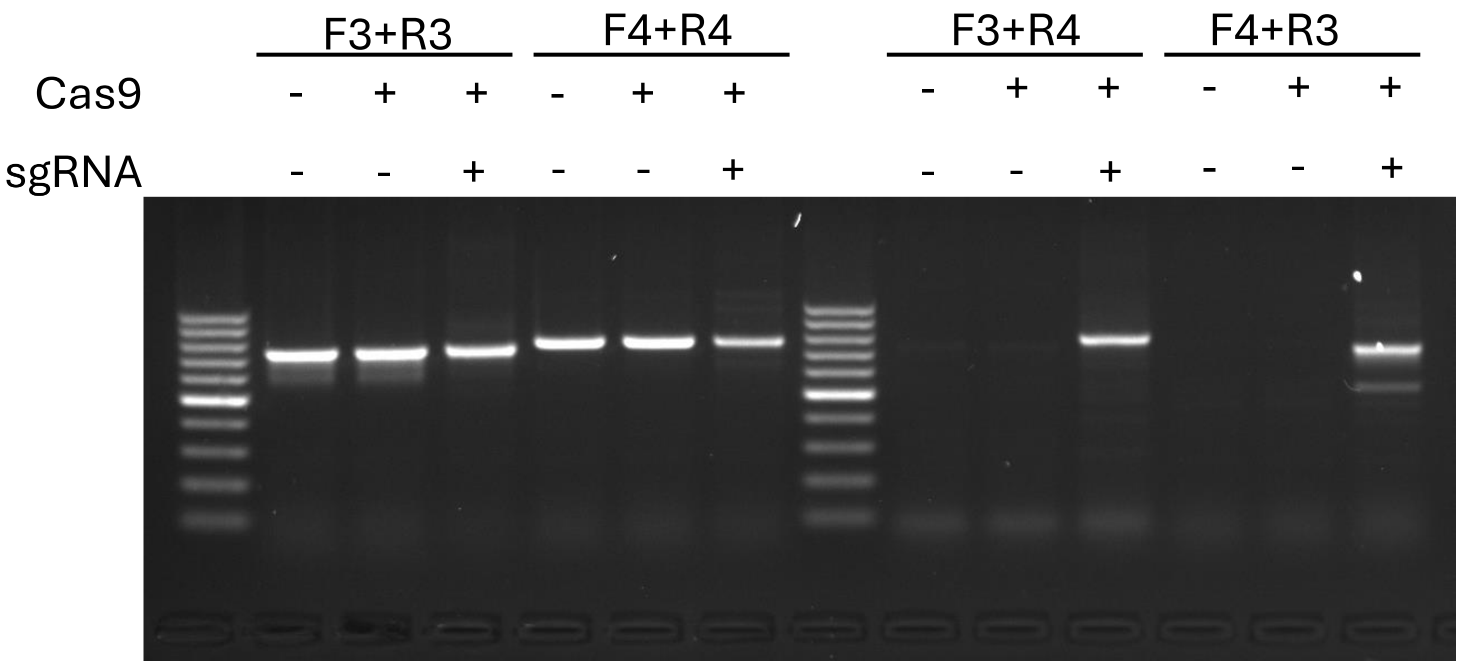

### Figure 6b

#### No colcemid 2 weeks

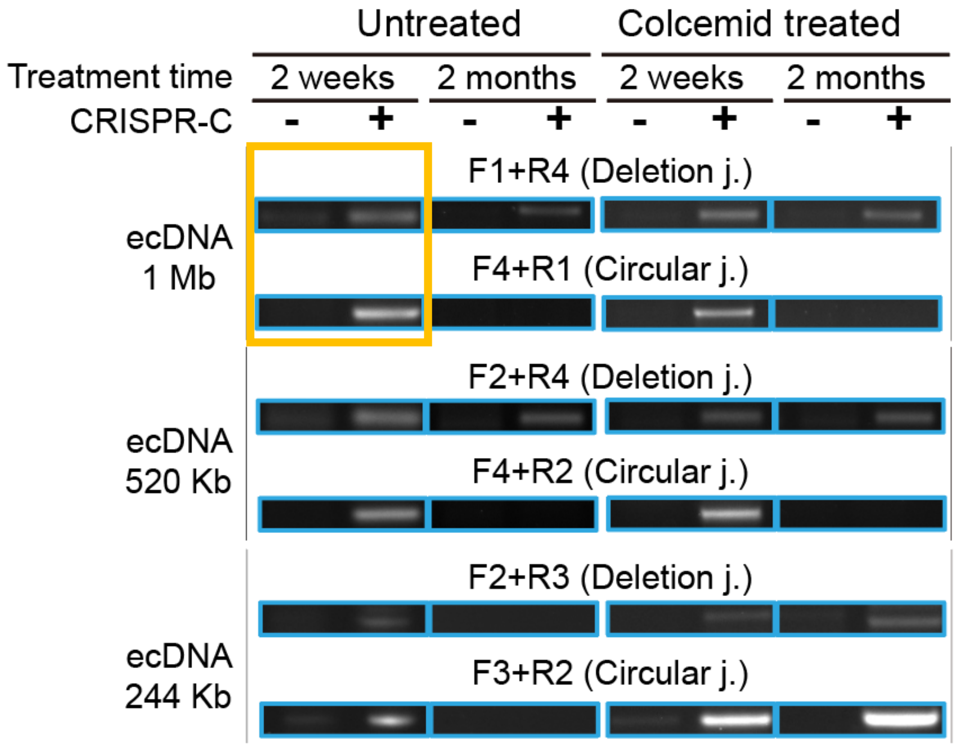


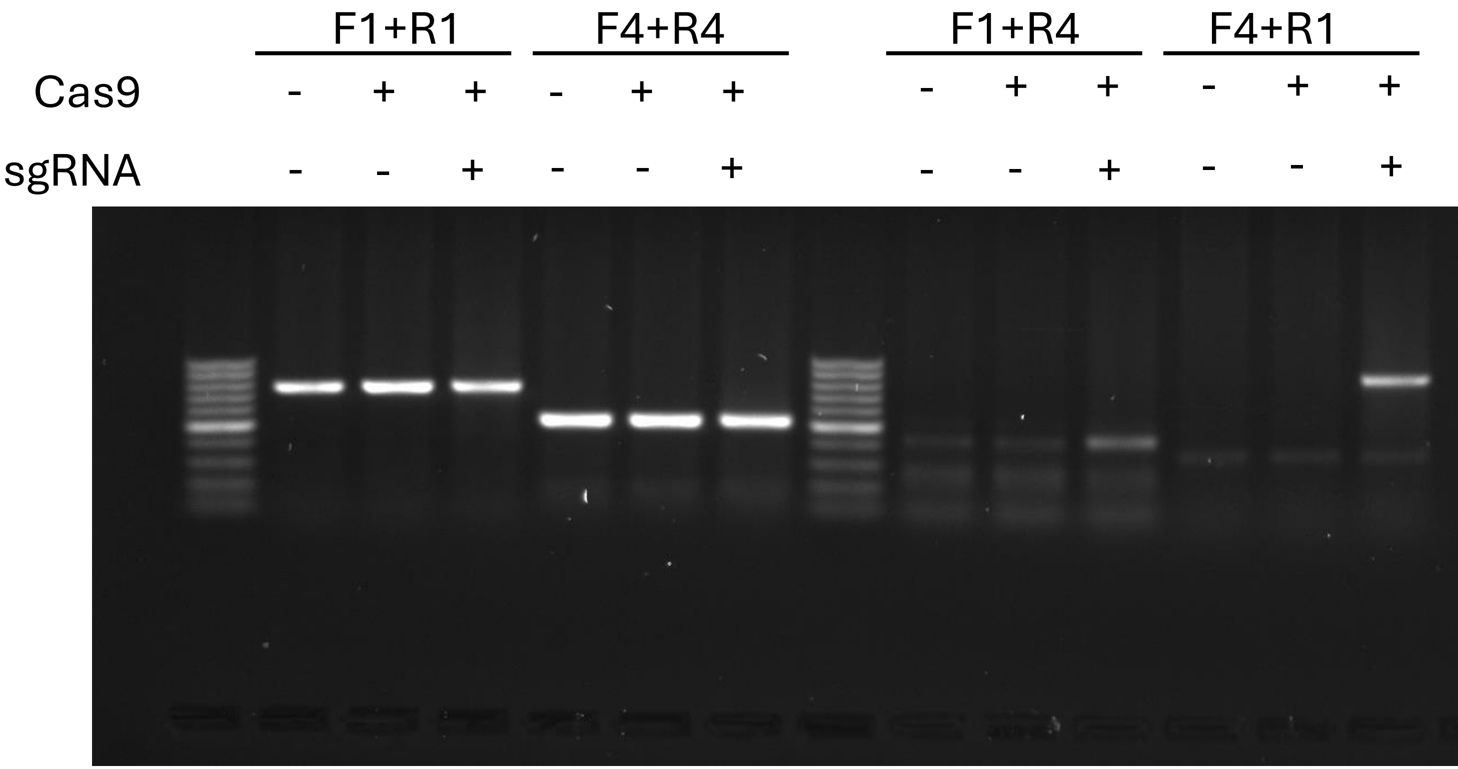

### Figure 6b

#### No colcemid 2 weeks

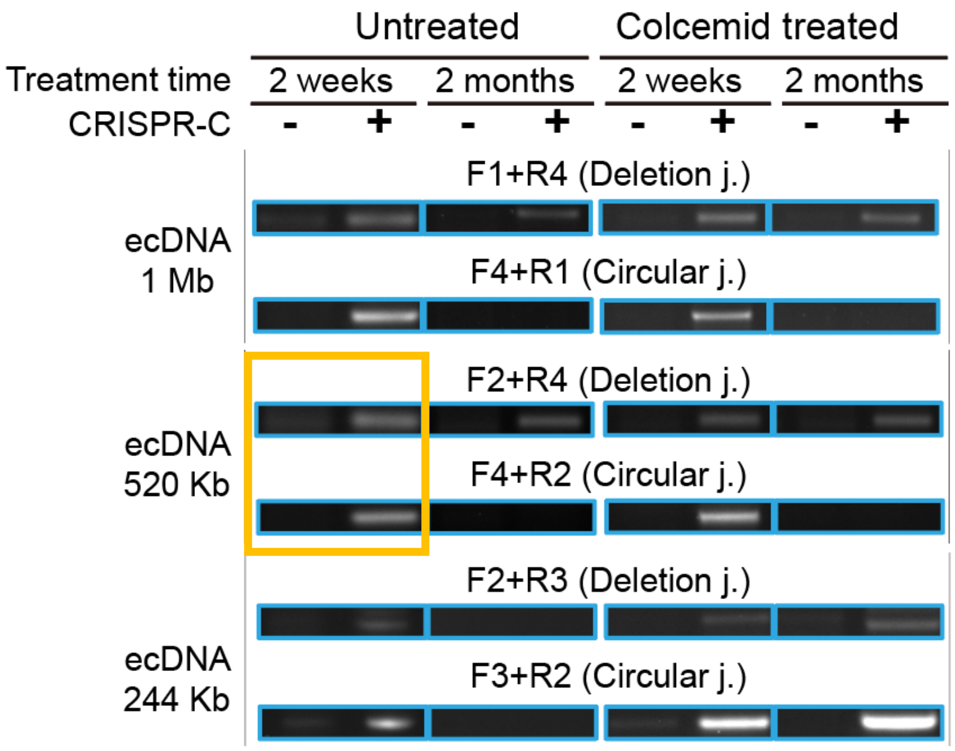


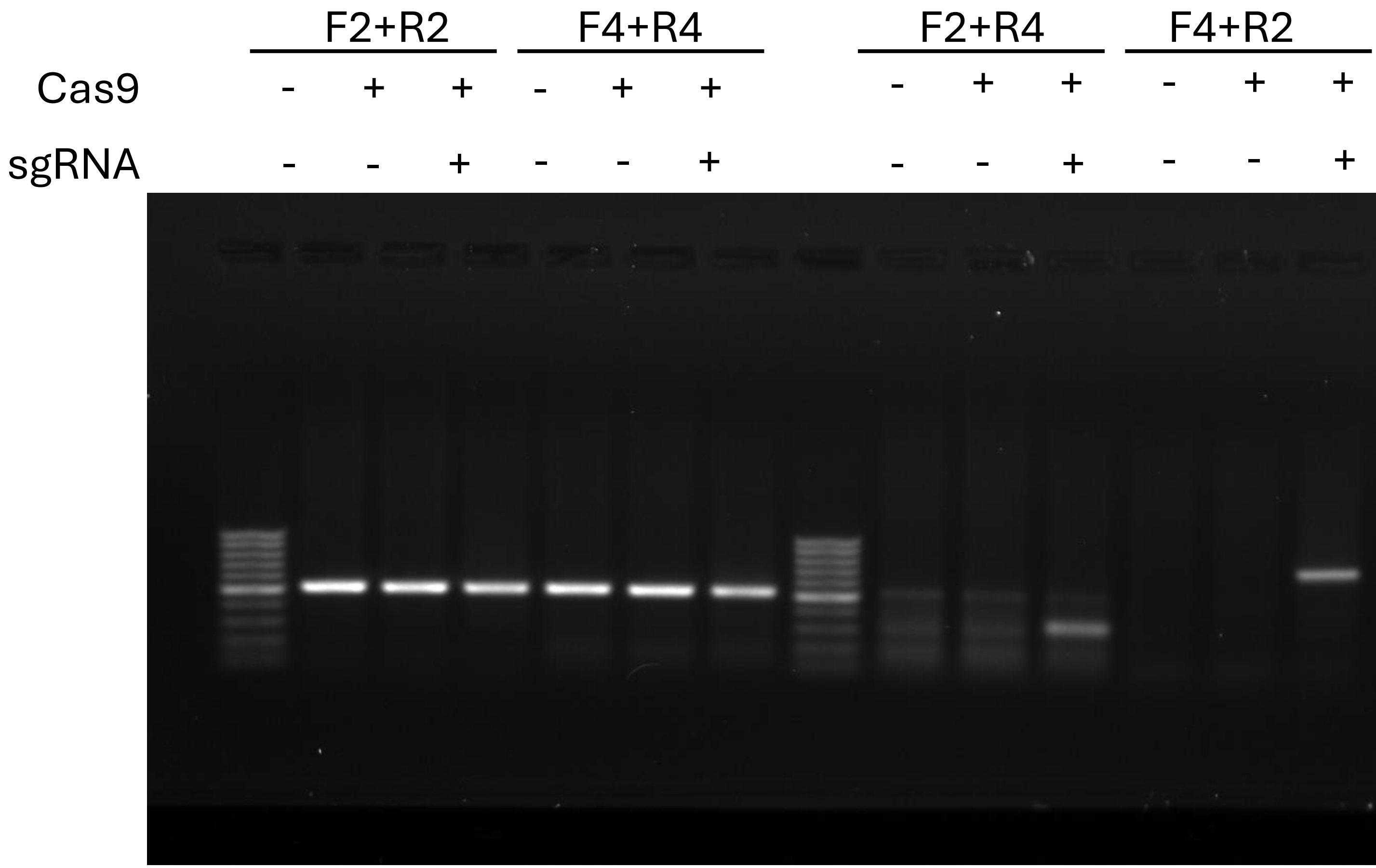

### Figure 6b

#### No colcemid 2 weeks

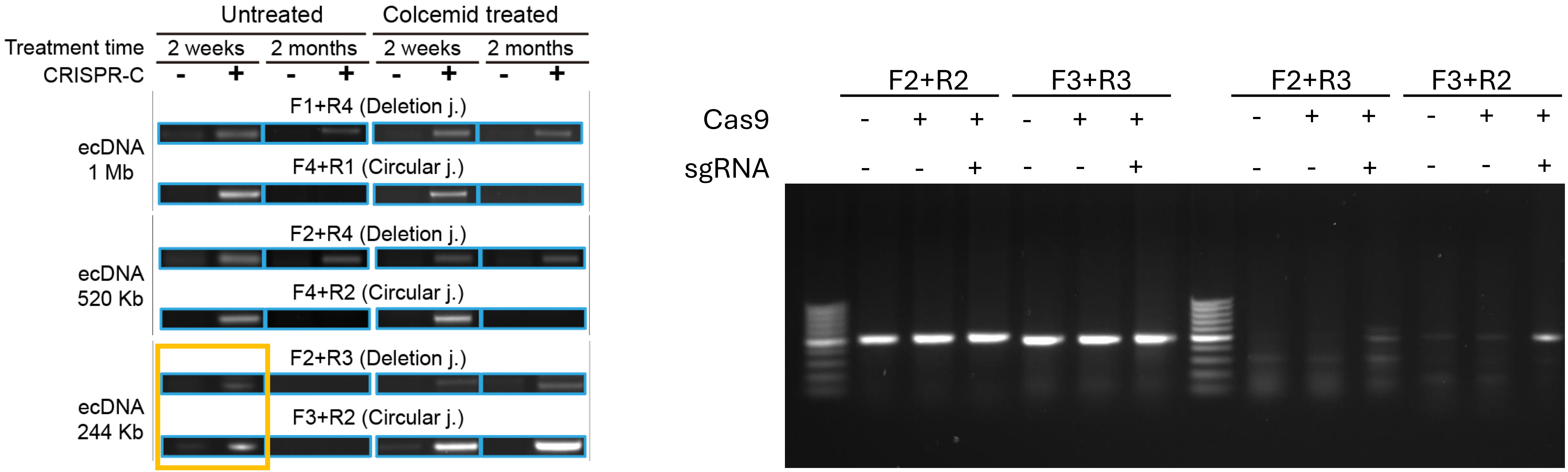

### Figure 6b

#### No colcemid 2 months

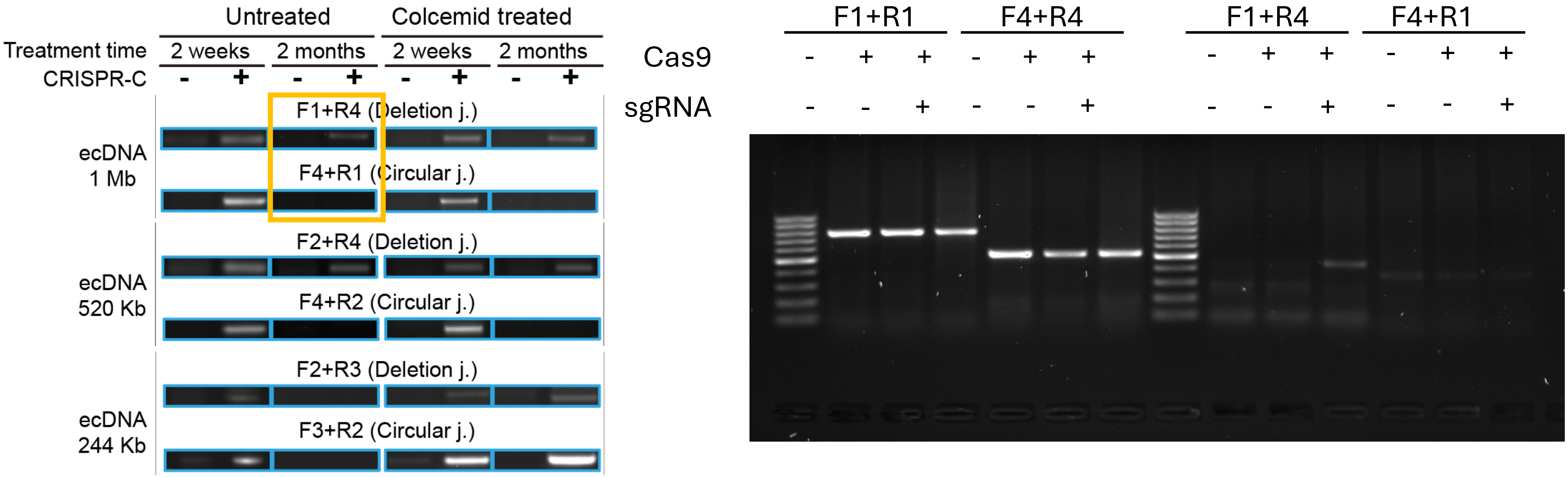

### Figure 6b

#### No colcemid 2 months

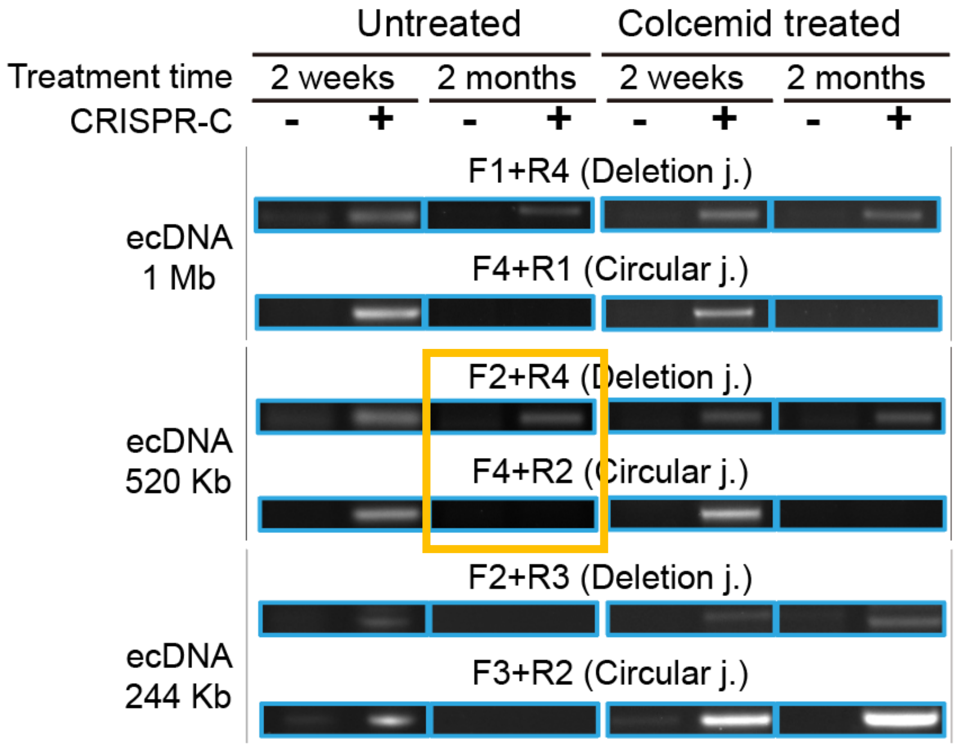


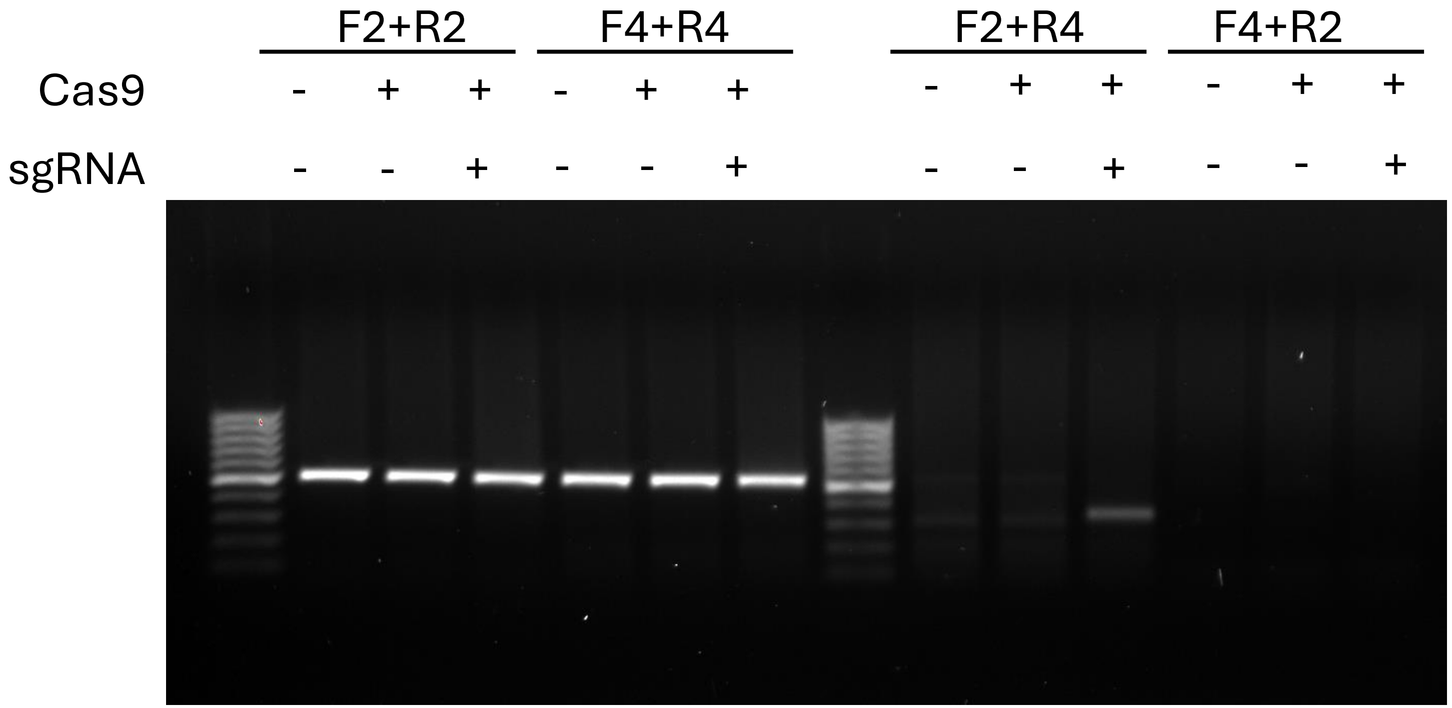

### Figure 6b

#### No colcemid 2 months

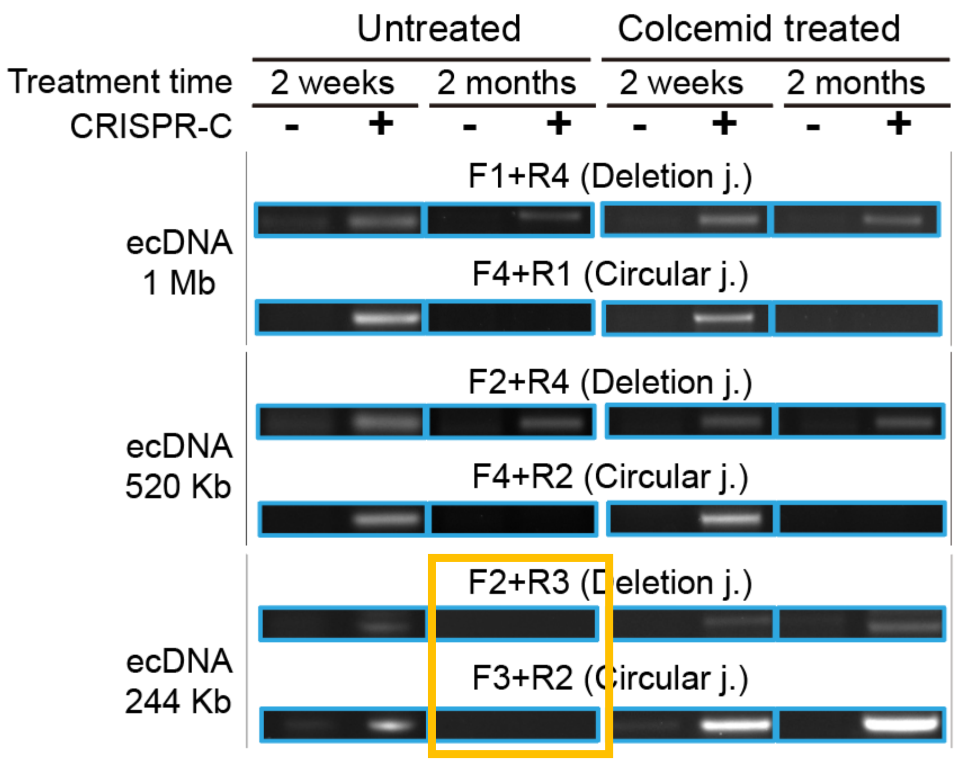


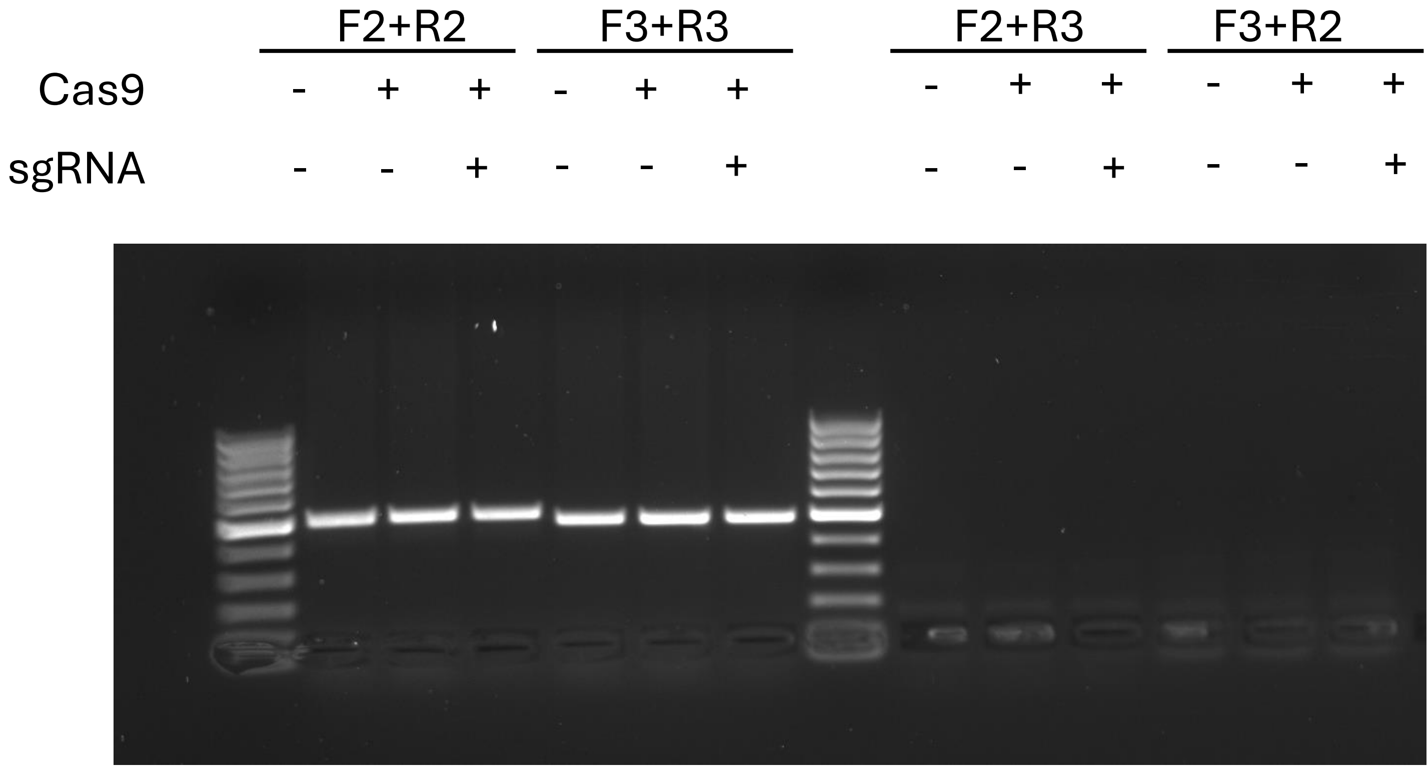

### Figure 6b

#### Colcemid 2 weeks

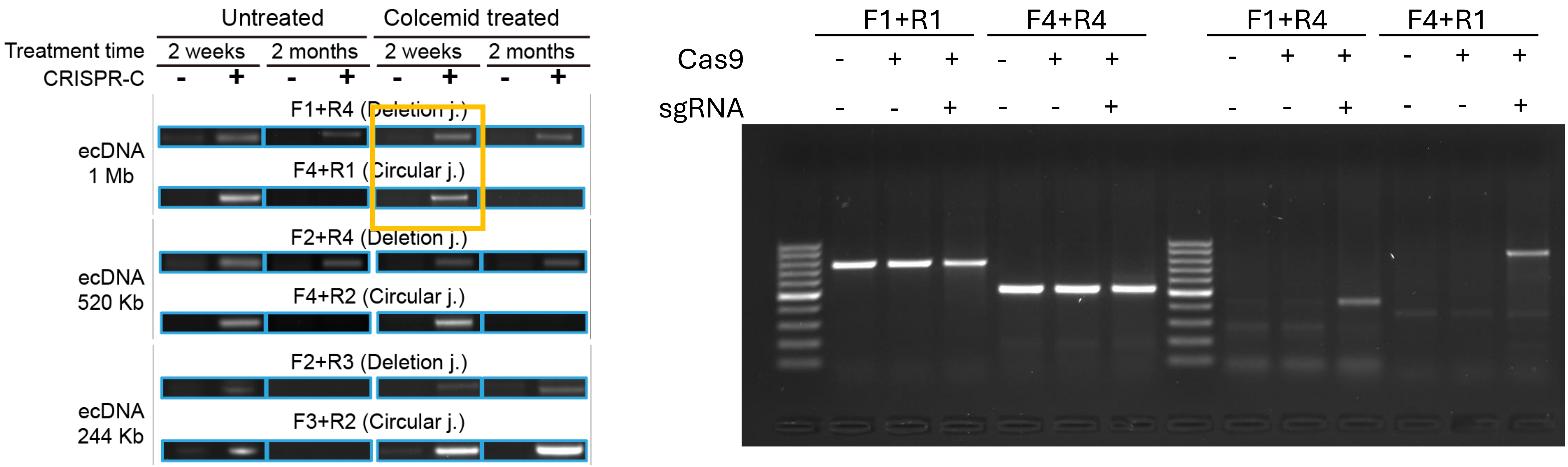

### Figure 6b

#### Colcemid 2 weeks

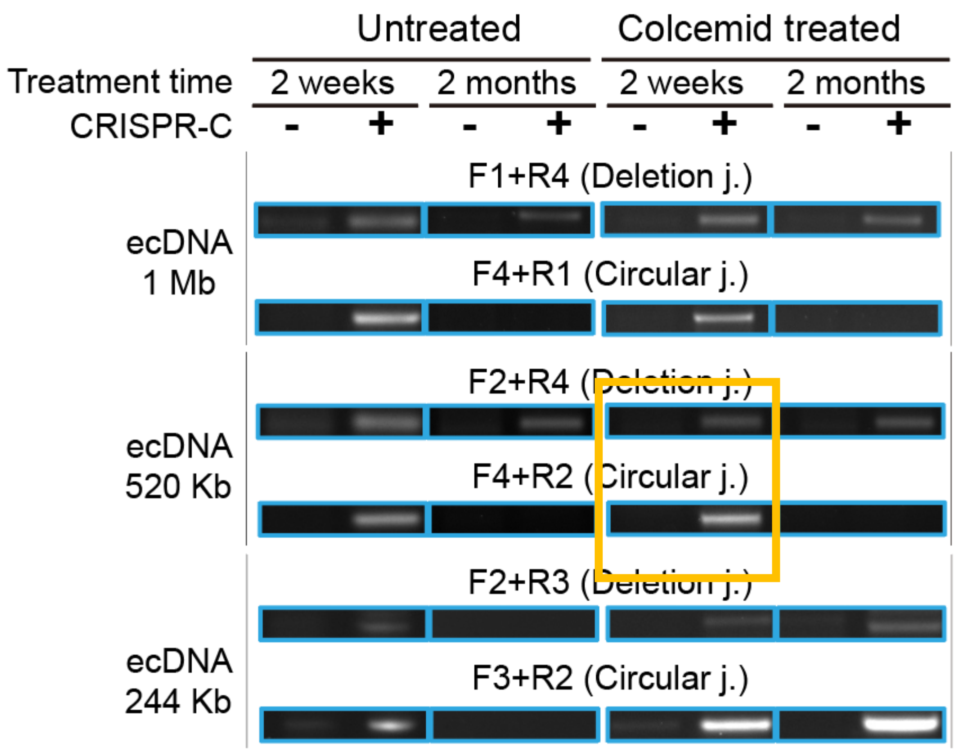


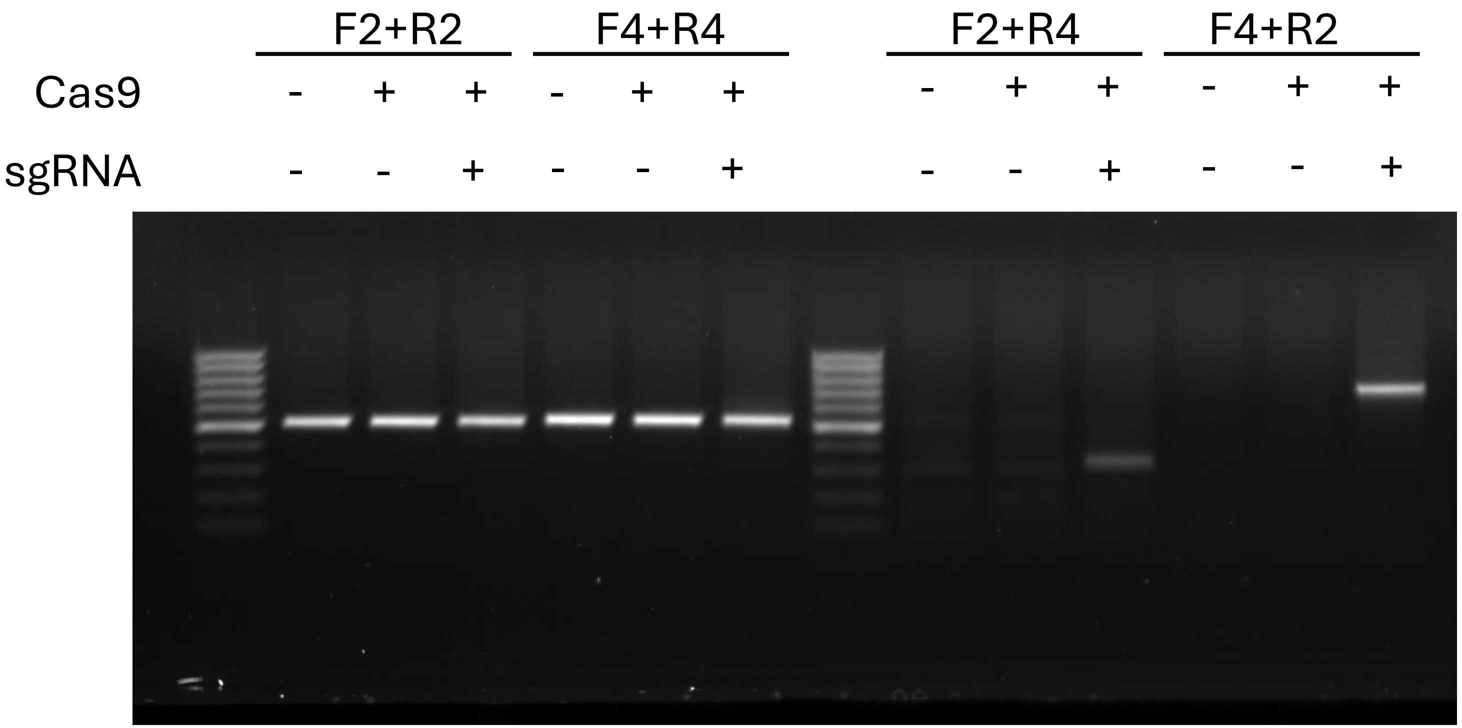

### Figure 6b

#### Colcemid 2 weeks

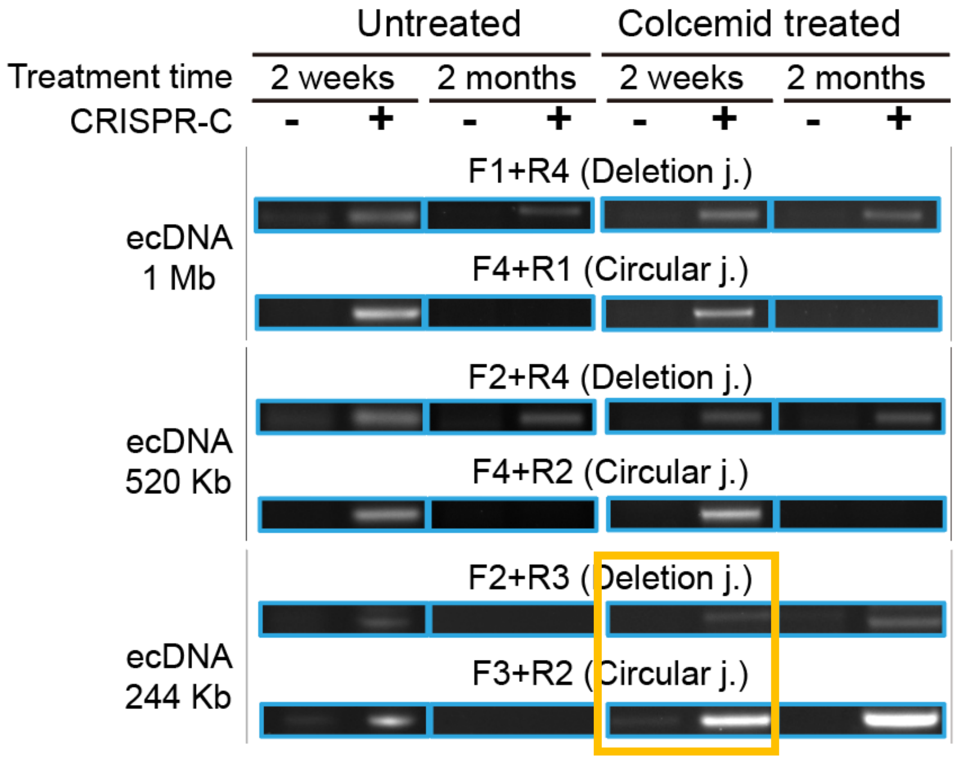


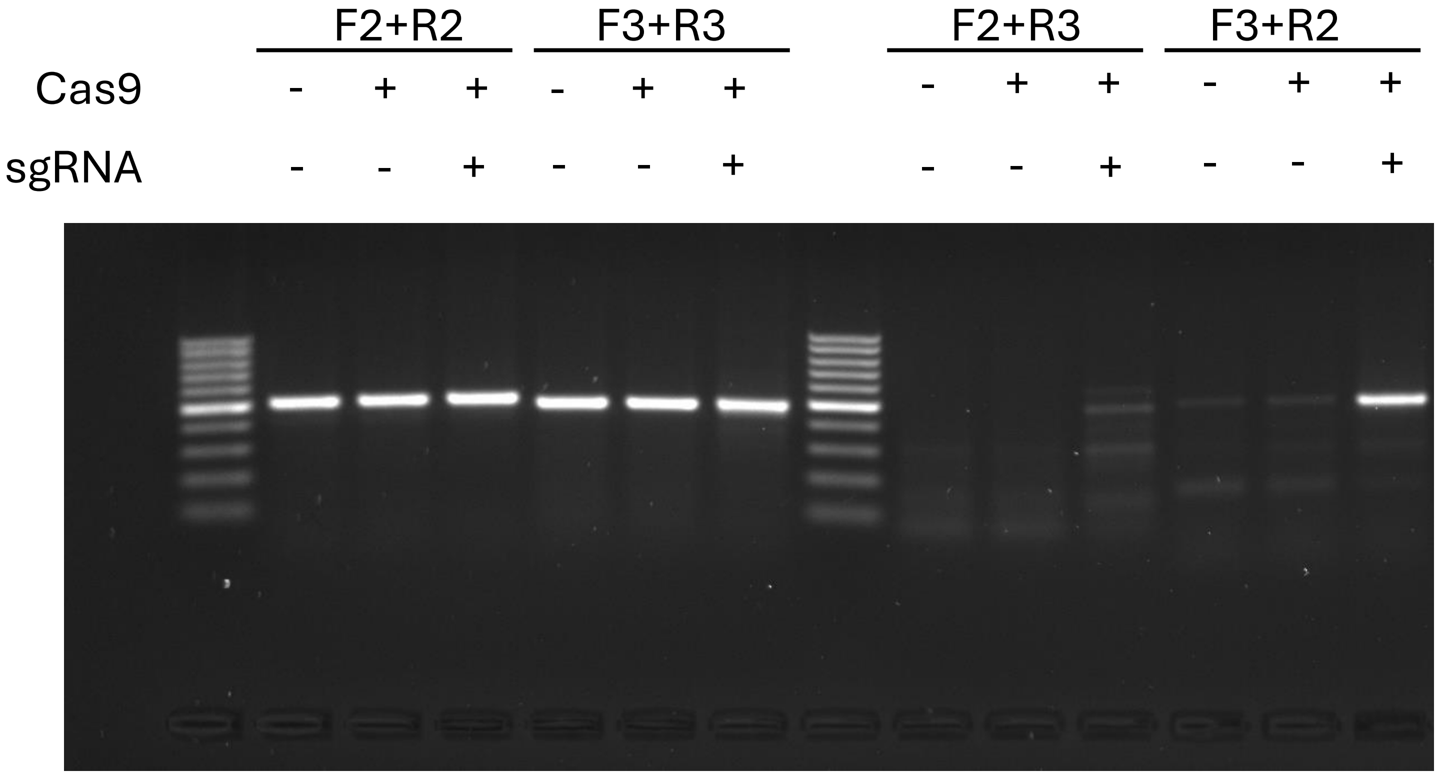

### Figure 6b

#### Colcemid 2 months

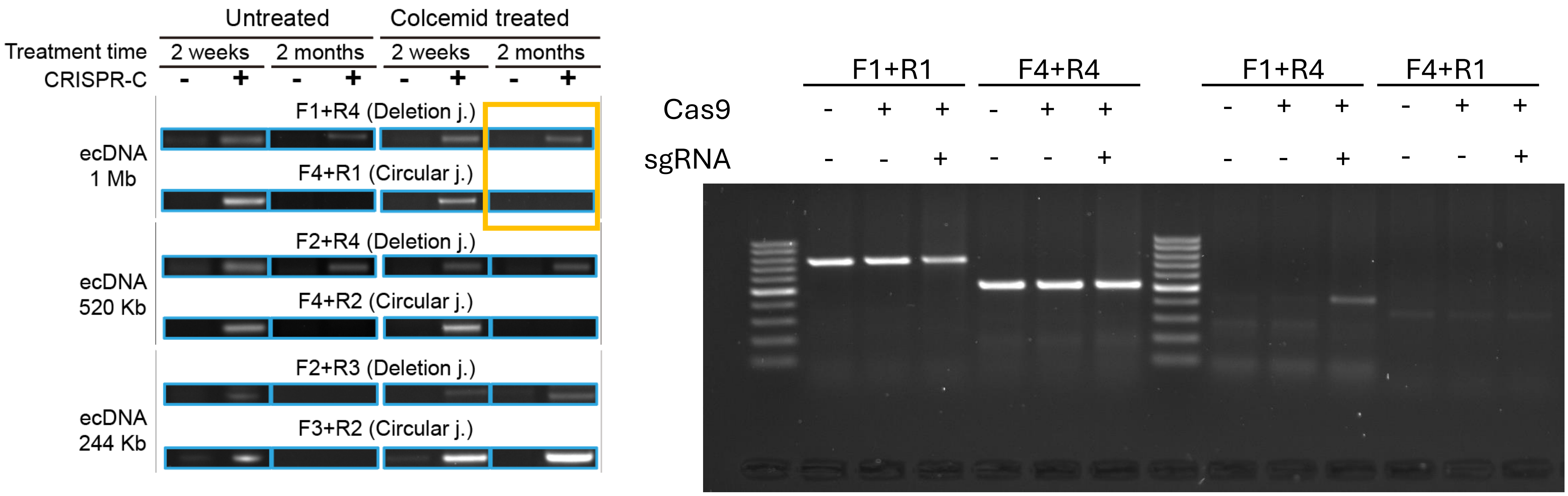

### Figure 6b

#### Colcemid 2 months

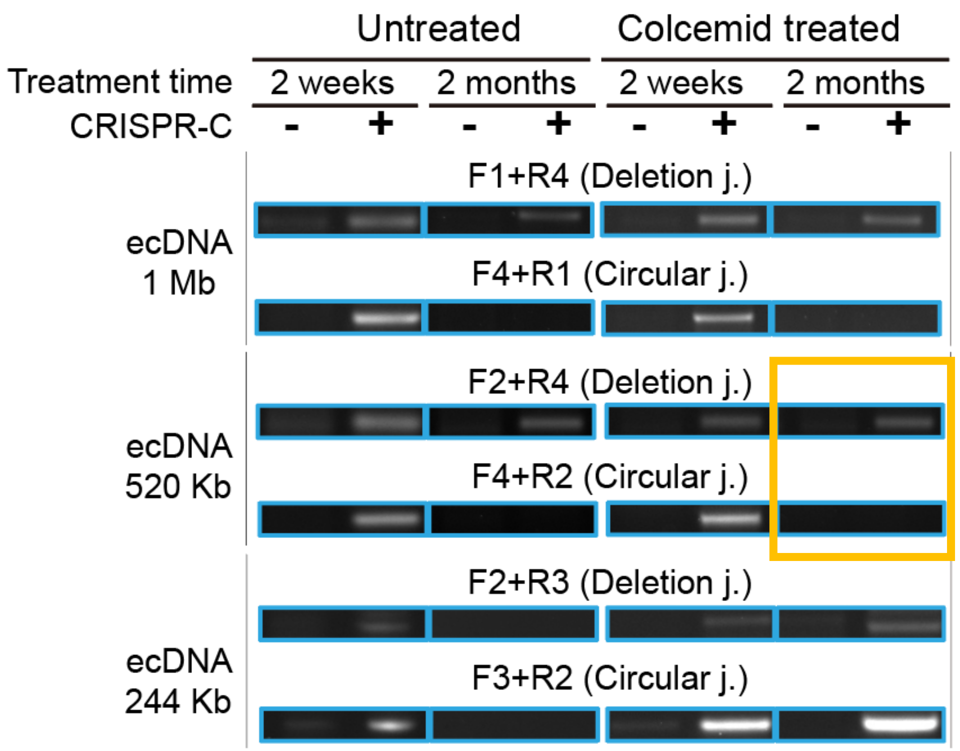


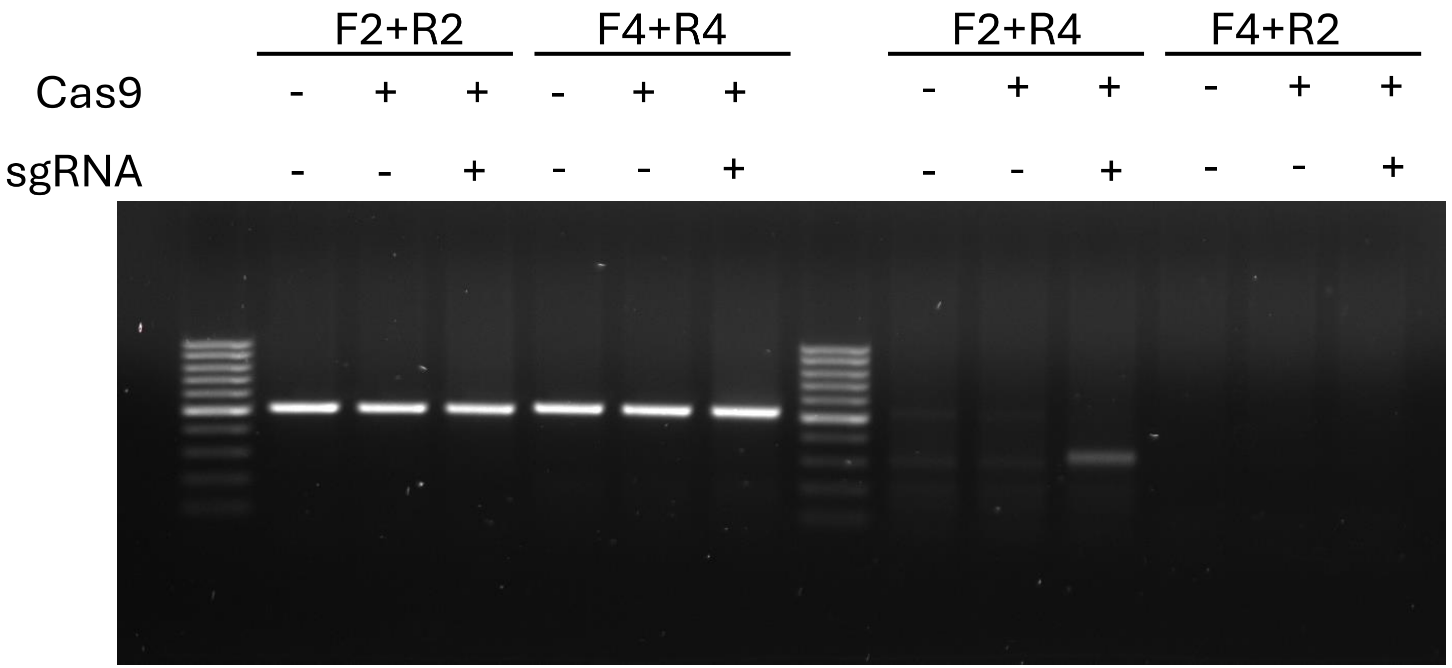

### Figure 6b

#### Colcemid 2 months

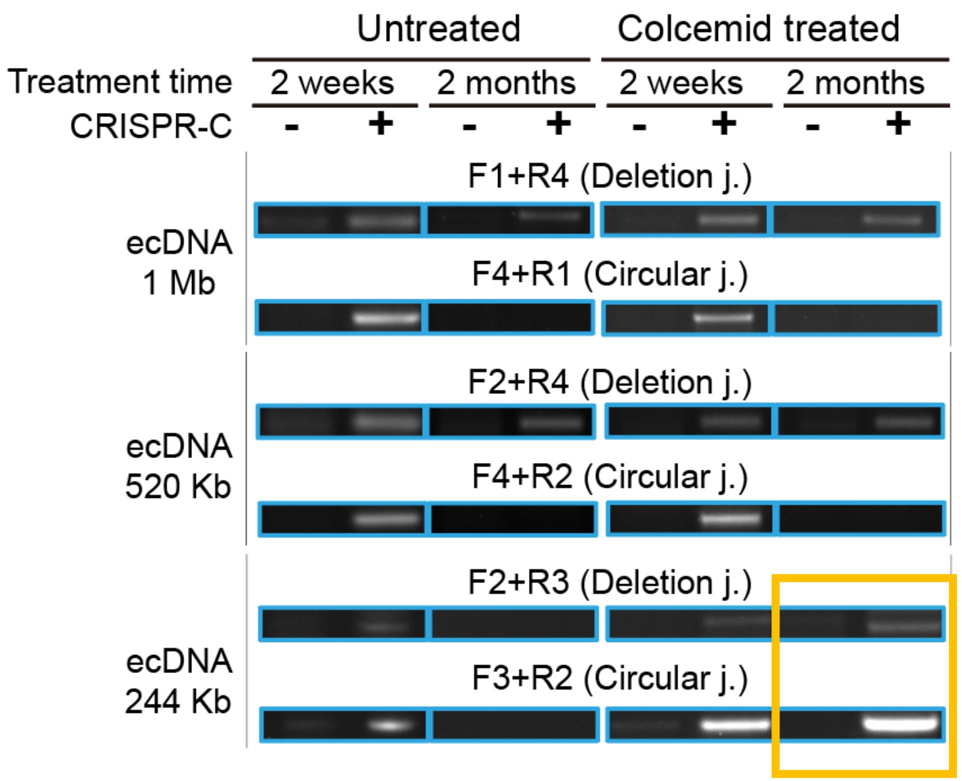


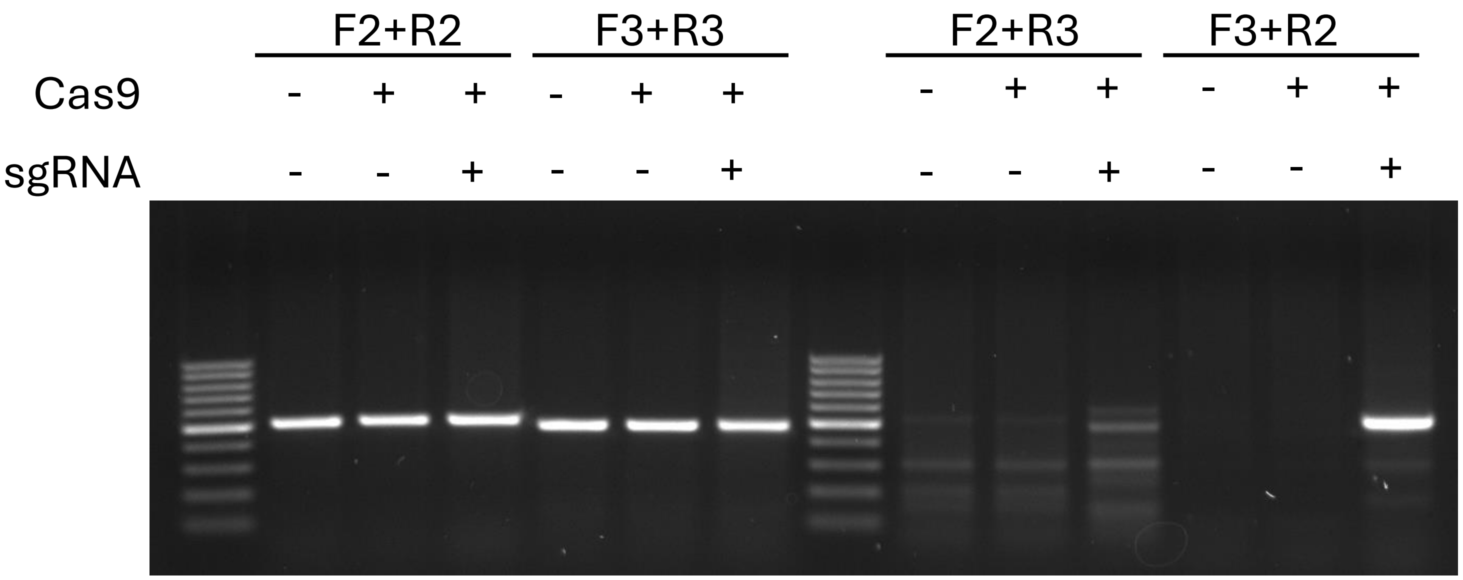

### Supplementary Figure S3

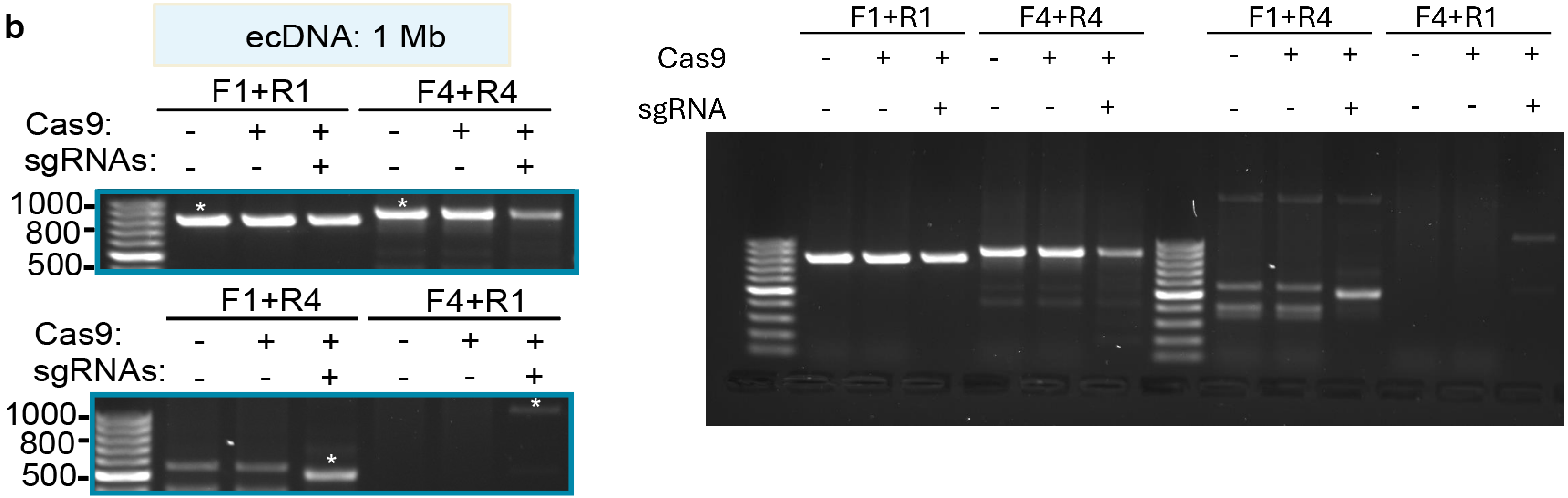

### Supplementary Figure S3

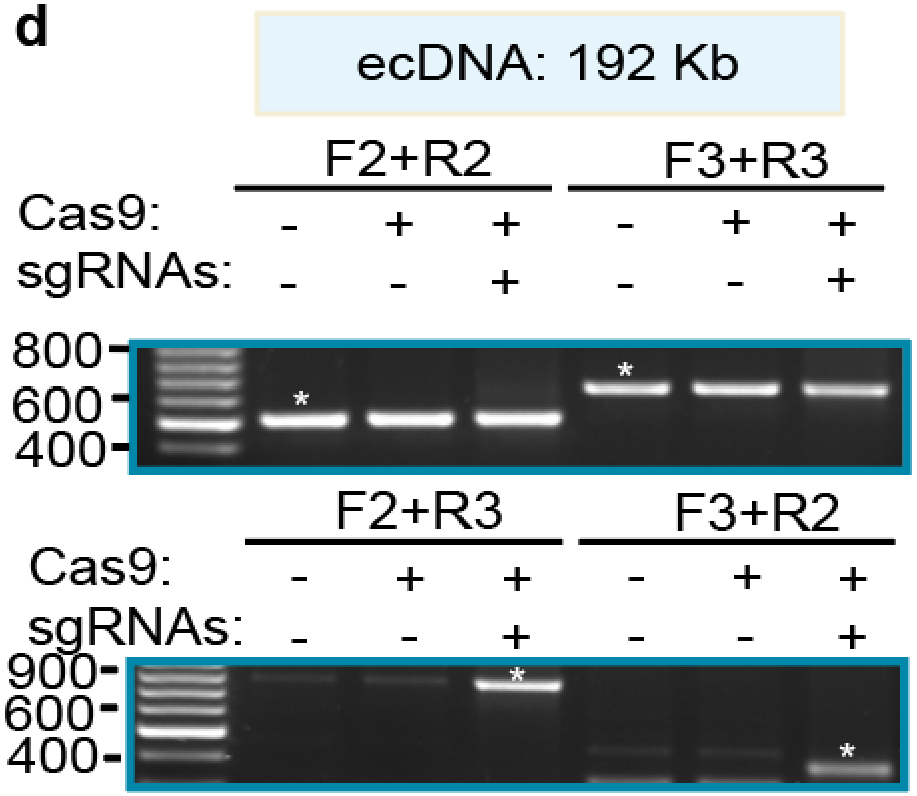


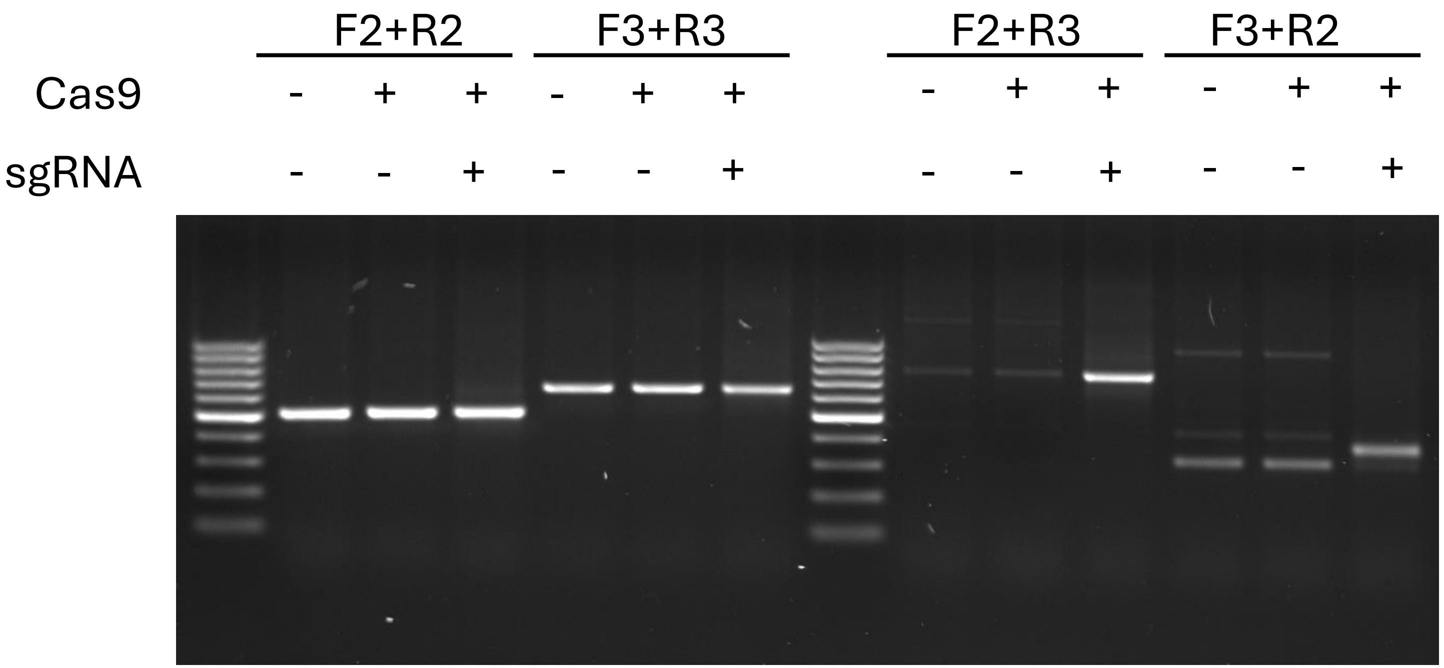

### Supplementary Figure S4

**b**

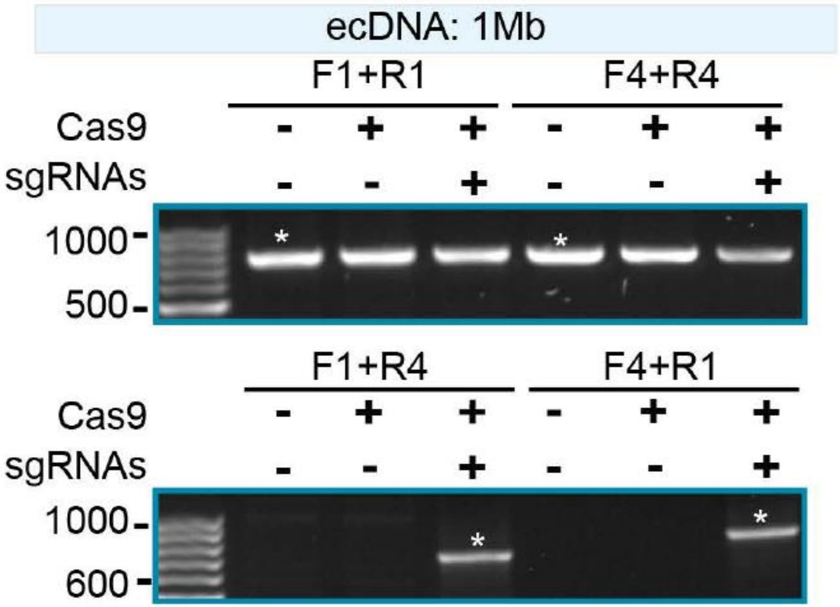


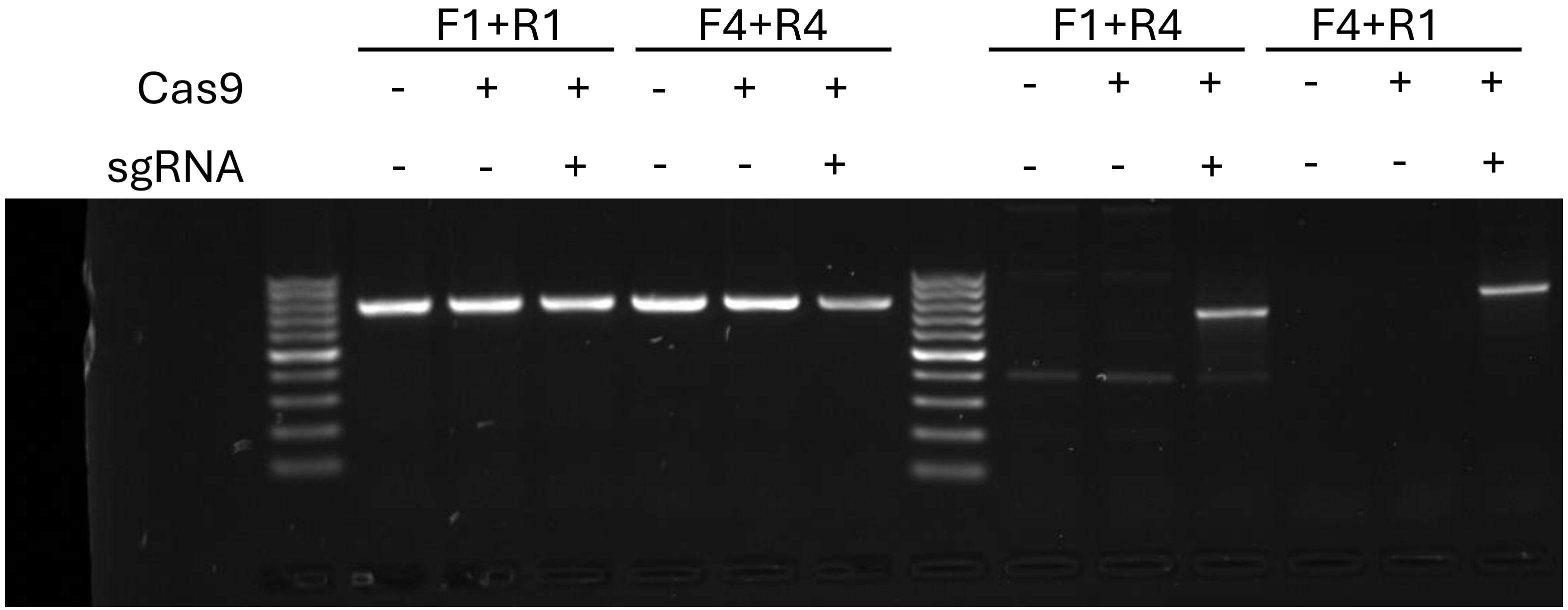

### Supplementary Figure S4

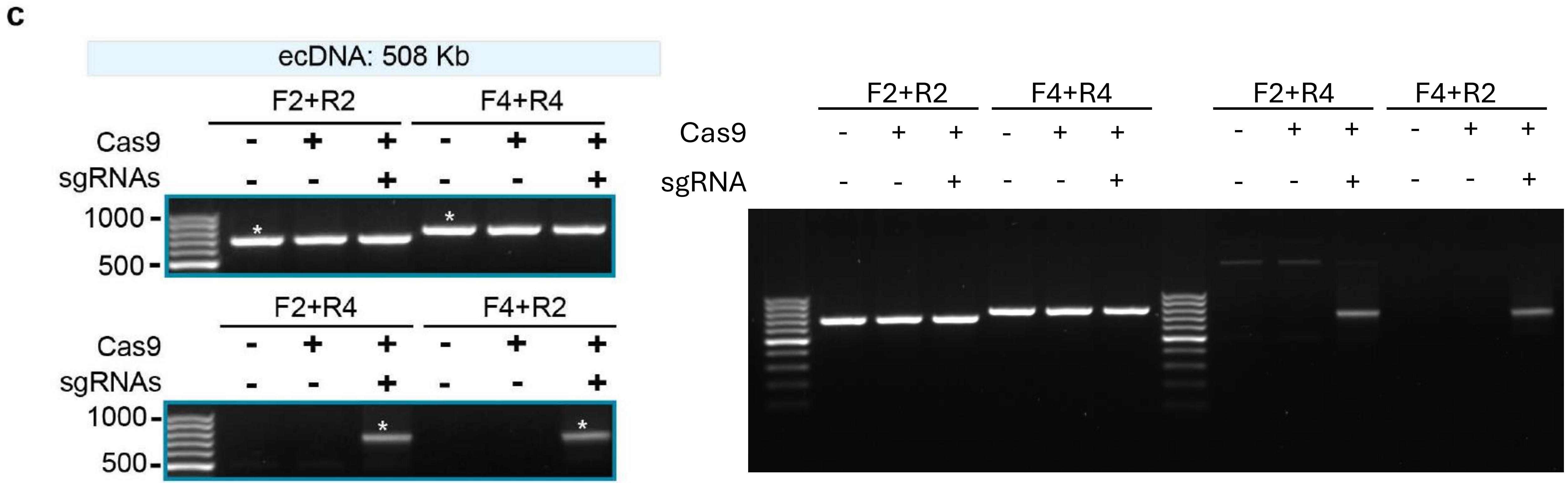

### Supplementary Figure S4

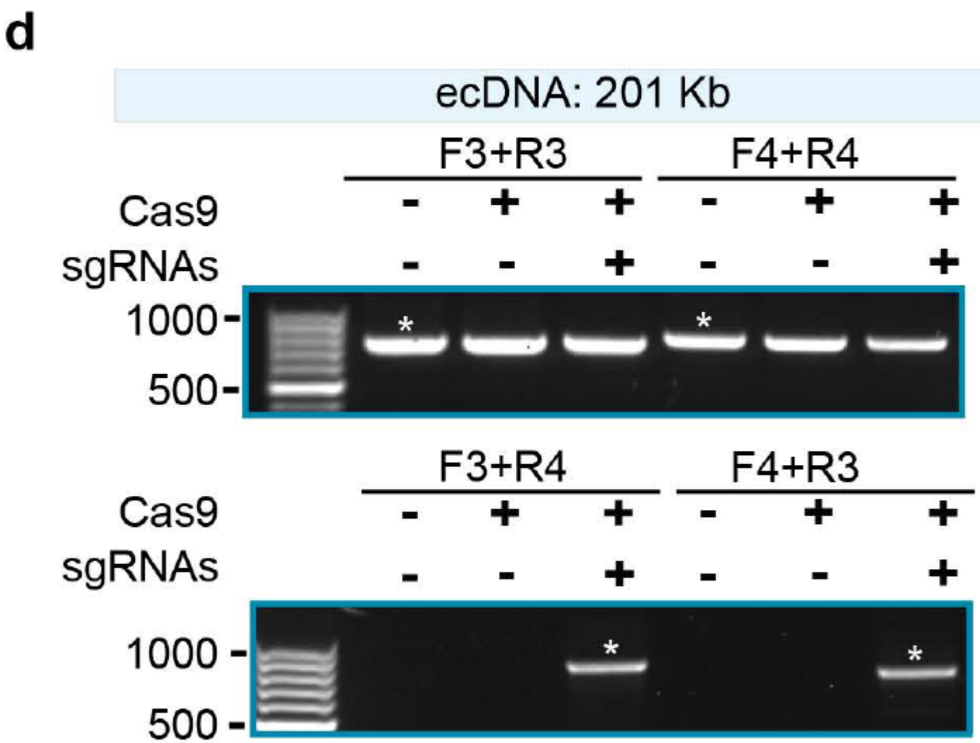


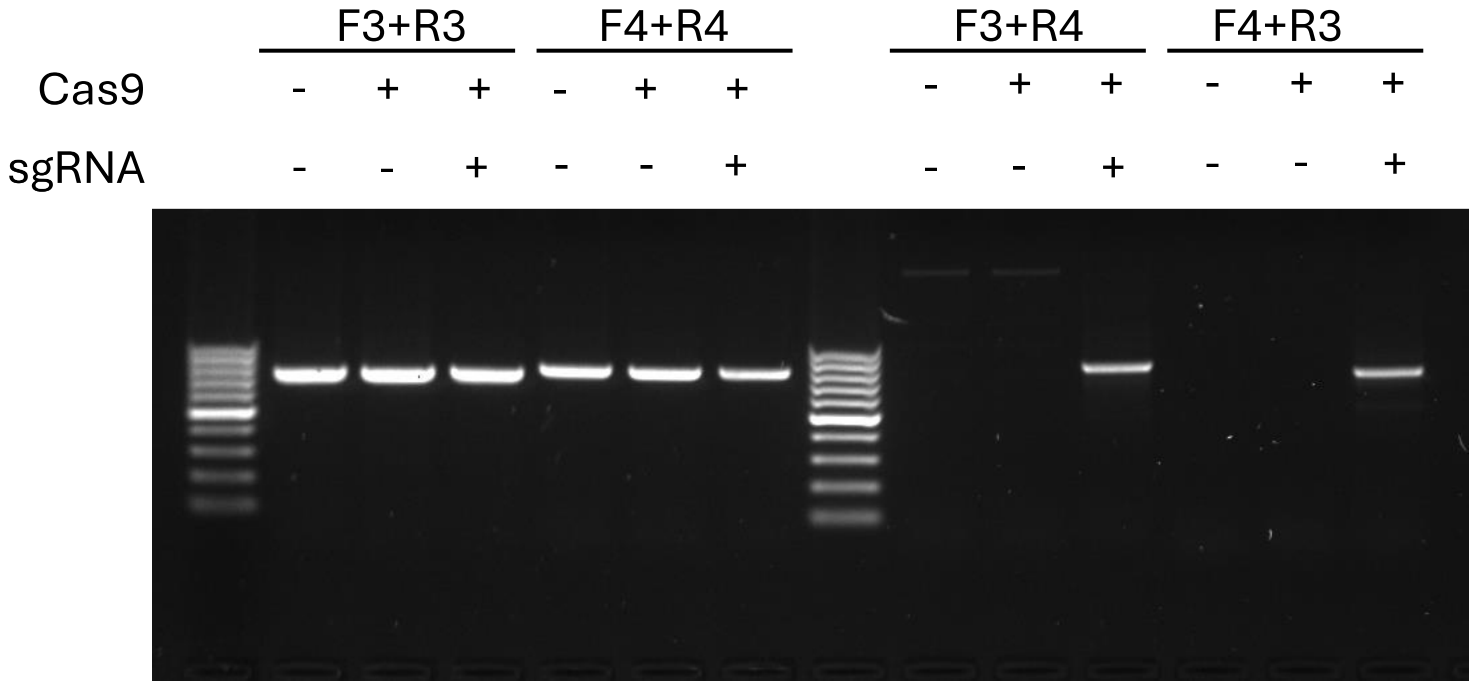

### Supplementary Figure S5

b

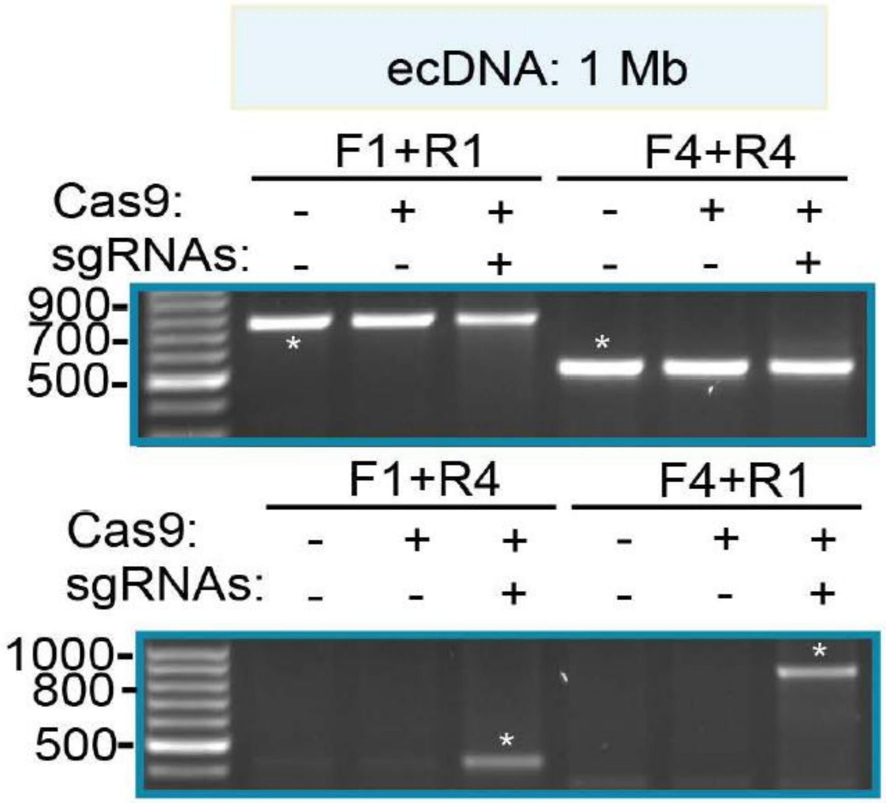


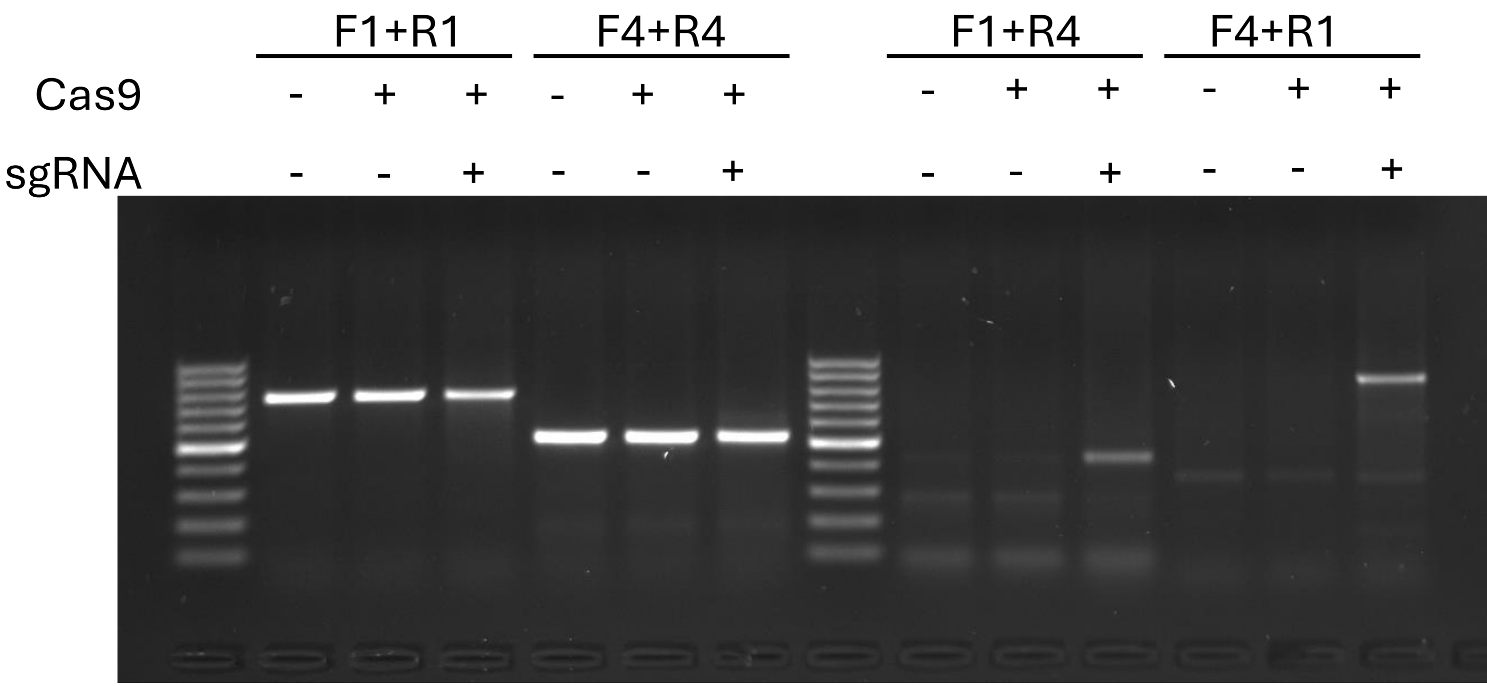

### Supplementary Figure S5

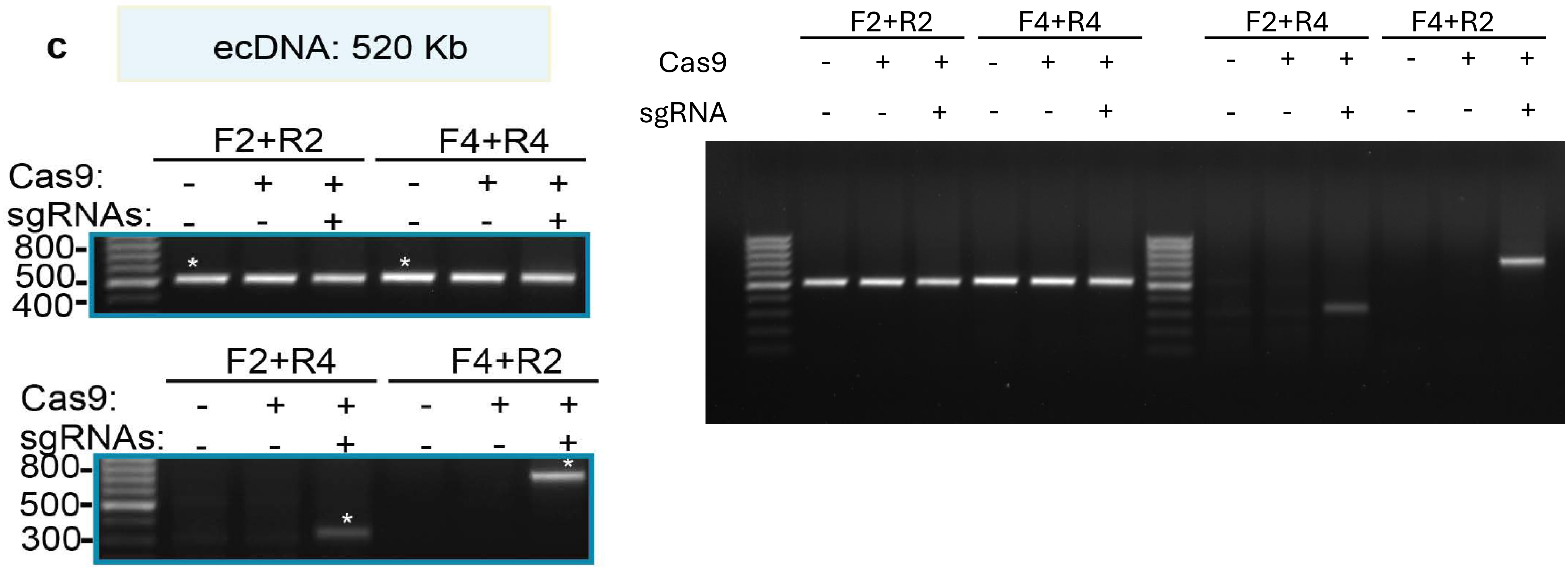

### Supplementary Figure S5

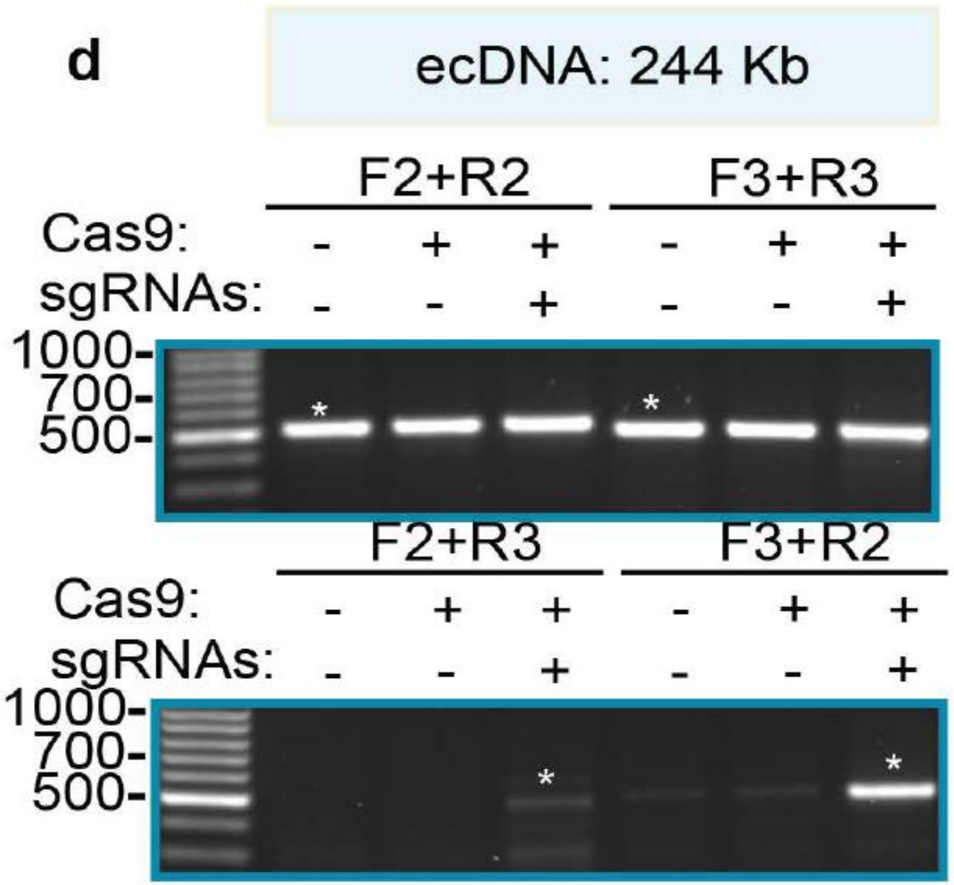


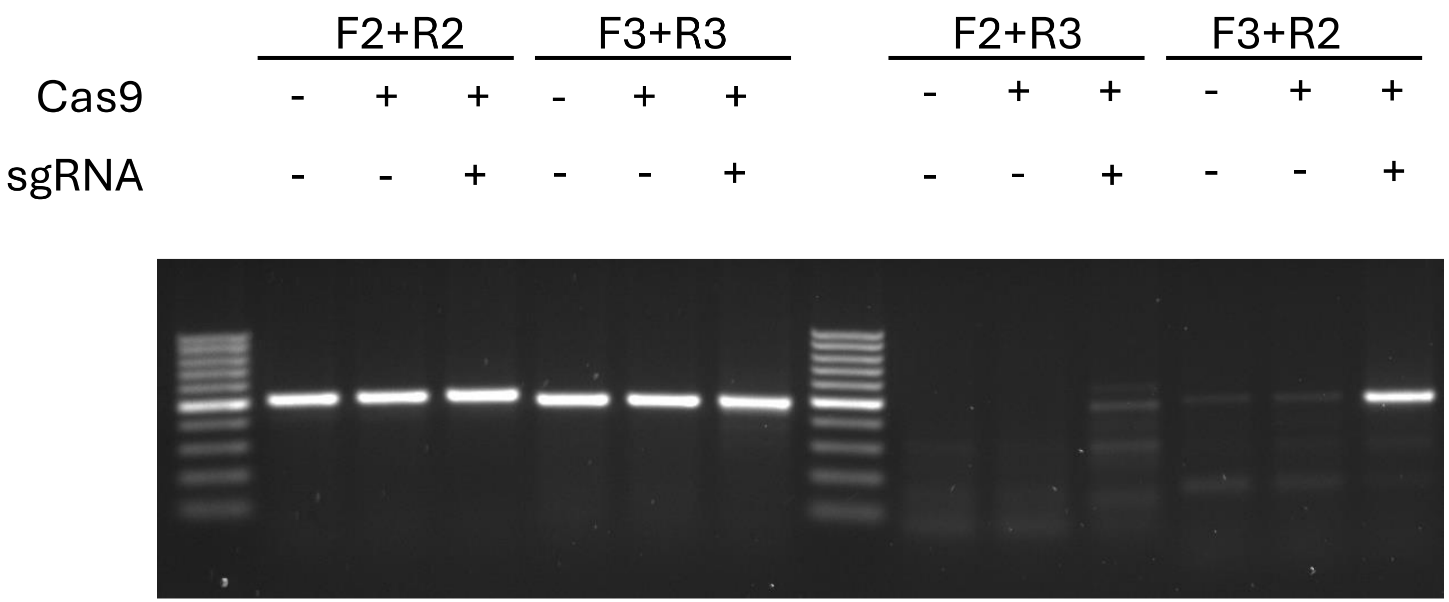

### Supplementary Figure S6

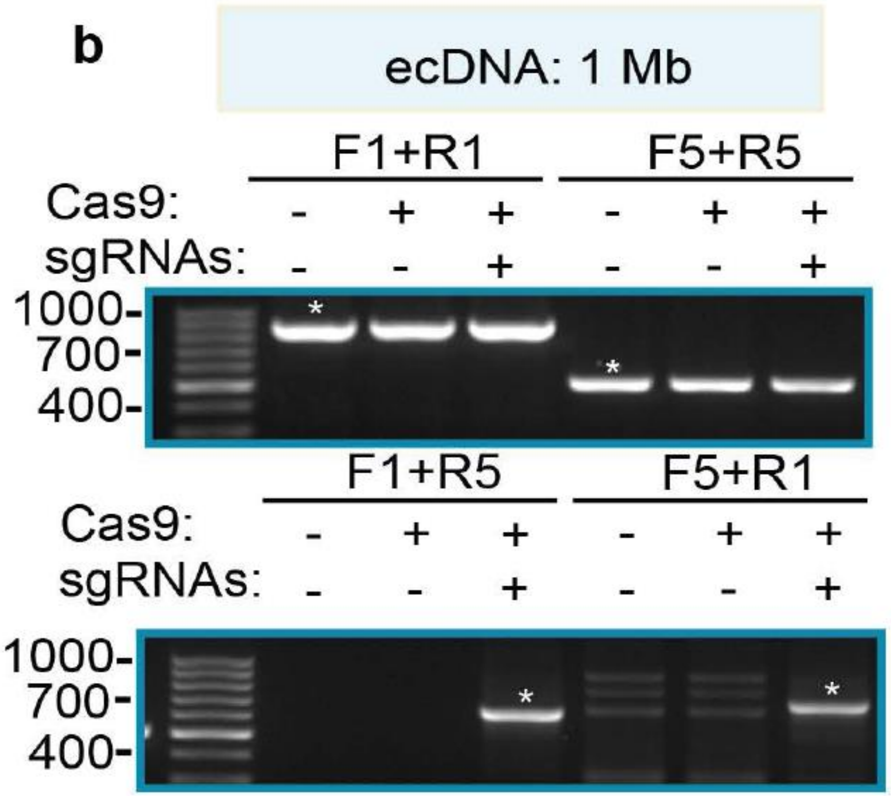


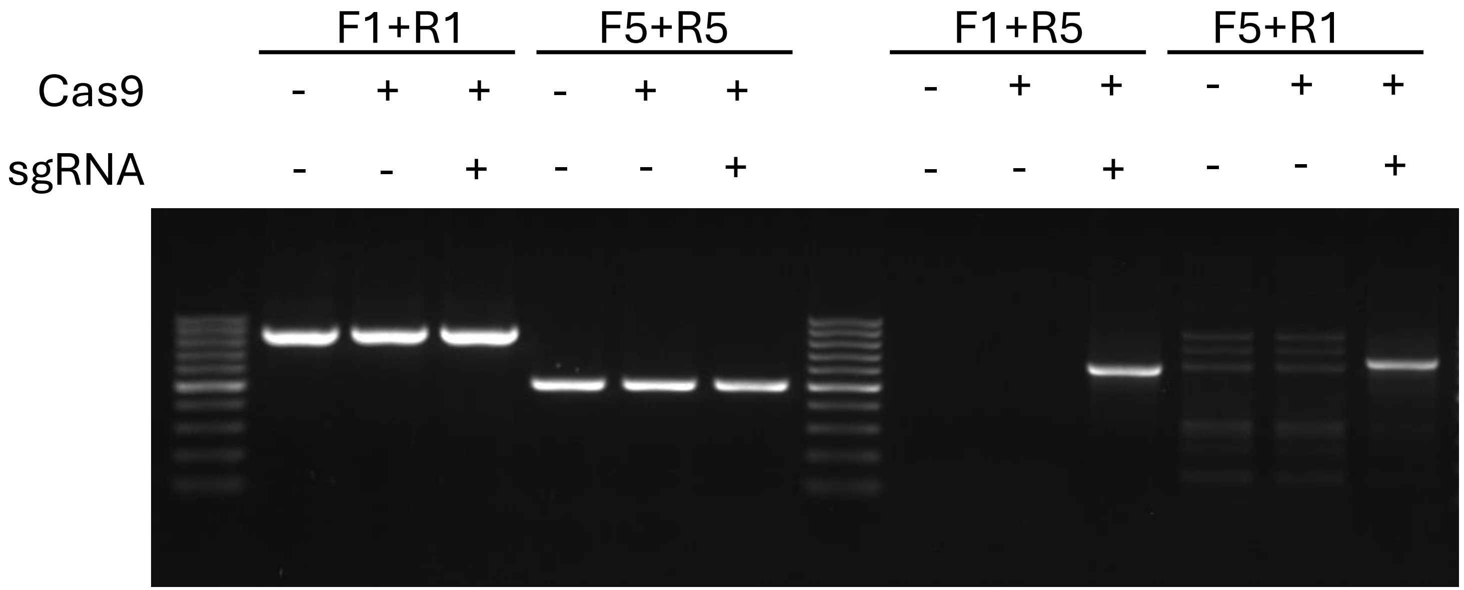

### Supplementary Figure S6

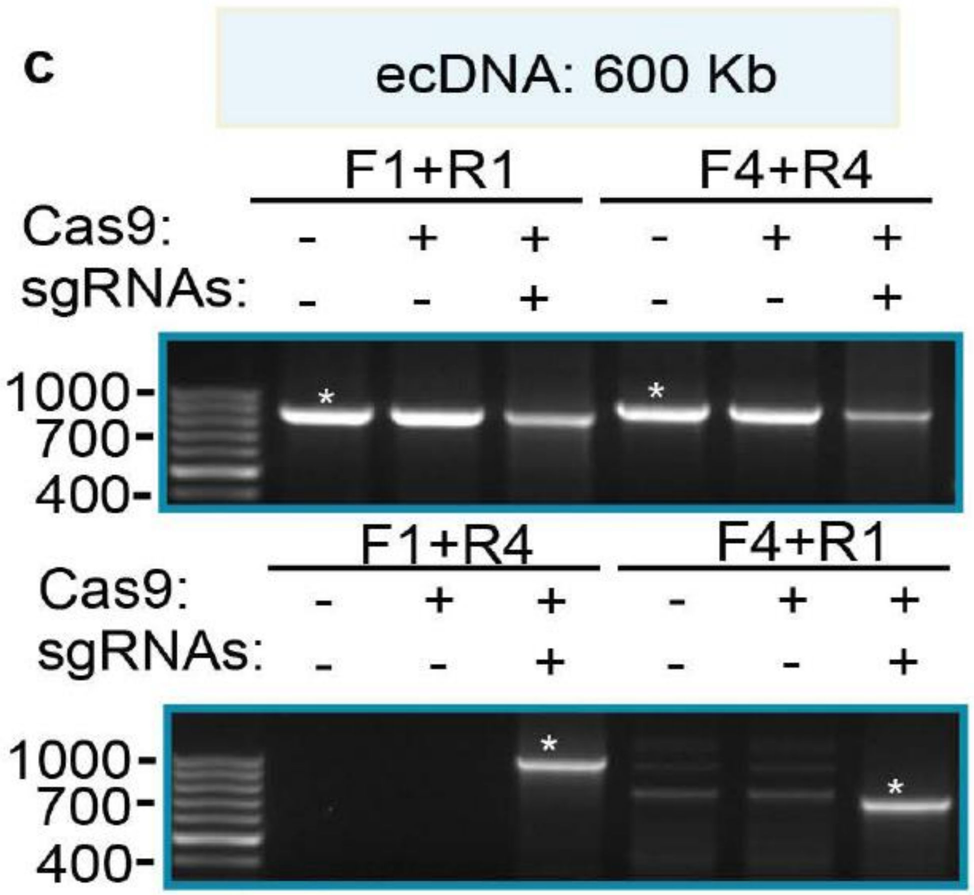


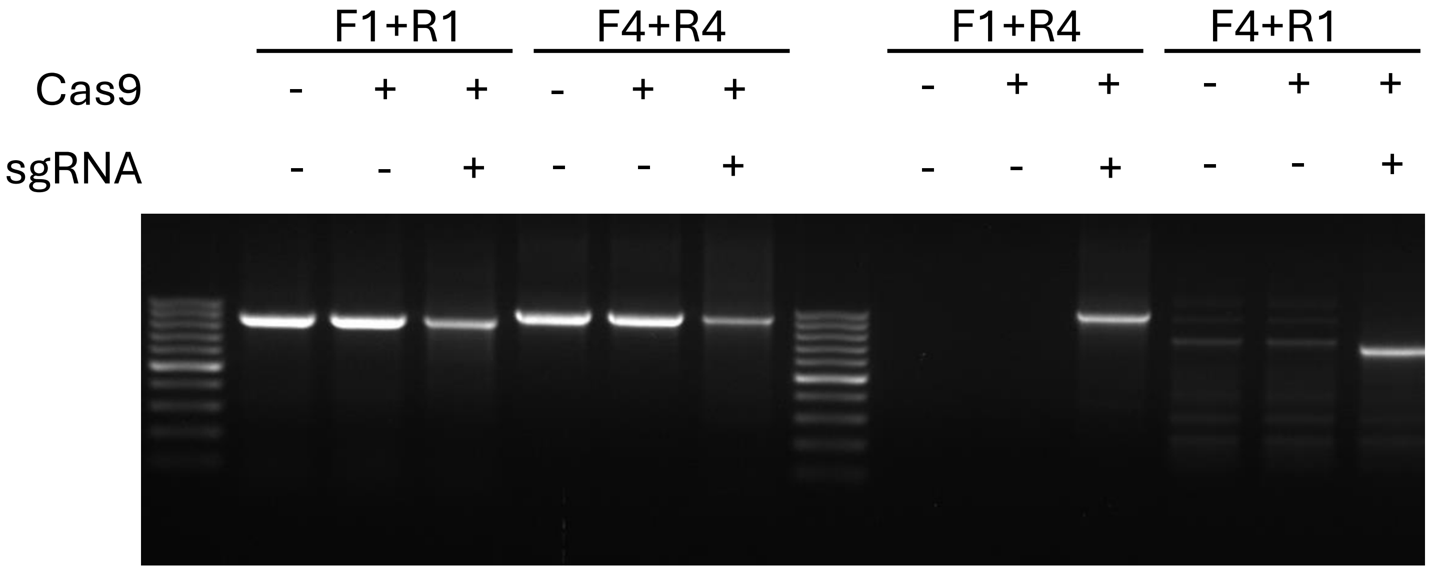

### Supplementary Figure S6

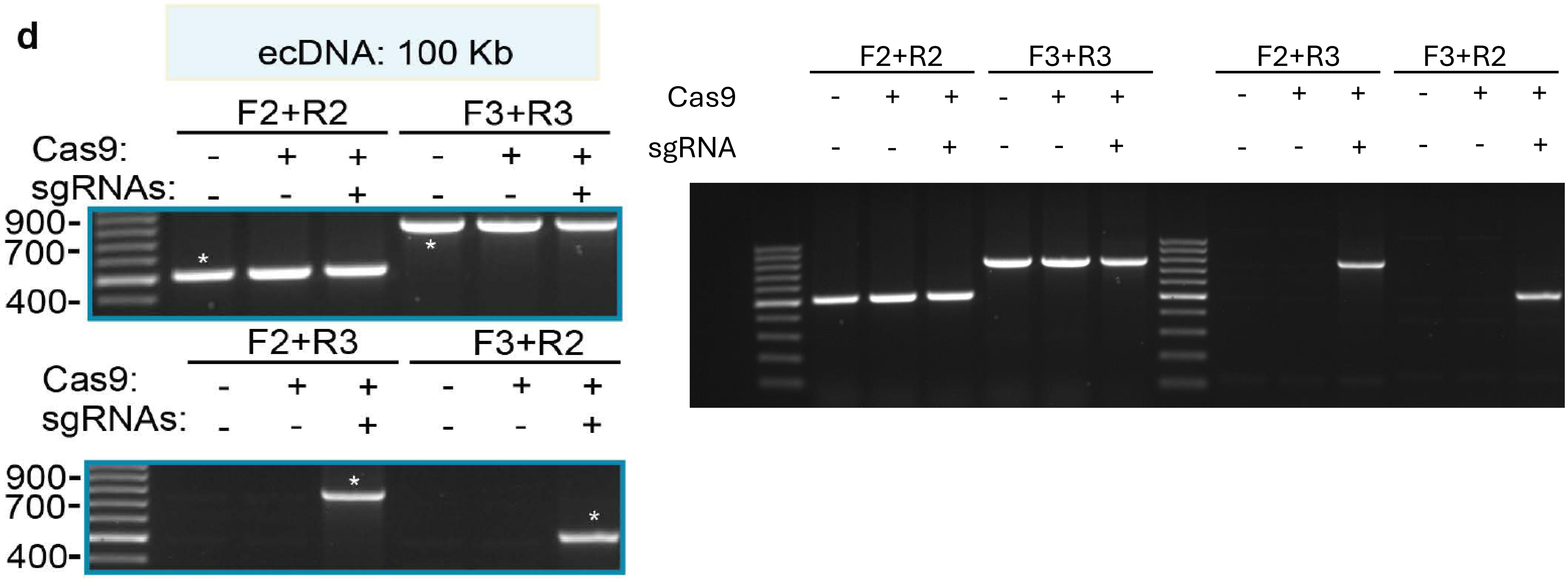

### Supplementary Figure S7

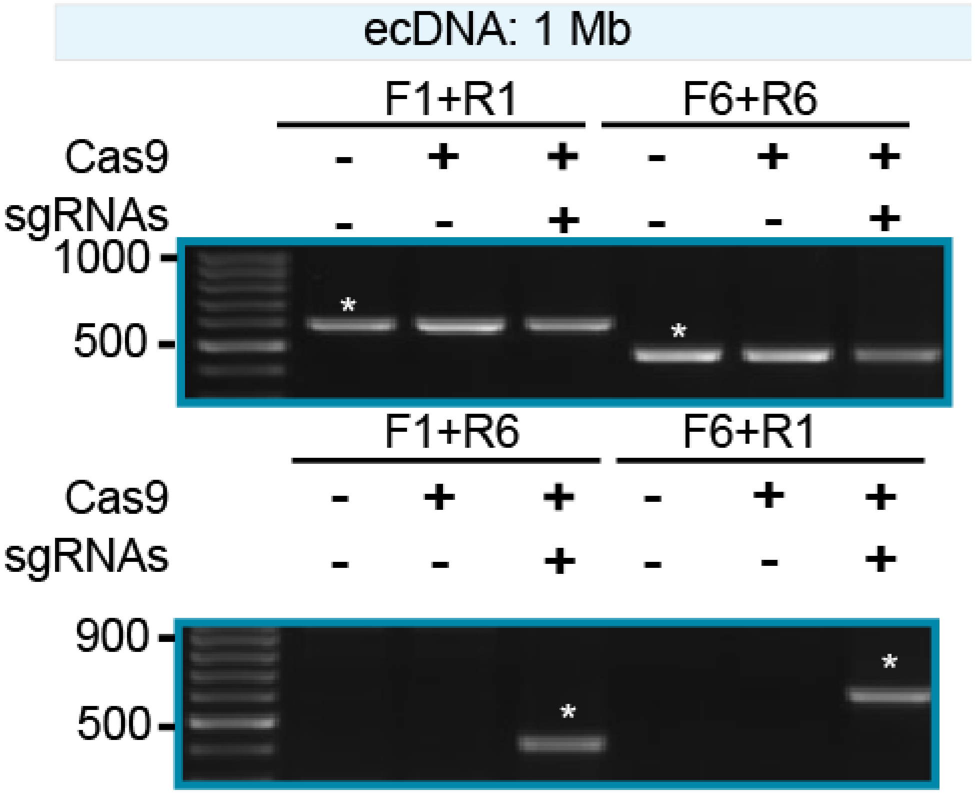


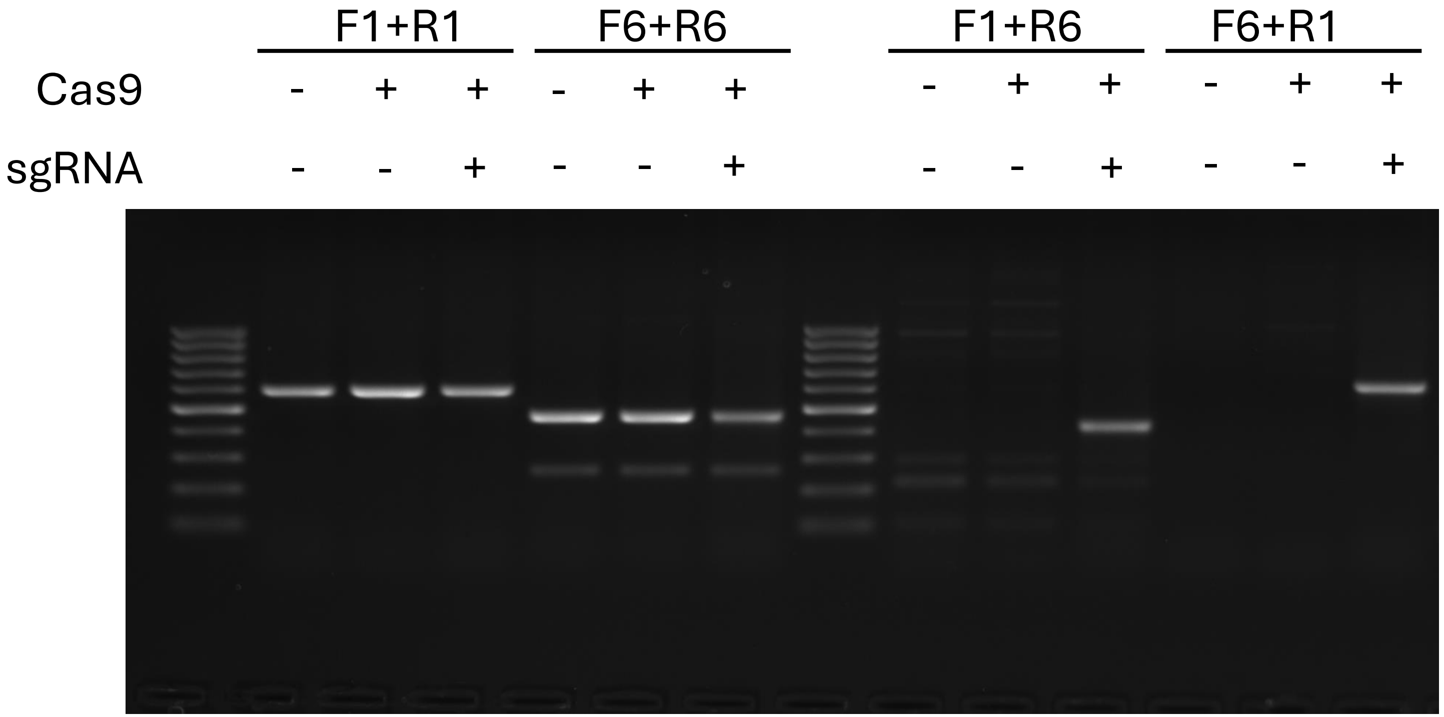

### Supplementary Figure S7

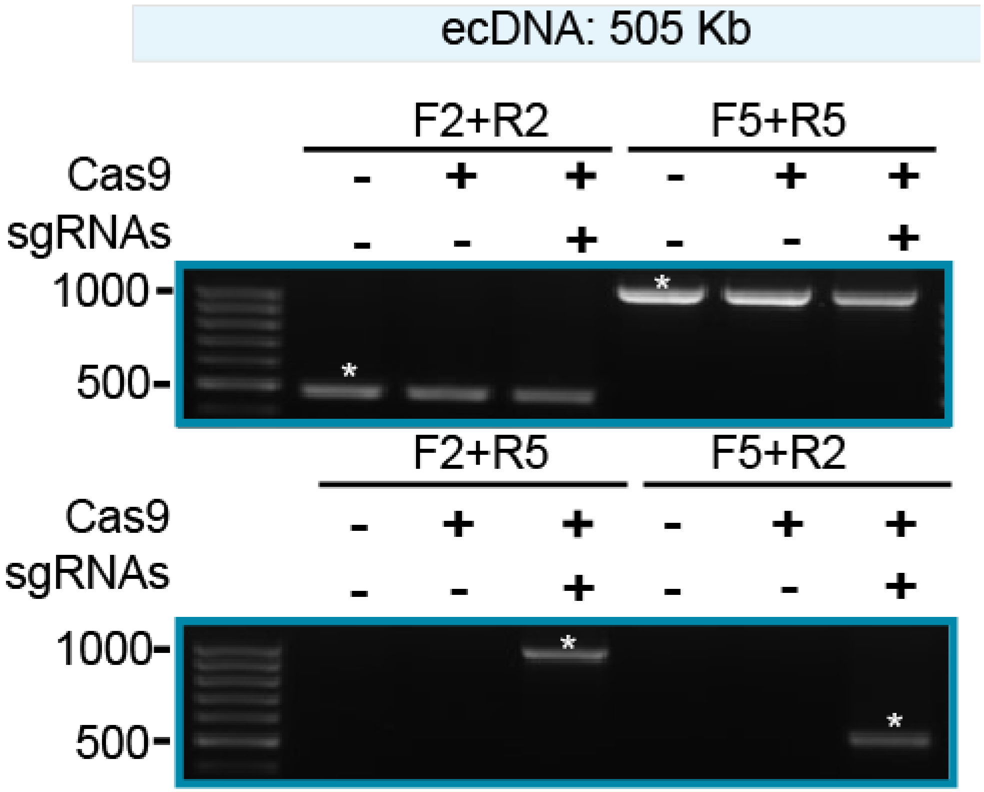


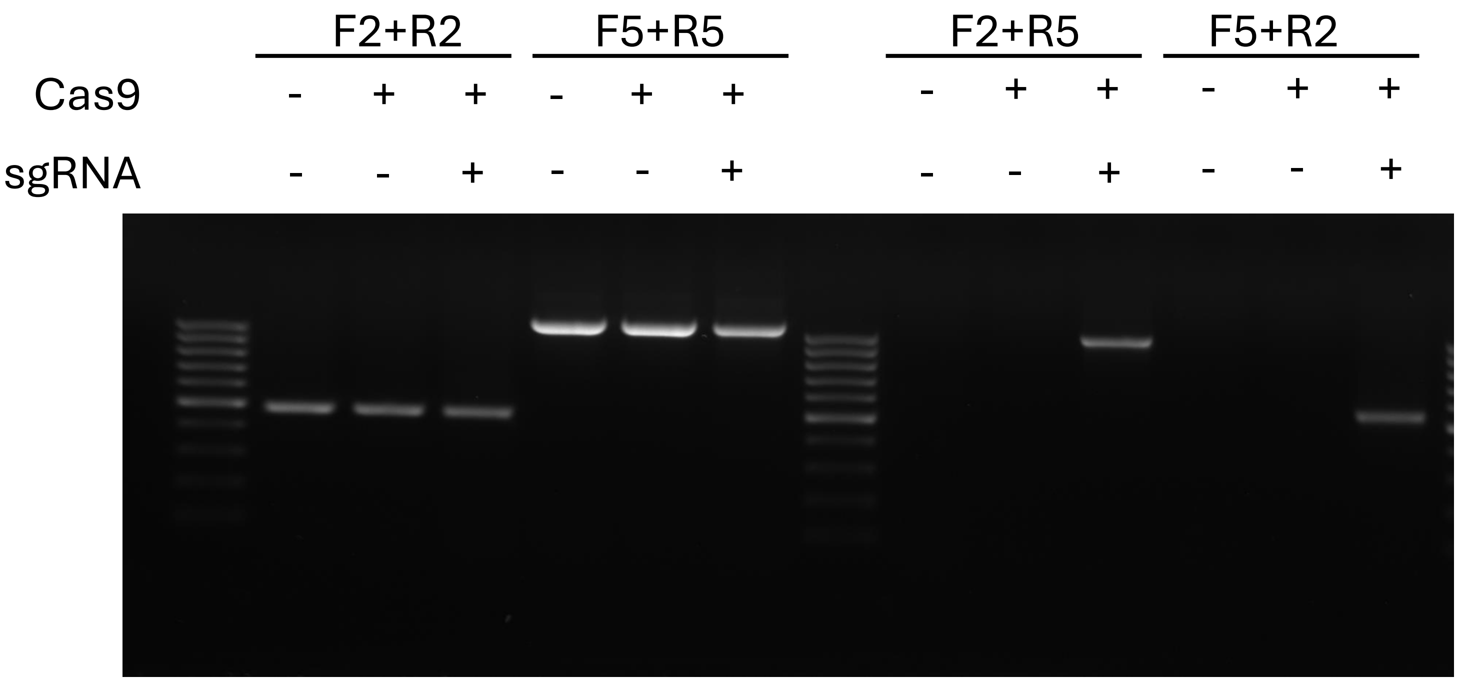

### Supplementary Figure S7

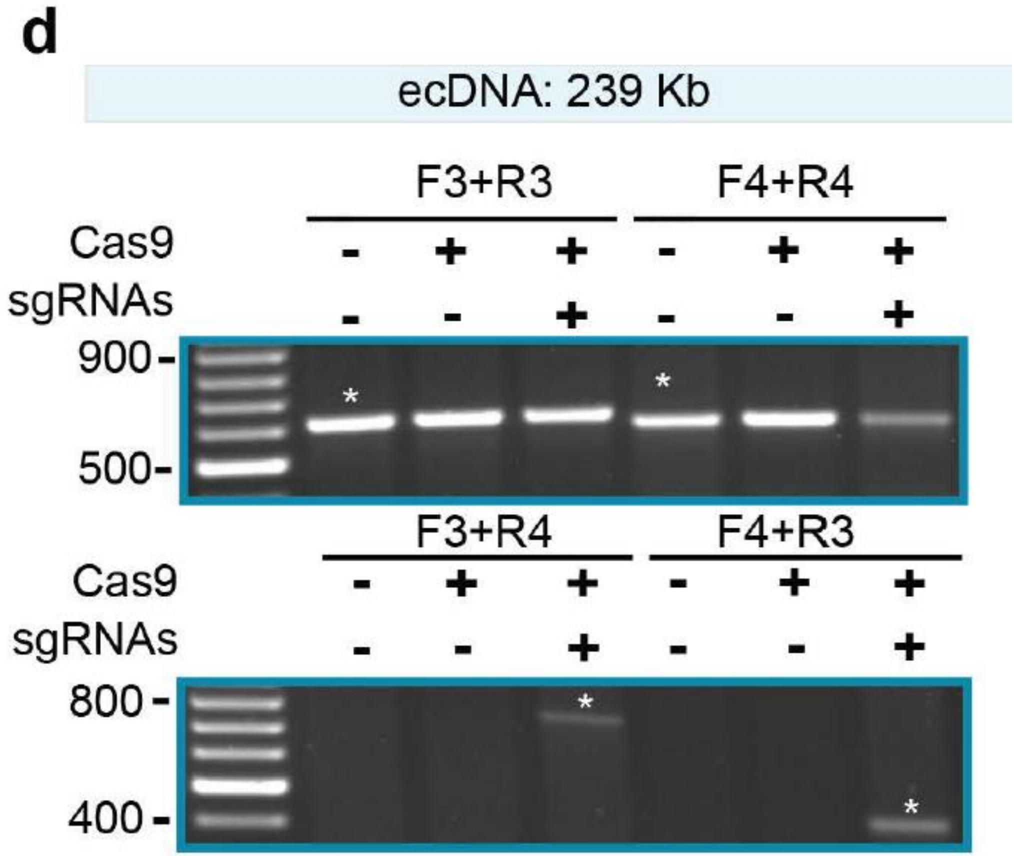


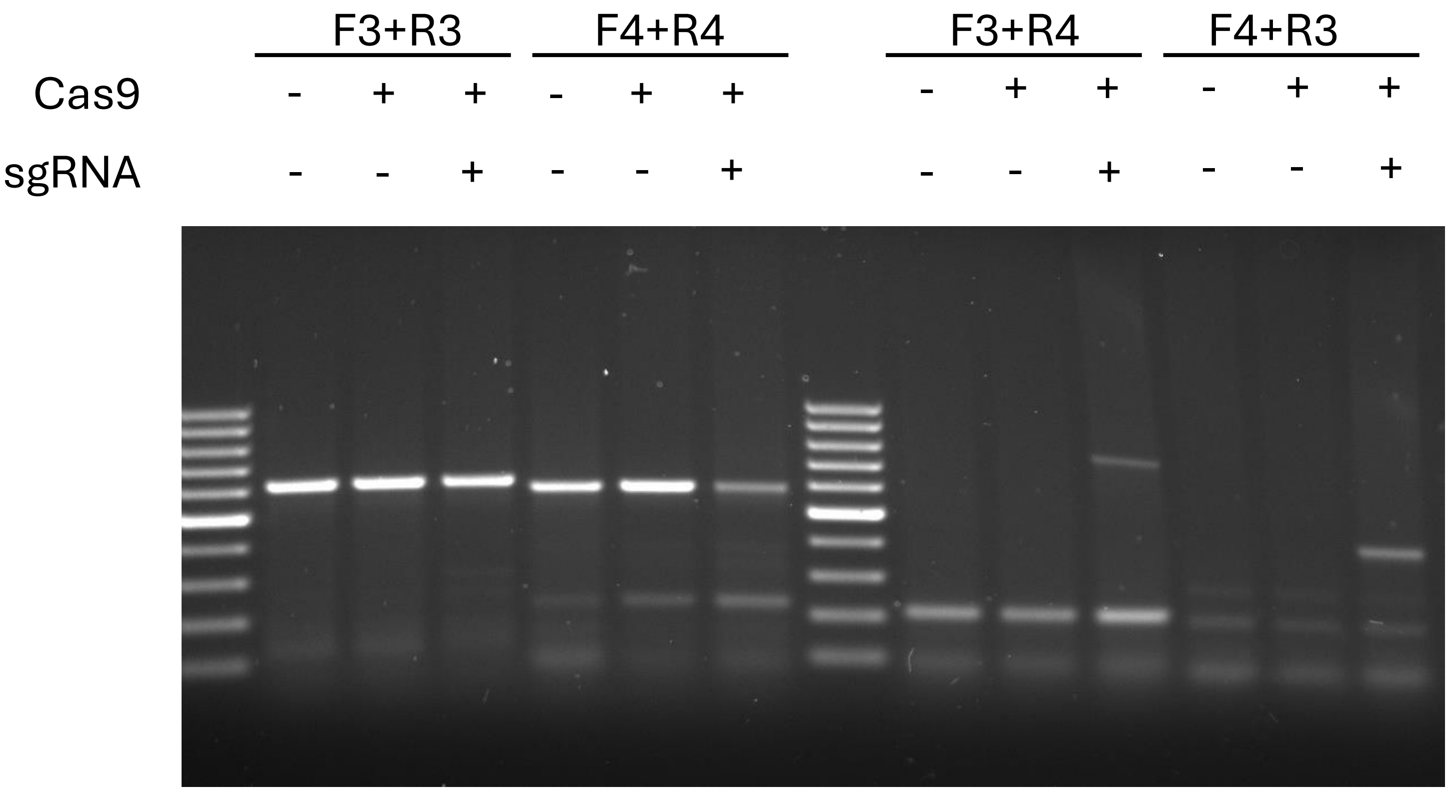

### Supplementary Figure S8

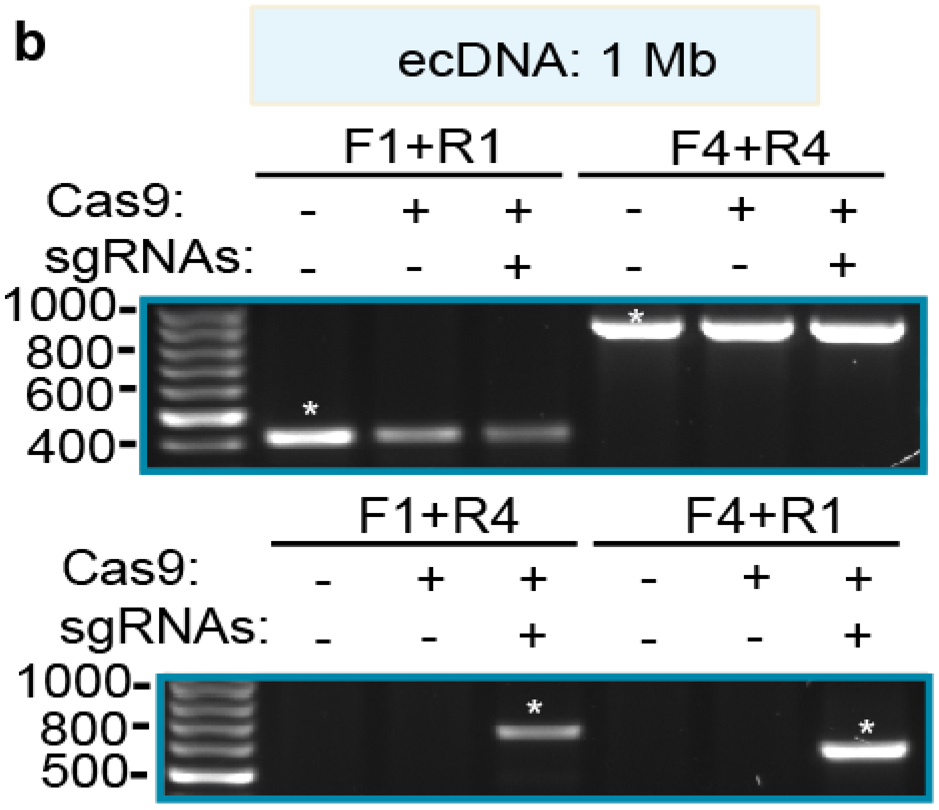


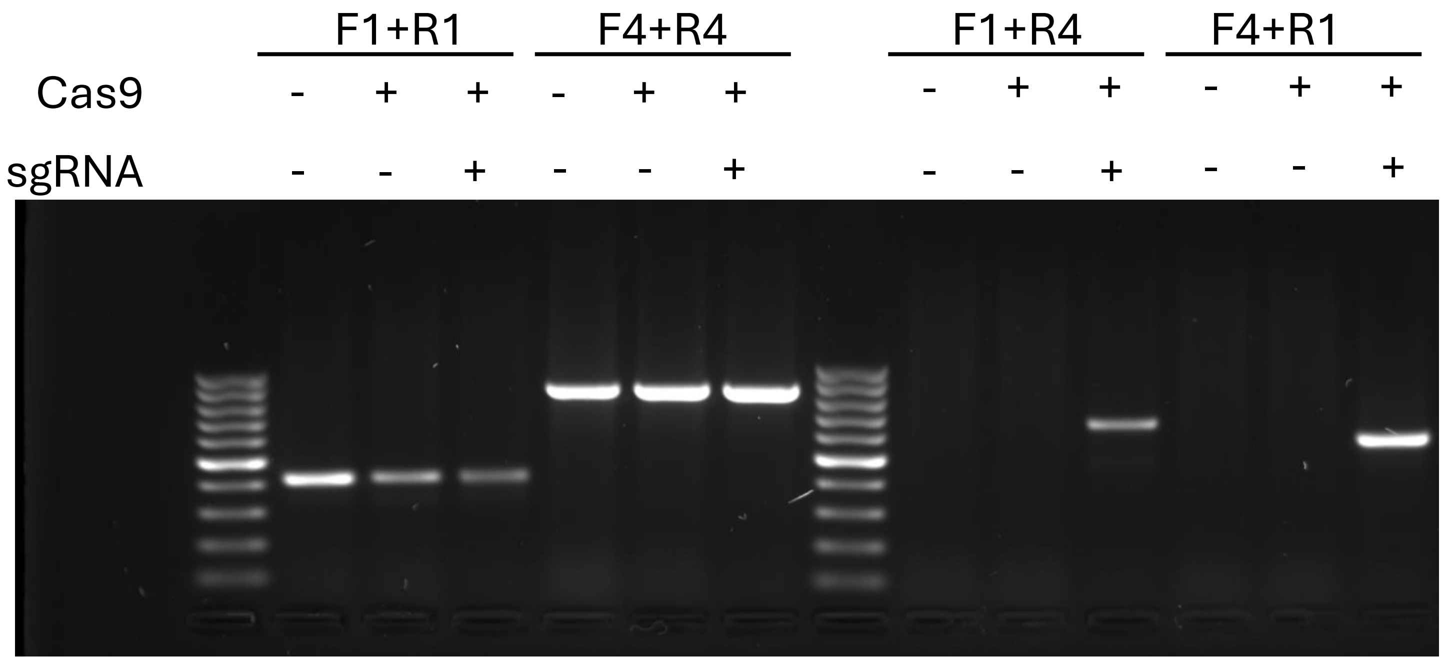

### Supplementary Figure S8

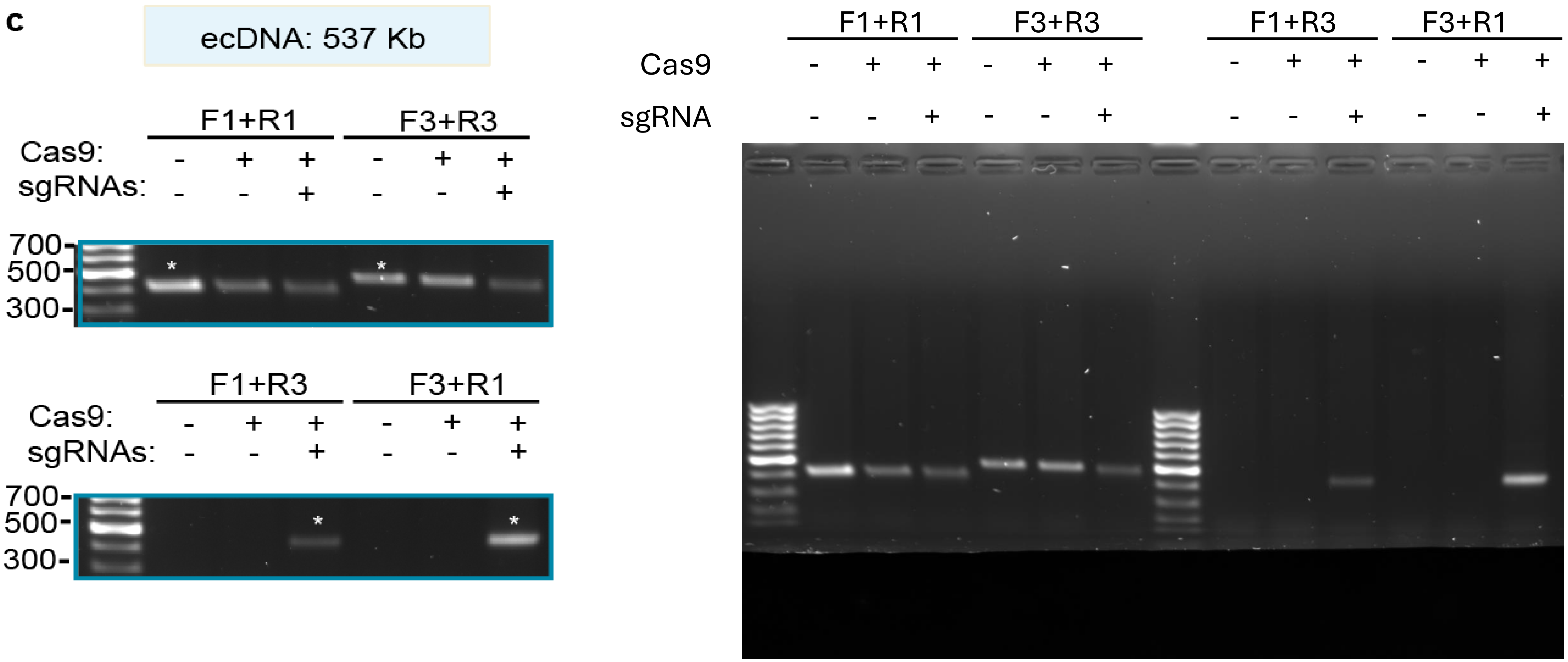

### Supplementary Figure S8

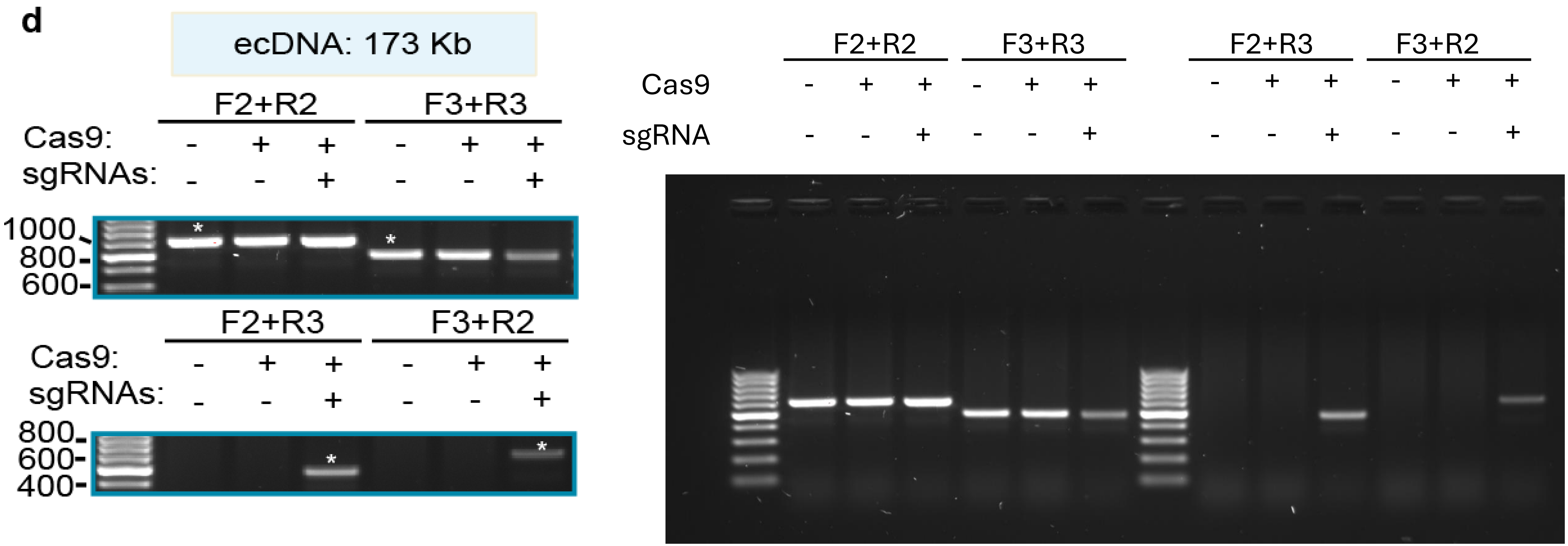

### Supplementary Figure S9

**b**

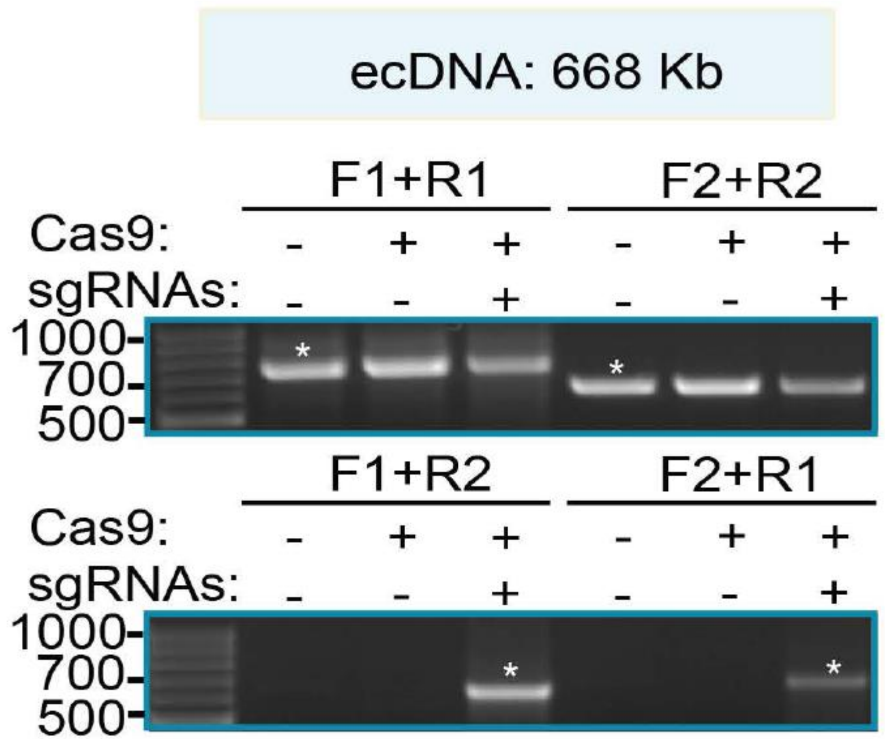


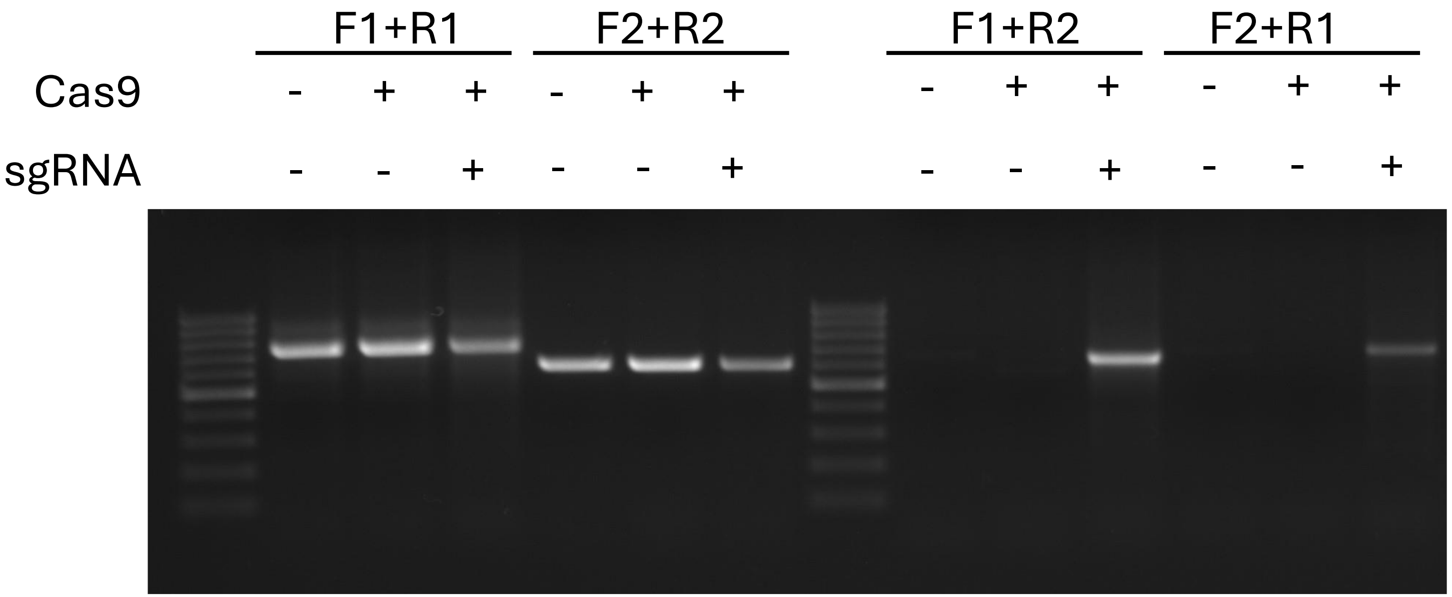

### Supplementary Figure S10

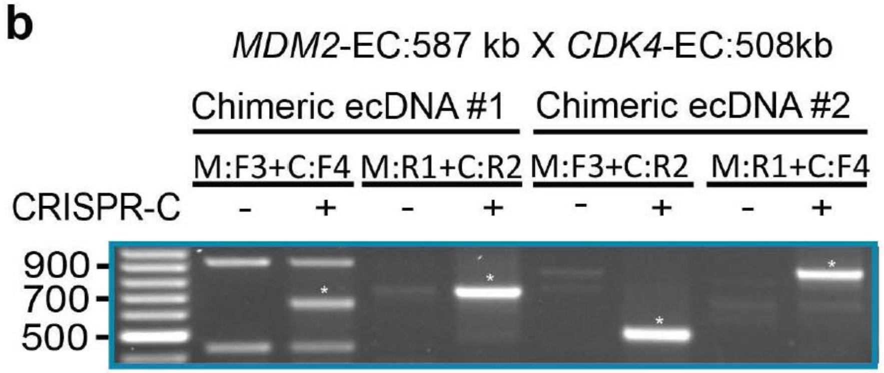


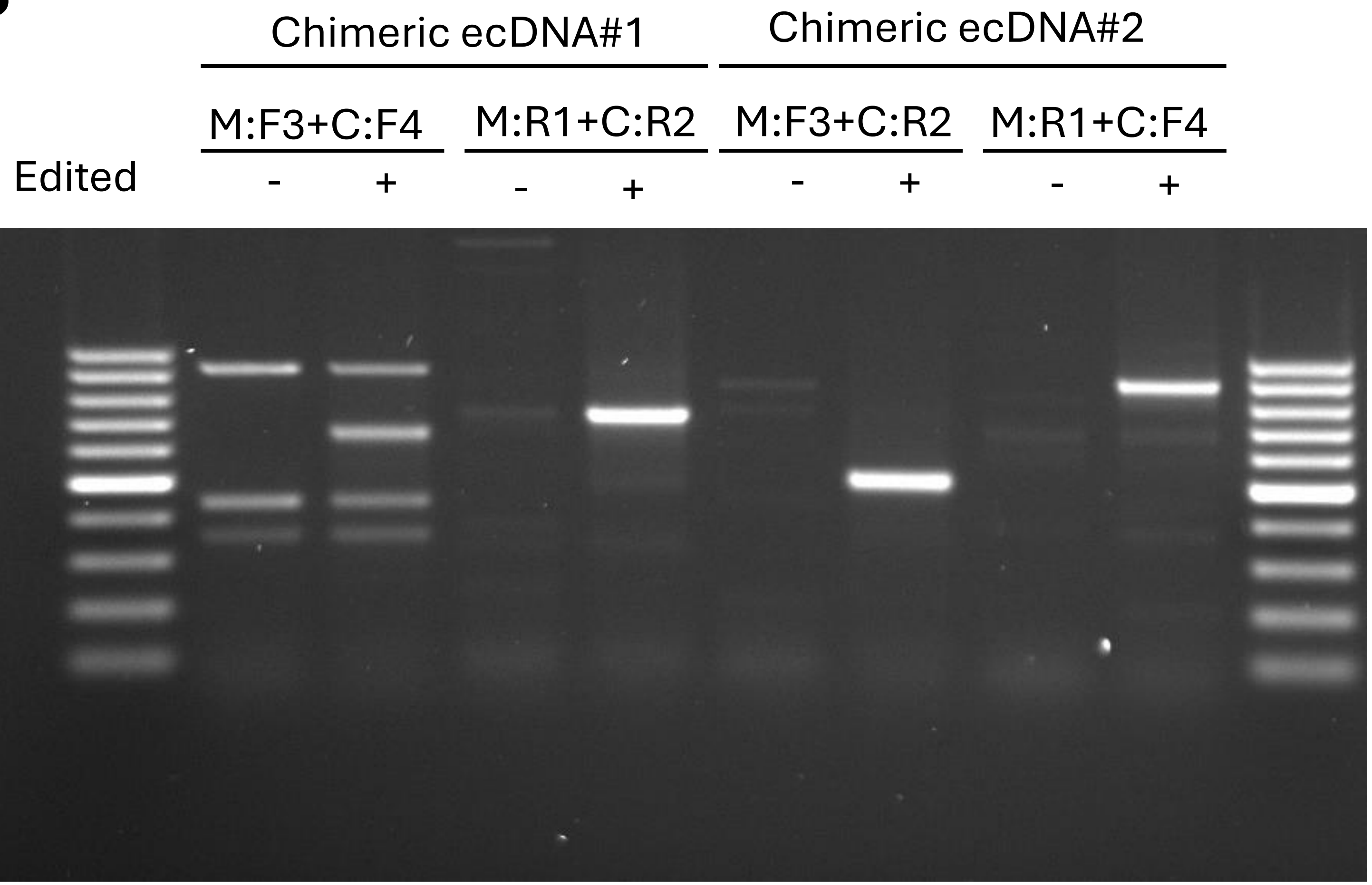

### Supplementary Figure S11

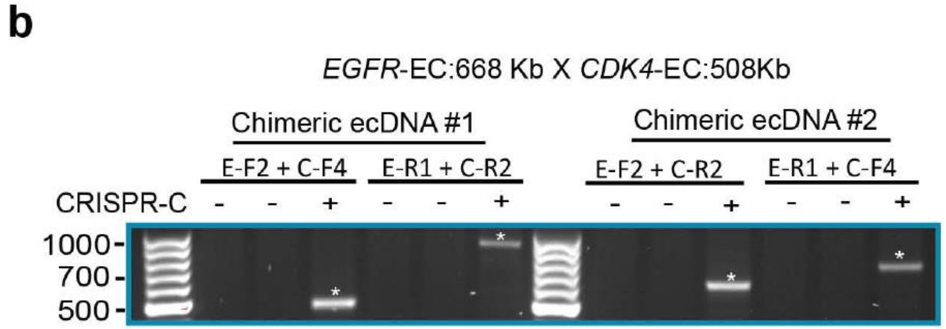


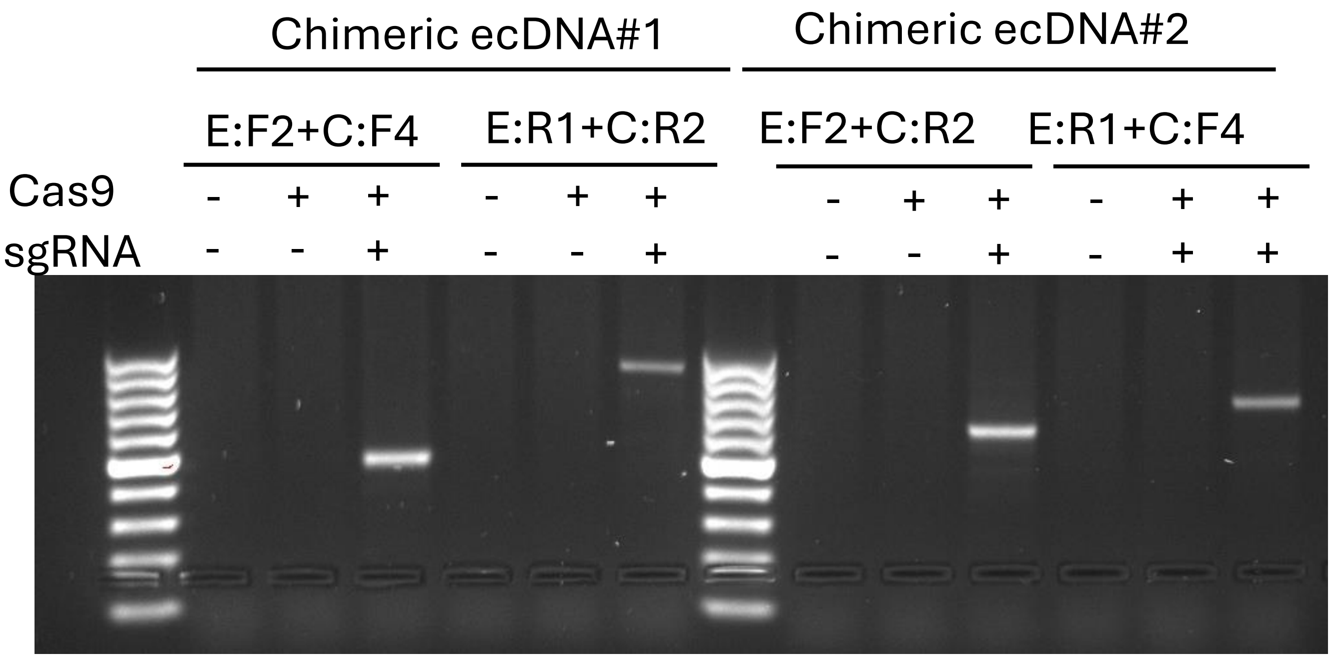

### Supplementary Figure S12

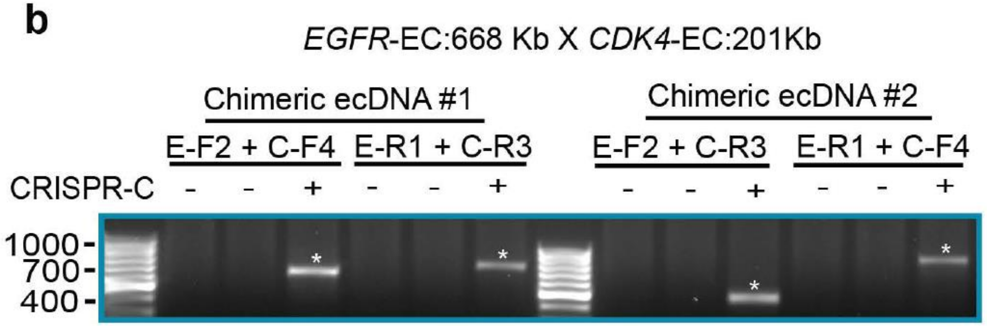


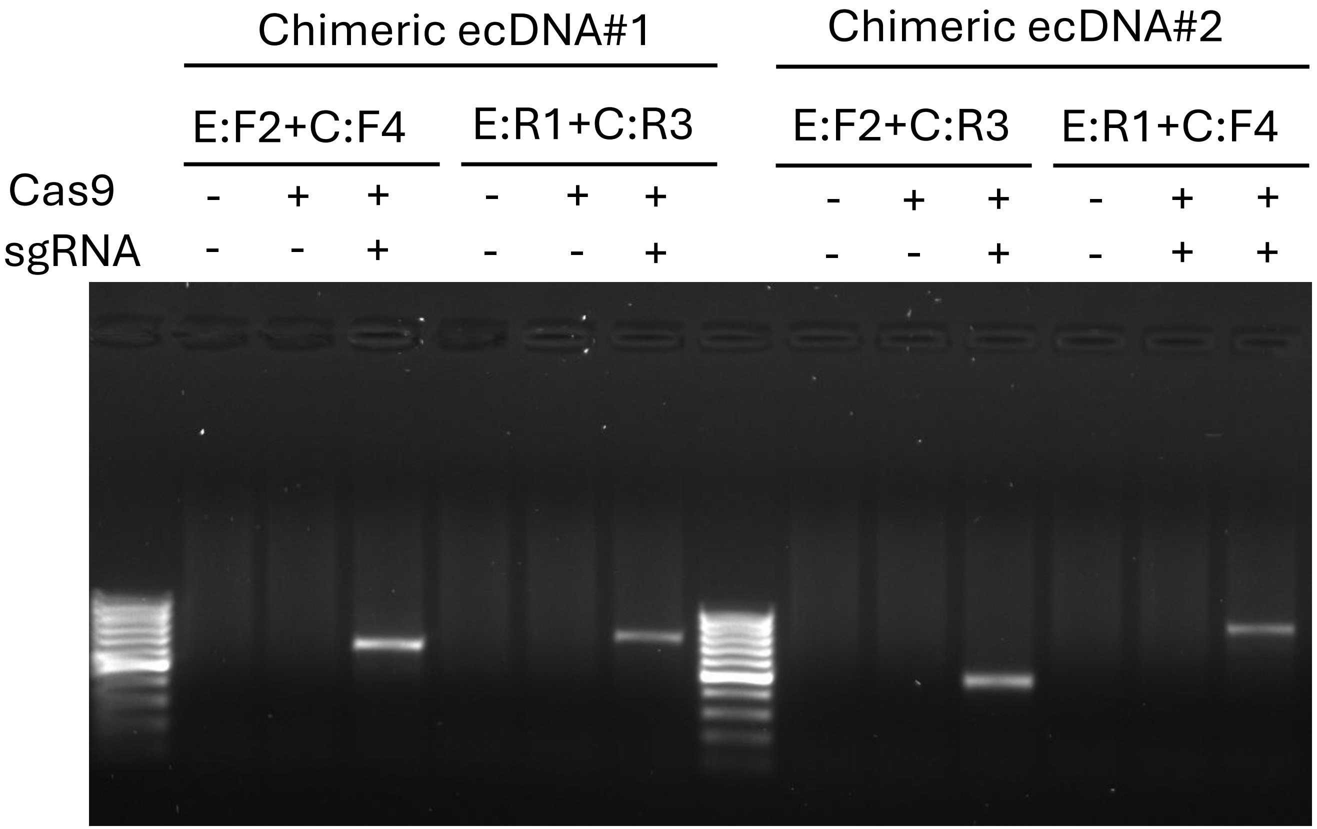

### Supplementary Figure S30

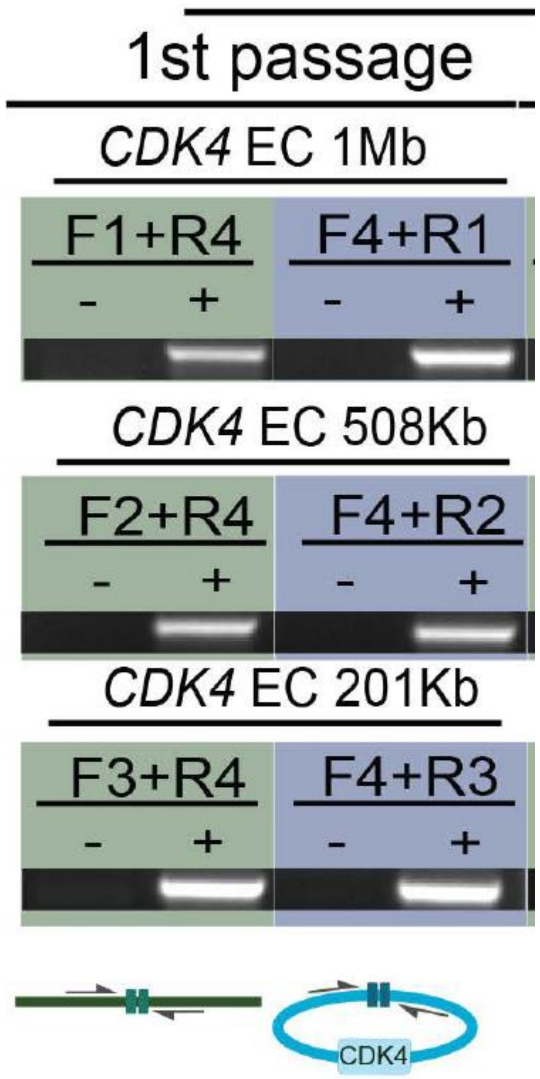


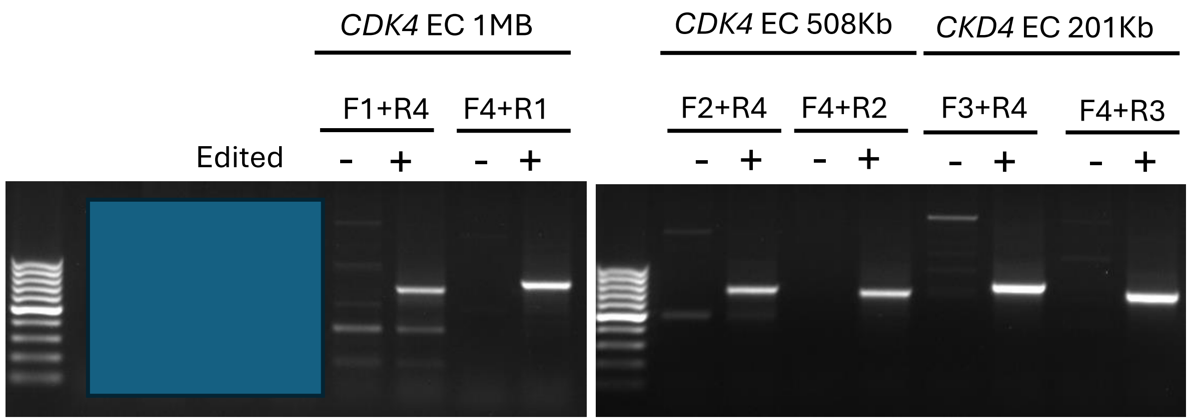


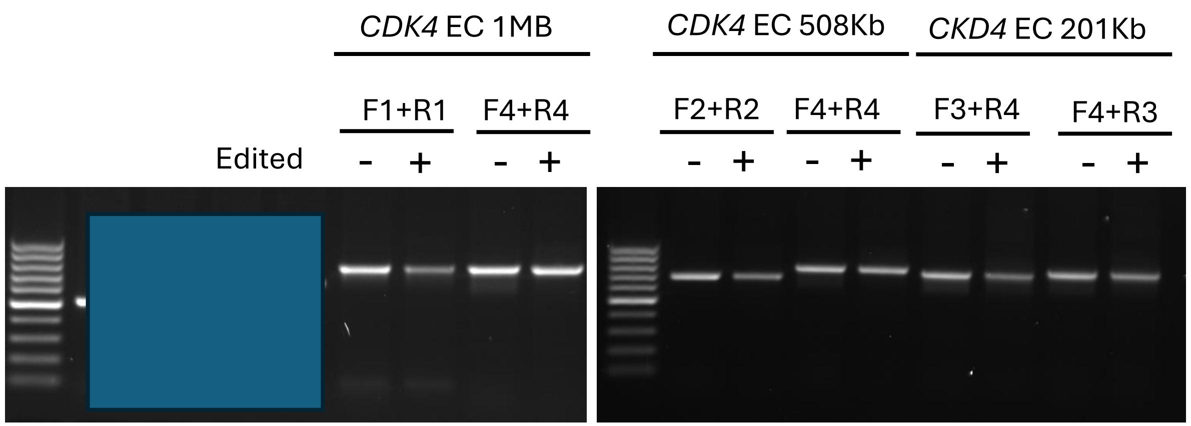


**Data not shown on Figure**

### Supplementary Figure S30

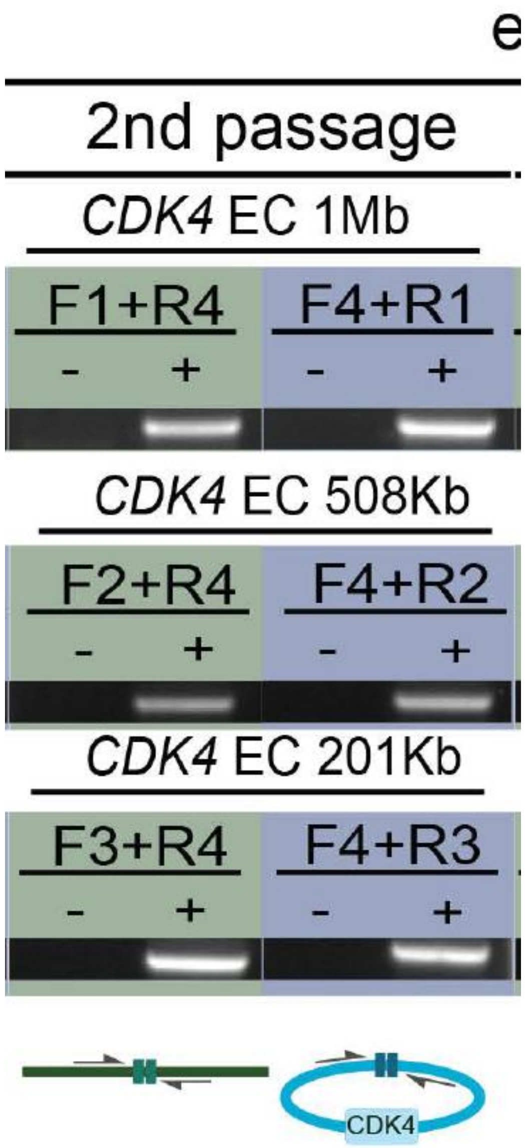


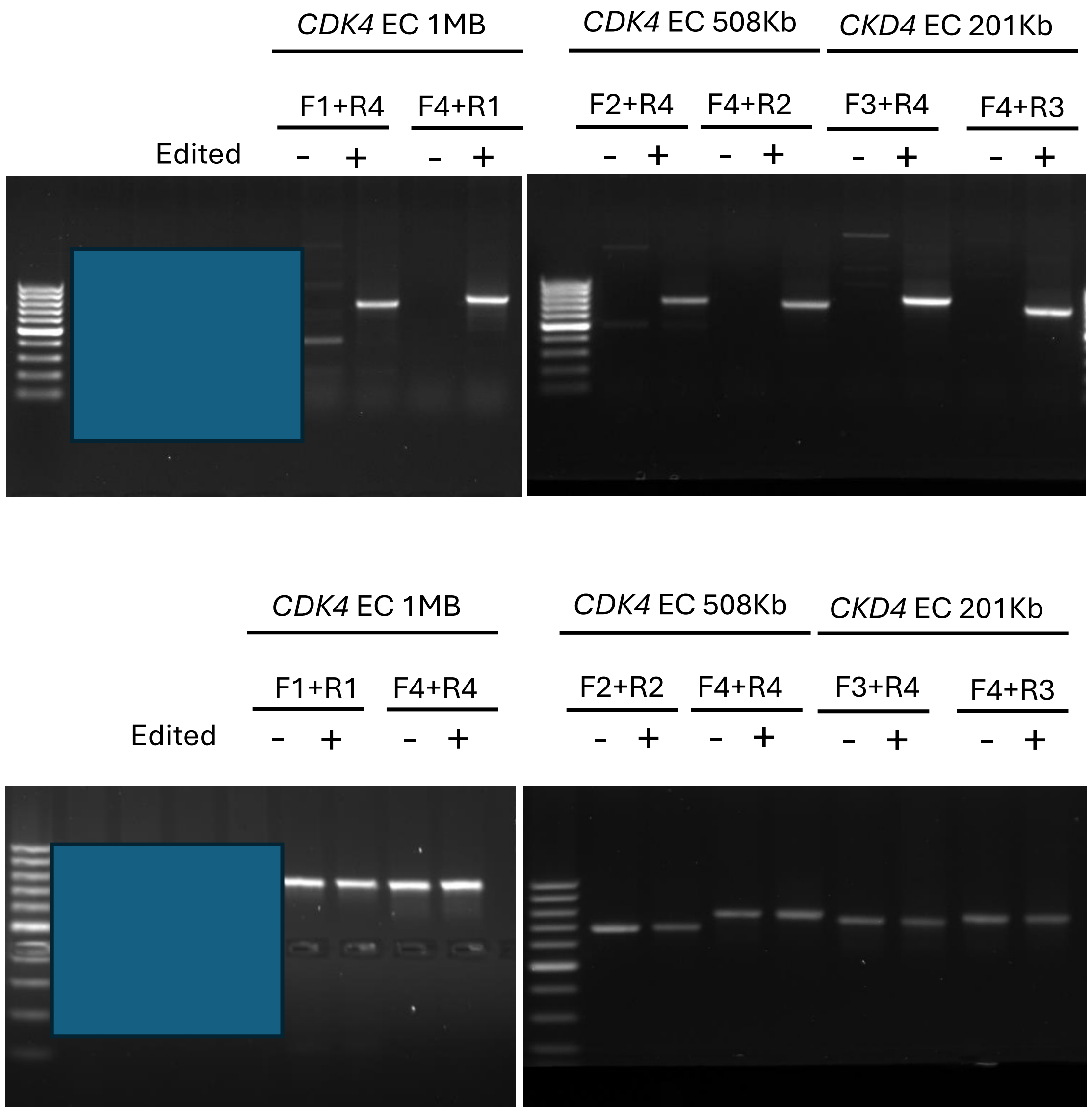


**Data not shown on Figure**

### Supplementary Figure S30

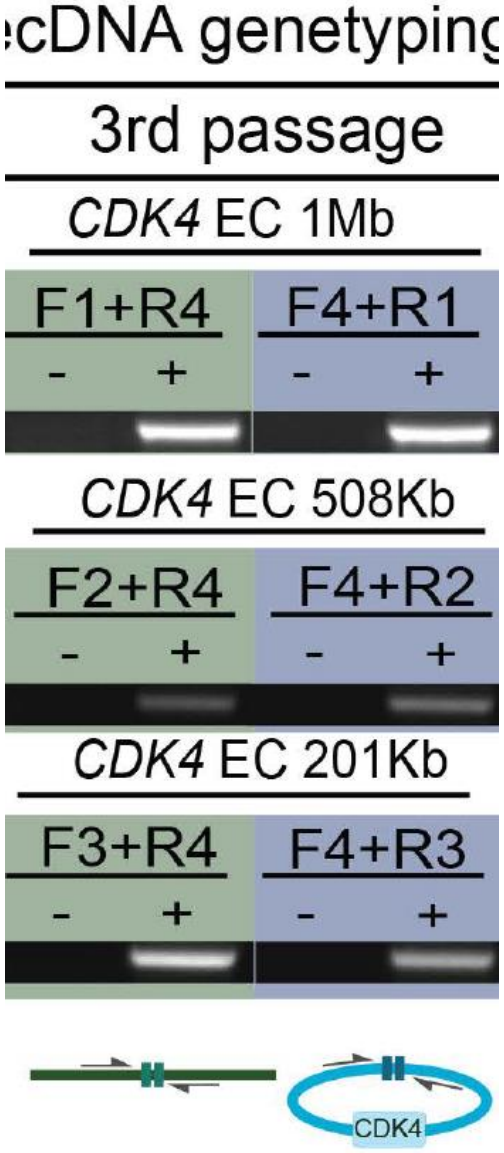


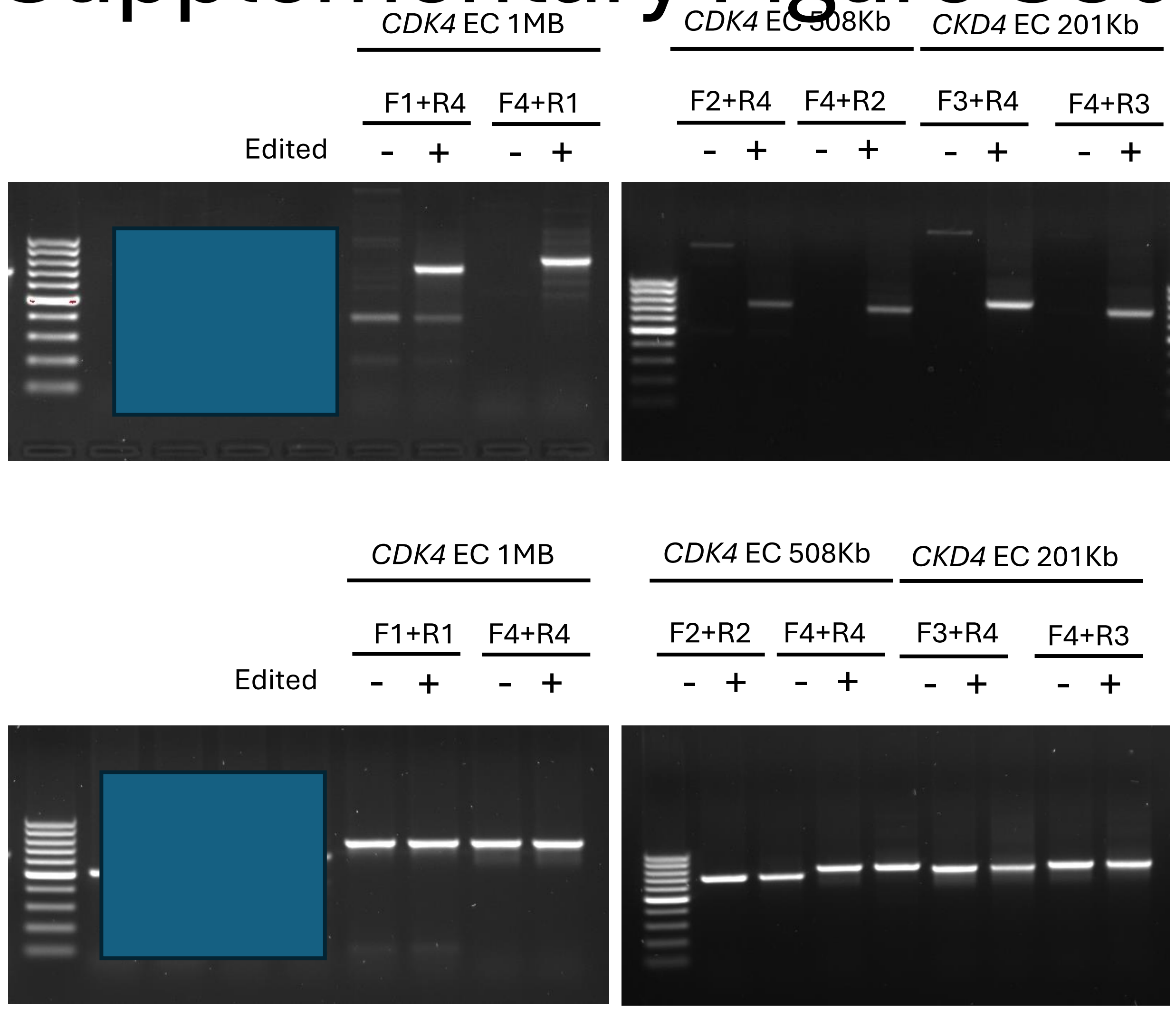


**Data not shown on Figure**

### Supplementary Figure S30

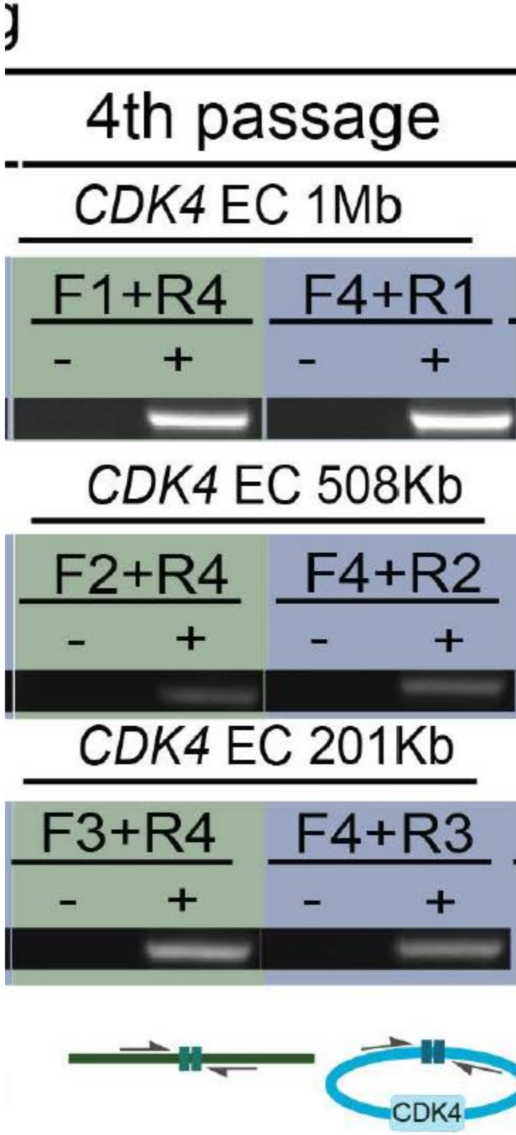


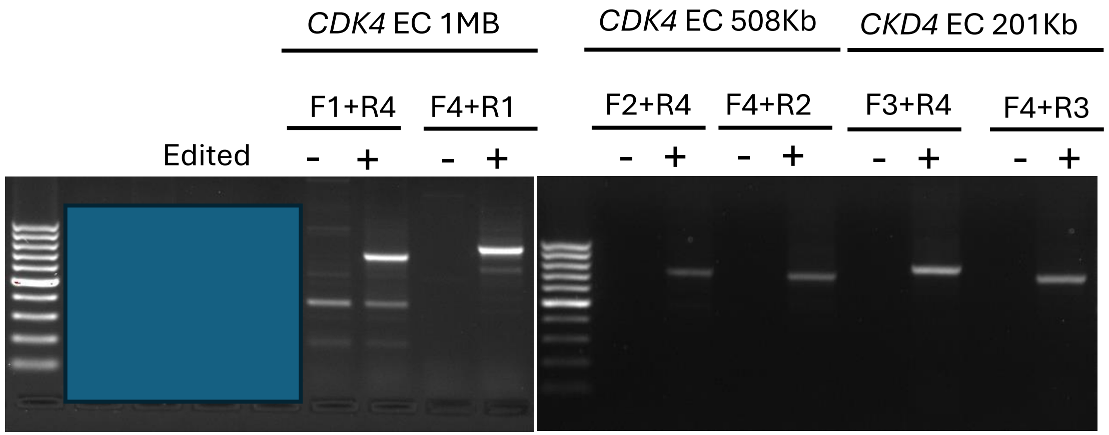


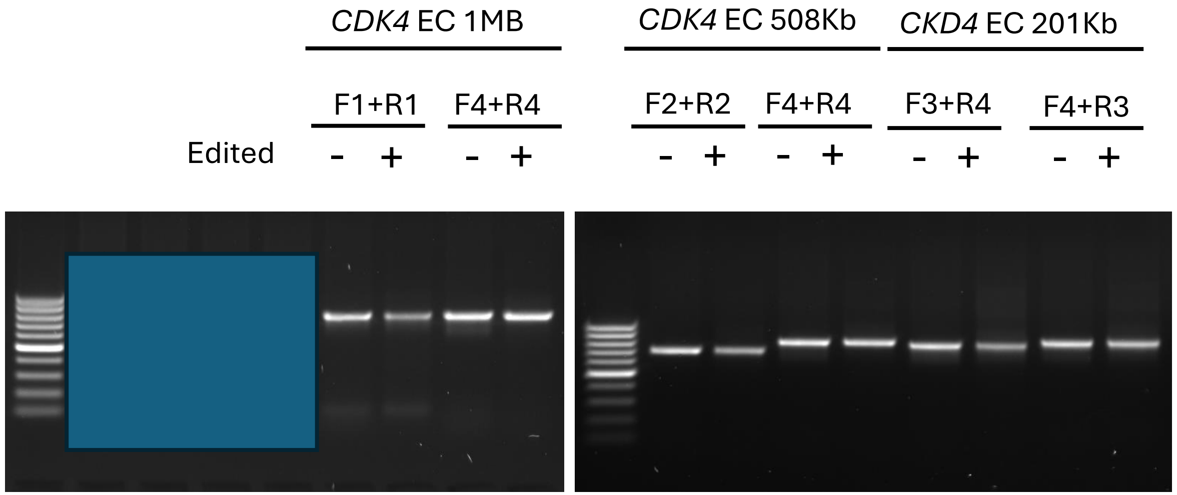


**Data not shown on Figure**

### Supplementary Figure S30

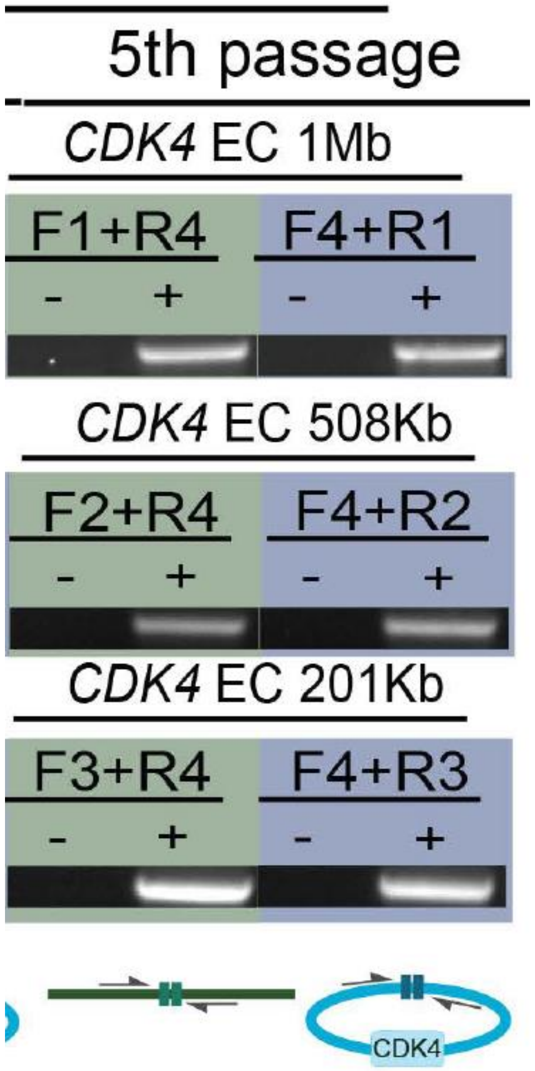


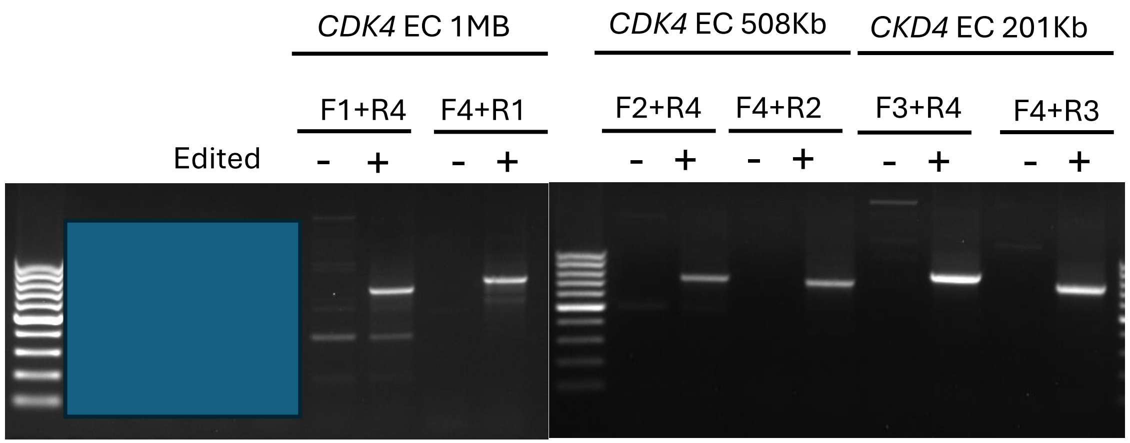


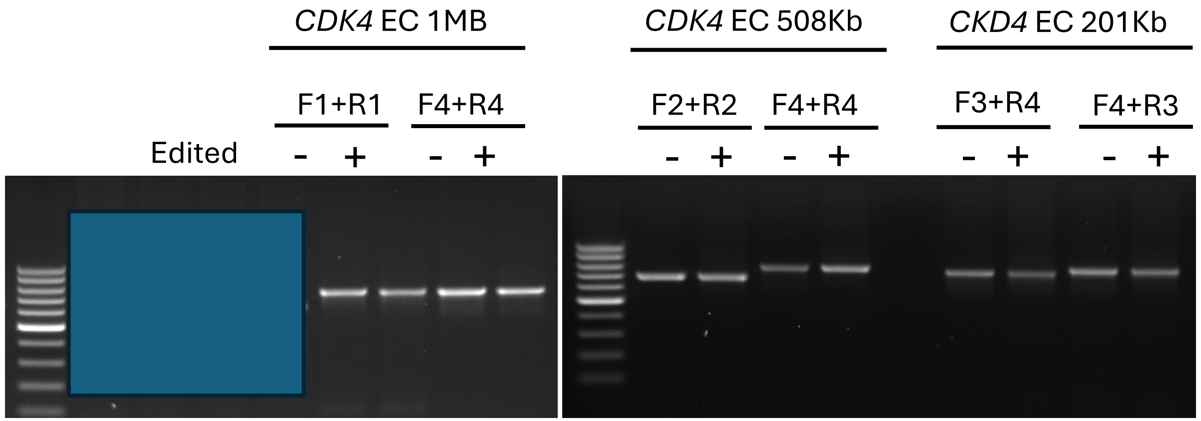


**Data not shown on Figure**

### Supplementary Figure S31

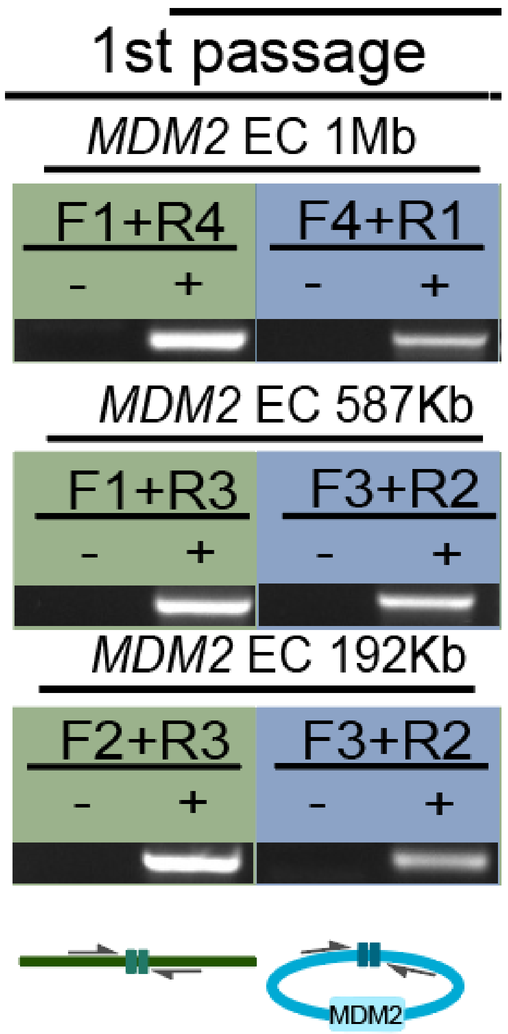


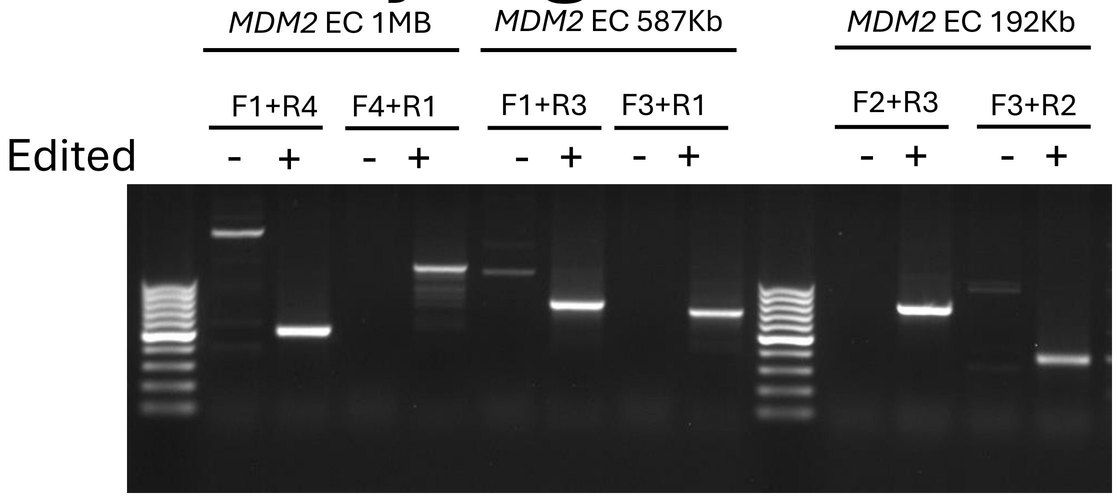


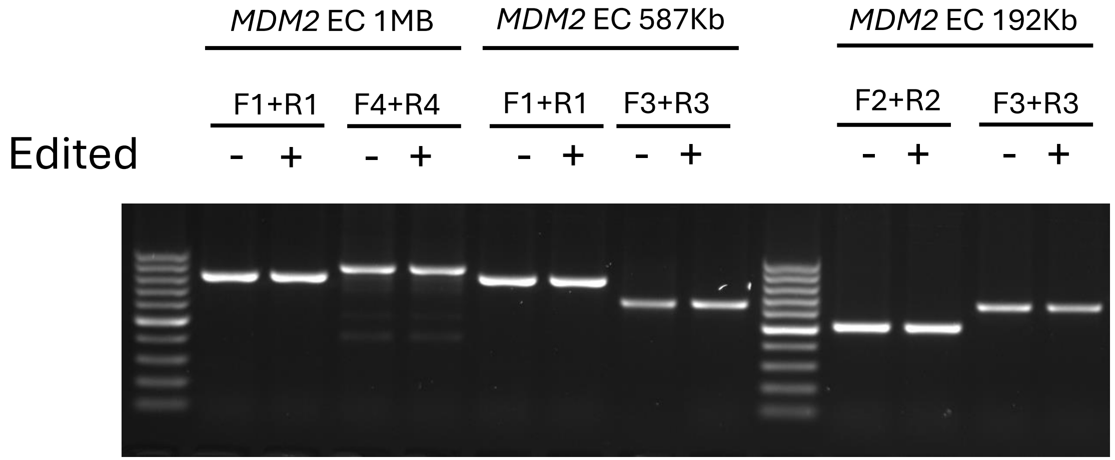


**Data not shown on Figure**

### Supplementary Figure S31

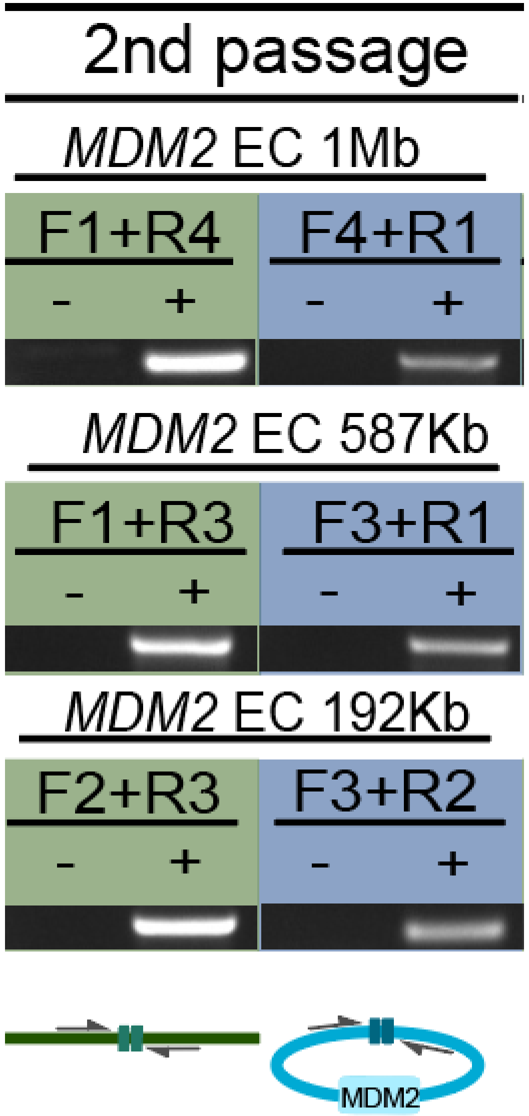


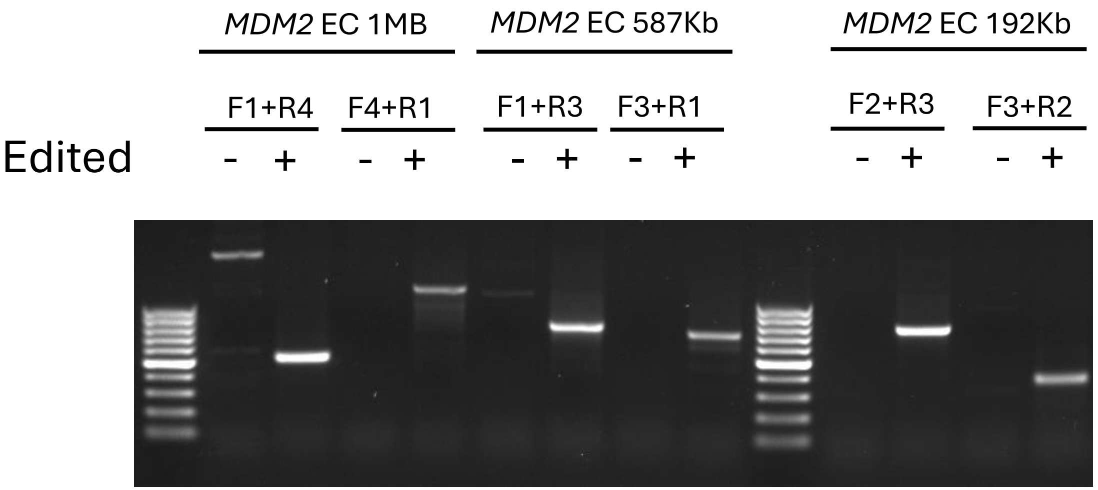


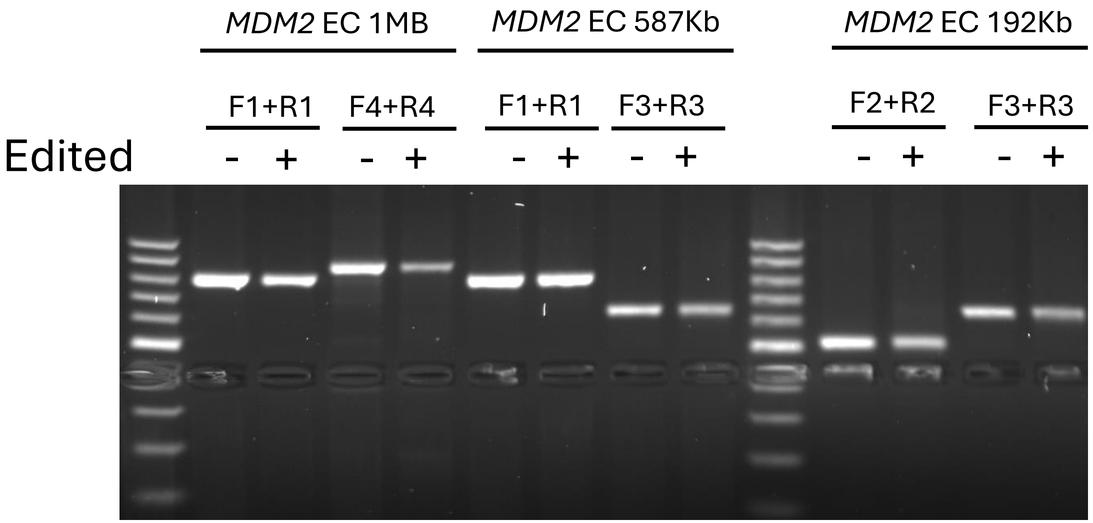


**Data not shown on Figure**

### Supplementary Figure S31

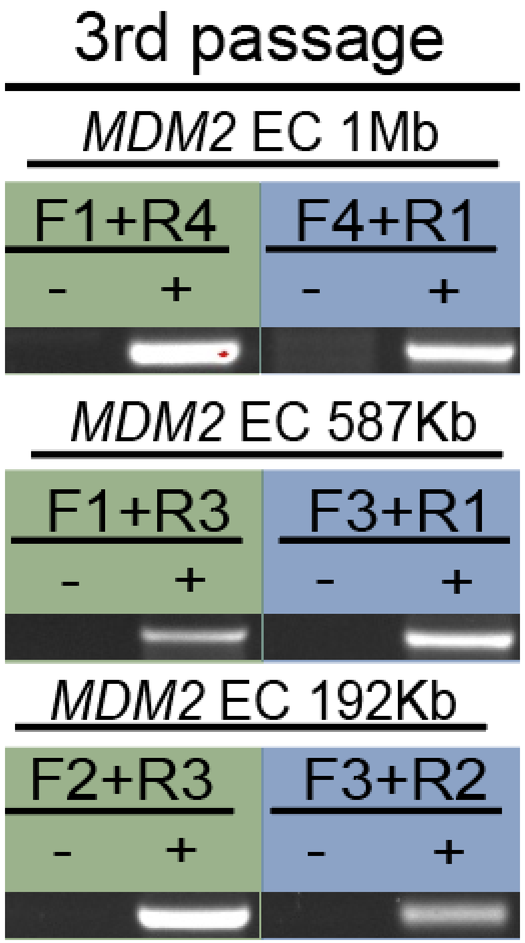


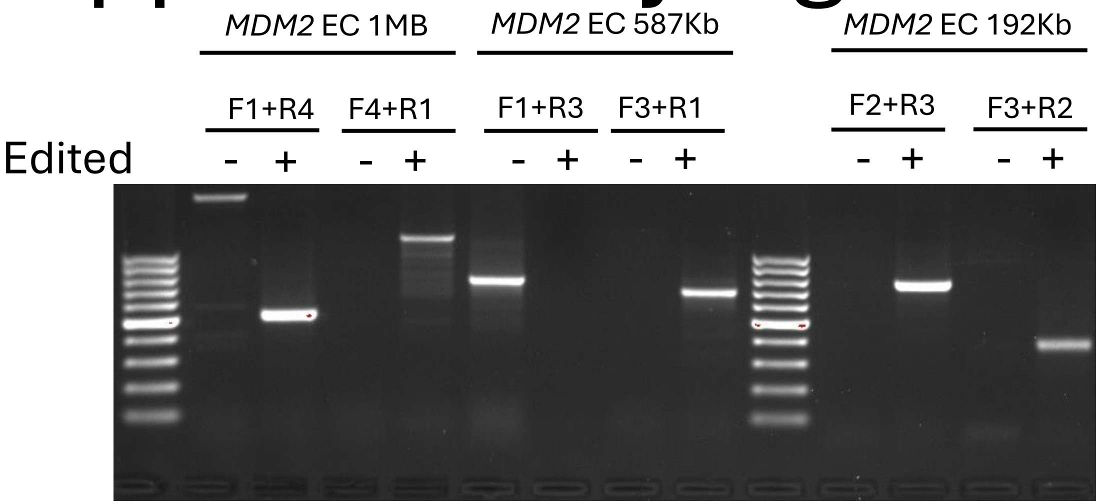


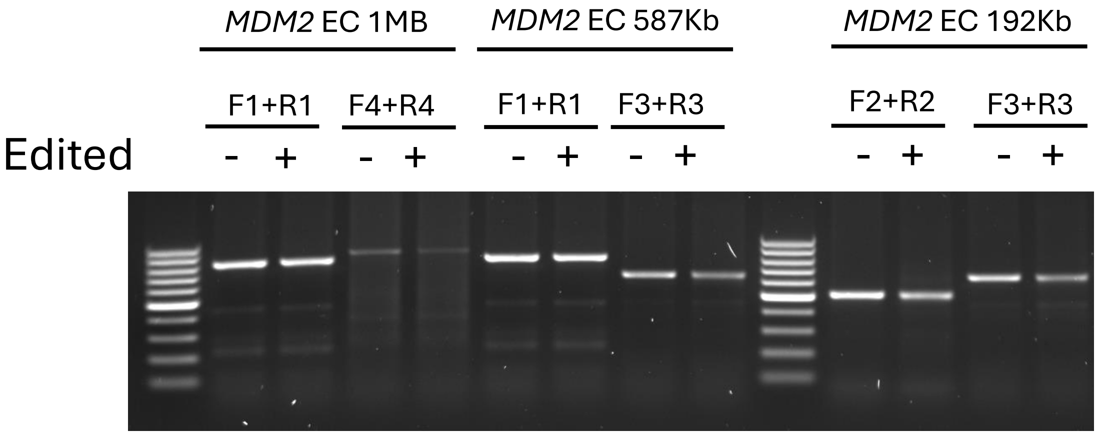


**Data not shown on Figure**

### Supplementary Figure S31

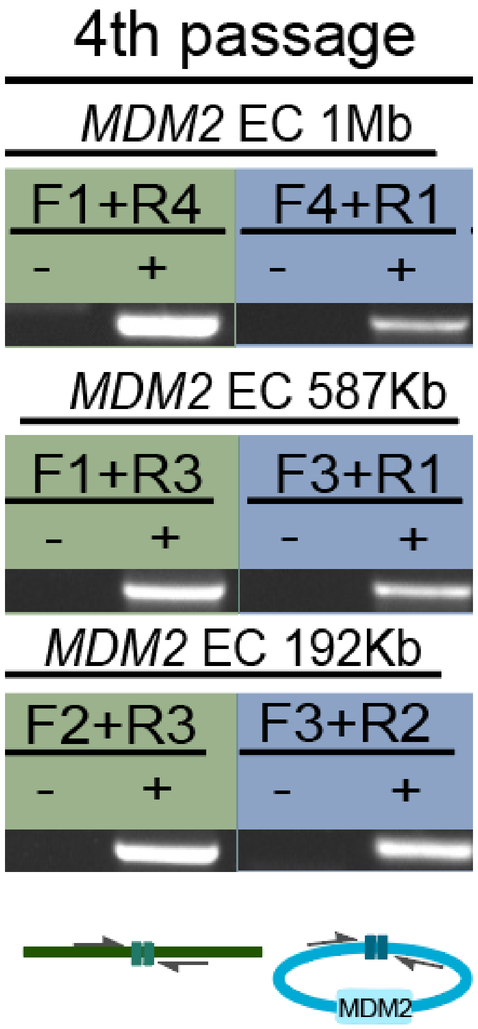


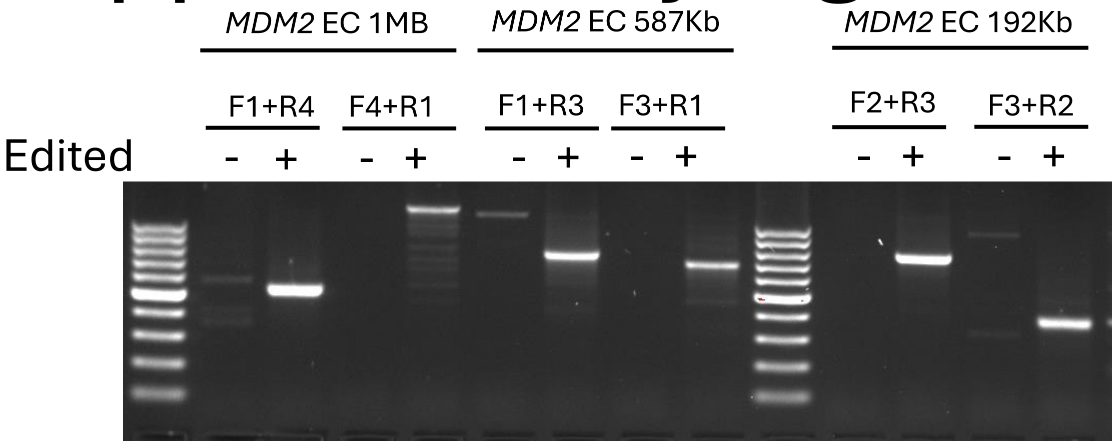


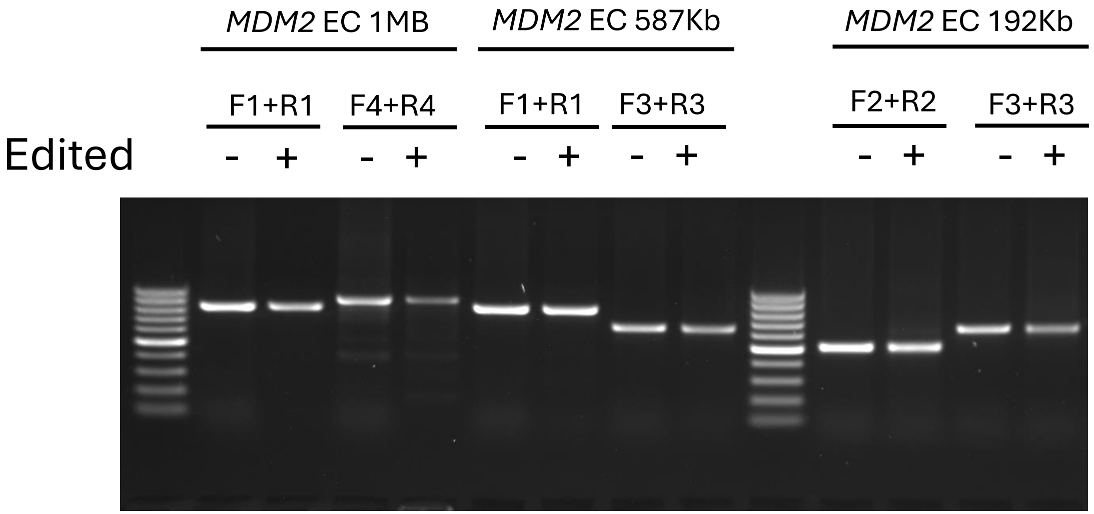


**Data not shown on Figure**

### Supplementary Figure S31

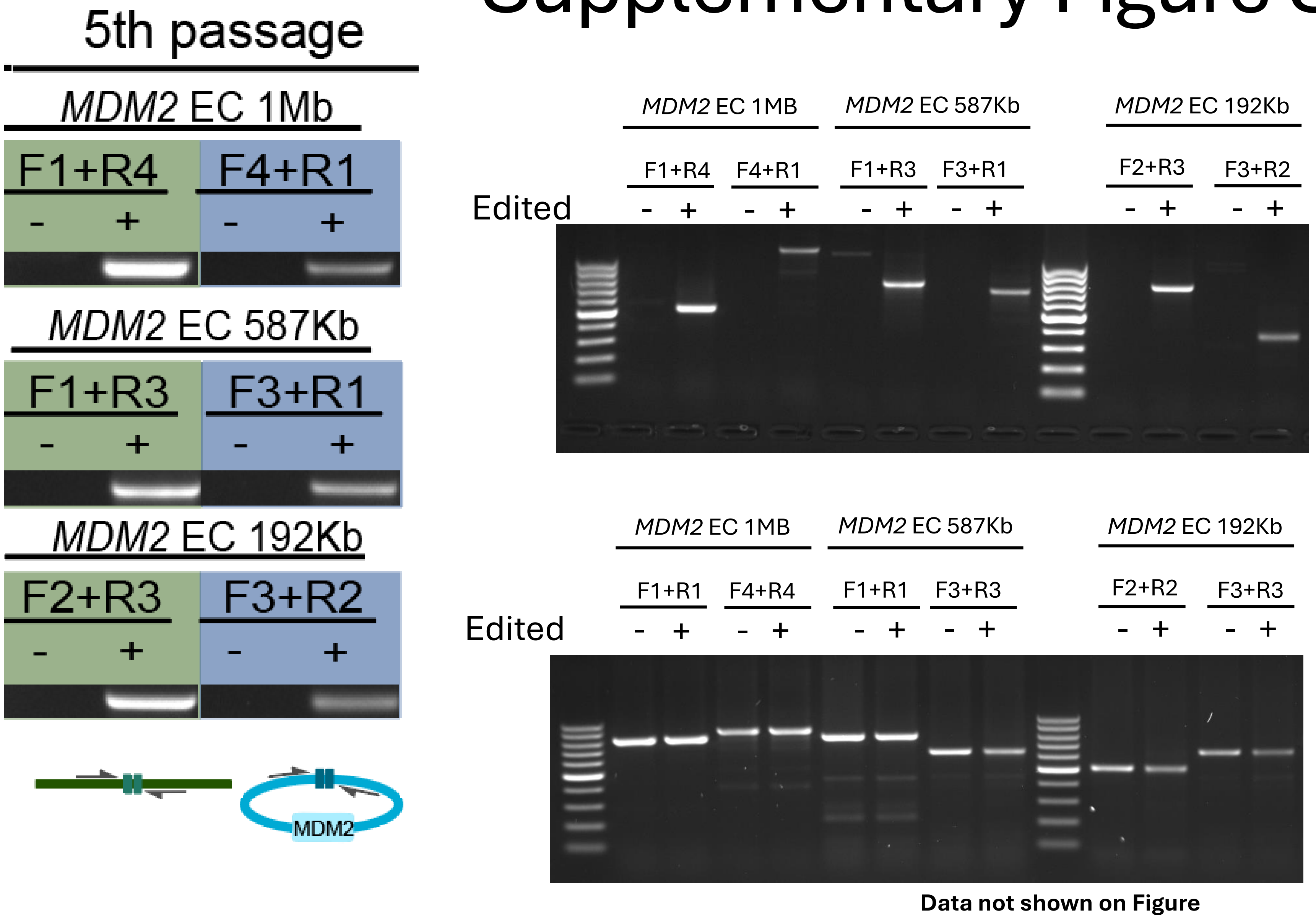

### Supplementary Figure S32

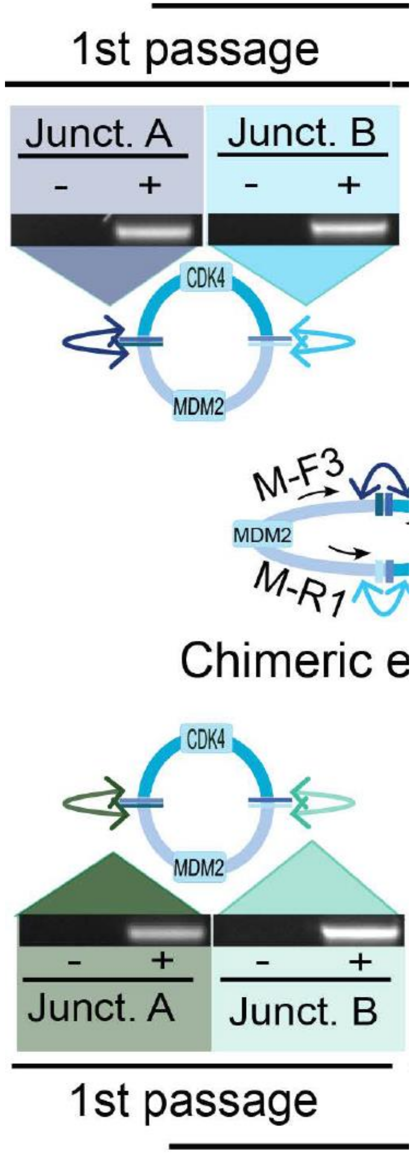


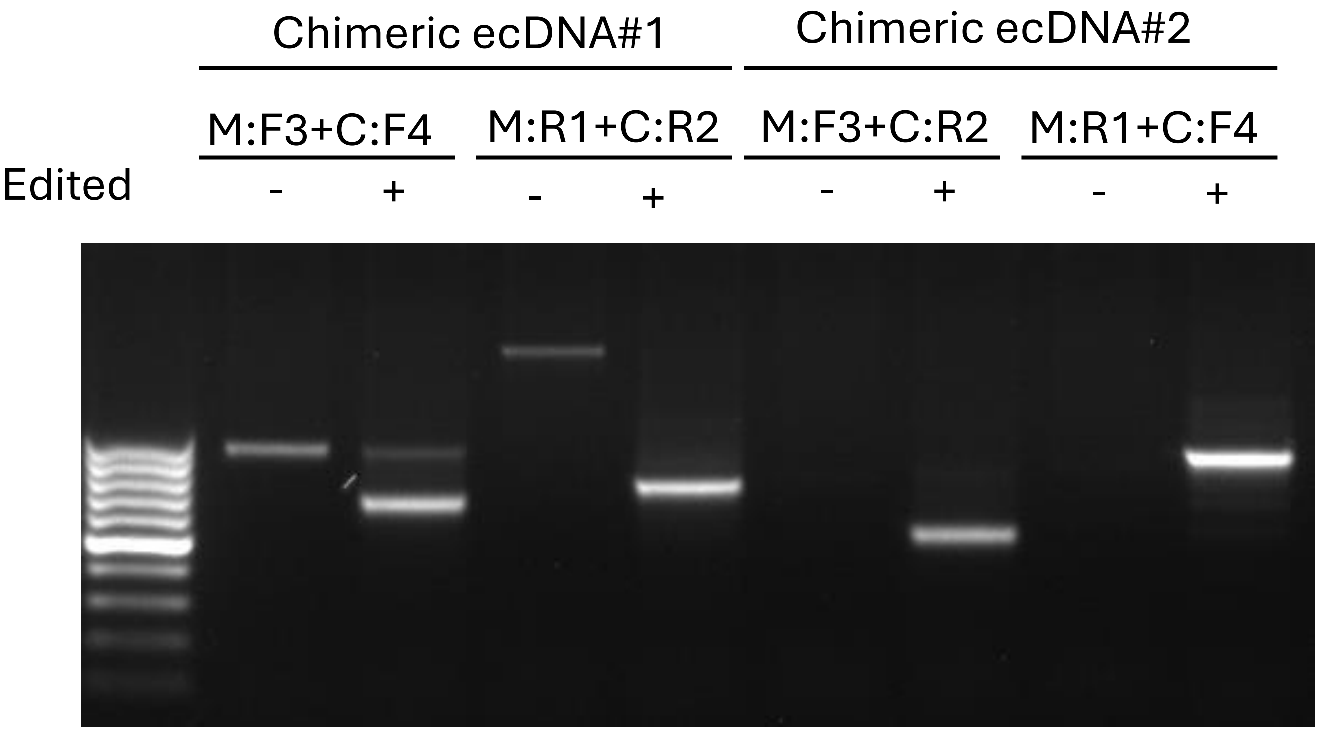

### Supplementary Figure S32

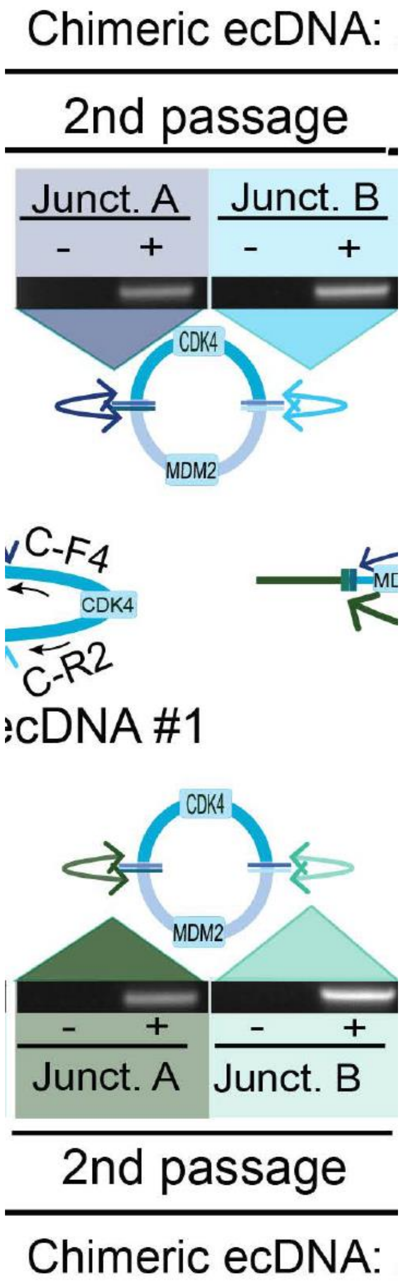


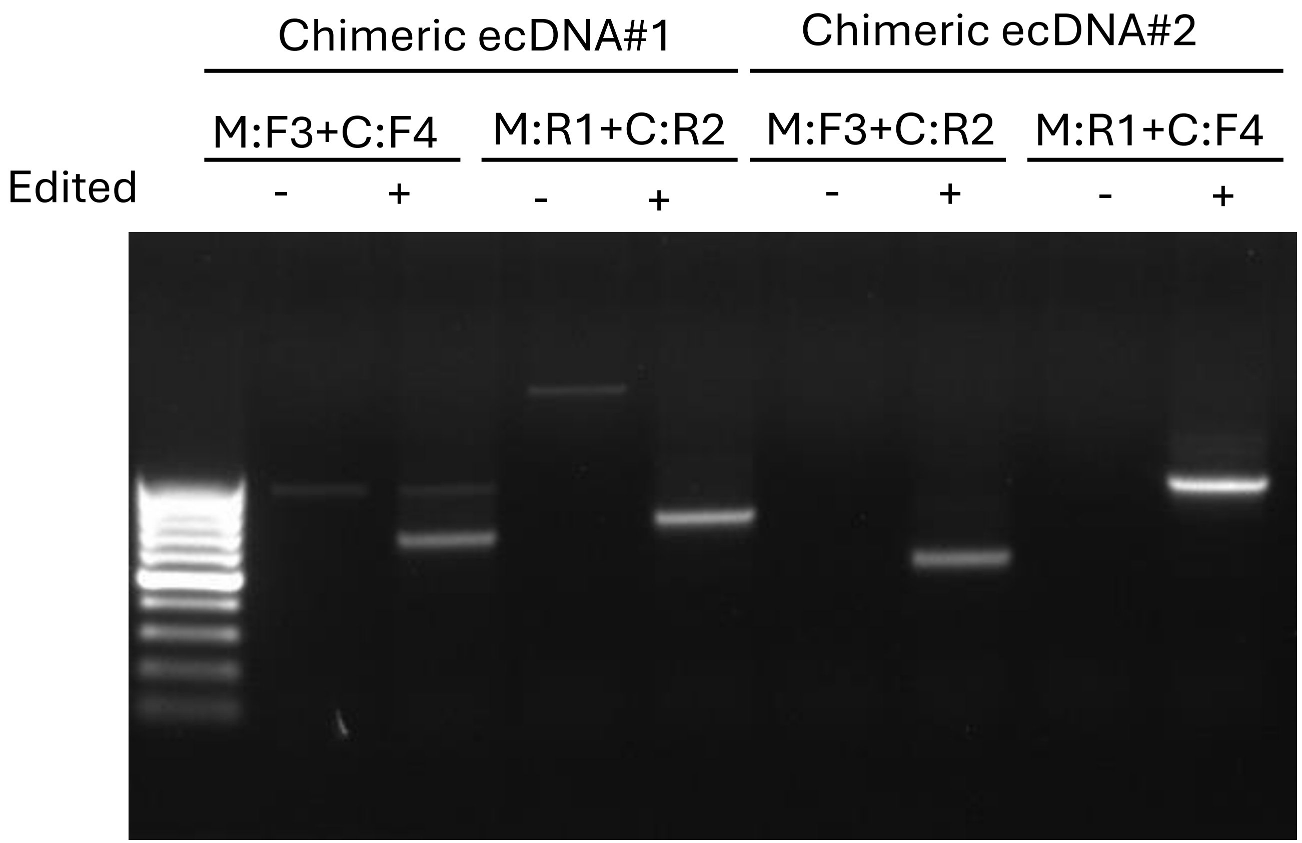

### Supplementary Figure S32

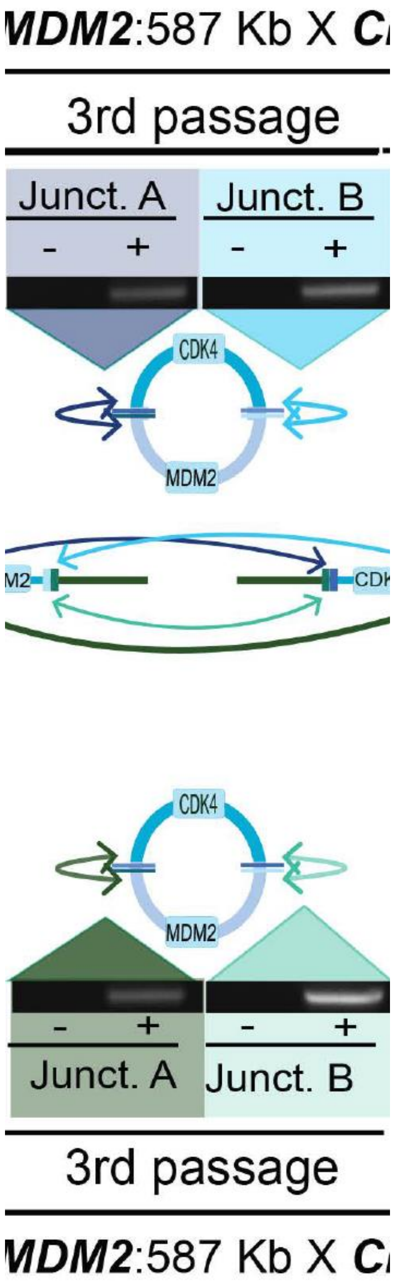


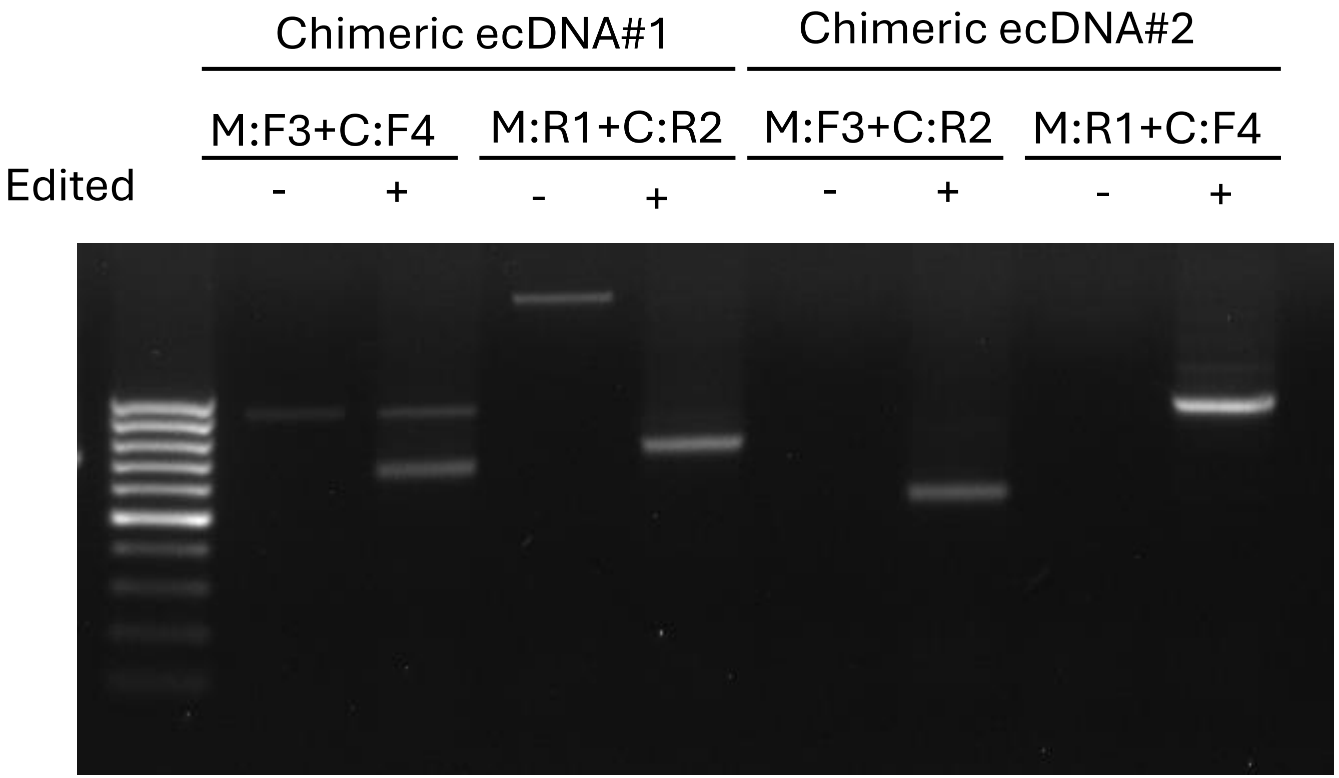

### Supplementary Figure S32

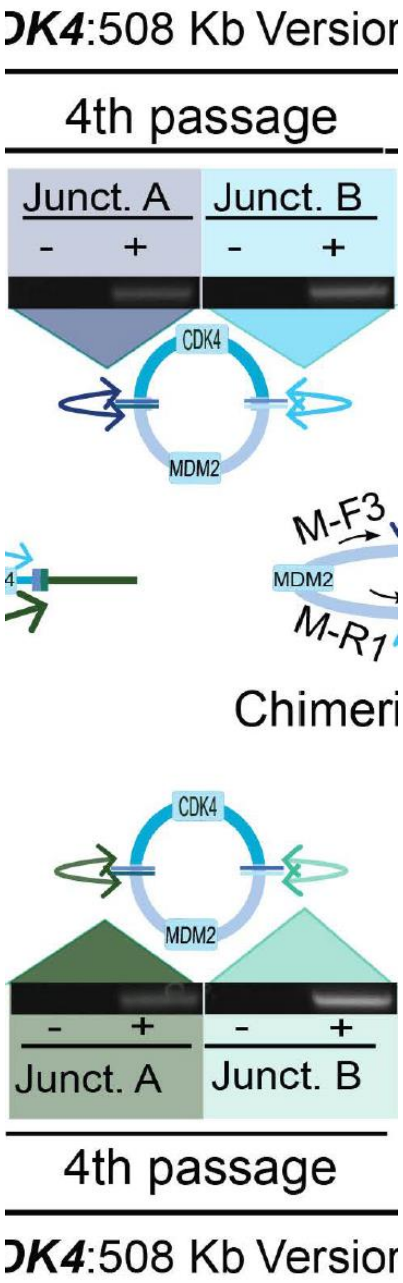


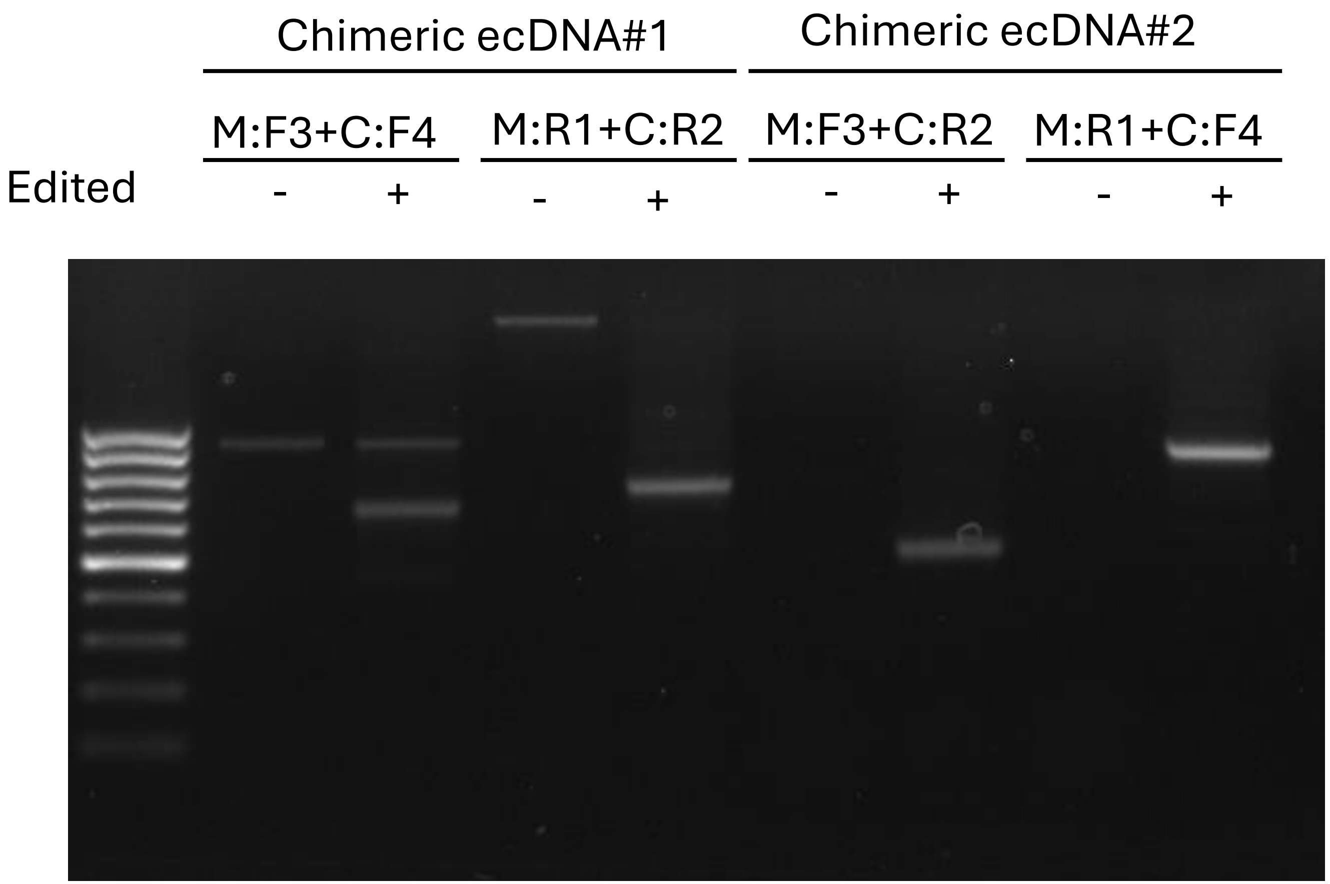

### Supplementary Figure S32

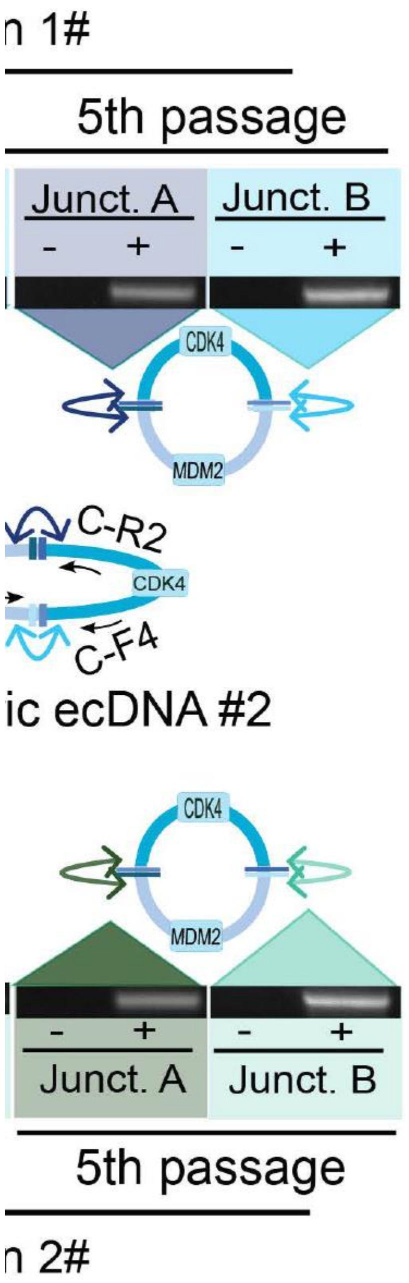


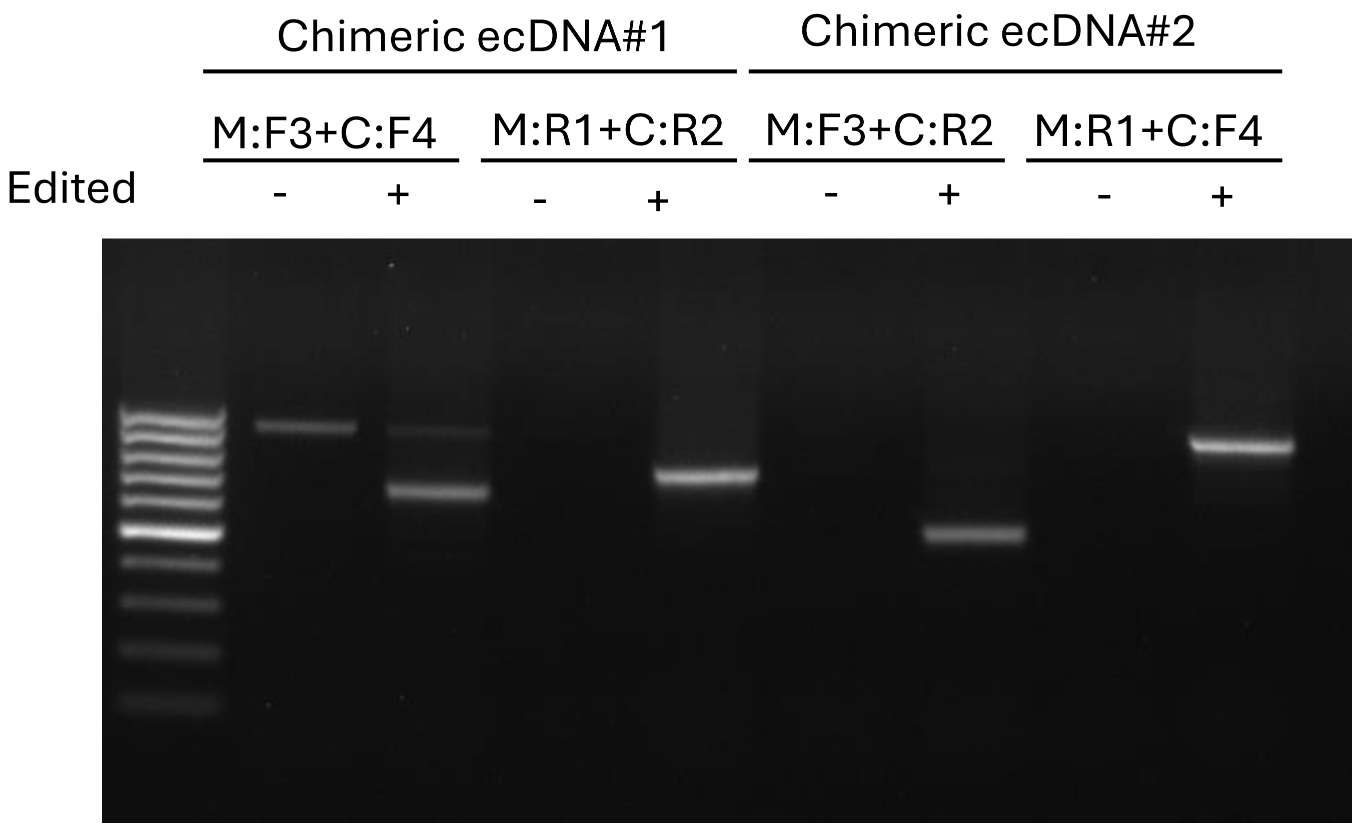
